# A global atlas of subsurface microbiomes reveals phylogenetic novelty, large scale biodiversity gradients, and a marine-terrestrial divide

**DOI:** 10.1101/2024.04.29.591682

**Authors:** S. Emil Ruff, Isabella Hrabe de Angelis, Megan Mullis, Jérôme P. Payet, Cara Magnabosco, Karen G. Lloyd, Cody S. Sheik, Andrew D. Steen, Anna Shipunova, Aleksey Morozov, Brandi Kiel Reese, James A. Bradley, Clarisse Lemonnier, Matthew O. Schrenk, Samantha B Joye, Julie A. Huber, Alexander J. Probst, Hilary G. Morrison, Mitchell L. Sogin, Joshua Ladau, Frederick Colwell

## Abstract

Subsurface environments constitute one of Earth’s largest habitats for microbial life, yet differences between surface and subsurface microbiomes and between marine and terrestrial microbiomes remain unclear. We analyzed 478 archaeal and 1054 bacterial amplicon sequence datasets and 147 metagenomes from diverse and globally distributed surface and subsurface environments. Taxonomic analyses show that archaeal and bacterial diversity is similar in marine and terrestrial microbiomes at local to global scales. However, community composition greatly differs between marine and terrestrial microbiomes, suggesting a horizontal phylogenetic divide between sea and land that mirrors patterns in plant and animal diversity. In contrast, community composition overlapped from surface to subsurface environments indicating vertical diversity continua rather than a discrete subsurface biosphere. Diversity of terrestrial microbiomes decreases from surface to subsurface environments. Marine subsurface diversity and phylogenetic novelty rivaled (bacteria) or even exceeded (archaea) that of surface environments, suggesting that the marine subsurface holds a considerable and underestimated fraction of Earth’s archaeal and bacterial diversity.

**Teaser:** Subsurface ecosystems within Earth’s crust and seafloor harbor diverse and distinct microbiomes featuring many unknown lineages.

## Introduction

Microbial life is pervasive in Earth’s hugely varied habitats and environments. Microorganisms have adapted to grow in acidic and alkaline springs, salterns, deserts, environments with temperature extremes greater than 122 °C or lower than −20 °C, and from ambient pressures up to pressures greater than those of abyssal oceanic trenches (*1–4*). Although surface ecosystems are thought to harbor the majority of Earth’s total biomass (*5*), most of Earth’s bacterial and archaeal biomass carbon is stored in subsurface ecosystems from less than one meter to many kilometers beneath the Earth’s surface. The local irregularity of boundaries between surface and subsurface ecosystems (*6*) due to the variability of physicochemical and biological parameters with depth, support only a loose definition of subsurface environments – which include soils, rocks, or sediments deeper than 1 to 8 m below Earth’s land surface or deeper than 0.1 to 1 m below the seafloor (mbsf) (*7–11*). In this study we use a stringent definition including only deep aquifers, rock fracture fluids, deep sediment, and rock cores.

At continental margins, especially at the mouths of large rivers, sediment thickness can exceed 10 km (*12*). Similarly, the habitable continental crust can exceed 15 km thickness in the Canadian, Fennoscandian and Siberian shields (*7*). Archaea, bacteria, and eukaryotic microorganisms inhabit this subsurface biosphere. Marine subsurface ecosystems, in particular, have relatively high proportions and abundances of *Archaea* compared to most other ecosystems (*8*, *13–16*). Subsurface ecosystems may host more than half of all microbial cells (∼5-12 × 10^29^) on Earth (*7*, *17–20*), despite sometimes very low cell densities, for example, in subsurface sediments of the oligotrophic South Pacific Gyre as low as approximately 100 cells per cubic centimeter of sediment (*21*).

Continuous deposition of organic matter from the overlying ocean and concomitant burial explains the introduction and presence of most microbes in deep sediment layers beneath the ocean floor. Life in marine deep subsurface sediments often bears a resemblance to the shallow subsurface life at a given location (*22–25*). Terrestrial subsurface ecosystems exhibit a similar connection to the surface world, with entrainment of shallow subsurface communities into the deeper realms via recharge and movement of fluids (*26*). This similarity exists despite the considerable environmental differences between surface and subsurface environments (*i.e.*, differences in pressure, light, oxygen, energy and nutrient availabilities, and available pore space within which cells might exist) that generally select for distinct microbial communities. Subsurface microbes can disperse via marine and terrestrial aquifers (*27*, *28*), fracture fluids (*29*), and porewaters (*30*). Hydrodynamic and hydrogeological processes such as eruptions of mud volcanoes (*31*, *32*) or fluid seepage (*33*, *34*) can reintroduce microbes from subsurface to surface environments.

Intrinsic or acquired capacities to metabolize subsurface energy, carbon, and nutrient sources, and/or improved survival and dormancy strategies allow microbes to survive and live in subsurface environments. The predominant occurrence of certain microbial lineages in subsurface ecosystems suggests that some organisms are better equipped for the subsurface than others. Coping strategies for survival may play a role, yet evidence also suggests that with increasing depth, *e.g.* in marine sediments, microbes change their expressed extracellular peptidases to those that specialize in highly degraded detrital proteins (*35*). Microbes can also increase cellular lifespan by slowing genome transcription (*36*), or actively expressing mRNA for DNA repair enzymes (*37*). Among many other processes acetogenesis (*38*, *39*), methanogenesis (*40*), fermentation of microbial biomass and necromass (*41–43*), serpentinization (*44*) and even radiolysis (*45*, *46*) might contribute to the subsistence of deep life. Indeed, while a portion of cells persist in a dormant state (*47*), many organisms actively metabolize (*21*, *48*, *49*) but often with generation times of decades to centuries (*23*, *50–53*).

While subsurface ecosystems harbor substantial biomass, diversity, and phylogenetic novelty, the degree to which the biodiversity of subsurface ecosystems directly compares with that of surface ecosystems remains unclear. Moreover, although standardized global meta-analyses exist for the marine (*16*) and the terrestrial subsurface (*54*), a standardized global dataset encompassing both marine and terrestrial subsurface environments was not available to date. Here, we provide a comparison between surface and subsurface as well as between marine and terrestrial environments. We investigate environments that span a broad range of depths from surface environments (under the influence of relatively fresh photosynthesis-derived organic matter) to very deep and isolated environments (cut off from photosynthetic primary production for at least centuries). We analyze and compare 1532 globally distributed taxonomic marker gene datasets and 147 metagenomes to investigate (i) whether microbial communities of marine and terrestrial biomes and of surface and subsurface environments fundamentally differ, (ii) whether subsurface environments are generally less diverse than surface ecosystems, (iii) whether subsurface environments harbor distinct clades, and iv) whether marine and terrestrial subsurface communities share a core microbiome.

## Results and Discussion

### The Census of Deep Life atlas of subsurface biodiversity

The Census of Deep Life (CoDL) under the auspices of the Deep Carbon Observatory organized a decadal effort (2010–2020) to characterize microbial diversity and function in subsurface ecosystems worldwide, carried out by more than two dozen groups and hundreds of researchers. After the completion of the CoDL, we compiled, re-analyzed, and compared 1054 bacterial and 478 archaeal 16S rRNA gene amplicon datasets from 35 individual globally distributed CoDL projects together with 17 additional non-CoDL projects that originated mainly from surface ecosystems (Fig. 1A, B; Table S1, S2). Archaeal and bacterial 16S rRNA gene amplicon sequence variants (ASVs, quasi strain-level microbial lineages) were amplified using domain-specific primers. We obtained 31099 archaeal and 415423 bacterial ASVs from 4.9 × 10^7^ archaeal and 13 × 10^7^ bacterial reads. On average we found 155 archaeal and 1257 bacterial ASVs per sample (Table S3, S4). Analyses of 147 metagenomes from 49 globally distributed CoDL subsurface projects complement the amplicon datasets. The metagenomes were used for taxonomic analyses based on two different taxonomic markers, 16S rRNA gene sequences retrieved by phyloFlash (pF16S; before read assembly) and ribosomal protein S3 genes (rpS3; after read assembly), as well as for a community analysis based on all predicted genes. We obtained a total of 1443 different archaeal and 22556 different bacterial pF16S genes, as well as 150 different archaeal and 1575 different bacterial rpS3 genes (Table S3, S4).

**Fig. 1.**
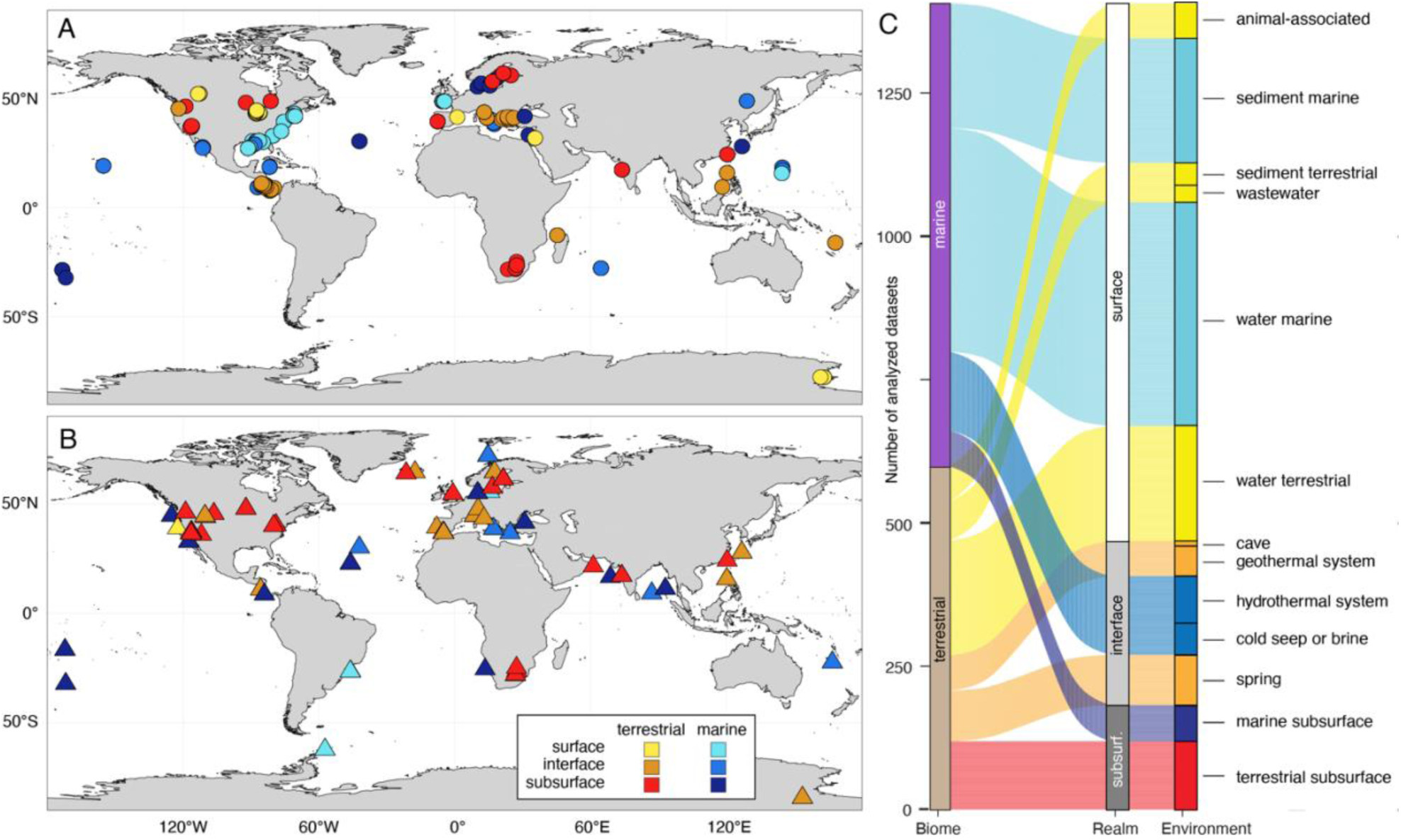
Geographic location and origin of samples. Maps of samples used for metabarcoding of 16S rRNA gene amplicon taxonomic marker genes (A) and shotgun metagenomic analyses of unassembled and de novo assembled taxonomic marker genes (B). Each symbol represents one project, which comprises multiple individual samples. Both terrestrial as well as marine samples may contain rock, sediment, or water samples. Further maps showing sample material, *p*H and temperature are included in the Supplementary Material (Fig. S1). (C) Overview of sample origins derived from marine and terrestrial biomes, depth realms, and environments.

We analyzed the communities first by grouping the amplicon datasets based on the biome from which the samples originated; marine biome (n_Archaea_=304 datasets, n_Bacteria_=505 datasets) and terrestrial biome (n_Archaea_=174, n_Bacteria_=549). Within those biomes we grouped samples based on the depth realm they originated from: surface, interface and subsurface. Surface datasets (n_Archaea_=183, n_Bacteria_=687) include water samples from oceans and lakes (at various depths in the water column), shallow (<0.1 mbsf) sediment samples from oceans, estuaries and lakes, animal-associated microbiomes, and wastewater. Surface ecosystems represent environments under the influence of relatively fresh photosynthesis-derived organic matter. Subsurface datasets (n_Archaea_=85, n_Bacteria_=122) originated from ecosystems that have likely been cut-off from photosynthetic primary production for a timescale of at least centuries. These ecosystems were accessed via boreholes and include deep sediments, aquifers, and fracture fluids. Marine subsurface sediments were further sub-divided based on depositional setting: shelf, slope, and abyssal domains. The location of each domain is defined by water depth (*55–57*). Shelf environments roughly correspond to water depths <200 m, except the Antarctic region where shelf area corresponds to water depths <500 m. The abyssal plain corresponds to areas of water depths >3500m. Sediments under other water depths are referred to as slopes. Interface datasets (n_Archaea_=210, n_Bacteria_=245) originated from environments that are subjected to influences from surface and subsurface environments and processes. Interface environments can be surface sites that are driven by energy from the subsurface, or vice versa, and included water and sediment samples from caves, hot springs, hydrothermal vents, and cold seeps. We also grouped and analyzed the samples based on the 13 major environments they originated from, as listed in Fig. 1C. To minimize batch effects and biases, we sequenced and analyzed all samples using the same primers, the same chemistry, the same Illumina instrument, and the same bioinformatic pipeline. We minimized the potential impact of contamination by including blanks and controls, and removing notorious contaminants (*58*, *59*). We minimized composition bias by normalizing template concentration and standardizing workflows (*60–62*).

### Substantial differences between marine and terrestrial microbiomes

We found substantial taxonomic differences in archaeal and bacterial community structure between marine and terrestrial biomes. For this analysis we grouped all datasets solely based on whether they originated from a marine or terrestrial environment, regardless of their association with a surface, interface, or subsurface environment. Alpha diversity (community diversity per locality/sample) and beta diversity (community diversity between localities/samples) of marine and terrestrial microbiomes showed the same trends across ASV, rpS3 and pF16S-based analyses. Richness, estimated richness (Chao1) and evenness (Shannon entropy, Inverse Simpson diversity) were similar for archaea but significantly higher for bacteria in marine relative to terrestrial microbiomes (Fig. 2A-F, Fig. S2). After subsampling the datasets to account for the unequal sampling effort, *i.e.*, different sequencing depth, we found on average 75 and 79 archaeal and 513 and 419 bacterial ASV in marine and terrestrial samples, respectively (Table S5, S6). Microbial community structure and composition was very different between the marine and terrestrial biomes. On average only 2-6 % of ASVs, 2 % of rpS3 genes and 8-9 % of pF16S genes were shared between any marine and terrestrial dataset (Fig. 2G, H; Fig. S3; S4). Gamma diversity (total community diversity per biome/environment) tended to be higher in the terrestrial biome (Fig. S3), with exception of bacterial ASVs which were higher in the marine biome. Analyses of all gene families found in the metagenomes, support a higher functional richness and evenness in the terrestrial microbiome (Fig S5A, B), yet the dissimilarity in functional genes between biomes was less pronounced, likely due to housekeeping genes that are found ubiquitously in most microorganisms (Fig. S5C, D).

**Fig. 2.**
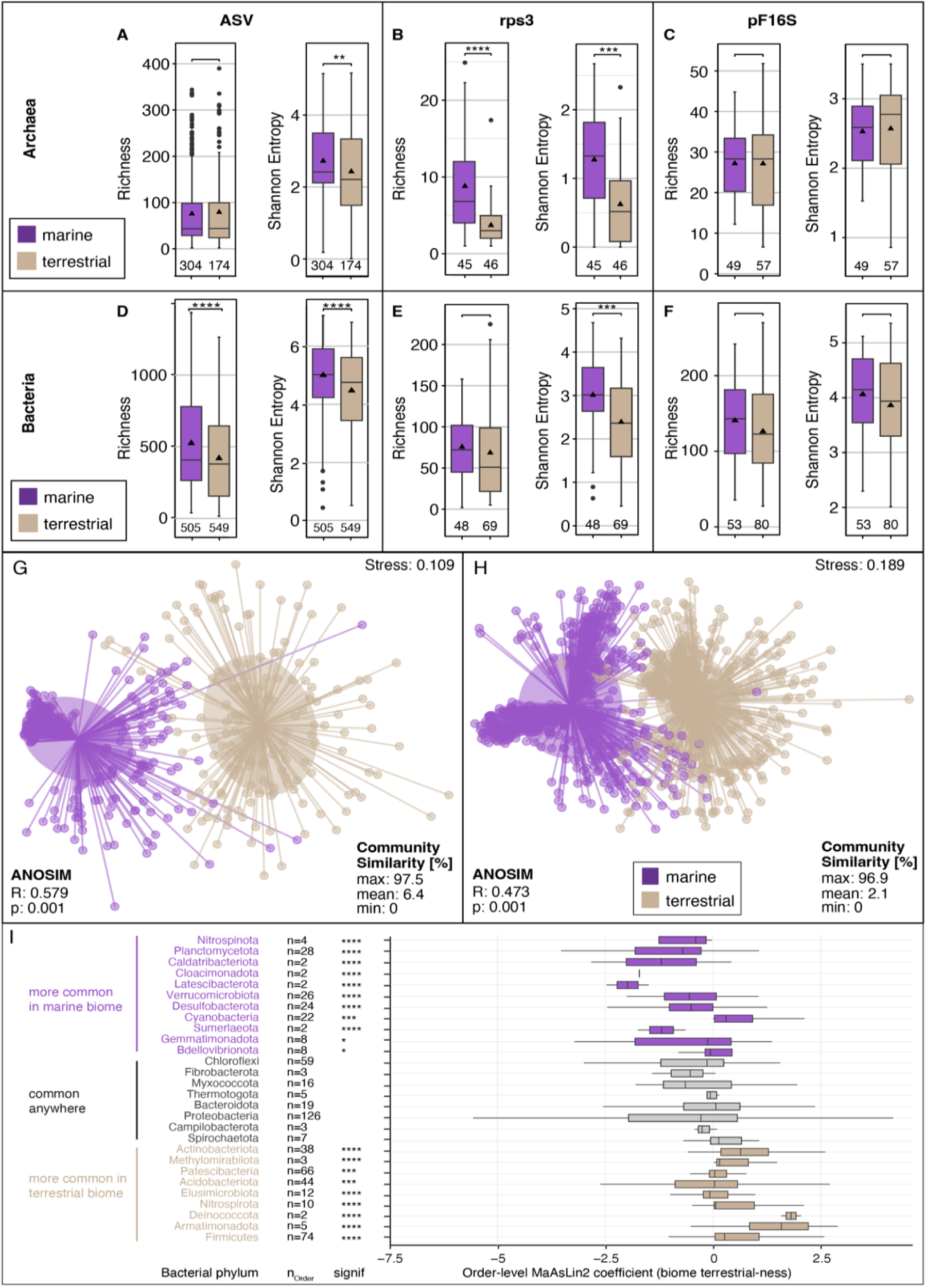
Microbial diversity in marine and terrestrial biome. Archaeal (A-C) and bacterial (D-F) alpha diversity (per sample community richness and evenness) in marine and terrestrial biomes using 16S rRNA gene amplicon sequence variants (ASVs; A, D), as well as metagenome-derived ribosomal protein S3 genes (rpS3; B, E) and 16S rRNA gene sequences detected by phyloFlash (pF16S). To allow comparison the datasets were subsampled to the same number of reads. Pairwise comparisons were performed using a Wilcoxon rank sum test. Significance: ***: p<0.001. The number of datasets is shown below the boxplots. Community dissimilarity between marine and terrestrial communities was shown by non-metric multidimensional scaling ordinations of ASV-based dissimilarity matrices using 469 archaeal (G) and 1105 bacterial datasets (H). Each circle represents the community structure of a dataset and is connected to the group centroid (weighted average mean of within-group distances), the ellipses depict one standard deviation of the centroid. Statistical testing using ANOSIM showed that the groups are overlapping but significantly different (R∼0.5, p<0.001). (I) Differential abundance analyses of marine vs terrestrial bacterial phyla. The phyla are ordered from top to bottom based on increasing phylum level MaAsLin2 coefficient, *i.e.* likeliness of their occurrence in terrestrial-derived samples (“terrestrial-ness”). Boxplots summarize order level MaAsLin2 coefficients, i. e., “terrestrial-ness”, within the listed phyla. Note: due to ease of visualization boxplots are also shown for very small number (n) of orders. Significance levels are: *: p<0.05, **: p<0.01, ***: p<0.001, ****: p<0.0001. Additional phyla are shown in Fig. S6.

A multivariate association analysis (*63*) of our dataset suggests that most bacterial phyla are significantly more abundant and prevalent in the marine biome (Fig. 2I, S6A). Phyla that are more common in marine environments include well-known *Cyanobacteria*, *Planctomycetota, Verrucomicrobiota,* and *Desulfobacterota*, but also less prominent phyla such as *Marinimicrobia, Nitrospinota, Fermentibacterota, Caldatribacteriota, Latescibacterota, Aerophobota* and *Gemmatimonadota.* In terrestrial environments we found more commonly *Firmicutes*, *Acidobacteriota*, *Actinobacteriota, Nitrospirota,* and *Patescibacteria*, as well as *Methylomirabilota, Armatimonadota and Elusimicrobiota* among others. A third group of bacterial phyla occupy both biomes at similar relative sequence abundances and prevalences. These included *Proteobacteria*, *Bacteroidota, Chloroflexi*, *Spirochaetota*, *Myxococcota, Campilobacterota* and *Thermotogota*. In contrast, most archaeal phyla were more abundant and prevalent in the terrestrial biome (Fig. S6B). Only *Thermoplasmatota* were significantly more common in marine environments, and *Crenarchaeota* and *Asgardarchaeota* were found in both biomes at similar abundances and prevalence. All other archaeal phyla including *Euryarchaeota, Halobacterota, Hadarchaeota, Nanoarchaeota, Aenigmarchaeota*, and *Altiarchaeota* were significantly more abundant and prevalent in the terrestrial biome. Relative sequence abundances of archaeal and bacterial phyla (Fig. S6C, D) support the trend revealed by the association analysis. Interestingly, some phyla contribute disproportionally to ASV-level diversity including *Nitrososphaeria, Bathyarchaeia, Chloroflexi and Planctomycetota* (Fig. S6E, F), being responsible for a larger part of the ASV-level diversity than their relative abundance suggests.

### Subsurface microbiomes can be as diverse as surface microbiomes

Within each biome, differences in alpha diversity between surface, interface, and subsurface environments, *i.e.*, depth realms, were less pronounced than between the biomes (Fig. S7, S8). Across depth realms, communities of both biomes – marine and terrestrial – mainly had a similar richness and shared many ASVs (Fig. S9, S10, Video S1, S2). Archaeal and bacterial communities in the three depth realms were distinct but with considerable overlaps. The larger differences in community structure between marine and terrestrial biomes than between depth realms, indicates that the divide between microbial life on land and in the sea is more pronounced than the divide between surface and subsurface communities. We found that species richness and diversity in many subsurface environments rival those in surface environments. This finding was consistent across the 13 investigated environment types – where levels of microbial diversity are often comparable across the surface, interface, and subsurface (Fig. 3A-D). Archaeal alpha diversity was overall highest in samples from cold seeps and brines, caves, springs, and the deep marine subsurface (Fig. 3A). Bacterial alpha diversity was very high in the cave samples and sediments, but also in seeps and vents. Marine subsurface bacterial diversity rivaled the diversity found in marine surface water, wastewater, or host-associated microbiomes (Fig. 3B). Total archaeal diversity (gamma diversity) was significantly higher in marine interface and subsurface environments than in the marine surface (Fig. 3C, S7A-D). Similarly, bacterial gamma diversity was highest in marine interface environments (Fig. 3D). In the terrestrial biome, archaeal gamma diversity was comparable across all depth realms (Fig. 3C), whereas bacterial diversity was highest in the surface (Fig. 3D).

**Fig. 3.**
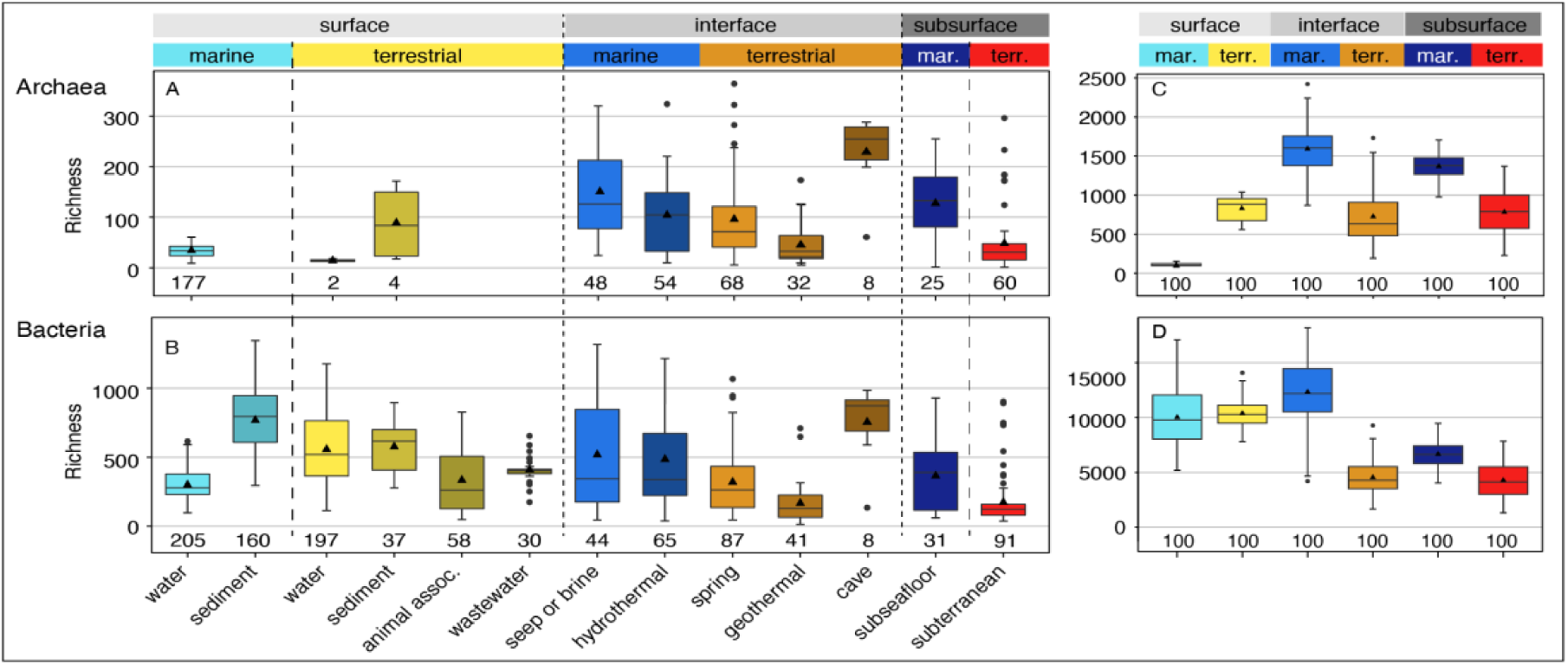
Observed archaeal and bacterial richness across environments based on 16S rRNA gene amplicon sequence variants. Archaeal (A) and bacterial (B) richness found in the 14 studied environments (n: number of included samples). Environments are grouped based on biome (marine, terrestrial) and depth realm (surface, interface, subsurface). Note: Archaeal terrestrial water and sediment samples contain too few datapoints for robust visualization using boxplots, yet the plots were retained for completeness. Total archaeal (C) and bacterial richness (D) using a subsampling approach to account for different group sizes (100 iterations using the smallest group size). Normalized gamma diversity corroborates that archaeal diversity is highest in marine interface and subsurface ecosystems even when group sizes are considered. All pairwise comparisons were significantly different except for gamma diversity between terrestrial interface and subsurface biomes (archaea and bacteria), and marine and terrestrial surface biomes (bacteria).

Interestingly, *Thermoplasmata*, *Nitrososphaeria*, and *Bathyarchaeia* together comprised more than half of the relative sequence abundance in any investigated environment, except the terrestrial subsurface (Fig. 4A; S11). The contribution of these three classes to global ASV richness was similarly outsized (Fig. 4B), while other clades that can occur at relatively high abundances contributed very little to overall richness, including *Hadarchaea*, ANME-1, *Methanobacteria*, *Thermoprotei* and *Methanococci* (Fig. 4A, B; S11). This suggests that a large part - if not most - of the global archaeal abundance and richness can be attributed to three archaeal classes *Thermoplasmata*, *Nitrososphaeria*, and *Bathyarchaeia*. Bacterial communities were largely dominated by *Proteobacteria* and *Bacteroidota*, except those of caves and the subseafloor where *Chloroflexi* played an important role (Fig. 4C). The contribution of the lineages to overall bacterial richness was often not proportional to their abundance. *Proteobacteria, Bacteroidota, Firmicutes, Cyanobacteria* and *Desulfobacterota* contributed less, while *Chloroflexi, Planctomycetota*, and *Verrucomicrobia* contributed more richness than expected based on their abundance (Fig. 4D). We found increasing archaeal richness and evenness with depth, supporting that archaea are well suited for life in subsurface and subsurface influenced, *i.e.* interface, environments. The importance of archaea in marine subsurface ecosystems has previously been demonstrated by studies assessing community structure using lipid analyses (*8*), cell abundances (*15*), and meta-omics (*13*, *64*). The high diversity of archaea in the terrestrial subsurface, rivaling that of terrestrial surface ecosystems is less well documented. Similarly surprising is the high diversity of bacteria in the marine subsurface in comparison to surface ecosystems, supporting previous findings that focused on marine sediments (*16*). This work collectively improves our understanding of the contribution of archaea and bacteria to global microbial diversity and ecosystem function. To further investigate the global trend and corroborate the high diversity of archaeal and bacterial diversity in marine subsurface ecosystems, future studies should include datasets of additional surface environments such as soils and freshwater sediments, as those were underrepresented in our datasets.

**Fig. 4.**
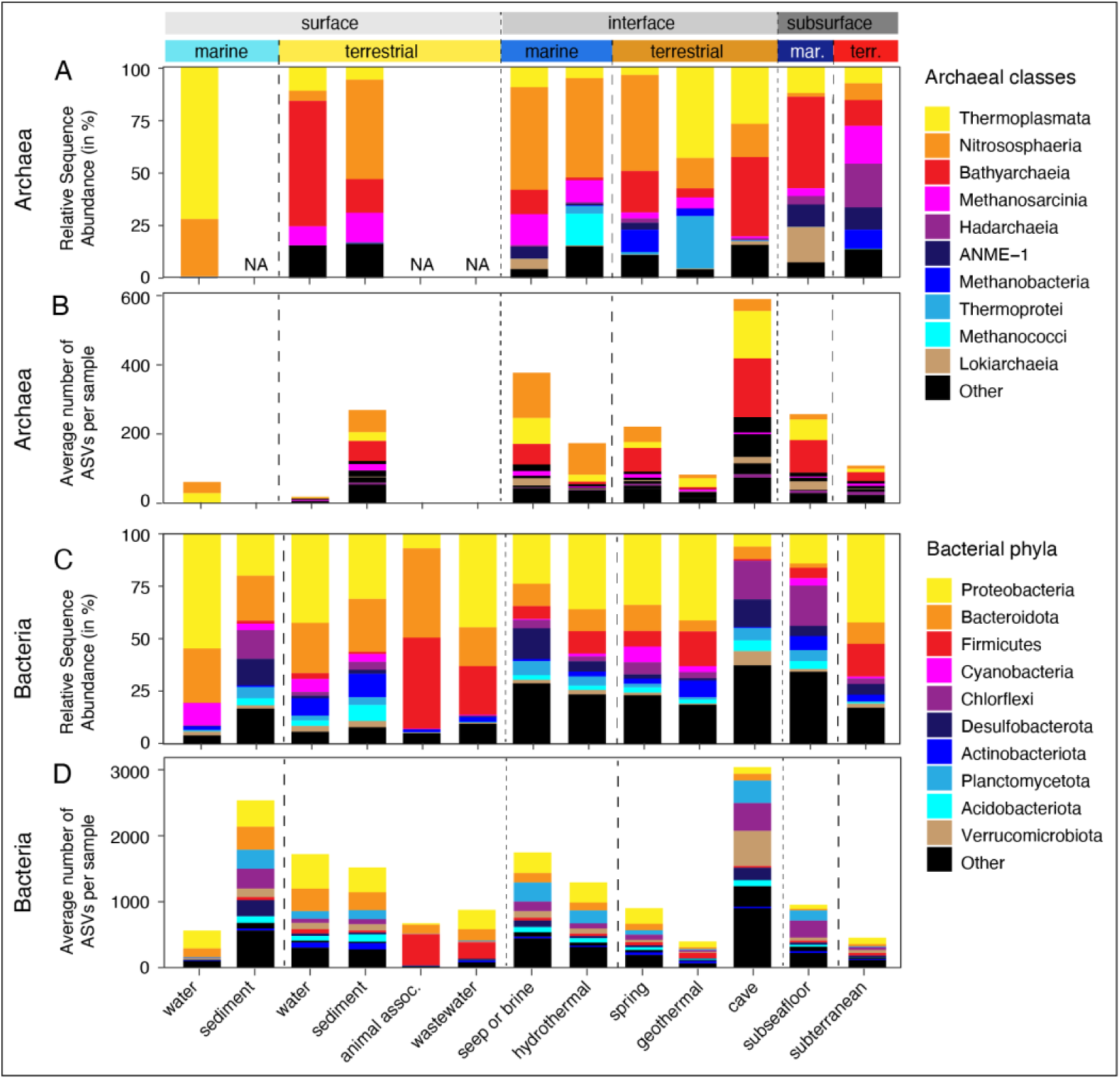
Relative sequence abundance and richness of most important lineages across environments. Relative sequence abundance of top 10 most abundant archaeal classes (A) and bacterial phyla (C) in the 13 studied environments. Contribution of most abundant lineages to average number of archaeal (B) and bacterial (D) ASVs per sample (richness).

### Interface and subsurface microbiomes are phylogenetically more diverse in marine than in terrestrial biomes

Microbial species-level diversity was significantly higher in marine interface and subsurface environments than in terrestrial interface and subsurface environments (Fig. 3C, D, S7). This was the case for archaea and bacteria, and for all tested alpha diversity indices concerning richness, evenness, and estimated richness, as well as gamma diversity. We corroborated this ASV-based trend through comparisons of the diversity of rpS3 and pF16S genes (Fig. S7), for which marine interface and subsurface microbiomes were also either significantly more diverse or statistically indistinguishable. Beta diversity between the marine and terrestrial subsurface also substantially differed based on ASVs (Fig. S9A, B), showing that analogous to surface ecosystems marine and terrestrial subsurface environments harbor many unique microbial species and represent fundamentally different habitats. Generally, the average community dissimilarity between terrestrial samples was greater than between marine samples. The processes that lead to lower richness and greater dissimilarity in the terrestrial subsurface cannot be explained with our dataset but could be due to fundamental properties of the environment, including high habitat heterogeneity, high disturbance, or low dispersal. The diversity trends were also supported by rpS3 and pF16S gene-based analyses (Fig. S9C, D).

### Abundant microbial lineages in the subsurface

Archaeal and bacterial community composition also differed between the surface and subsurface environments in terms of lineage identity, relative sequence abundance and prevalence, *i.e.*, the percentage of samples in which a lineage occurred (Fig. 5, 6, S11-13). Certain archaeal and bacterial lineages were significantly more prevalent in samples from subsurface than surface ecosystems. Most archaeal phyla predominantly occurred in the subsurface (Fig. 5B), except for *Thermoplasmatota*, which due to their abundance in the ocean are surface-associated, and *Crenarchaeota,* which have low statistical support for a preferred surface occurrence, likely caused by the class *Bathyarchaeia* which occurs in almost all marine and terrestrial subsurface samples (Fig. 6, S11). Among the *Bacteria* the phyla *Firmicutes*, *Caldatribacteriota, Nitrospirota, Patescibacteria, Aerophobota, Chloroflexi, Desulfobacterota, Methylomirabilota,* and *Spirochaetota* are among those that are more likely found in the subsurface (Fig. 5A, Fig. S12, S14). This finding supports previous reports of microbes that are abundant in deep subsurface environments including the sulfate-reducing *Candidatus* Desulforudis audaxviator (*65–67*), *Spirochaeta* (*68*) or organisms affiliating with (*Cald)atribacteriota* (*69–75*). In contrast, *Cyanobacteria, Bdellovibrionota, Fusobacteriota, Verrucomicrobiota, Planctomycetota, Bacteroidota, Campilobacterota*, and *Proteobacteria* are more widespread in surface ecosystems.

**Fig. 5.**
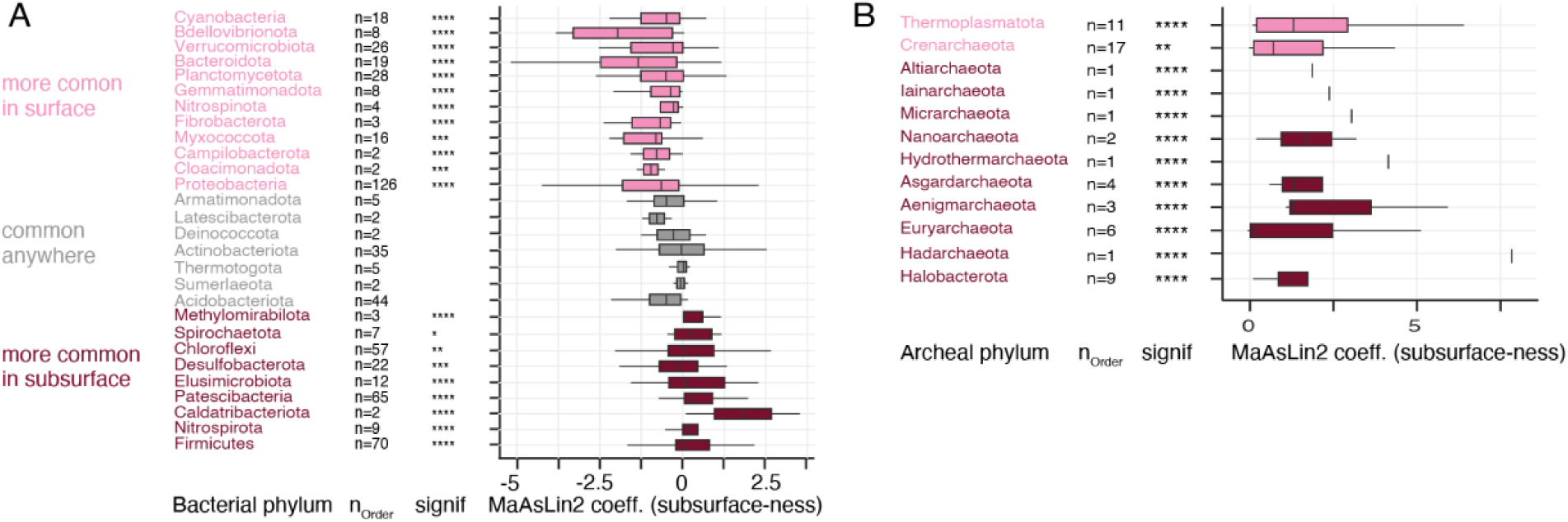
Multivariate association analyses of microbial lineages. Analyses compare the occurrence of bacterial (A) and archaeal (B) phyla in surface vs subsurface realms. The phyla are ordered from top to bottom based on increasing likeliness of their occurrence in subsurface-derived samples (increasing MaAsLin2 coefficients; “subsurface-ness”). Boxplots summarize order level MaAsLin2 coefficients within the listed phyla. Note: due to ease of visualization boxplots are even shown for very small number (n) of orders. The significance of the MaAsLin2 phylum level coefficient is shown in the column denoted “signif”. Significance levels are: *: p<0.05, **: p<0.01, ***: p<0.001, ****: p<0.0001. Additional bacterial phyla are shown in Fig. S14.

**Fig. 6.**
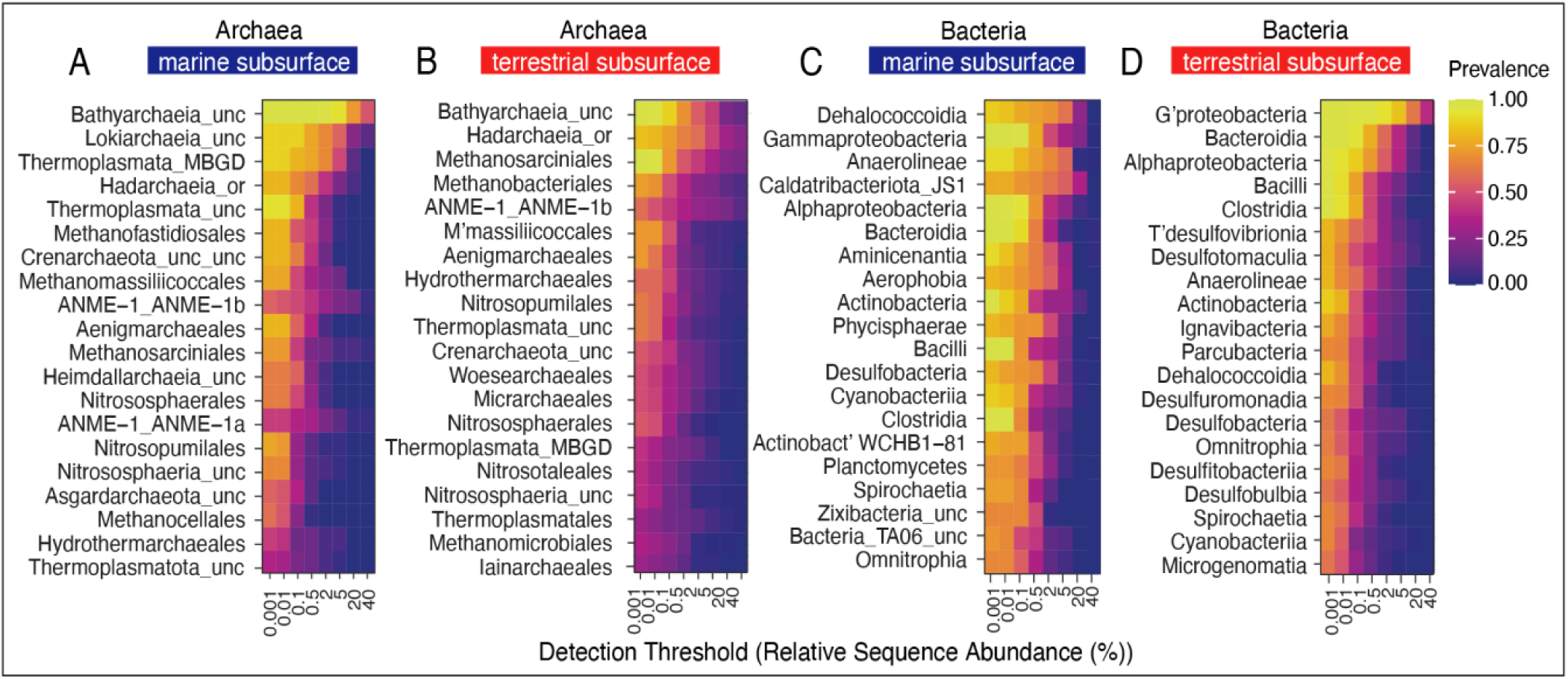
Subsurface core microbiomes. The heatmaps show potential archaeal order level core microbiomes in the marine (A) and terrestrial subsurface (B), and bacterial class level core microbiomes in the marine (C) and terrestrial subsurface (D). The 20 most relevant lineages, regarding their prevalence and relative sequence abundances, are shown as rows, each column color represents the lineages prevalence at a certain RSA threshold. Uncultured/unclassified lineages are denoted by “_unc” and the closest known phylogenetic level or the closest phylogenetic level with an isolated representative is shown. For example, Bathyarchaeia_unc are all uncultured/unclassified/ order level clades in the class Bathyarchaeia.

We have grouped the most abundant subsurface class and order level lineages based on their relative sequence abundances (RSAs) and prevalence. The first group comprises lineages with high RSAs and a high prevalence, *e.g.*, *Bathyarchaeia* in the marine subsurface or *Gammaproteobacteria* in the terrestrial subsurface. Both lineages occur in all samples (prevalence = 1) at up to 2 % RSA (Fig. 6A, D). *Bathyarchaeia* still occur at more than half of all marine samples (prevalence >0.5) with RSAs of >40 % (Fig. 6A). The second group comprises lineages that have a high prevalence and low RSAs, *i.e.*, they occur at almost all sites but are mostly rare. This group comprises terrestrial subsurface *Bathyarchaeia* and *Methanosarciniales*, as well as *Bacilli, Clostridia, Bacteroidia, Alphaproteobacteria,* and *Actinobacteria* in both the marine and terrestrial subsurface (Fig. 6C, D). The third group features those with low prevalence and high abundance, which means they do not occur at all sites, but when they occur, they are very abundant or even dominant. This group includes marine-associated *Lokiarchaeia* and ANME-1a, as well as *Hadarchaeia* and ANME-1b, which exhibit this distribution in both the marine and terrestrial subsurface. In the bacterial domain, examples include JS1 *Caldatribacteriota*, which constitute up to 20 % of the community in about half of the marine subsurface samples (Fig. 6C), as well as marine *Dehalococcoidia, Aminicenantia, Aerophophia, Phycisphaerae,* and *Desulfobacteria.* The fourth group represents organisms with a low prevalence and low abundance. These lineages are generally rare, and their occurrence may be stochastic. Depending on an emphasis on prevalence or abundance, the lineages in the first three groups can be interpreted as a core microbiome of the marine or terrestrial subsurface. Although there is little community overlap at species-level, there are lineages at higher phylogenetic levels that occur in both biomes, including *Bathyarchaeia*, *Hadarchaeia*, ANME-1b, *Alpha*- and *Gammaproteobacteria*, *Bacteroidia, Bacilli,* and *Clostridia* which can be considered a subsurface core microbiome.

### Phylogenetic novelty in the subsurface

Many lineages that are abundant in subsurface environments belong to branches in the tree of life that have few or no cultured representatives, *e.g.*, *Loki*-, *Bathy*-, and *Hadarchaeia*, and *Caldatribacteriota*. Together with their often outsized contribution to richness (Fig 3F) it is likely that the subsurface holds a substantial degree of phylogenetic and likely genomic novelty. To test whether the described high richness in interface and subsurface environments indeed represents phylogenetic novelty, we analyzed the phylogenetic proximity of the detected lineages to their closest isolated relatives. For this, we mapped ASV against a database of cultivated isolates. Our analyses show that most archaeal ASVs in the marine subsurface are only 80-90 % sequence identical to the closest isolated relative (Fig. 7A), on average 88 % (Fig. S15A), indicating that novel family-, order- or potentially even class-level clades can be expected (*76*). In the marine surface and interface and in terrestrial environments archaeal ASVs are on average more closely related to a cultured isolate (95-97 % sequence identity, Fig. 7A, S15), nevertheless representing potential novel lineages. In the case of bacteria, average sequence identity to the closest isolate is also lowest in the marine subsurface (88 % on average), supporting the notion of extensive phylogenetic novelty in the seafloor (*16*). However, bacterial phylogenetic novelty is similarly high in all other environments, with most ASV being only ∼90 % sequence identical to the closest isolate (Fig. 7B), ranging on average from 89 % - 92 % (Fig. S15). Notably, many of the ASVs that are phylogenetically distant from their closest cultured relative have high relative sequence abundances (Fig. 7). This is especially true for *Archaea*, for which several phylogenetically distant ASV had relative sequence abundances between 1-10 % (Fig. 7A). These findings support the notion of a largely untapped reservoir of uncultivated archaeal diversity in the subsurface (*77*). But our findings highlight extensive novelty in most other environments, and in the bacterial domain.

**Fig. 7.**
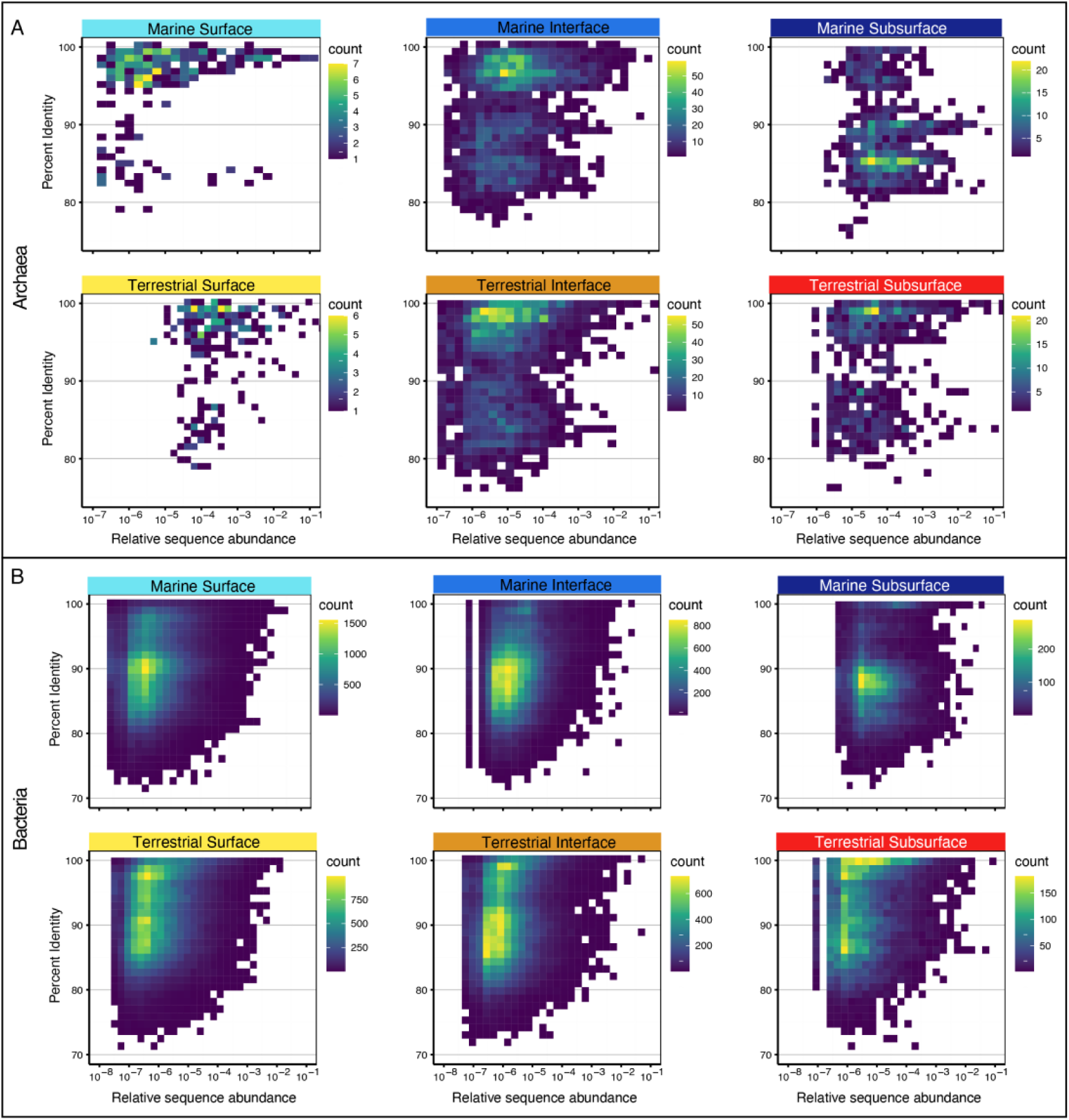
Microbial phylogenetic novelty. Percent identity values (PIVs) of archaeal (A) and bacterial (B) amplicon sequence variants (ASVs) relative to their closest cultured relative for the studied biomes and realms. Each square in the density plot represents one ASV and depicts its PIV (y-axis) and relative sequence abundance (RSA, x-axis, logarithmic). The color gradient shows how many ASVs have identical PIV and RSA. The more yellow, the more ASVs are represented by the square.

### Global continua and great divides

Overall, we demonstrate that communities in the marine and terrestrial subsurface can be as phylogenetically diverse as those on the surface, harboring a substantial part of Earth’s biodiversity. This finding confirms previous insights from global marine sediments (*16*) and the terrestrial subsurface (*54*) yet bridges the two distinct biomes in one global analysis. Our comparison of marine and terrestrial microbiomes and depth realms shows that microbial diversity is particularly high in interface environments that are influenced by surface and subsurface processes. The communities at interface ecosystems shared community membership with those of surface and subsurface environments, which reflects the location, as well as the exchange of fluids and other materials between the depth realms. Mud volcanoes and methane seeps have been shown to connect surface and subsurface environments (*31–33*), as have marine crustal fluids (*78*, *79*). Archaeal interface communities shared more genus-level clades with the subsurface (Fig. 8A, B), whereas bacterial interface communities shared more genus-level clades with the surface (Fig. 8C, D), suggesting that different ecological processes shape archaeal and bacterial communities.

**Fig. 8.**
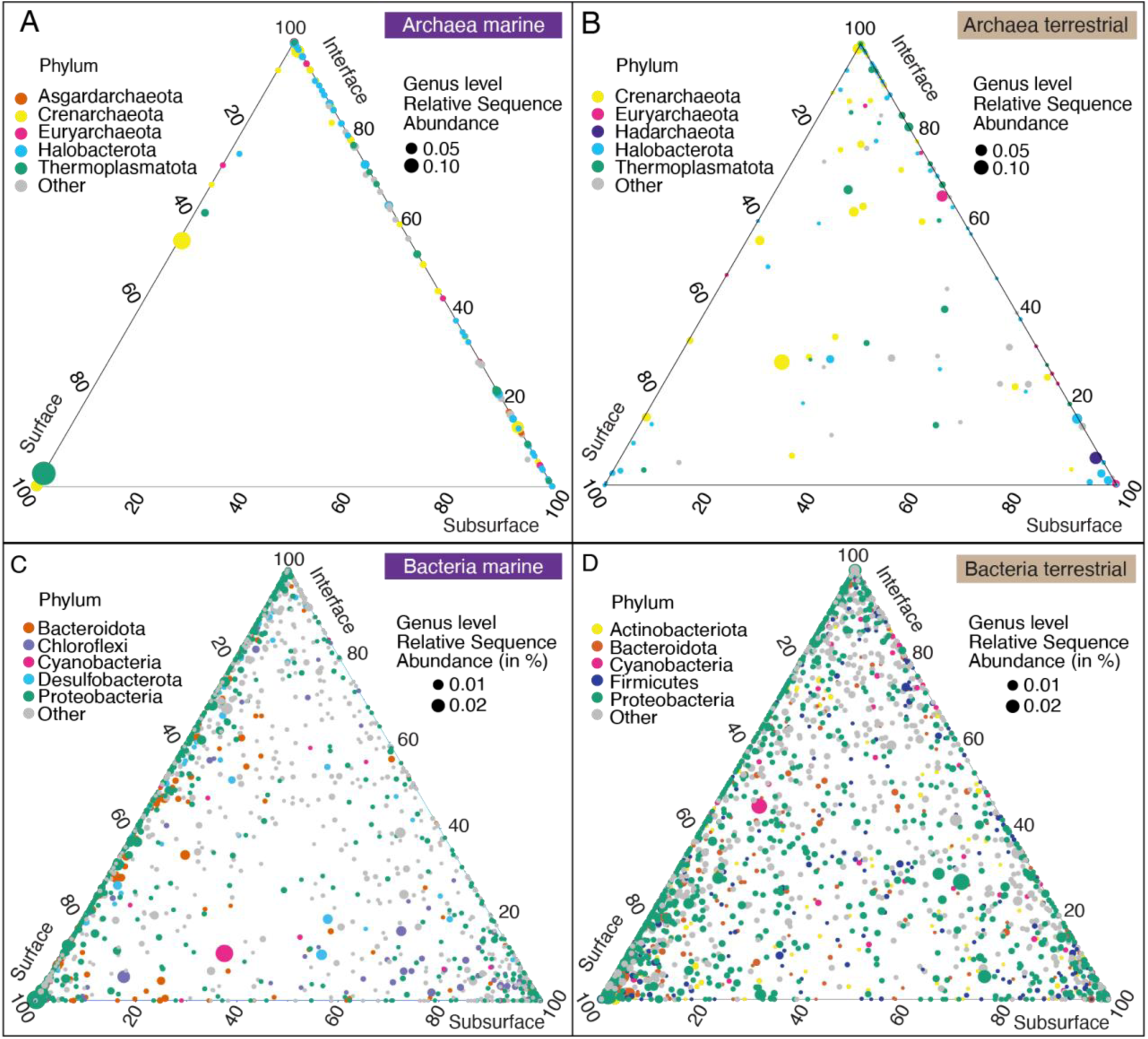
Genus-level diversity within and between depth realms. Ternary plots show archaeal (A, B) and bacterial genus-level diversity (C, D) in the marine (A, C) and terrestrial biome (B, D). Each circle is a genus-level clade, circle size is average relative sequence abundance in the respective biome, circle colors represent to which of the top 5 phyla the lineage belongs. The location of the circle shows the average relative sequence abundance in each depth realm (surface, interface and subsurface). For example, if a circle lies exactly in the center of the plot, the respective lineage is equally abundant (33.3 %) in each of the three depths, if it lies at one of the corners (*e.g.*, surface) it means it occurs only in the surface, circles that lie on the sides of the plot occur in only two of the three depths.

Despite harboring many distinct microbial lineages, the microbiomes of surface, interface, and subsurface environments showed a relatively high overlap in community composition, suggesting a diversity continuum with depth (Fig. S9, Video S1, S2) that connects surface and subsurface communities across environments and deep time. Moreover, many of the environments we studied cannot be easily defined as surface or subsurface environment. Subsurface or subsurface-impacted ecosystems are not necessarily energy depleted and, in fact, can be relatively energy rich - such as hydrocarbon seeps (*80*), brines (*81*), hydrothermal vents (*82*), oil, coal and shale deposits (*51*, *83*, *84*), serpentinizing systems (*85–87*), and caves (*88*). The subsurface is often replete with methane, hydrogen, and other energy and carbon sources (*45*, *89–92*). An absence of oxidants, nutrients, trace metals, or extreme physical or chemical conditions, however, can shape the specific community structures and hinder microbial growth or activity in relatively energy-rich ecosystems (*93*, *94*). Depth and oxygen content are also not always reliable proxies for subsurface conditions, *e.g.*, in marine sediments with very low organic matter content, oxic conditions can prevail throughout the sediment column (*95*).

Sediment depth and sediment age can be vastly different between locations (*56*), with past surface conditions or depositional environments (Fig. S16) apparently being imprinted into a given subsurface sample (*24*, *25*). Defining environments and samples by a few select parameters such as depth, oxygen content, or age is useful, but to fully understand the subsurface geobiosphere we may need to consider large-scale gradients and continua, analogous to the concept of the critical zone in soil science (*96*, *97*).

Our analyses further reveal a major divide between marine and terrestrial microbiomes, showing significant differences in local and overall richness. In contrast to the studied depth realms, the differences between marine and terrestrial microbiomes were striking, with little overlap in community structure (Fig. 2, S2). This divide was also found based on the analysis of metagenome-derived 16S rRNA and rpS3 genes (Fig. 2, S4). The taxonomic differentiation between marine and terrestrial microbial communities mirrors the differences between animal communities in these biomes (*98*) and is likely caused by the very different chemical and physical properties that shape these environments (*99*). The environmental drivers responsible for the gradients, continua, and divides remain undefined but they likely include factors such as salinity (*100*, *101*), energy availability (*57*), geological activity (*102*, *103*), hydrologic conditions and recharge rates (*104*, *105*), and constrictions of void space within which microbial cells might exist (*94*). Improvements in global scale modeling and increasing availability of data can provide the tools to describe the surface and subsurface on large scales, refine our insights into microbial provinces in the subsurface (*106*), and potentially define regions of similar subsurface biogeochemical regimes analogous to Longhurst provinces for the surface ocean (*107*).

## Material and Methods

### Dataset specifications

To investigate the microbial diversity and composition of surface, interface, and subsurface microbiomes, we used 478 archaeal and 1054 bacterial 16S rRNA gene amplicon datasets sequenced at the W. M. Keck Ecological and Evolutionary Genetics Facility of the Marine Biological Laboratory (Woods Hole, MA, USA). Most archaeal and bacterial datasets from subsurface and interface environments were part of the Census of Dep Life (CoDL), and most datasets from surface ecosystems were not collected during the CoDL but were kindly shared by numerous other investigators (see Acknowledgements). All sequencing datasets were prepared using identical primers, nearly identical library preparation protocols, identical sequencing chemistries, and bioinformatic analyses to minimize batch effects, biases and ensure comparability of communities based on amplicon sequence variants (ASVs). We removed datasets from enrichment samples, blanks, and controls, as well as datasets that we considered as failed runs (< 2000 bacterial reads or <1000 archaeal reads) and subsurface datasets that contained lineages belonging to known contaminants of subsurface samples (*59*). Contextual data for each sample/sequence dataset is listed in Table S2. Dataset 1 and 2, representing the archaeal and bacterial analysis logs using the “biome” grouping as a representative workflow). We are aware that certain environments are overrepresented (*e.g.*, marine surface), while others are underrepresented (*e.g.*, caves) or missing (*e.g.*, soils). However, the highlighted global trends are likely reliable due to the size and breadth of the dataset, the comparability of environments and robustness of the results (*e.g.*, towards the removal of reads, Fig. S8).

### 16S rRNA gene library preparation and amplicon sequencing

Genomic DNA was extracted by the laboratories responsible for each project (Table S1). Sequencing was performed at the W. M. Keck Ecological and Evolutionary Genetics Facility. The bacterial 16S rRNA gene V4-V5 variable region was amplified using the forward primer 518F (3’-CCAGCAGCYGCGGTAAN-5’) and reverse primers 926R (3’-CCGTCAATTCNTTTRAGT-5’, 3’-CCGTCAATTTCTTTGAGT-5’, 3’-CCGTCTATTCCTTTGANT-5’) (*108*). The archaeal 16S rRNA gene V4-V5 variable region was amplified using the forward primers 517F (3’-GCCTAAAGCATCCGTAGC-5’, 3’-GCCTAAARCGTYCGTAGC-5’, 3’-GTCTAAAGGGTCYGTAGC-5’, 3’-GCTTAAAGNGTYCGTAGC-5’, 3’-GTCTAAARCGYYCGTAGC-5’) and reverse primer 958R (3’-CCGGCGTTGANTCCAATT) (*109*). Amplicons were sequenced using Illumina’s v3 600-cycle (paired-end) reagent kit on a MiSeq (Illumina Inc., San Diego, CA, USA). Reads were demultiplexed based on the combination of index (CASAVA 1.8) and barcode (custom python scripts).

### 16S rRNA gene amplicon-based community analyses

Raw sequences were analyzed using DADA2 (*110*) following the DADA2 Pipeline Tutorial v1.16 (https://benjjneb.github.io/dada2/tutorial.html). Each Illumina run was analyzed separately to use run-specific error profiles, as is recommended best practice for big data (https://benjjneb.github.io/dada2/bigdata.html). In brief, forward and reverse reads were quality-trimmed to 275 bp and 205 bp, respectively, and primer sequences (17 bp forward, 21 bp reverse) were removed. Reads with more than two expected errors were discarded and paired reads were merged. All runs were combined, and chimeric sequences were removed. Species level taxonomy was assigned with the silva_nr_v138_train_set and silva_species_assignment_v138 based on the Silva small subunit reference database SSURef v138 (release date: 16-Dec-2019; (*111*)). After quality control and the removal of enrichments, blanks, biological and technical replicates, we obtained 478 archaeal and 1054 bacterial amplicon datasets. Archaeal datasets contained a total of 4.9 × 10^7^ sequence reads belonging to 31099 unique amplicon sequence variants (ASVs). Archaeal samples had on average 1.02 × 10^5^ reads and 155 ASVs (Table S3). Bacterial datasets comprised a total of 13 × 10^7^ sequence reads belonging to 415423 unique ASVs. Bacterial samples had on average 1.2 × 10^5^ reads and 1257 unique ASVs (Table S4). Amplicon datasets may not quantitatively represent the sampled community yet are reliable to compare community structure and make ecological interpretations (*62*). Alpha diversity (richness, Shannon entropy, Inverse Simpson Diversity and Chao1 estimated richness) was calculated from the ASV-by-sample table using a subsampling approach to account for unequal sampling effort. We used 1142 and 2216 randomly chosen reads from each archaeal and bacterial sample, respectively. Even when using very stringent subsampling conditions of 50000 randomly chosen archaeal and bacterial sequences, respectively, the trends in alpha and beta diversity did not change substantially, hence we decided to include as many samples as possible. Bray-Curtis dissimilarities (*112*) between all samples were calculated and used for two-dimensional nonmetric multidimensional scaling (NMDS) ordinations with 20 random starts (*113*). All analyses were carried out with VisuaR (https://github.com/EmilRuff/VisuaR) - a workflow based on the R statistical environment, custom R scripts and several R packages including *vegan* (*114*) and *ggplot2*. Exemplary analysis logs (DatasetS1, S2), the used DADA2 script (DatasetS3), the VisuaR v40 script (DatasetS4) and an example VisuaR user input file (DatasetS5) for this study are available as supplementary materials.

### Gene sequence identity analyses

To determine the relatedness of detected 16S rRNA gene sequences to those of the closest isolated strain, we performed a BLASTn search of each 16S rRNA gene amplicon sequence against a database of sequences from cultured archaeal and bacterial isolates, selected to yield only the single best hit for each amplicon. The isolate database was created with *makeblastdb* using sequences from cultured isolates in the SILVA NR Ref database v123 available at (https://www.arb-silva.de/). We included recently cultured (*Cald)atribacteria* (*73*). Only alignment lengths > 350 (97 % of bacterial amplicons and 64 % of archaeal amplicons) were analyzed.

### Metagenomic DNA sequencing, contig assembly, gene annotation and count

Paired-end libraries (2×151 bp) were prepared for a subset of samples using Illumina TruSeq DNA library preparation kit (Illumina, San Diego, CA, USA) and sequenced on the Illumina NextSeq500 platform. Raw demultiplexed reads were quality processed using bbduk.sh in BBMap v.38.71 (Bushnell B. – https://sourceforge.net/projects/bbmap/) in two successive steps, by first removing Illumina adapters from read termini (with options: ktrim=r minlen=40 mink=11 tbo tpe k=23 hdist=1 hdist2=1 ftm=5) prior to quality filtering against PhiX genome to an average quality threshold of 12 (with options: maq=12 trimq=12 qtrim=rl maxns=3 minlen=40 k=31 hdist=1). Quality trimmed reads mapping to the human HG19 genome with over 95% identity were discarded using bbmap.sh in BBMap (with options: local minratio=0.9 usemodulo maxindel=3 bwr=0.16 bw=12 quickmatch fast minhits=2 qtrim=rl trimq=12 untrim idtag printunmappedcount kfilter=25 maxsites=1 k=14). Cleaned reads were used to construct de novo assemblies for each sample using metaSPAdes (v3.13.0) with default parameters (*115*). Open reading frames (ORFs) were predicted on assembled contigs ≥ 1kb using Prodigal (v2.6.3) (setting -p meta) (*116*). Predicted genes were clustered at ≥ 95% sequence identity and ≥ 80% overlap using dedupe.sh in BBMap (with option arc=t am=t ac=t mid=95 mlp=80) to produce a unique gene catalog of 15,663,206 non-redundant genes. The unigene was functionally assigned to a KEGG Orthology (KO) using kofamscan (v1.3) (*117*). Ribosomal protein S3 (rpS3) sequences were extracted from all predicted ORFs (*118*) using hmmsearch (*119*) against custom Hidden Markov Models (HMMs) for Archaea and Bacteria (https://github.com/AJProbst/rpS3_trckr), yielding a total of 8,691 rpS3 sequences. RpS3 amino acids were de-replicated at ≥ 99% identity and ≥ 80% overlap with dedupe.sh (with options: arc=t am=t ac=t mid=99 mlp=80) to produce 4,392 rpS3 single gene sequences, equivalent to microbial species. Taxonomic classification of rpS3 genes was performed using Kaiju (v1.7.3, with option: greedy-5 mode) (*120*).

For taxonomic and gene count analyses, cleaned paired-end reads were subsampled to a depth of 2M reads per sample using reformat.sh in BBMap (with options: samplereadstarget=2000000 sampleseed=121) and then mapped back to the annotated genes using BBMap with read alignment of ≥95% for the unigene (with options: minid=0.95 idfilter=0.95 ambiguous=random) and ≥99% for rpS3 genes (with options: minid=0.99 idfilter=0.99 ambiguous=random). Gene count tables were normalized to gene per million (*e.g.*, equivalent to transcript per million, TPM), following (*121*). Metagenomic short reads were mapped to the SILVA SSU reference database (*111*) to assign nearest taxonomic units, as well as full-length 16S/18S rRNA gene sequences were reconstructed from metagenomes using phyloFlash v3.4 (*122*). Metagenome-derived rpS3 and 16S rRNA genes were subsampled (Table S3, S4), analyzed and visualized analogous to the amplicon-derived 16S rRNA genes using the same VisuaR community analysis workflow.

### Statistical analyses

Differences in alpha diversity metrics between conditions were tested using the Wilcoxon signed rank-test (*ggsignif*) as implemented in *ggplot2* (*123*). P values were corrected stringently using the Bonferroni method. We used Analysis of Similarity (*124*) as implemented in *vegan* to test whether community structure between conditions was significantly different, as visualized in NMDS plots. Multivariate associations were tested as implemented in the MaAsLin2 method (*63*).

## Supporting information

Table S1 - Dataset IDs and Accession

Table S3 - Archaeal Diversity Indices

Table S4 - Bacterial Diversity Indices

Movie S1 - 3D-NMDS Archaea

Movie S2 - 3D-NMDS Bacteria

Dataset S1 - Community Analysis Log Archaea

Dataset S2 -Community Analysis Log Bacteria

Dataset S3 - DADA2 R Script

Table S2 - Sample ID and Contextual Data

Dataset S4 - VisuaR Analysis R script

Dataset S5 - VisuaR User Input R script

## Acknowledgments

We are indebted to all our colleagues that shared their (sometimes unpublished) datasets with us: William Brazelton (U of Utah), Zoe Cardon (Marine Biological Laboratory) and Nuria Fernandez-Gonzalez (U de Valladolid), Matthew Saxton (Miami U), Lois Maignien (U of Western Brittany), Katja Meyer (U Willamette), Brooke Osborne (Brown U), Jeremy Rich (U of Maine), Adelaide Roguet, Ryan Newton, Deborah Dila and Sandra McLellan (U of Wisconsin Milwaukee), Igor Tiago (U of Coimbra), Elizabeth Trembath-Reichert (Arizona State U), Florian Trigodet (U Chicago), and Linda Amaral Zettler (Royal Netherlands Institute for Sea Research). Without including outgroup datasets generously provided by these researchers the comparison between surface and subsurface biomes would not have been possible. We are also grateful to all colleagues participating in the Census of Deep Life (CoDL), who organized field work, and collected and processed samples, as well as the expeditions and programs that enabled the sample collection. For further details on specific CoDL projects, results, site descriptions, funding and publications refer to the Supplementary Information and Table S1. Funding for sequencing was provided by the CoDL within the larger framework of the Deep Carbon Observatory. Among CoDL Principal Investigators and colleagues we are especially grateful to Panagiotis Adam, Till Bornemann, Brandon Briggs, Steve D’Hondt, Ahmed Eleish, Eric Gaidos, Adrienne Hoarfrost, Fang Huang, Tom Kieft, Rosa Leon-Zayas, Ken Locey, Magdalena Osburn, Lotta Purkamo, Karyn Rogers, and Barbara Sherwood Lollar for helpful discussions. Above all we want to thank our friend, colleague, and mentor Jan Amend for being a continuous source of inspiration, an ally of early career scientists, and a driving force in geobiology and subsurface science. This study was conceived during the Deep Carbon Observatory funded “Origins and Movement of Life” workshop, which Jan co-organized and co-hosted in 2016.

## Funding

Simons Foundation (824763, SER)

Human Frontier Science Program (RGEC34/2023, SER)

Deep Carbon Observatory (Alfred P. Sloan Foundation G-2016-7206, KGL)

Office of Biological and Environmental Research of the U.S. Department of Energy (DE-SC0020369, KGL)

CNRS Chaires de Professeur Junior (JAB)

Center for Dark Energy Biosphere Investigations (C-DEBI OCE-0939564, JAH) Deep Life I (JAH, MOS; 2011-12-01).

## Author contributions

Conceptualization: SER, AJP, MLS, JL, FC

Methodology: SER, IHDA, KGL, CS, ADS, JAB, JAH, AJP, HGM, JL, FC

Investigation: SER, IHDA, MM, JP, CM, KGL, CS, ADS, AS, AM, BKR, JAB, CL, MOS, SBJ, HGM, JL, FC

Visualization: SER, IHDA Supervision: SER, BKS, HGM, FC

Writing—original draft: SER, IHDA, KGL, JAH

Writing—review & editing: SER, IHDA, KGL, CS, ADS, JAB, MOS, JAH, AJP, MLS, JL, FC

## Competing interests

Authors declare that they have no competing interests.

## Data and materials availability

16S rRNA gene amplicon datasets and metagenomic datasets are available under the NCBI umbrella Bioproject PRJNA183206 and the individual project accessions listed in Table S1. Contextual data is available in Table S2. The R-based workflow to analyze marker gene data, *VisuaR*, is available on GitHub (https://github.com/EmilRuff/VisuaR) as well as in Supplementary Dataset S4, S5 (exemplary user input and workflow for archaeal ASV analysis grouped by “biome”).

## Notes

### Competing Interest Statement

The authors have declared no competing interest.

