## Supplementary Text and Figures for "A global atlas of subsurface microbiomes reveals phylogenetic novelty, large scale biodiversity gradients, and a marine-terrestrial divide"

S. Emil Ruff *et al.*

**This PDF file includes:**

Supplementary Text

Figs. S1 to S16

References (125 to 158)

**Other Supplementary Materials for this manuscript include the following:**

Tables S1 to S4

Movies S1 to S2

Datasets S1 to S5

Supplementary Text

*The Census of Deep Life synthesis (continued)*

Further details concerning the datasets that we used for this global synthesis project can be found on the VAMPS website (<https://vamps2.mbl.edu/>) as well as in research studies that have already been published using NCBI project accession numbers (Table S1). Datasets that were part of the Census of Deep Life (CoDL) have featured in investigations of volcanoes (*125*, *126*), deep terrestrial subsurface (*7*, *127*–*130*), methane-rich springs (*131*–*133*), mineral water springs (*134*, *135*), ultradeep bedrock (*136*), rock fracture fluids (*137*–*140*), caves (*88*), marine seafloor (*141*, *142*), marine sediments (*143*–*145*), hydrothermal sediments (*43*, *146*–*148*), hydrothermal vents (*149*), marine hydrocarbon plumes (*150*), and marine mud volcanoes (*32*, *151*). We also included amplicon datasets that were not part of the CoDL, *e.g.*, investigating marine sediments (*152*–*154*), polluted sediments and waters (*155*–*157*).

Due to the large number of datasets, and the very different interests of the researchers leading each project, only a limited set of environmental parameters was available for each sample. In addition to latitude, longitude, and depth (Fig. 1A, B), we have information on the sample material (Fig. S1A), *p*H (Fig. S1B), and temperature (Fig. S1C) for most samples.

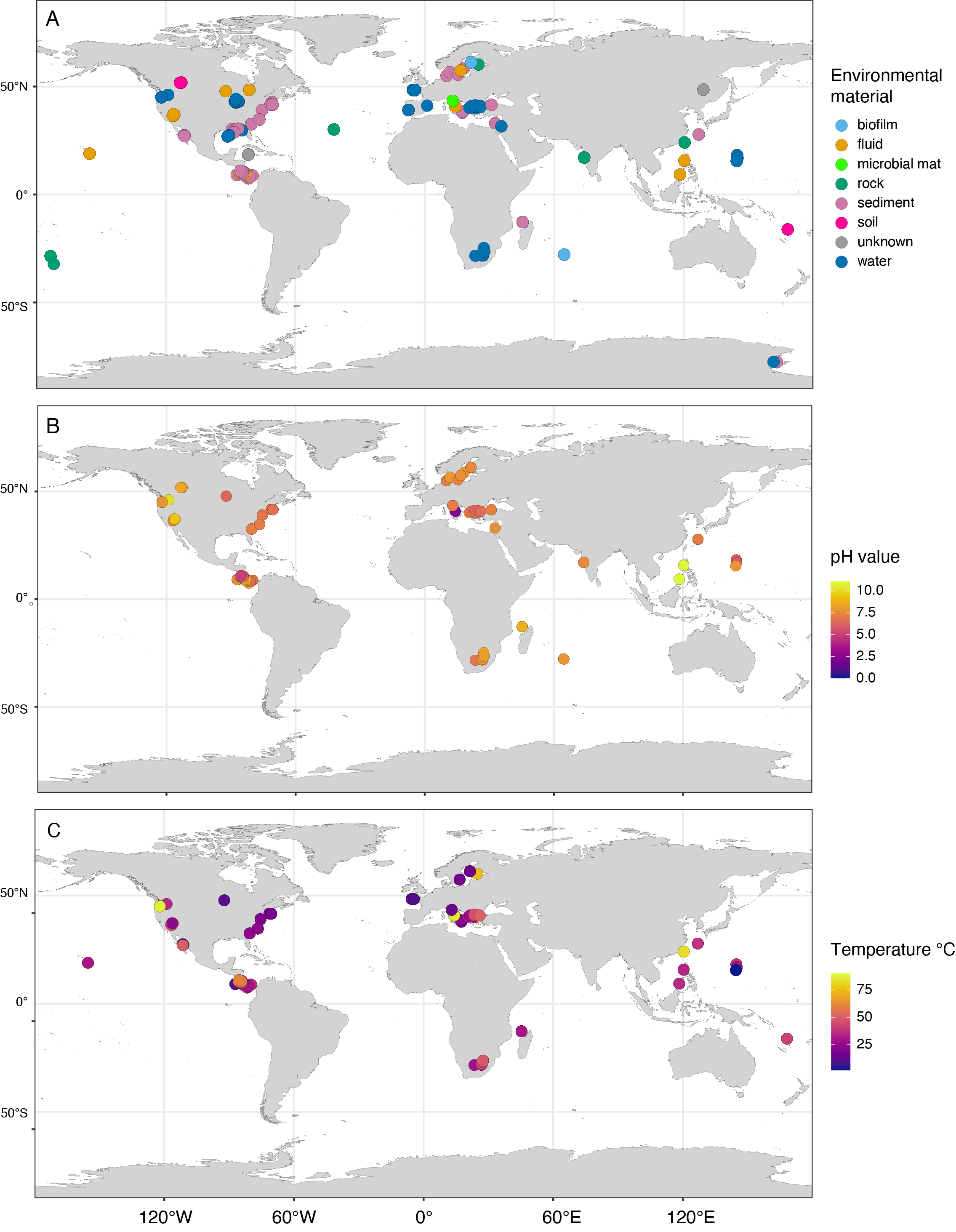

**Fig. S1: Maps of sampling sites**. Maps show the environmental material (A), *p*H (B), and temperature (C) of the collected samples.

*Substantial differences between marine and terrestrial microbiomes (continued)*

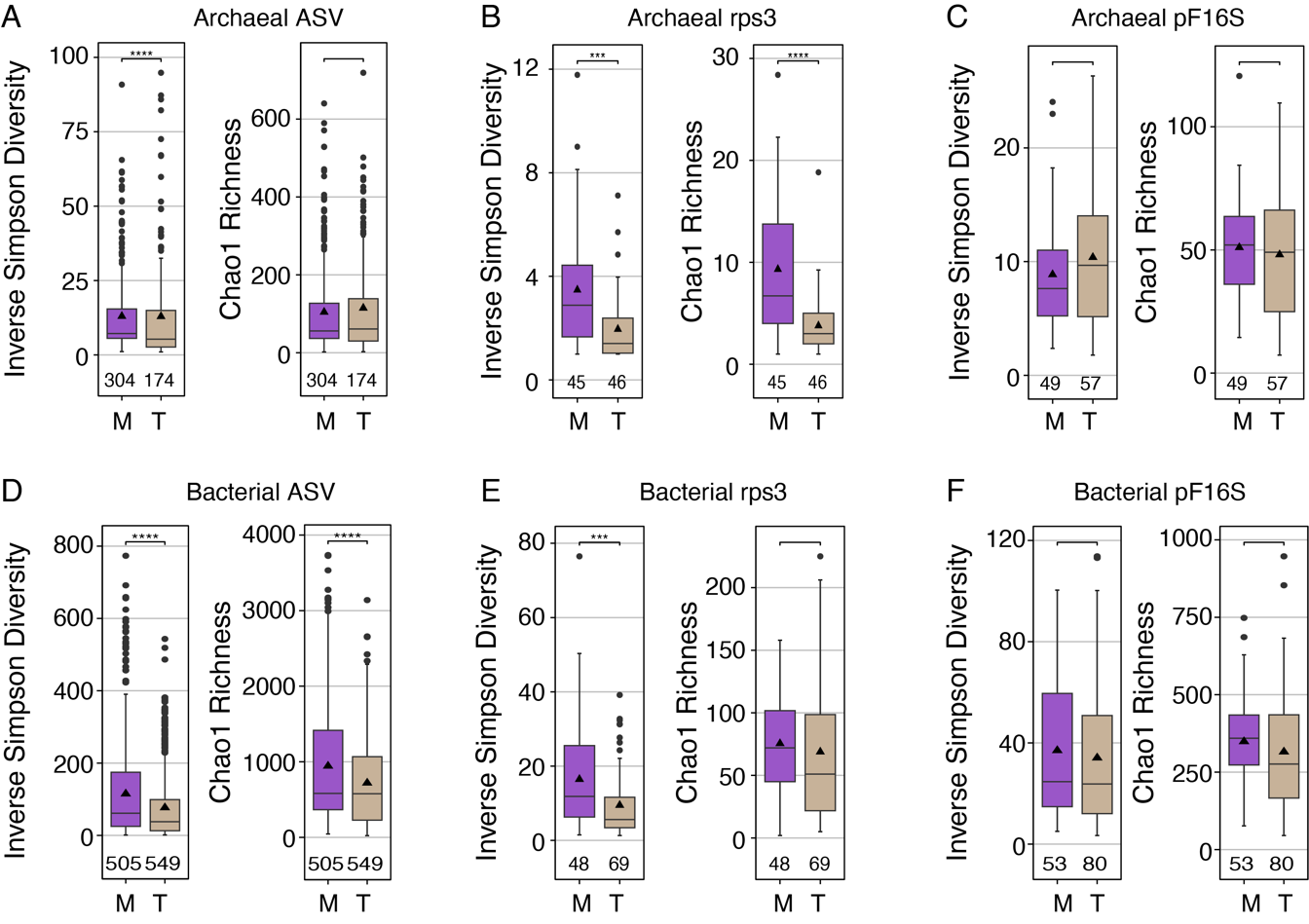

**Fig. S2: Microbial diversity indices**. Archaeal (A-C) and bacterial (D-F) alpha diversity (per sample community evenness and estimated richness) in marine (M) and terrestrial (T) microbiomes using 16S rRNA gene amplicon sequence variants (ASV; A, D), as well as metagenome-derived ribosomal protein S3 genes (rpS3; B, E) and 16S rRNA gene sequences detected by phyloFlash (pF16S). Archaeal and bacterial community evenness (inverse Simpson diversity) and estimated richness (Chao1) is similar or higher in marine biomes than in terrestrial biomes in all three methods.

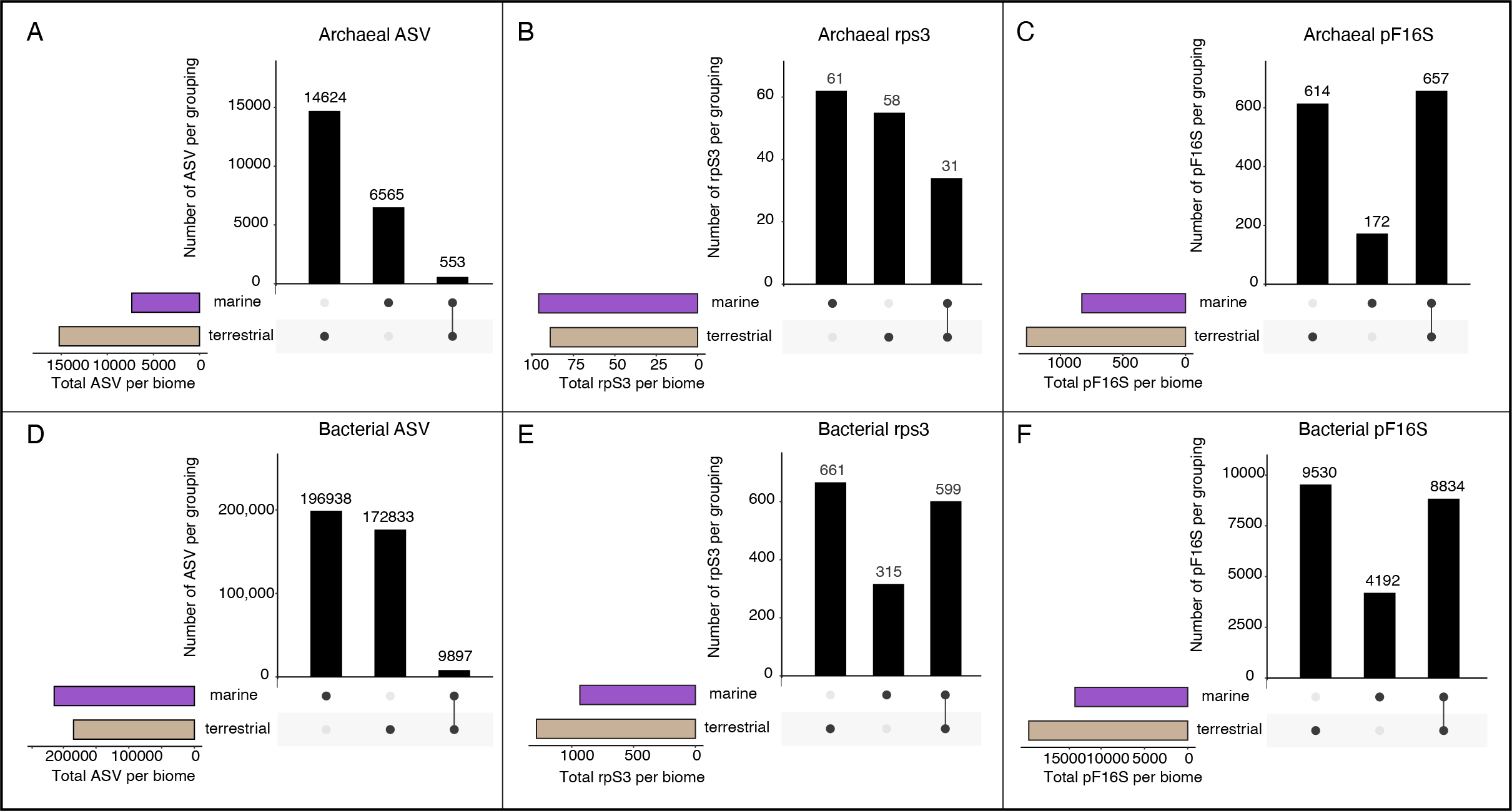

**Fig. S3: Community overlap**. Upset plots of archaeal (A-C) and bacterial communities (D-F) in marine and terrestrial microbiomes using 16S rRNA gene amplicon sequence variants (ASV; A, D), metagenome-derived ribosomal protein S3 genes (rpS3; B, E) and 16S rRNA gene sequences detected by phyloFlash (pF16S). The plots depict the same information as Venn diagrams and show how many unique sequences (ASV, rpS3, or pF16S) exclusively occur in a group or grouping (vertical bars). The total number of unique sequences in a group (gamma diversity) is shown as horizontal bars. For example, in panel A, the total number of archaeal ASV is higher in terrestrial ecosystems (light brown bar) than in marine biomes (purple bar). Within terrestrial biomes 14624 ASV occur exclusively there (first vertical bar), 6565 occur exclusively in marine biomes, and only 553 are shared between both biomes. The ASV data for panel A and D was subsampled, *i.e.*, corrected for the different number of datasets from each biome, to allow direct comparison between biomes.

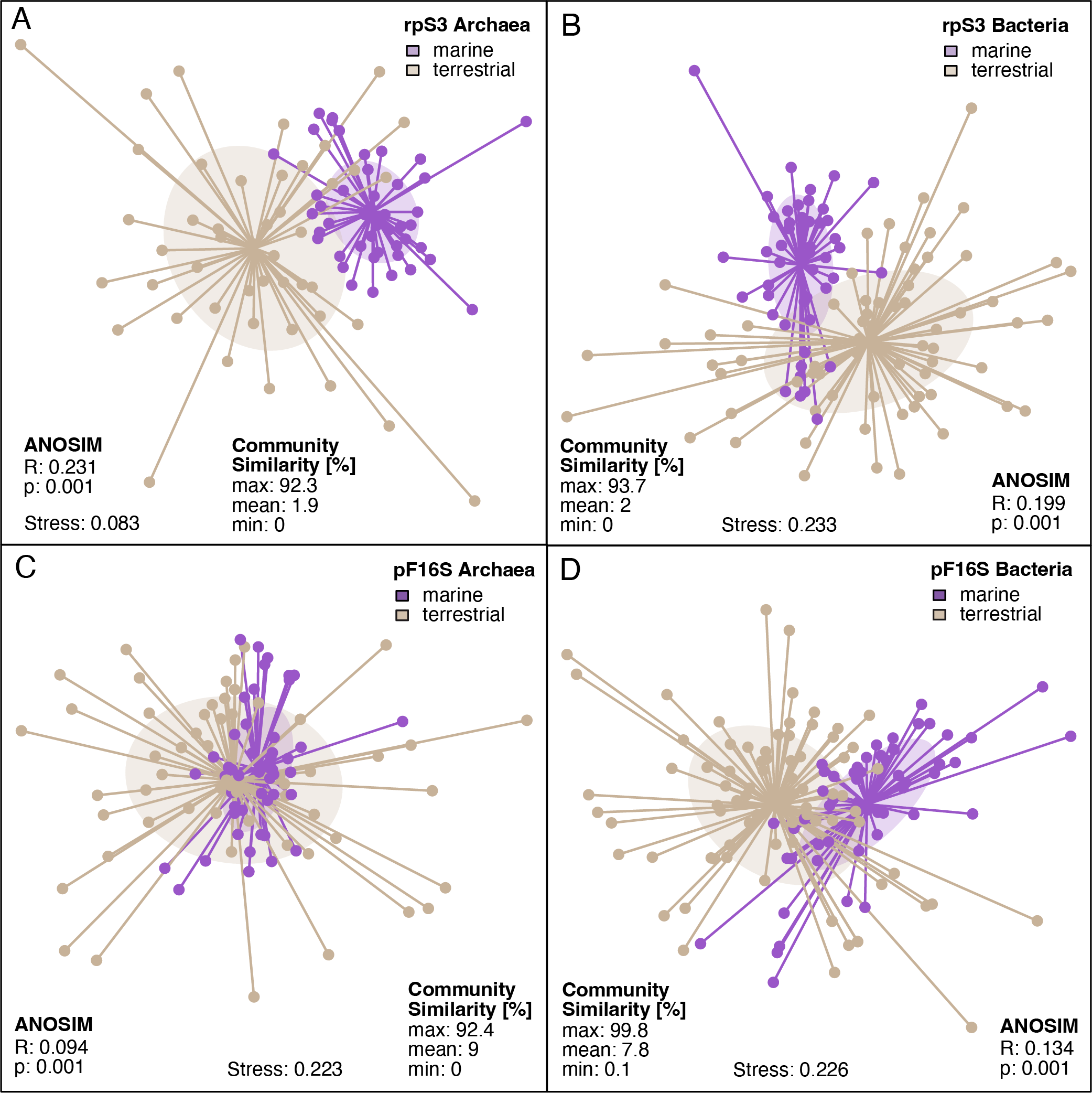

**Fig. S4: Community structure in marine and terrestrial biomes**. Plots are based on archaeal (A) and bacterial rpS3 genes (B) as well as on archaeal (C) and bacterial metagenome-derived 16S rRNA genes (pF16S; D). Apart from archaeal pF16S, the biomes show a rather small overlap, corroborating the substantial differences seen with ASV analyses, which is striking given that metagenomes are not ideal in detecting community diversity and structure, but rather useful for investigating the most abundant populations.

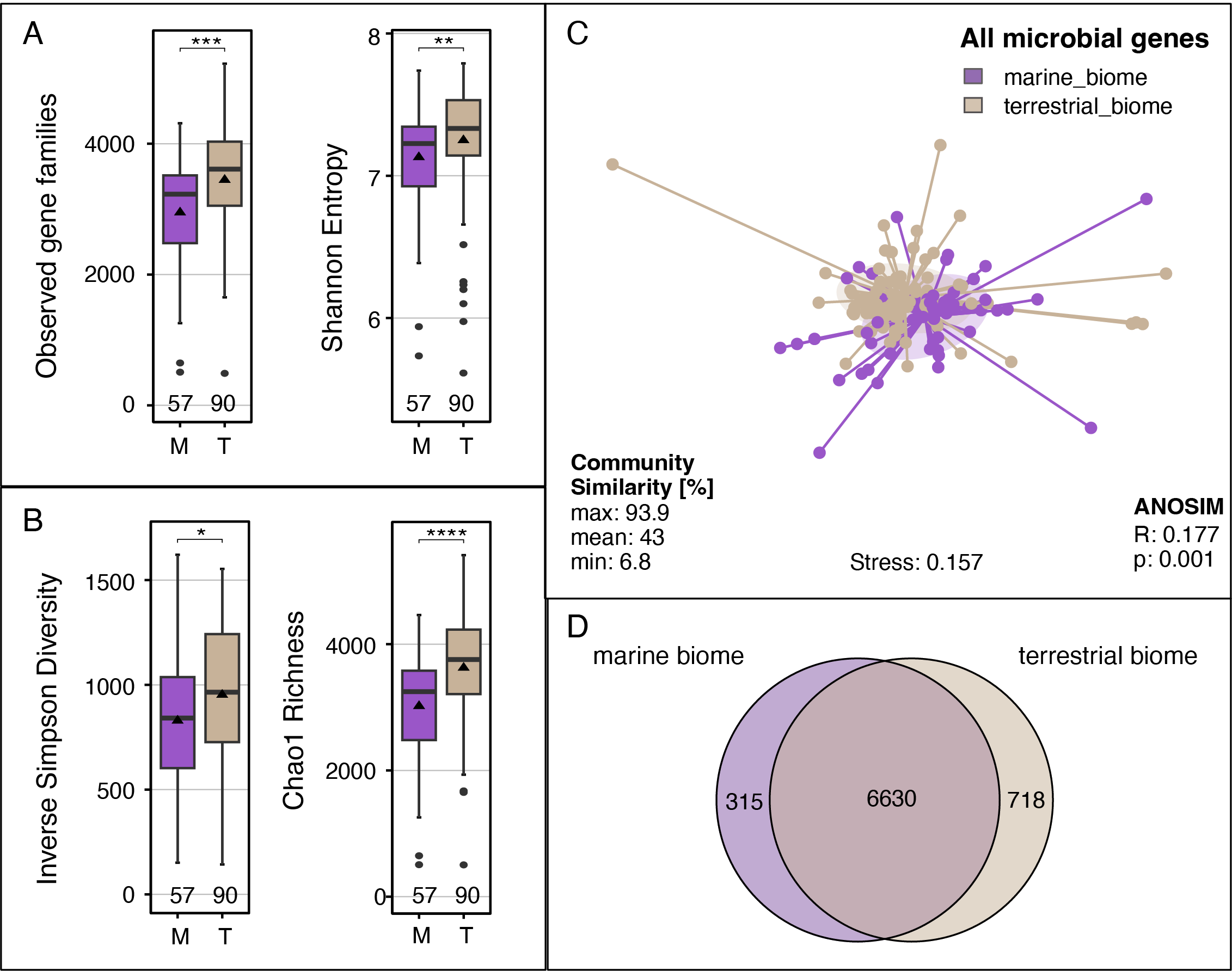

**Fig. S5: Alpha diversity in marine versus terrestrial biomes based on gene families**. Gene families were retrieved from 71 marine and 88 terrestrial metagenomes. Shown by the number of observed gene families and Shannon Entropy (A), as well as Inverse Simpson Diversity and estimated richness (B). Observed and estimated richness are very similar indicating that we have captured most of the metabolic capabilities present in these analyzed samples. The studied terrestrial ecosystems comprised significantly more gene families per sample, *i.e.* per metagenome, than marine, as tested using a Wilcoxon rank sum test. Significance levels were: *: p<0.05; **: p<0.01; ***: p<0.001; ****: p<0.0001. The number of used metagenomes is shown below the boxplots. Despite the differences, most gene families were shared among the two biomes, suggesting the communities largely contain a common set of metabolisms and housekeeping pathways, as shown in the non-metric dimensional scaling (NMDS) ordination in (C) and the Venn diagram (D).

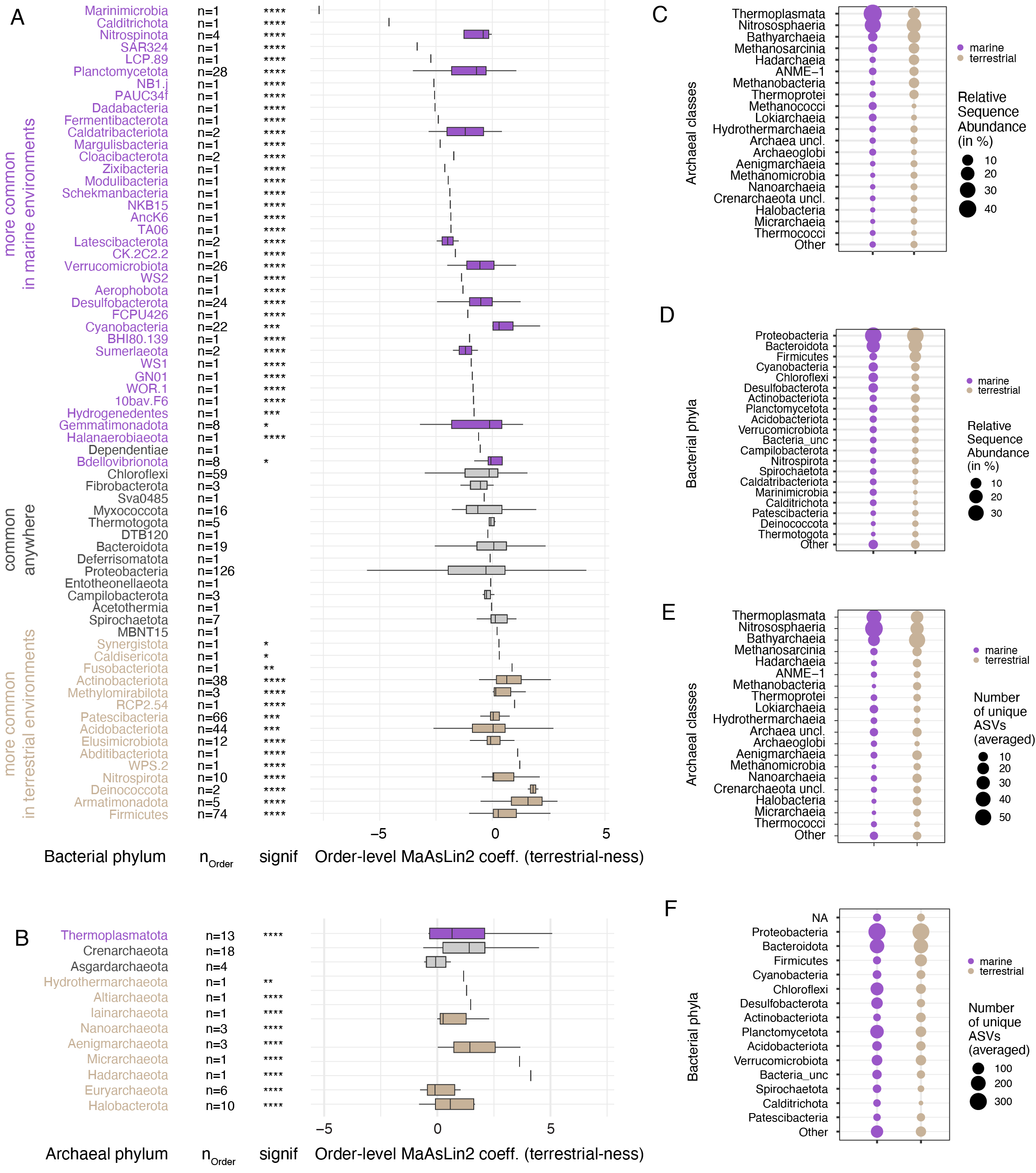

**Fig. S6: Abundance analyses of marine vs terrestrial lineages.** Differential abundance analysis of bacterial (A) and archaeal (B) phyla. The phyla are ordered from top to bottom based on increasing phylum level MaAsLin2 coefficient, *i.e.* likeliness of their occurrence in terrestrial-derived samples (“terrestrial-ness”). Boxplots summarize MaAsLin2 coefficients, i. e., “terrestrial-ness”, of orders withing the listed phyla. Note: due to ease of visualization boxplots are also shown for very small number (n) of orders; phyla and orders are based on the SILVA reference tree. The significance of this occurrence is shown in the column denoted “signif”. Significance levels are: *: p<0.05, **: p<0.01, ***: p<0.001, ****: p<0.0001. Phyla in brown are found significantly more often in the terrestrial biome, while phyla in purple are found significantly less often in the terrestrial biome, *i.e.*, occur more often in the marine biome. The analysis was done using MaAsLin2 on phylum and order level (*63*). Relative abundance of archaeal classes (C) and bacterial phyla (D) found in marine and terrestrial biome (C). The 20 most abundant classes are shown, remaining classes are shown as “Other”. (D) Total number of unique ASVs found in respective classes (E) and phyla (F).

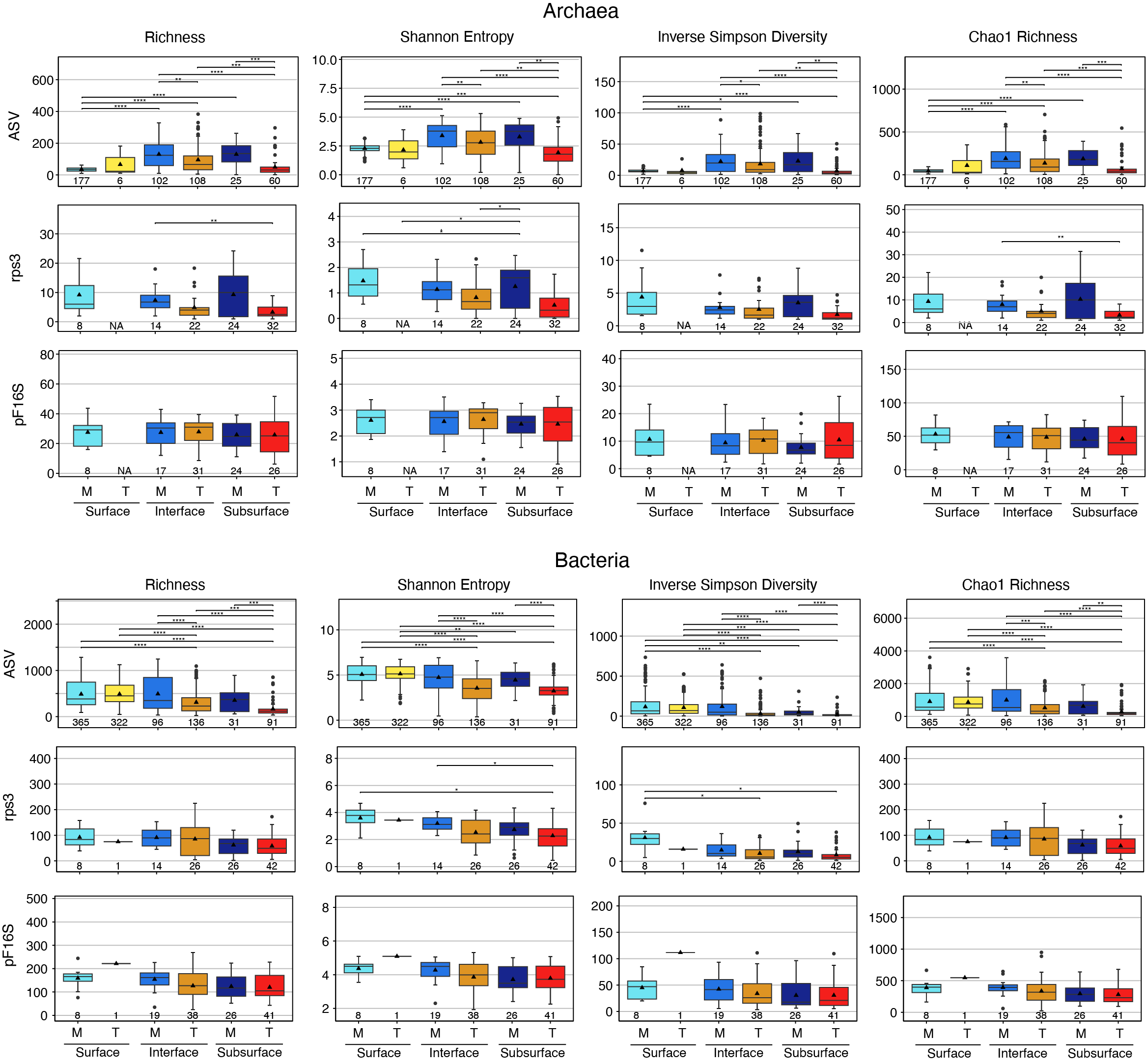

**Fig. S7: Alpha diversity indices.** Alpha diversity of archaeal (A-D) and bacterial communities (E-H) in sequencing datasets from marine (M) and terrestrial (T) surface, interface, and subsurface ecosystems. Columns show four major diversity indices (observed richness, Shannon Entropy, Inverse Simpson Diversity and Chao1 estimated richness), rows show which data was used to calculate diversity indices (metabarcoding-derived 16S amplicon sequence variants – ASVs, metagenome-derived rpS3 genes, and metagenome-derived 16S rRNA gene reads mapped to a reference tree using phyloFlash – pF16S). The number (n) of used samples for each category is shown. Significance was tested using a Wilcoxon rank sum test, significance values are: *: p<0.05; **: p<0.01, ***: p<0.001, ****:p<0.0001. p values were corrected using the Bonferroni method. Note: Shannon and Simpson indices are relatively independent of the total number of ASVs analyzed. The presented values derive from an analysis of 2000 subsampled reads per sample but are basically indiscernible from an analysis using 50000 reads per sample (Fig. S8). In contrast, the number of observed ASVs (richness) does change with the number of subsampled reads analyzed, however the relative differences between groups are retained. Almost all pairs that were significantly different were between biomes, very few within biomes, corroborating the great differences between marine and terrestrial communities.

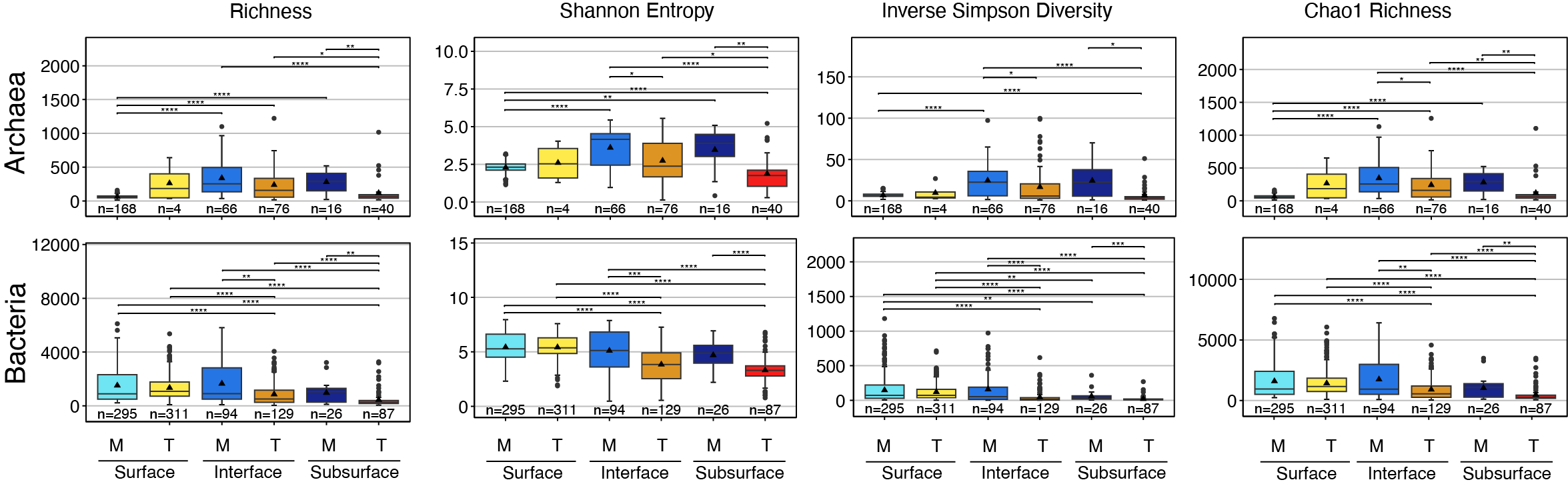

**Fig. S8: Alpha diversity indices of subsampled data**. To investigate the influence of subsampling on alpha diversity we used a very conservative cutoff using only samples that had at least 50000 archaeal or bacterial randomly chosen reads per sample. The difference in number of samples per group (n) to the ASV rows in Fig. S7 is due to samples that had less than 50000 archaeal or bacterial reads and were thus discarded. Overall trends in alpha diversity and significant differences between groups were very similar to the results obtained by analyzing 1142 archaeal and 2216 bacterial reads, the cutoff that was used for all other analyses.

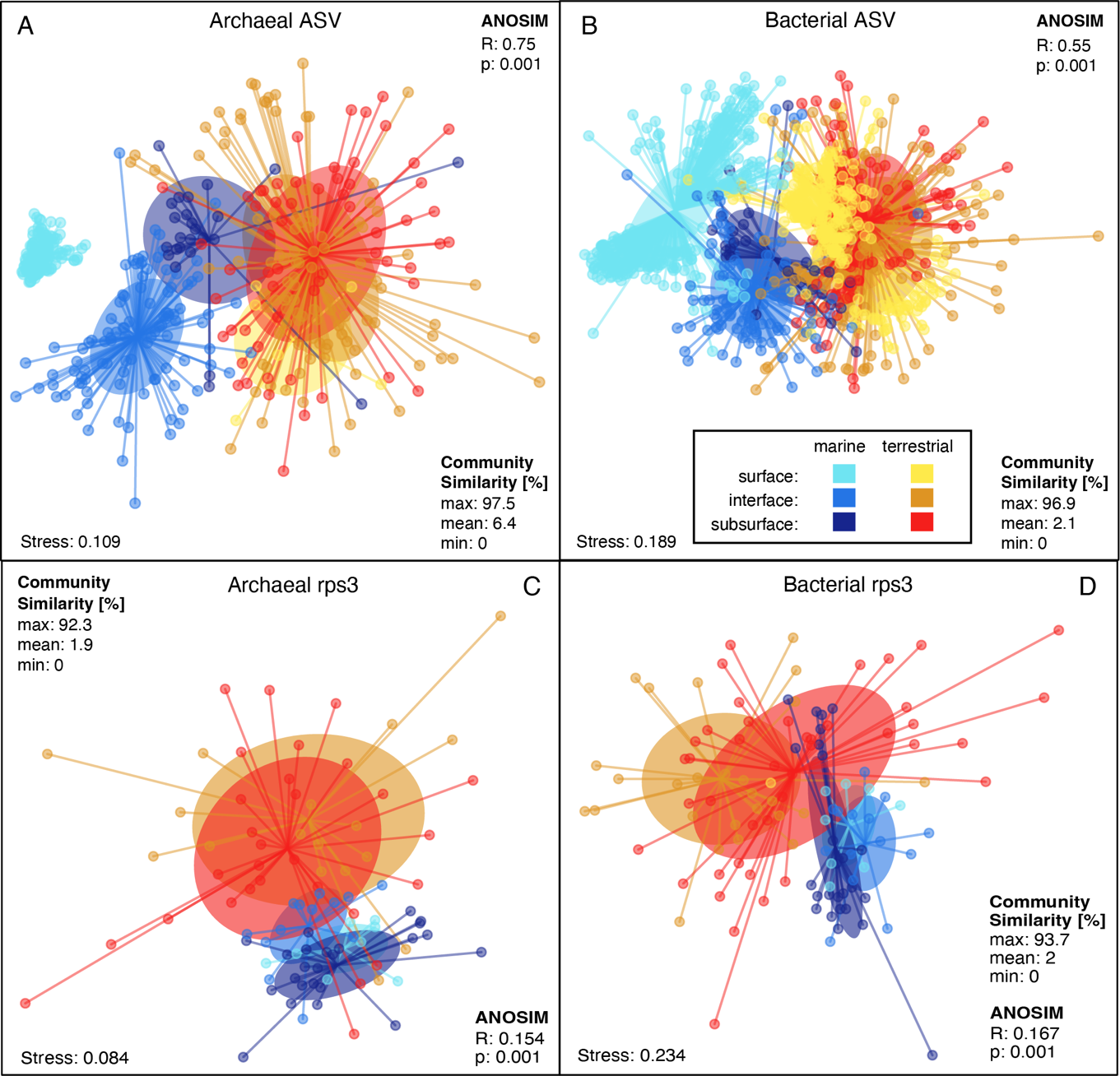

**Fig. S9: Microbial beta diversity**. Non-metric multidimensional scaling (NMDS) of archaeal (A, C) and bacterial (B, D) communities based on amplicon sequence variants (A, B) and rpS3 genes (C, D). Each dot represents the microbial community structure of a sample, colors represent six groups – three depths in both biomes. The plots show 469 archaeal amplicon datasets (37 projects, A), and 1105 bacterial amplicon datasets (51 projects, B), 91 archaeal rpS3 (C), and 117 bacterial rpS3 gene datasets (D). In A and B, the groups are overlapping, yet significantly different based on an Analysis of Similarity (ANOSIM). The separation between groups in A and B is easier to visualize in a 3-dimensional space, for this we have calculated and animated a 3D-NMDS (Video S1, S2) visualizing that each of the ecosystem types harbored distinct archaeal (ANOSIM: R=0.63; p=0.001) and bacterial communities (ANOSIM: R=0.54; p=0.001). In C and D, the groups are overall not significantly different, however the clear separation of the centroids, *i.e.* the average within-group distances, and ellipses (depicting 1 standard deviation from the centroid) corroborate the ASV-based trend. Community differences are particularly clear in marine vs terrestrial realms, *e.g.*, the blue (marine interface) and orange (terrestrial interface) ellipses, or the dark blue (marine subsurface) and red (terrestrial subsurface) ellipses hardly overlap, suggesting very distinct communities in these realms.

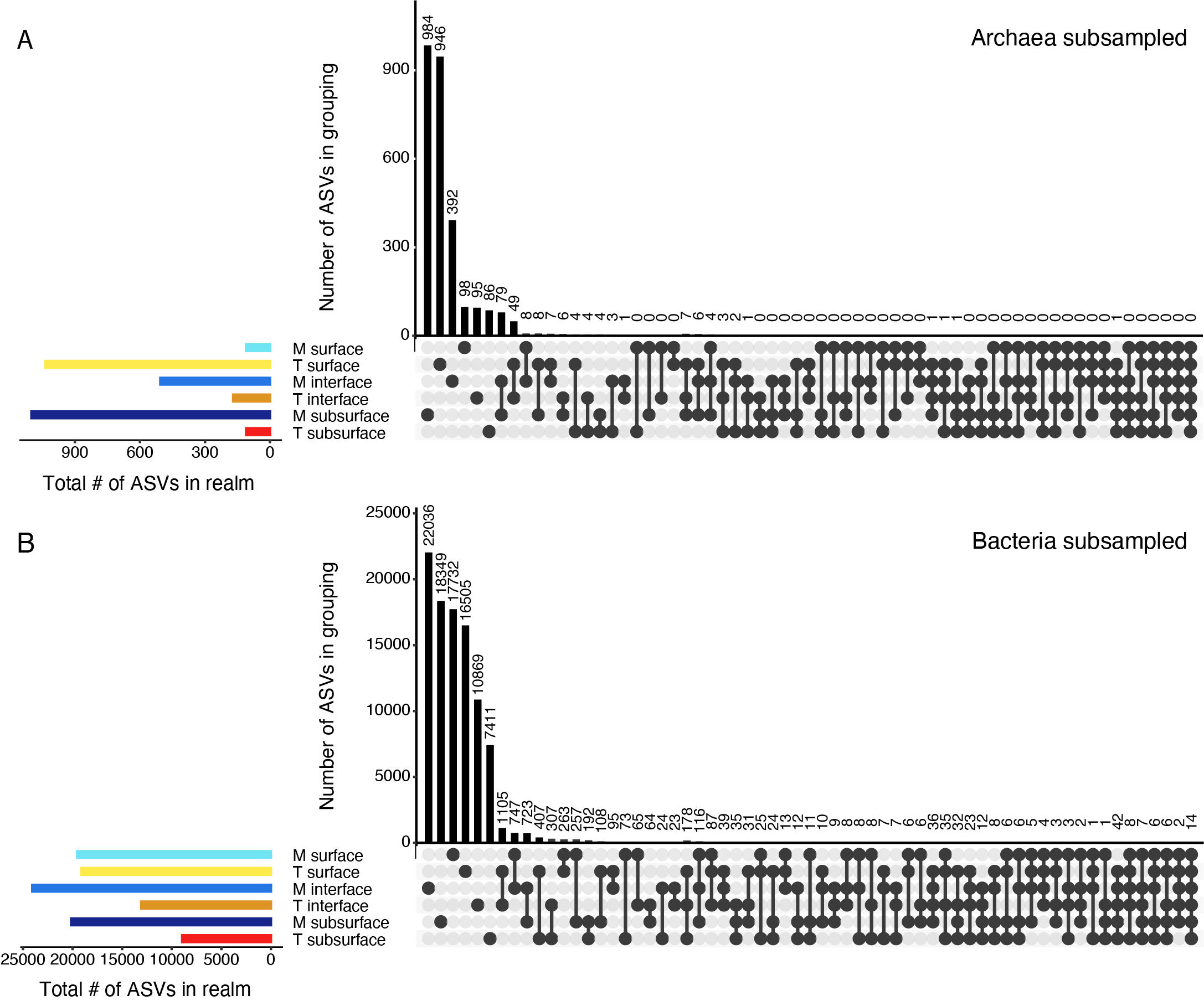

**Fig. S10: Community overlaps between analyzed biomes and depth realms**. Upset plots show subsampled archaeal (A) and bacterial communities (B). Subsampling was done to account for different numbers of samples per group and thus resulting unequal sampling effort. The plots depict the same information as Venn diagrams and show how many amplicon sequence variants (ASV) exclusively occur in a group or grouping (vertical bars). The total number of ASV of a group (gamma diversity) is shown as horizontal bars. For example, the highest number of total archaeal ASV in the subsampled dataset is found in marine subsurface environments. This group contains more than 1000 archaeal ASVs in total (gamma diversity, dark blue horizontal bar in A) of which 984 archaeal ASV occur exclusively in this group (1st vertical bar in A). Another example, there are 1105 bacterial ASV that occur exclusively in terrestrial surface AND interface datasets, but nowhere else (7^th^ vertical bar in B), this is depicted by the two connected black dots in the 2^nd^ and 4^th^ row. Archaeal and bacterial richness is decreasing with depth in terrestrial environments, but in marine environments richness is similar across depth for bacteria and even increasing with depth for archaea. Corroborating the trends seen with alpha diversity indices, and NMDS plots, it is clear from the upset plots that more ASVs are shared between realms of the same biome than between biomes.

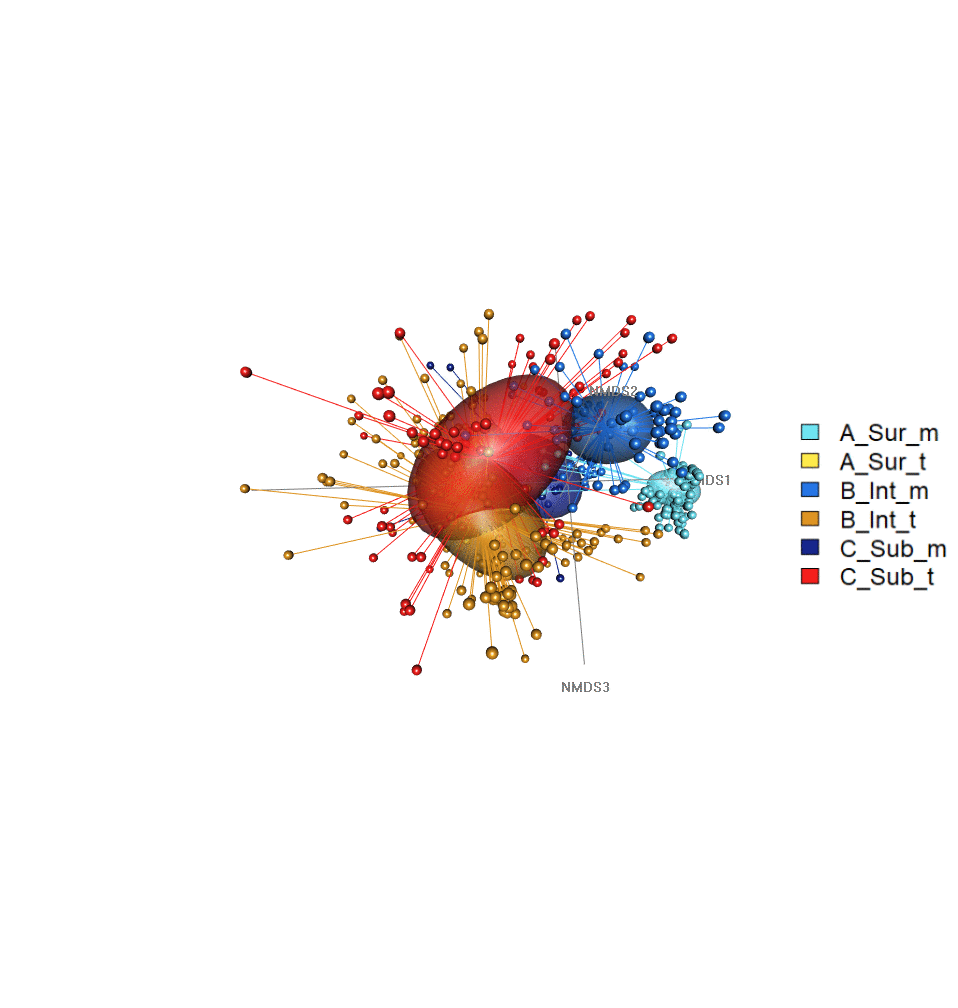

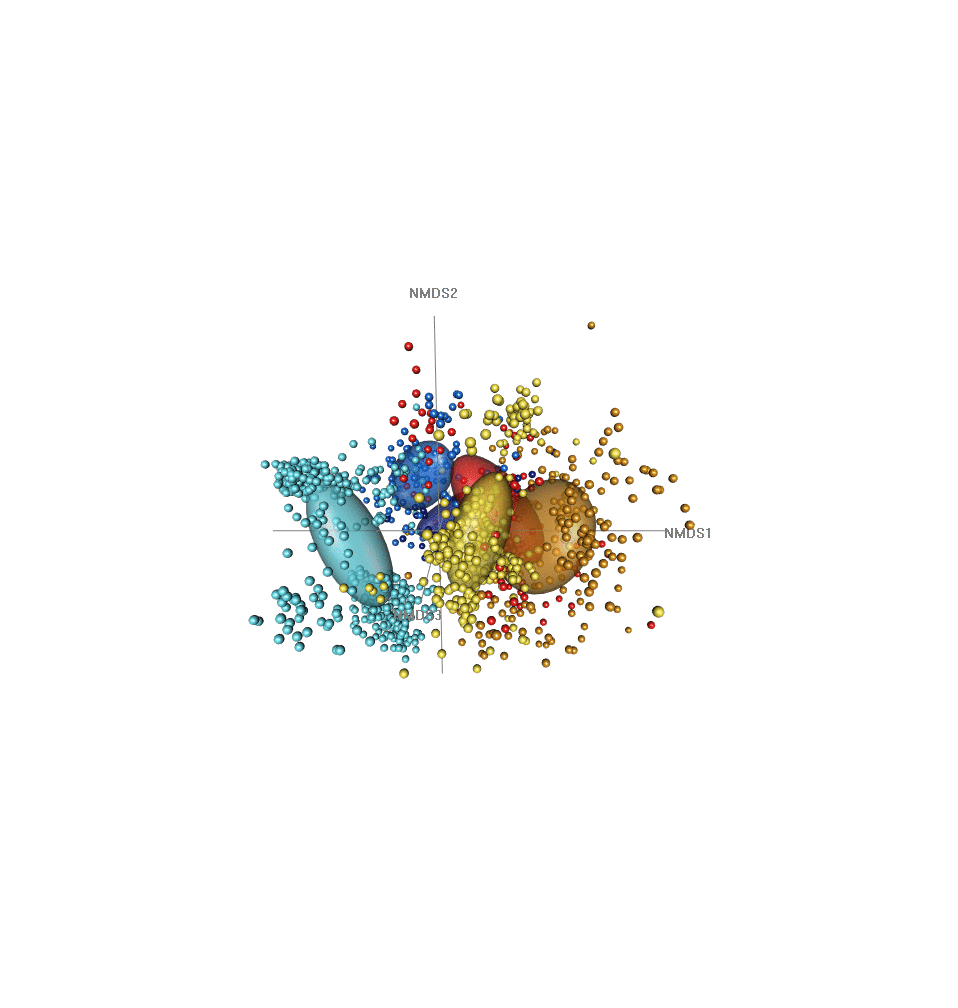

Bacteria

Archaea

**Movie S1, S2: Video-animated three-dimensional non-metric multidimensional scaling**. Ordination of archaeal (Video S1) and bacterial community structure (Video S2) based on 16S rRNA gene amplicon sequence variants (ASVs). Each dot is one sample, dot size represents Shannon entropy. The spheres depict the volume enclosing one standard deviation from the weighted average mean of within-group distances (centroid). Each sample is connected to the centroid of the group it belongs to.

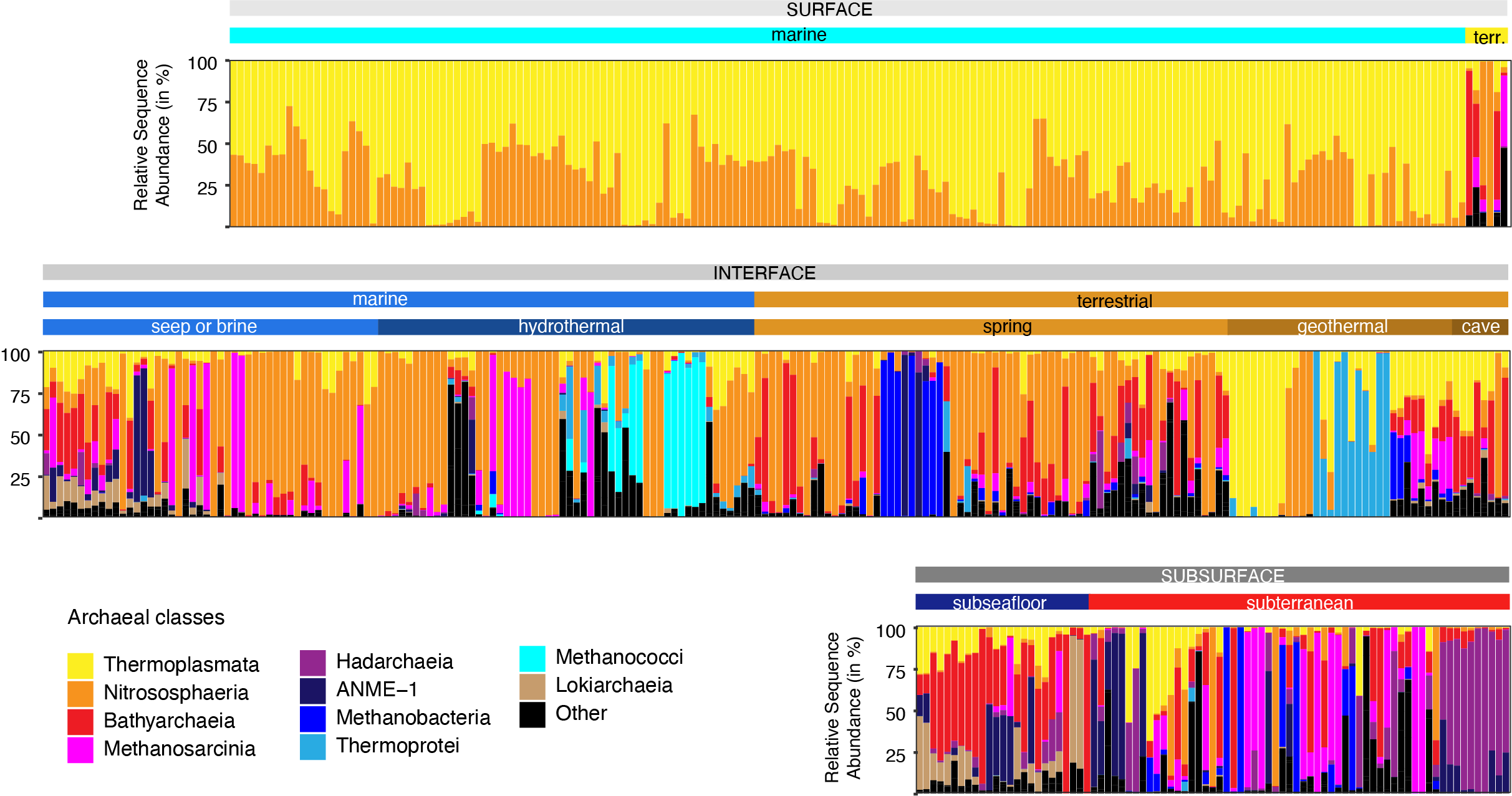

**Fig. S11: Archaeal class-level composition across all samples.** Each bar represents one sample and shows the relative sequence abundance of the top 10 most abundant archaeal classes. All remaining classes are summed and shown as “Other”. Relative abundances are based on the subsampled ASV × Sample table that was used for all community analyses. The samples are grouped by depth, biome, and environments (vertical bars above) and ordered analogous to Fig. 3.

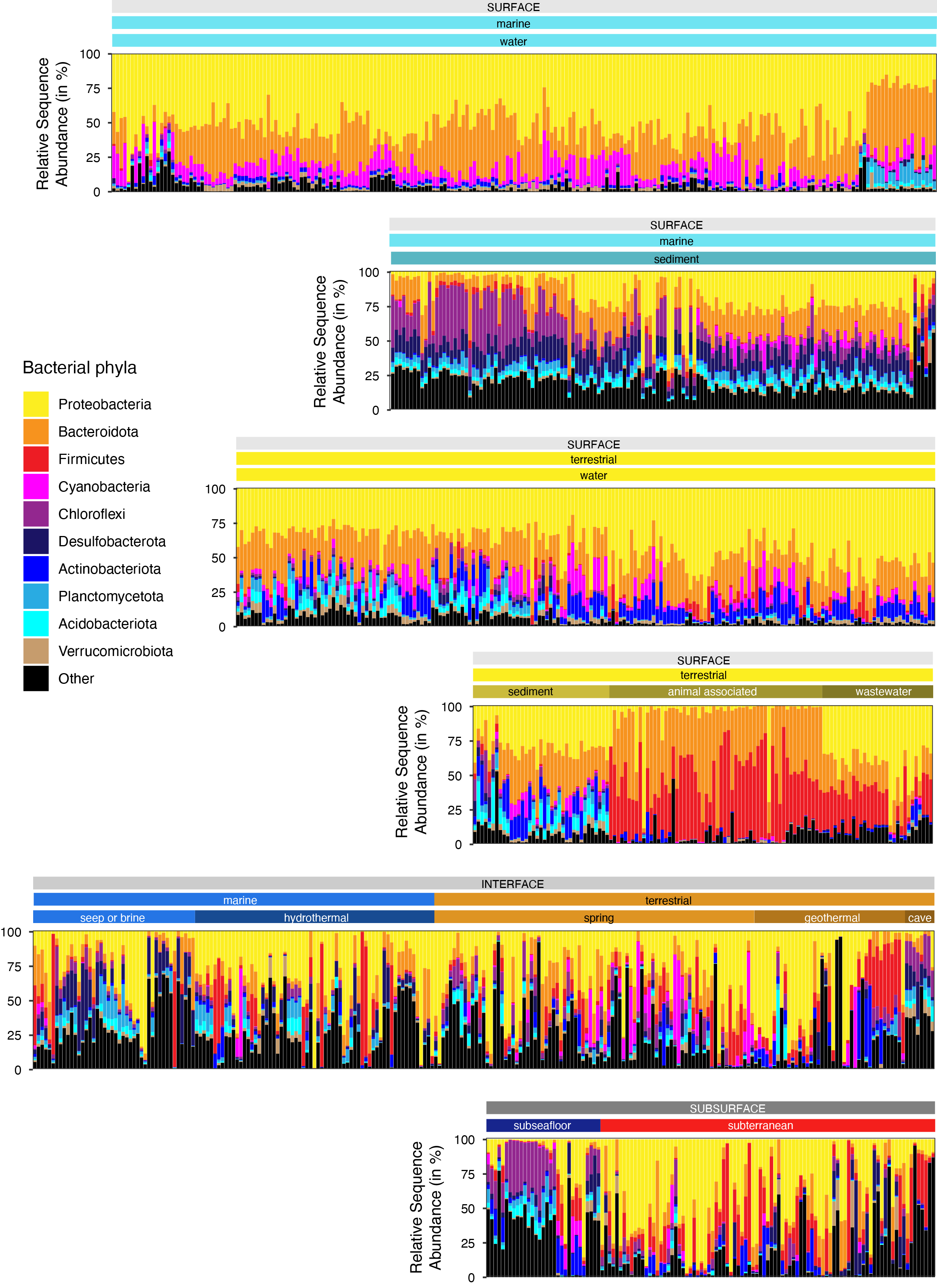

**Fig. S12: Bacterial phylum-level composition across all samples.** Each bar represents one sample and shows the relative sequence abundance of the top 10 most abundant bacterial phyla. All remaining phyla are summed and shown as “Other”. Relative abundances are based on the subsampled ASV × Sample table that was used for all community analyses. The samples are grouped by depth, biome, and environments (vertical bars above) and ordered analogous to Fig. 3.

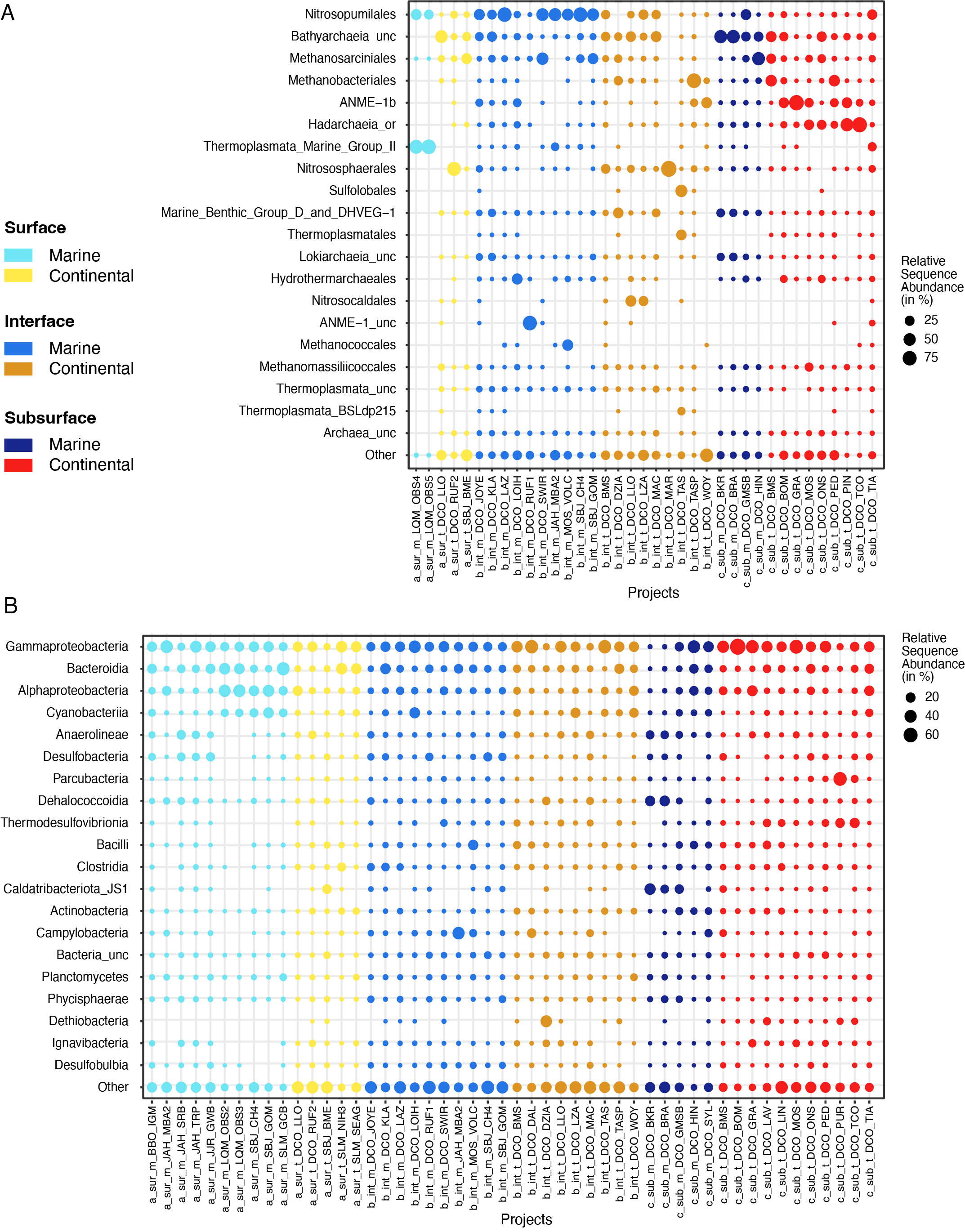

**Fig. S13: Relative sequence abundance.** Top 20 order-level archaeal clades (A) and class-level bacterial clades (B), averaged across samples for each project.

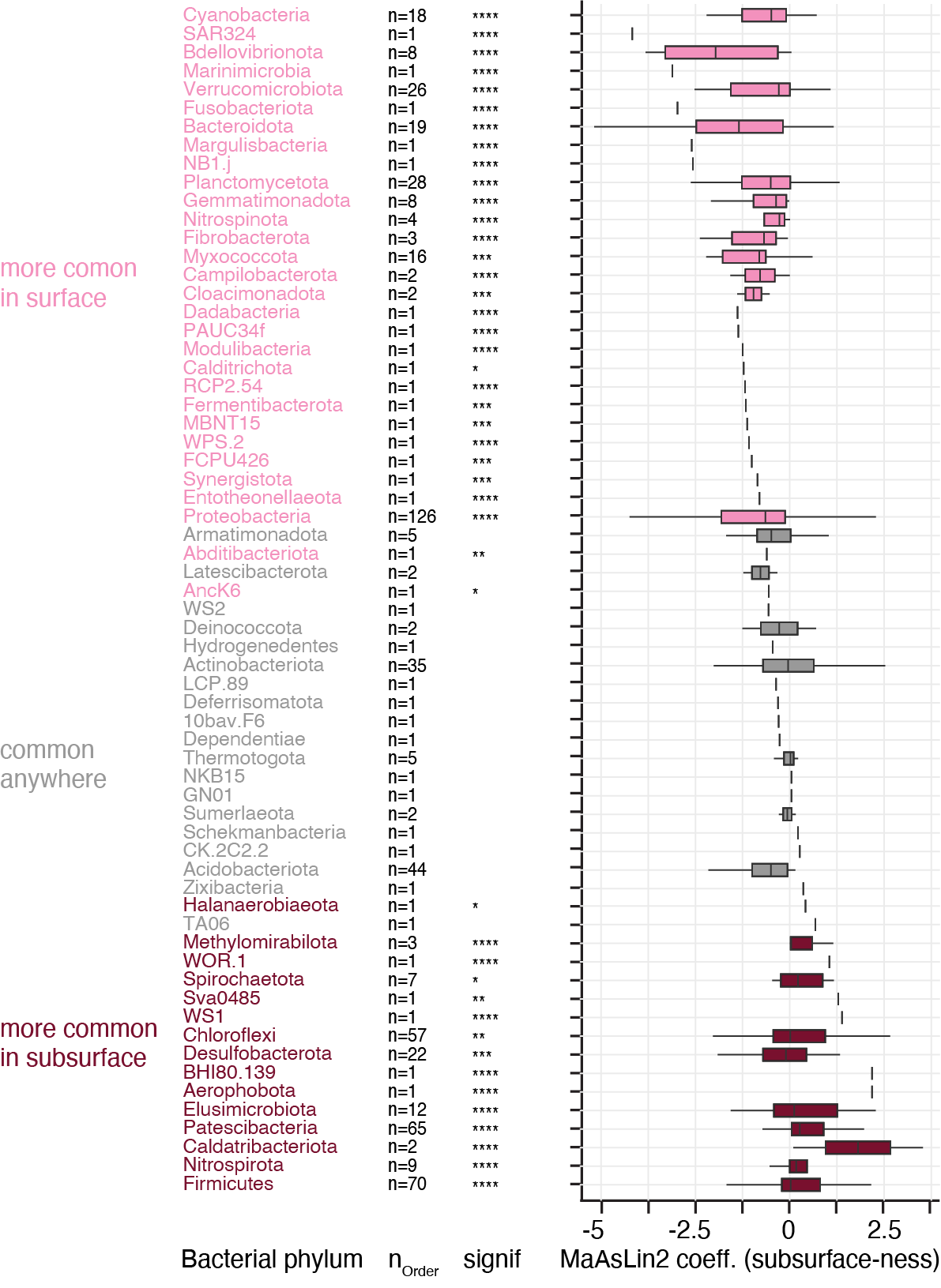

**Fig. S14: Differential abundance analyses.** Analyses comparing the occurrence of bacterial phyla in subsurface vs surface realms. The phyla are ordered from top to bottom based on increasing phylum level MaAsLin2 coefficient, *i.e.* likeliness of their occurrence in subsurface-derived samples (“subsurface-ness”). Boxplots summarize MaAsLin2 coefficients, i. e., “subsurface-ness”, of orders withing the listed phyla. Note: due to ease of visualization boxplots are even shown for very small number (n) of orders. The significance of this occurrence is shown in the column denoted “signif”. Significance levels are: *: p<0.05, **: p<0.01, ***: p<0.001, ****: p<0.0001. Phyla in in pink are found significantly more often in the surface realm, while phyla in maroon are found significantly less often in the surface realm, *i.e.*, occur more often in the subsurface.

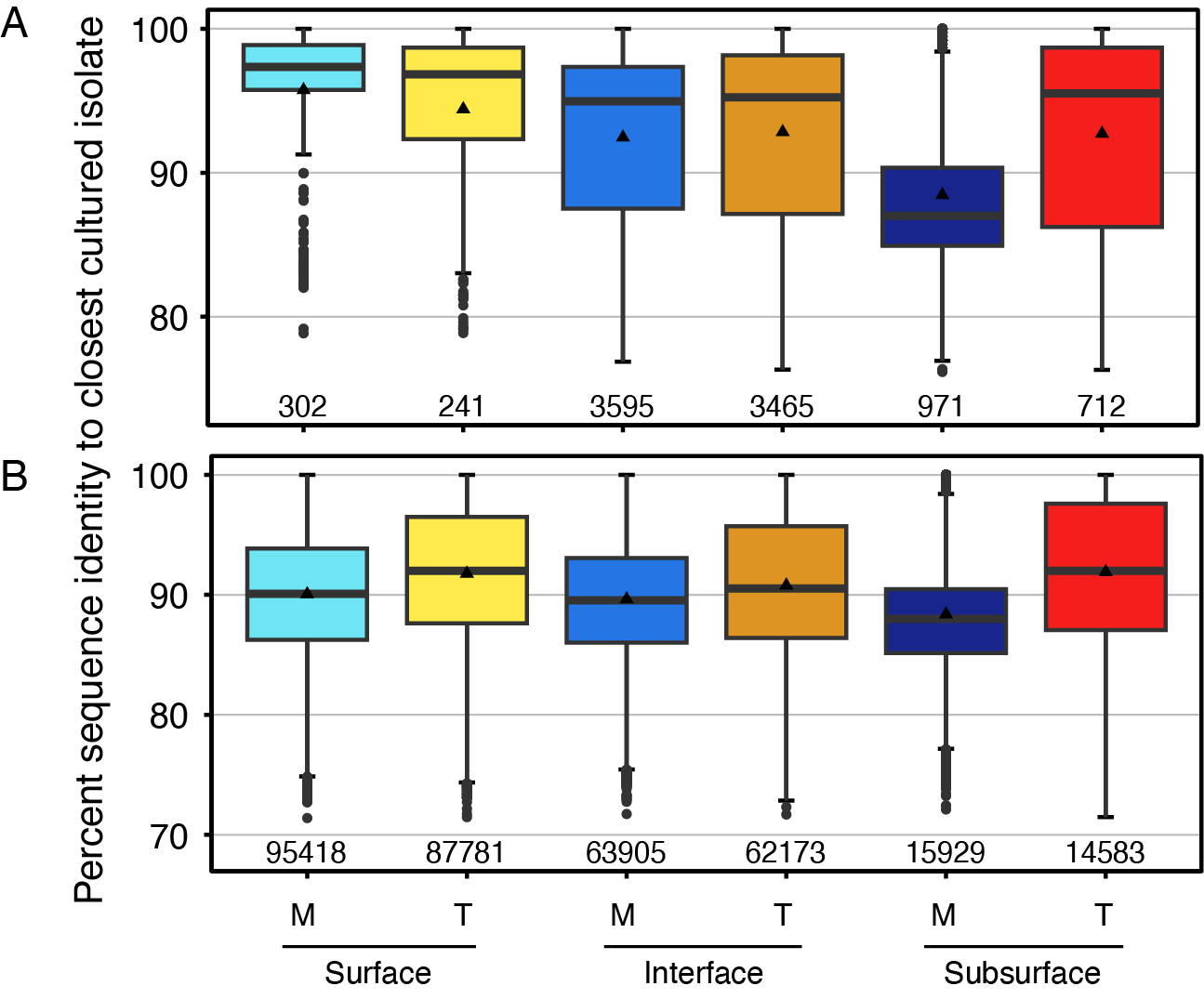

**Fig. S15: Phylogenetic novelty.** Summary of the percent identity values of n (below each boxplot) archaeal (A) and bacterial (B) amplicon sequence variants (ASVs) for the six studied realms. To determine the percent identity, each 16S rRNA gene amplicon was aligned to a database of cultured isolates. Every pairwise comparison was highly significant using a Wilcoxon rank sum test (p<0.01), except marine and terrestrial surface (archaea) and terrestrial interface and subsurface groups (archaea), significances not shown.

*Marine surface conditions may be imprinted in subsurface sediments*

We show differences in the archaeal and bacterial communities in marine subseafloor shelf, slope, and abyssal sedimentary environments (Fig. S16), consistent with the differences in cell-specific energy utilization across these different depositional settings (*158*). The differences in archaeal and bacterial richness and evenness between abyssal, slope and shelf subsurface sediments (Fig. S16A, C) are corroborated by differences in the community structure at respective environments (Fig. S16B, D). These findings support earlier studies that show similarities of surface and subsurface sedimentary communities suggesting that subsurface communities are shaped and imprinted by the shallow seafloor communities from which they are derived from (*23*–*25*). However, as the samples are not all from the same margin, and as the number of samples in each category is rather low the trend should be interpreted with caution.

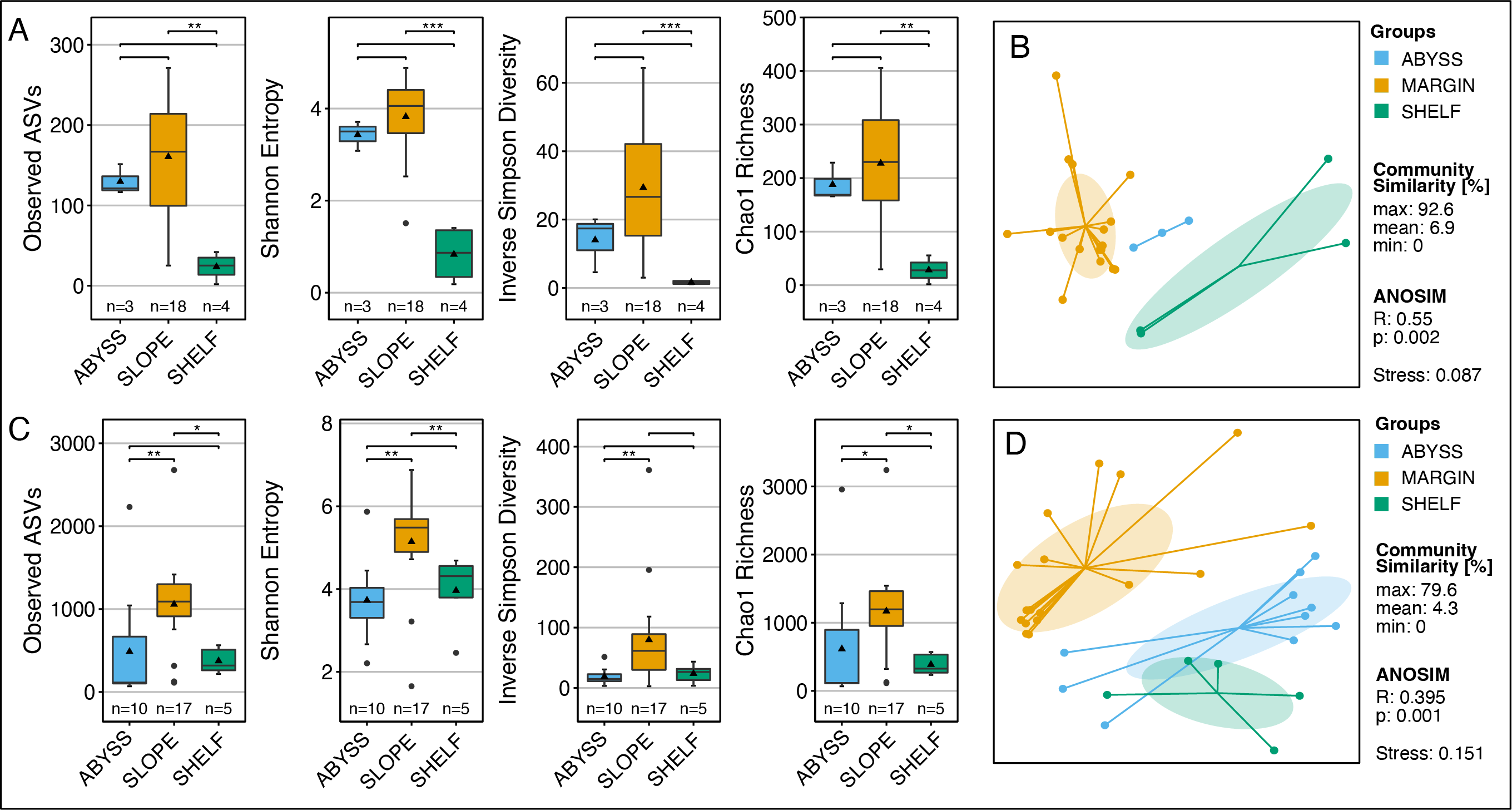

**Fig. S16: Microbial diversity in subsurface sediments from different depositional environments**. Archaeal (A, B) and bacterial (C, D) alpha diversity and beta diversity are significantly different between shelf and slope subsurface sediments. Significances were calculated with a Wilcoxon Rank sum test. Levels are: *: p<0.05, **: p<0.01, ***: p<0.001. Note: The number of archaeal datasets for abyssal and shelf environments is low providing limited statistical support.
