## Supplementary material for "A global atlas of subsurface microbiomes reveals phylogenetic novelty, large scale biodiversity gradients, and a marine-terrestrial divide": Dataset S3 - DADA2 R Script: DatasetS3_DADA2_v3_script.R

```
basic.packages <- c("package:stats","package:graphics","package:grDevices","package:utils","package:datasets","package:methods","package:base")
package.list <- search()[ifelse(unlist(gregexpr("package:",search()))==1,TRUE,FALSE)]
package.list <- setdiff(package.list,basic.packages)
if (length(package.list)>0)  for (package in package.list) detach(package, character.only=TRUE)
rm(list=ls()); closeAllConnections(); gc()

############ DADA2 - Exemplary workflow ###############################

# # Q1: Do your sequences derive from different runs? If yes -> Create sequence tables separately and merge afterwards.
# # Q2: Do your sequences contain Adapters/Primer? If yes -> truncate sequences accordingly -> change trim.lefty (used as trimLeft in dada2::filterAndTrim())
# # Q3: How much do your runs overlap? How much can you truncate at the end of each run? -> consider to change trim.righty (used as truncLen in dada2::filterAndTrim())
# # Q4: Do you have single- or paired end reads? Set PairedReads accoridngly
# # Q5: How big is your dataset. To reduce computational cost and analysis time, consider setting workflow to "bigdata" instead of "smalldata". This might introduce some loss of resolution, but decrease the memory required and computational cost (https://benjjneb.github.io/dada2/bigdata.html)

# # This workflow uses dada2; please cite 
# # Callahan BJ, McMurdie PJ, Rosen MJ, Han AW, Johnson AJA, Holmes SP (2016). "DADA2: High-resolution sample inference from Illumina amplicon data." _Nature Methods_, *13*, 581-583. doi: 10.1038/nmeth.3869 (URL: https://doi.org/10.1038/nmeth.3869)
# # DADA2 workflow using sample inference for small data (computational intensive for big datasets). This workflow uses unzipped and zipped forward and reverse reads (or only forward or only reverse).
# # For help check out https://benjjneb.github.io/dada2/index.html

# # dada2 installation instructions: https://benjjneb.github.io/dada2/dada-installation.html
# if (!requireNamespace("BiocManager", quietly = TRUE))
#   install.packages("BiocManager")
# BiocManager::install("dada2", version = "3.17")
# BiocManager::install("DECIPHER")

if (!require('dada2')) {install.packages('dada2')}; library(dada2)
if (!require('DECIPHER')) {install.packages('DECIPHER')}; library(DECIPHER)
if (!require('data.table')) {install.packages('data.table')}; library(data.table)

source("C:/Users/i.hrabedeangelis/Dropbox/scripts/R/aafunctions.R")
# source("X:/EugenSeibold/Topics/a04/rp_2/v01/aafunctions.R")
getN <- function(x) sum(getUniques(x))

# # ==== User Input ==== 
# # In addition to the input provided here, you have to set trim.righty to truncate your reads after assessing the quality profiles
workflow <- 'smalldata' # set to 'bigdata' or 'smalldata' depending on your dataset and your research question
PairedReads <- 'paired'                          # set to 'single' or 'paired'
ProjectName <- 'dada2test'                          # Give your project a name.

# # # calculate length of overlap:
FrwdPrmrStart <- 515  # start of your forward primer, e.g. 515 for 515F
RvrsPrmrStart <- 909  # start of your reverse primer, e.g. 909 for 909R
ReadLength <- 301     # length of your reads

if(PairedReads == "paired") {
  message(paste("Your reads have an overlap of",(ReadLength-((RvrsPrmrStart-FrwdPrmrStart+1)/2))*2,"bases (not accounting for biological length variation)."))
} else if (PairedReads == "single") {
  message("You are using single reads.")
}

PathToDadaInputFiles=file.path("C:/path/to/sequencing/data")                                      # Select the folder where dada can find your sequences (fastq files)
PathToDadaOutputFiles=file.path("C:/path/to/folder/where/results/should/be/saved",paste(ProjectName,"_dada2",sep=''))  # Select a folder where dada should save your analysis. Inside this folder a folder with the name of your analysis and _dada2 attached to it will be created
TaxonomyFolder=file.path("X:/EugenSeibold/Topics/a04/rp_2/v01/DatabasesForDada2/dada")                                            # Select the folder where you have saved the SILVA taxonomy files 
classmeth <- "dada2" # "dada2" use dada2 algorithm (naive Bayesian classifier) or "dec" for the decipher algorithm. Be aware that depending on your choice you'll need different databases
database="silva138" 

# which database you want to use 
# for dada2 classifying: 
# database = "silva1381" # for silva_nr_v138.1 # optimized for bac and arc, not for eukaryotes
# database = "silva138" # for silva_nr_v138
# database = "silva132" # for silva_nr_v132 # use this one for eukaryotes
# database = "rdp18" # for RDP_18
# database = "pr2" # for PR2
# database = "unite" # for unite

# for decipher classifiying: 
# database = "silva138" # for SILVA_SSU_r138_2019
# database = "rdp18" # for RDP_v18
# database = "pr2v4" # for PR2_v4
# database = "unite2021" # for UNITE_v2021_May2021

# # If you only have forward or reverse reads leave the other field empty
PatternForwardReads="_R1_001.fastq" # Change to the Pattern at the end of your forward reads, if e.g. SAMPLENAME_PATTERN.fastq, provide '_PATTERN.fastq'
PatternReverseReads="_R2_001.fastq" # Change to the Pattern at the end of your reverse reads

MultiThread=F               # Use MultiThread=F (Windows) or MultiThread=TRUE

trim.lefty <- c(17,17)      # If you need to cut off primers or adapters from your reads. the first number is what will be cut off at the start of the forward reads, the second number is what will be cut off at the start of the reverse reads. if you have single reads (e.g. only forward or only reverse) then only use one number

maxerror <- c(2,2) # for maxEE in dada2::filterAndTrim(). Sets the maximum number of "expected errors" allowed in a read. Default c(2,2) for merged reads and c(2) for single reads. " If you want to speed up downstream computation, consider tightening maxEE. If too few reads are passing the filter, consider relaxing maxEE, perhaps especially on the reverse reads (eg. maxEE=c(2,5)), and reducing the truncLen to remove low quality tails. Remember though, when choosing truncLen for paired-end reads you must maintain overlap after truncation in order to merge them later." (dada2 homepage) 

qualityscore <- 2 # for truncQ in dada2::filterAndTrim(). Sets the quality score. truncQ=2 will truncate the read at the first nucleotide with a quality score of 2 (if there is one). Read will only  thrown away if it is then shorter than truncLen

reversecomplement=F         # for tryRC in taxonomy assignment. Set T if you want to try if the reverse complement better fits. Might be necessary for reverse reads.

Pool=F                      # Set T, F, or "pseudo". Pooling can increase the sensitivity to rare per-sample variants. Needs more memory and computation time. Pseudo-pooling approximates pooling with less time requirement.

if(PairedReads=="paired"){
  message(paste("Your merged reads will have a length of",(RvrsPrmrStart-FrwdPrmrStart-sum(trim.lefty)+1) ,"bases."))
} else if (PairedReads=="single") {
  message(paste("You're reads will have a length of",ReadLength-trim.lefty,"bases."))
}

# # ==== Create analysis directories and log file ====

dir.create(PathToDadaOutputFiles)
dir.create(file.path(PathToDadaOutputFiles,"plots"))
dir.create(file.path(PathToDadaOutputFiles,"filtered"))

workspaceName=paste(ProjectName,"_dada2",".RData",sep="")

cat('dada2 Analysis\n',file=(file.path(PathToDadaOutputFiles,paste(ProjectName,'_SessioInfo.txt',sep=''))),append=TRUE)
sink(file=(file.path(PathToDadaOutputFiles,paste(ProjectName,'_SessioInfo.txt',sep=''))),append=TRUE)
cat('dada2 ProjectName: ',ProjectName,'\n',sep='')
print(paste('AnalysisDate: ',Sys.Date()))

sink(stdout(),type='message')
sessionInfo()
closeAllConnections()


cat('dada2 Analysis\n',file=(file.path(PathToDadaOutputFiles,paste(ProjectName,'_Analysis_log.txt',sep=''))),sep='',append=TRUE)
sink(file=(file.path(PathToDadaOutputFiles,paste(ProjectName,'_Analysis_log.txt',sep=''))),append=TRUE,type='output')
print(paste('AnalysisDate: ',Sys.Date()))
cat('dada2 ProjectName: ',ProjectName,'\n\n',sep='')

cat('Information on input sequences','\n',sep='')
cat('You are analysing ',PairedReads,' reads.','\n',sep='')
cat('The sequencing data was read in from the folder ',PathToDadaInputFiles,'.','\n',sep='')
cat('Your forward primer starts at base ',FrwdPrmrStart,'.','\n',sep='')
if(PairedReads == "paired") {
  cat('Your reverse primer starts at base ',RvrsPrmrStart,'.','\n',sep='')
}
cat('Your reads have a length of ',ReadLength,' bases.','\n',sep='')
if(PairedReads =="paired") {
  cat('Your untruncated reads have an overlap (without accounting for biological length variation) of ',(ReadLength-((RvrsPrmrStart-FrwdPrmrStart+1)/2))*2,' bases.','\n\n',sep='')
}
sink()


# # ==== Reads fastq's ==== 

list.files(PathToDadaInputFiles) # here, all your fastq files should be listed

if (PairedReads=='paired') {
  fnFs <- sort(list.files(PathToDadaInputFiles, pattern=PatternForwardReads, full.names = TRUE))  # creates a list containing all your forward reads
  fnRs <- sort(list.files(PathToDadaInputFiles, pattern=PatternReverseReads, full.names = TRUE))
  if(length(fnFs) != length(fnRs)) stop("Forward and reverse files do not match.")
  sample.names <- sapply(strsplit(basename(fnFs), PatternForwardReads), `[`, 1)                   # Extracts sample names
  sample.names <- sapply(strsplit(basename(fnRs), PatternReverseReads), `[`, 1)
} else if (PairedReads=='single') {
  fnFs <- sort(list.files(PathToDadaInputFiles, pattern=PatternForwardReads, full.names = TRUE))
  sample.names <- sapply(strsplit(basename(fnFs), PatternForwardReads), `[`, 1)
}

# # ==== Plots and Saves Quality profiles ==== 

if (PairedReads=='paired') {
  if(length(fnFs) > 16) {
    pdf(file.path(PathToDadaOutputFiles,"plots",paste(ProjectName,"_Quality_Forward_16.pdf",sep="")),width=10,height=8,useDingbats=FALSE)
    print(plotQualityProfile(fnFs[1:16]))
    dev.off()
    pdf(file.path(PathToDadaOutputFiles,"plots",paste(ProjectName,"_Quality_Reverse_16.pdf",sep="")),width=10,height=8,useDingbats=FALSE)
    print(plotQualityProfile(fnRs[1:16]))
    dev.off()  
  }
  pdf(file.path(PathToDadaOutputFiles,"plots",paste(ProjectName,"_Quality_Forward_all.pdf",sep="")),width=20,height=20,useDingbats=FALSE)
  print(plotQualityProfile(fnFs))
  dev.off()
  pdf(file.path(PathToDadaOutputFiles,"plots",paste(ProjectName,"_Quality_Reverse_all.pdf",sep="")),width=20,height=20,useDingbats=FALSE)
  print(plotQualityProfile(fnRs))
  dev.off()
} else if (PairedReads=='single') {
  if(length(fnFs) > 16) {
    pdf(file.path(PathToDadaOutputFiles,"plots",paste(ProjectName,"_Quality_Forward_16.pdf",sep="")),width=10,height=8,useDingbats=FALSE)
    print(plotQualityProfile(fnFs[1:16]))
    dev.off()
  }
  pdf(file.path(PathToDadaOutputFiles,"plots",paste(ProjectName,"_Quality_Forward_all.pdf",sep="")),width=20,height=20,useDingbats=FALSE)
  print(plotQualityProfile(fnFs))
  dev.off()
}
gc()

# # ==== INPUT NEEDED - truncLen and pontentially add primer info ==== 
# # Your reads should at least have an overlap of 20 + biological length variation nucleotides.
# # the v3v4 region is bimodal with a peak at ~440 and one of ~460 nucleotides. (https://github.com/benjjneb/dada2/issues/857)

# removePrimers(fnRs,filtRs,'GGACTACHVGGGTWTCTAAT',orient = T)
stop("Set your truncation length.")

trim.righty <- c(250,200)   # First number gives the length of the forward reads you want to keep (after which base the sequence will be discarded), the second number is the same for the reverse reads. Keep in mind that you will still need to have overlap! For single reads only use one number.

if(PairedReads == "paired") {
  message(paste("If you cut your reads like that you'll have",as.character((FrwdPrmrStart+trim.righty[1]-1)-(RvrsPrmrStart-trim.righty[2]+1)+1),"bases overlap."))
  message(paste("The merged reads will have a length of",as.character(RvrsPrmrStart-trim.lefty[2]-(FrwdPrmrStart+trim.lefty[1])+1),"bases (not accounting for biological length variation)."))
  if(((FrwdPrmrStart+trim.righty[1]-1)-(RvrsPrmrStart-trim.righty[2]+1)+1) < 0) {
    stop("You truncated your reads too much, you're paired reads do not overlap anymore. Change trim.righty() acordingly")
  }
  if(((FrwdPrmrStart+trim.righty[1]-1)-(RvrsPrmrStart-trim.righty[2]+1)+1) < 30) {
    warning("You're reads have an overlap of ",((FrwdPrmrStart+trim.righty[1]-1)-(RvrsPrmrStart-trim.righty[2]+1)+1),". This might be too short to account for the recommended overlap of 20 bases plus the biological length variation of nucleotides.")
  }
} else if(PairedReads=="single") {
  message(paste("If you cut your reads like that they'll have a length of",as.character(ReadLength-trim.lefty-(ReadLength-trim.righty)),"bases."))
}


# # ==== Logfile ==== 
sink(file=(file.path(PathToDadaOutputFiles,paste(ProjectName,'_Analysis_log.txt',sep=''))),append=TRUE,type='output')
cat('The following parameters were used for dada2::filterandTrim().','\n',sep='')
if(PairedReads == 'paired') {
  cat('truncLen: ',trim.righty[1],',',trim.righty[2],'\n',sep='')
  cat('This truncation length results in a final overlap of ',as.character((FrwdPrmrStart+trim.righty[1]-1)-(RvrsPrmrStart-trim.righty[2]+1)+1),' bases.\n',sep='')
  cat('maxEE: ',maxerror[1],',',maxerror[2],'\n',sep='')
  cat('trimLeft: ',trim.lefty[1],',',trim.lefty[2],'\n',sep='')
} else {
  cat('truncLen: ',trim.righty[1],'\n',sep='')
  cat('This truncation length results in a final read length of ',as.character(trim.righty[1]-trim.lefty[1]),' bases.\n',sep='')
  cat('maxEE: ',maxerror[1],'\n',sep='')
  cat('trimLeft: ',trim.lefty[1],'\n',sep='')
  
}
cat('truncQ: ',qualityscore,'\n\n',sep='')

cat('The following parameters were used for dada2::dada().','\n',sep='')
cat('pool: ',Pool,'\n',sep='')
if(workflow == 'smalldata') {
  cat('Sample inference was calculated for all input data at once (small data protocol). \n\n',sep='')
} else if (workflow == 'bigdata') {
  cat('Sample inference was calculated using the big data protocol. \n\n',sep='')
}

cat('The following algorithm, database and parameters were used for dada2::assignTaxonomy() and dada2::addSpecies().','\n',sep='')
cat('algorithm: ',classmeth,'\n',sep='')
cat('database: ',database,'\n',sep='')
cat('tryRC: ',reversecomplement,'\n',sep='')
sink()

# # ==== dada2::filterAndTrim ==== 
# Considerations for your own data: The standard filtering parameters are starting points, not set in stone. 
# If you want to speed up downstream computation, consider tightening maxEE. 
# If too few reads are passing the filter, consider relaxing maxEE, perhaps especially on the reverse reads (eg. maxEE=c(2,5)), and reducing the truncLen to remove low quality tails. 
# truncLen cuts the reads off after the given base position (truncLen=c(190,235) cuts off the forward reads after the 190th base and the reverse reads after the 235th base)
# Remember though, when choosing truncLen for paired-end reads you must maintain overlap after truncation in order to merge them later. 

if (PairedReads=='paired') {
  filtFs <- file.path(PathToDadaOutputFiles, "filtered", paste0(sample.names, "_F_filt.fastq.gz")) # Assigns the filenames for the filtered fastq.gz files
  filtRs <- file.path(PathToDadaOutputFiles, "filtered", paste0(sample.names, "_R_filt.fastq.gz"))
  names(filtFs) <- sample.names
  names(filtRs) <- sample.names
  out <- filterAndTrim(fnFs, filtFs, fnRs, filtRs, truncLen=trim.righty,                            # change the parameters according to your sequencees 
                       maxN=0, maxEE=maxerror, truncQ=qualityscore, rm.phix=TRUE,
                       compress=TRUE, multithread=MultiThread,trimLeft = trim.lefty, verbose = T)
} else if (PairedReads=='single') {
  filtFs <- file.path(PathToDadaOutputFiles, "filtered", paste0(sample.names, "_F_filt.fastq.gz")) # Assigns the filenames for the filtered fastq.gz files
  gc()
  names(filtFs) <- sample.names
  out <- filterAndTrim(fnFs, filtFs,  truncLen=trim.righty,
                       maxN=0, maxEE=maxerror, truncQ=qualityscore, rm.phix=TRUE,
                       compress=TRUE, multithread=MultiThread,trimLeft = trim.lefty, verbose = T)
}

gc()

out<-as.data.frame(out)
out$ratio <- out$reads.out/out$reads.in

head(out) # shows reads in and reads out for the first 6 samples

message('ratio filtered/input')
summary(out$ratio)

# # ==== dada2::learnErrors ==== 

if(workflow == 'smalldata') {
  if (PairedReads=='paired') {
    errF <- learnErrors(filtFs, multithread=MultiThread)
    errR <- learnErrors(filtRs, multithread=MultiThread)
    pdf(file.path(PathToDadaOutputFiles,"plots",paste(ProjectName,"_Error_Forward.pdf",sep="")),width=8,height=8,useDingbats=FALSE)
    print(plotErrors(errF, nominalQ=TRUE))
    dev.off()
    pdf(file.path(PathToDadaOutputFiles,"plots",paste(ProjectName,"_Error_Reverse.pdf",sep="")),width=8,height=8,useDingbats=FALSE)
    print(plotErrors(errR, nominalQ=TRUE))
    dev.off()
    filtFs <- filtFs[file.exists(filtFs)] # in case some samples were excluded previously they will be filtered out here from the list
    filtRs <- filtRs[file.exists(filtRs)]
  } else if (PairedReads=='single') {
    errF <- learnErrors(filtFs, multithread=MultiThread)
    pdf(file.path(PathToDadaOutputFiles,"plots",paste(ProjectName,"_Error_Forward.pdf",sep="")),width=8,height=8,useDingbats=FALSE)
    print(plotErrors(errF, nominalQ=TRUE))
    dev.off()
    filtFs <- filtFs[file.exists(filtFs)] # in case some samples were excluded previously they will be filtered out here from the list
  }
  gc()
} else if (workflow == 'bigdata') {
  set.seed(100)
  if (PairedReads=='paired') {
    errF <- learnErrors(filtFs, nbases=1e8, multithread=MultiThread)
    errR <- learnErrors(filtRs, nbases=1e8, multithread=MultiThread)
    pdf(file.path(PathToDadaOutputFiles,"plots",paste(ProjectName,"_Error_Forward.pdf",sep="")),width=8,height=8,useDingbats=FALSE)
    print(plotErrors(errF, nominalQ=TRUE))
    dev.off()
    pdf(file.path(PathToDadaOutputFiles,"plots",paste(ProjectName,"_Error_Reverse.pdf",sep="")),width=8,height=8,useDingbats=FALSE)
    print(plotErrors(errR, nominalQ=TRUE))
    dev.off()
    filtFs <- filtFs[file.exists(filtFs)] # in case some samples were excluded previously they will be filtered out here from the list
    filtRs <- filtRs[file.exists(filtRs)]
  } else if (PairedReads=='single') {
    errF <- learnErrors(filtFs, nbases=1e8, multithread=MultiThread)
    pdf(file.path(PathToDadaOutputFiles,"plots",paste(ProjectName,"_Error_Forward.pdf",sep="")),width=8,height=8,useDingbats=FALSE)
    print(plotErrors(errF, nominalQ=TRUE))
    dev.off()
    filtFs <- filtFs[file.exists(filtFs)] # in case some samples were excluded previously they will be filtered out here from the list
  }
  gc()
}


# # ==== dada2::dada - Applies Sample Inference algorithm ==== 

if(workflow == 'smalldata') {
  if (PairedReads=='paired') {
    dadaFs <- dada(filtFs, err=errF, multithread=MultiThread,pool=Pool)
    gc()
    dadaRs <- dada(filtRs, err=errR, multithread=MultiThread,pool=Pool)
    gc()
    sapply(dadaFs, getN)/out$reads.out
    sapply(dadaRs, getN)/out$reads.out
    message('ratio denoisedF/filtered')
    print(summary(sapply(dadaFs, getN)/out$reads.out))
    message('ratio denoisedR/filtered')
    print(summary(sapply(dadaRs, getN)/out$reads.out))
  } else if (PairedReads=='single') {
    dadaFs <- dada(filtFs, err=errF, multithread=MultiThread,pool=Pool)
    sapply(dadaFs, getN)/out$reads.out
    message('ratio denoisedF/filtered')
    print(summary(sapply(dadaFs, getN)/out$reads.out))
  }
  save.image(file.path(PathToDadaOutputFiles,workspaceName))
  gc()
} else if(workflow == 'bigdata') {
  # Sample inference and merger of paired-end reads
  elementno=1
  if (PairedReads == "paired") {
    mergers <- vector("list", length(sample.names))
    names(mergers) <- sample.names
    for(sam in sample.names) {
      cat("Processing:", sam, "\n")
      derepF <- derepFastq(filtFs[[sam]])
      dadaFs <- dada(derepF, err=errF, multithread=MultiThread)
      derepR <- derepFastq(filtRs[[sam]])
      dadaRs <- dada(derepR, err=errR, multithread=MultiThread)
      merger <- mergePairs(dadaFs, derepF, dadaRs, derepR)
      mergers[[sam]] <- merger
      # add info to output overview
      out[elementno,4] <- sum(dadaFs$denoised)
      out[elementno,5] <- out[elementno,4]/out[elementno,2]
      out[elementno,6] <- sum(dadaRs$denoised)
      out[elementno,7] <- out[elementno,6]/out[elementno,2]
      out[elementno,8] <- sapply(mergers[elementno],getN)
      rm(derepF); rm(derepR); rm(merger)
      message('ratio denoisedF/filtered')
      print(summary(out$V4/out$reads.out))
      message('ratio denoisedR/filtered')
      print(summary(out$V6/out$reads.out))
      message('ratio merged/denoised')
      print(summary(sapply(mergers[elementno], getN)/((out$V4+out$V6)/2)))
      elementno <- elementno+1
      gc()
    }
  } else if (PairedReads == "single") {
    dadaFs <- vector("list", length(sample.names))
    names(dadaFs) <- sample.names
    for(sam in sample.names) {
      cat("Processing:", filtFs, "\n")
      derepF <- derepFastq(filtFs[[sam]])
      dadaFs[[sam]] <- dada(derepF, err=errF, multithread=MultiThread)
      # add info to output overview
      out[elementno,4] <- sum(dadaFs[[sam]][["denoised"]])
      # out[elementno,5] <- out[elementno,4]/out[elementno,2]
      elementno <- elementno+1
      rm(derepF)
      message('ratio denoisedF/filtered')
      print(summary(out$V4/out$reads.out))
      gc()
    }
  }
  rm(elementno); rm(sam)
  save.image(file.path(PathToDadaOutputFiles,workspaceName))
}


# # ==== Merges paired reads ==== 

if(workflow == 'smalldata') { # for big data, merging already happens while we go through the sample inference
  if (PairedReads=='paired') {
    mergers <- mergePairs(dadaFs, filtFs, dadaRs, filtRs, verbose=TRUE)
    head(mergers[[1]]) # Inspect the merger data.frame from the first sample
    message('ratio merged/denoised')
    print(summary(sapply(mergers, getN)/((sapply(dadaFs, getN)+sapply(dadaRs, getN))/2)))
  } 
  gc()
}


# # ==== Construct sequence table ==== 

if (PairedReads=='paired') {
  seqtab <- makeSequenceTable(mergers)
} else if (PairedReads=='single') {
  seqtab <- makeSequenceTable(dadaFs)
}
gc()
dim(seqtab)
table(nchar(getSequences(seqtab))) # Inspect distribution of sequence lengths

saveRDS(seqtab, file.path(PathToDadaOutputFiles,paste(ProjectName,"_seqtab.rds",sep="")))

# # ==== Optional: Filter out non-target-length sequences ==== 
# seqtab <- seqtab[,nchar(colnames(seqtab)) %in% 250:256] 

# # ==== Removes chimeras ==== 

seqtab.nochim <- removeBimeraDenovo(seqtab, method="consensus", multithread=MultiThread, verbose=TRUE)
gc()
dim(seqtab.nochim)
dim(seqtab.nochim)[2]/dim(seqtab)[2]
sum(seqtab.nochim)/sum(seqtab)
saveRDS(seqtab.nochim,file.path(PathToDadaOutputFiles,paste(ProjectName,"_seqtab_nochim.rds",sep=""))) 

if(PairedReads=="paired") {
  message('ratio nochim/merged')
  summary(rowSums(seqtab.nochim)/sapply(mergers, getN))
}


# # ==== Track your reads through the pipeline ==== 

if (PairedReads=='paired') {
    track <- out; rownames(track) <- sample.names
    if(workflow == "smalldata") {
      track <- merge(track,as.data.frame(sapply(dadaFs, getN)) ,by="row.names", all=T); rownames(track) <- track[,1]; track <- track[,-1]
      track[,5] <- track[,4]/track[,2]
      track <- merge(track,as.data.frame(sapply(dadaRs, getN)) ,by="row.names", all=T); rownames(track) <- track[,1]; track <- track[,-1]
      track[,7] <- track[,6]/track[,2]
    }
    track <- merge(track,as.data.frame(sapply(mergers, getN)) ,by="row.names", all=T); rownames(track) <- track[,1]; track <- track[,-1]
    track[,9] <- track[,8]/((track[,6]+track[,4])/2)
    track <- merge(track,as.data.frame(rowSums(seqtab.nochim)) ,by="row.names", all=T); rownames(track) <- track[,1]; track <- track[,-1]
    track[,11] <- track[,10]/track[,8]
    track[,12] <- track[,10]/track[,1]
    colnames(track) <- c("input", "filtered","ratio-filtered-input", "denoisedF","ratio-denoisedF-filtered", "denoisedR","ratio-denoisedR-filtered", "merged","ratio-merged-denoised", "nonchim","ratio-nochim-merged",'ratio-out-in')
} else if (PairedReads=='single') {
    track <- out; rownames(track) <- sample.names
    track <- merge(track,as.data.frame(sapply(dadaFs, getN)) ,by="row.names", all=T); rownames(track) <- track[,1]; track <- track[,-1]
    track[,5] <- track[,4]/track[,2]
    track <- merge(track,as.data.frame(rowSums(seqtab.nochim)) ,by="row.names", all=T); rownames(track) <- track[,1]; track <- track[,-1]
    track[,7] <- track[,6]/track[,4]
    track[,8] <- track[,6]/track[,1]
    colnames(track) <- c("input", "filtered","ratio-filtered-input", "denoisedF","ratio-denoisedF-filtered","nonchim","ratio-nochim-denoisedF","ratio-out-in")
} 

message("For the following samples no reads remained:")
rownames(track)[rowSums(is.na(track))>0]

head(track)
message('ratio filtered/input')
summary(track$`ratio-filtered-input`)
if (PairedReads=='paired') {
  message('ratio denoisedF/filtered')
  print(summary(track$`ratio-denoisedF-filtered`))
  message('ratio denoisedR/filtered')
  print(summary(track$`ratio-denoisedR-filtered`))
  message('ratio merged/denoised')
  print(summary(track$`ratio-merged-denoised`))
  message('ratio nochim/merged')
  print(summary(track$`ratio-nochim-merged`))
} else if (PairedReads=='single') {
  message('ratio denoisedF/filtered')
  print(summary(track$`ratio-denoisedF-filtered`))
  message('ratio nochim/denoisedF')
  print(summary(track$`ratio-nochim-denoisedF`))
}
message('ratio out/in')
print(summary(track$`ratio-out-in`))

data.table::fwrite(as.data.frame(track),file=file.path(PathToDadaOutputFiles,paste(ProjectName,"_track.csv",sep="")),col.names = T,row.names = T)

track.overview <- do.call(cbind, lapply(track, summary))
track.total <- do.call(cbind, lapply(track, sum))
track.overview <- rbind(track.overview,track.total)
rownames(track.overview)[nrow(track.overview)] <- "Total"

track.overview[nrow(track.overview),3] <- track.overview[nrow(track.overview),2]/track.overview[nrow(track.overview),1]
track.overview[nrow(track.overview),5] <- track.overview[nrow(track.overview),4]/track.overview[nrow(track.overview),2]
if(PairedReads=="paired") {
  track.overview[nrow(track.overview),7] <- track.overview[nrow(track.overview),6]/track.overview[nrow(track.overview),2]
  track.overview[nrow(track.overview),9] <- track.overview[nrow(track.overview),8]/((track.overview[nrow(track.overview),4]+track.overview[nrow(track.overview),6])/2)
  track.overview[nrow(track.overview),11] <- track.overview[nrow(track.overview),10]/track.overview[nrow(track.overview),8]
  track.overview[nrow(track.overview),12] <- track.overview[nrow(track.overview),10]/track.overview[nrow(track.overview),1]
} else if (PairedReads=="single") {
  track.overview[nrow(track.overview),7] <- track.overview[nrow(track.overview),6]/track.overview[nrow(track.overview),4]
  track.overview[nrow(track.overview),8] <- track.overview[nrow(track.overview),6]/track.overview[nrow(track.overview),1]
}

data.table::fwrite(as.data.frame(track.overview),file=file.path(PathToDadaOutputFiles,paste(ProjectName,"_track-overview.csv",sep="")),col.names = T,row.names = T)

save.image(file.path(PathToDadaOutputFiles,workspaceName))
gc()


# # ==== Assigns taxonomy ==== 

taxass <- assignDifTaxonomy(algorithm=classmeth, databases=database, i.sequencenochim=seqtab.nochim, i.taxonomyfolder=TaxonomyFolder, mthread=MultiThread, RC=reversecomplement, outputfolder=PathToDadaOutputFiles, nameproject=ProjectName)

if(classmeth=="dada2") {
  if(database == "unite" | database == "pr2") {
    taxa.species <- as.data.frame(taxass)
  } else {
    taxa <- as.data.frame(taxass)
  }
} else if (classmeth=="dec") {
  taxa.species <- as.data.frame(taxass)
}
rm(taxass)
gc()

## save taxonomy files
if(classmeth == "dada2") {
  if(database == "unite" | database == "pr2") {
    saveRDS(taxa.species, file.path(PathToDadaOutputFiles,paste(ProjectName,"_",classmeth,"_",database,"_taxa_species.rds",sep="")))
  } else {
    saveRDS(taxa, file.path(PathToDadaOutputFiles,paste(ProjectName,"_",classmeth,"_",database,"_taxa.rds",sep="")))
  }
} else if (classmeth == "dec") {
  saveRDS(taxa.species, file.path(PathToDadaOutputFiles,paste(ProjectName,"_",classmeth,"_",database,"_taxa_species.rds",sep="")))
}

# # species assignment only necessary for dada2 algorithm when using silva or rdp
taxasss <- assignDifTaxonomy.species(i.taxonomy=taxa, databases=database, i.taxonomyfolder=TaxonomyFolder, RC=reversecomplement, outputfolder=PathToDadaOutputFiles, nameproject=ProjectName, numrecords= 100000) # change numrecords to something lower to control peak meomry

if(classmeth=="dada2") {
  if(database == "silva132" | database == "silva138"| database == "silva1381"| database == "rdp18") {
    taxa.species <- as.data.frame(taxasss)
  }
}
rm(taxasss)
gc()

## save taxonomy files
if(classmeth == "dada2") {
  if(database == "silva132" | database == "silva138"| database == "silva1381"| database == "rdp18") {
    saveRDS(taxa.species, file.path(PathToDadaOutputFiles,paste(ProjectName,"_",classmeth,"_",database,"_taxa_species.rds",sep="")))
  } else {}
} else if (classmeth == "dec") {}

# # ==== taxonomy summary ==== 
if (!exists("taxa.species")) {
  taxa.species <- taxa
}
# # number of classified ASVs at each level
out.ASVs <- rep(nrow(taxa.species),ncol(taxa.species)) # number of different ASVs
out.ASVs <- rbind(out.ASVs,colSums(!is.na(taxa.species))) # number of ASVs classified at each level
out.ASVs <- rbind(out.ASVs, out.ASVs[2,]/out.ASVs[1,]*100) # percentage of assigned ASVs per level
rownames(out.ASVs) <- c("NumberOfASVs","NumberofAssignedASVs","percentageAssignedASVs")

out.asv.kingdom.unique <- unique(na.omit(taxa.species[,1]))

if(any(!is.na(taxa.species[,1]))) { 
  for (num in 1:length(out.asv.kingdom.unique)) {
    out.ASVs.phyl <- vector("numeric",length = ncol(taxa.species))
    for (i in 1:ncol(taxa.species)) {
      for (j in 1:nrow(taxa.species)) {
        if((!is.na(taxa.species[j,1]) & taxa.species[j,1]==out.asv.kingdom.unique[num]) & !is.na(taxa.species[j,i])) {
          out.ASVs.phyl[i] <- out.ASVs.phyl[i] + 1
        }
      }
    }
    out.ASVs <- rbind(out.ASVs,out.ASVs.phyl)
    rownames(out.ASVs)[nrow(out.ASVs)] <- paste0("NumberAssignedASVs",out.asv.kingdom.unique[num])
    out.ASVs <- rbind(out.ASVs,out.ASVs[nrow(out.ASVs),]/out.ASVs[1,]*100)
    rownames(out.ASVs)[nrow(out.ASVs)] <- paste0("PercentAssignedASVs",out.asv.kingdom.unique[num])
  }
}


# # number of classified ASVs at each level per sample (and first column total number of ASVs)
out.sample.ASVs <- matrix(nrow=nrow(seqtab.nochim),
                          ncol = ncol(taxa.species)+1,
                          dimnames = list(rownames(seqtab.nochim),c("TotalASVs-inclUnassigned",colnames(taxa.species))))

out.sample.ASVs[,1] <- rowSums(seqtab.nochim != 0) # total different ASVs

for (i in 1:ncol(taxa.species)){ # goes through each taxonomic level
  for (j in 1:nrow(seqtab.nochim)){ # goes through all samples
    count = 0
    for (k in 1:ncol(seqtab.nochim)){ # goes through all ASVs
      if(seqtab.nochim[j,k] != 0 & !is.na(taxa.species[k,i])) {
        count = count +1
      }
    }
    out.sample.ASVs[j,i+1] <- count
  }
}

# # number of classified reads at each level
out.reads <- rep(sum(seqtab.nochim),ncol(taxa.species))# total number of reads

noNA.tax <- lapply(taxa.species, function(column) which(!is.na(column))) # finds positions which are not NA in taxonomy file

out.reads.assigned <- numeric(length=ncol(taxa.species))

# if (any(!is.na(taxa.species[,1]))) { # for certain taxonomic databases reads can be unclussified at kingdom level but classified at lower levels
  for (i in 1:ncol(taxa.species)) { # goes through each taxonomic level
    count <- 0
    for (k in 1:ncol(seqtab.nochim)) { # goes through all ASVs
      if(!is.na(taxa.species[k,i])) {
        count = count + colSums(seqtab.nochim)[k]
      }
    }
    out.reads.assigned[i] <- count
  }
# }
out.reads <- rbind(out.reads,out.reads.assigned) # number of assigned reads
rownames(out.reads)[1] <- "NumberOfReads"
rownames(out.reads)[nrow(out.reads)] <- "NumberAssignedReads"
out.reads <- rbind(out.reads,out.reads[2,]/out.reads[1,]*100) # percentage of assigned reads
rownames(out.reads)[nrow(out.reads)] <- "PercentAssignedReads"

# # number of classified reads at each kingdom, for all taxonomic levels
if(any(!is.na(taxa.species[,1]))) { # here important because we are now going through kingdom levels only.
  for (num in 1:length(out.asv.kingdom.unique)) {
    out.reads.phyl <- vector("numeric",length = ncol(taxa.species))
    if (any(grepl(out.asv.kingdom.unique[num],taxa.species[,1]))) {
      for (i in 1:ncol(taxa.species)) { # goes through each taxonomic level
        count <- 0
        for (k in 1:ncol(seqtab.nochim)) { # goes through all ASVs
          if((!is.na(taxa.species[k,1]) & taxa.species[k,1]==out.asv.kingdom.unique[num]) & !is.na(taxa.species[k,i])) {
            count = count + colSums(seqtab.nochim)[k]
          }
        }
        out.reads.phyl[i] <- count
      }
    }
    out.reads <- rbind(out.reads,out.reads.phyl)
    rownames(out.reads)[nrow(out.reads)] <- paste0("NumberAssignedReads",out.asv.kingdom.unique[num])
    out.reads <- rbind(out.reads, out.reads[nrow(out.reads),]/out.reads[1,]*100)
    rownames(out.reads)[nrow(out.reads)] <- paste0("PercentAssignedReads",out.asv.kingdom.unique[num])
  }
} 

overview <- rbind(out.ASVs,rep(NA,ncol(out.ASVs)),out.reads)

# # number of classified reads at each level per sample (and first column total number of reads)
out.sample.reads <- matrix(nrow=nrow(seqtab.nochim),
                           ncol = ncol(taxa.species)+1,
                           dimnames = list(rownames(seqtab.nochim),c("TotalReads-inclUnassigned",colnames(taxa.species))))

out.sample.reads[,1] <- rowSums(seqtab.nochim) # total different ASVs

for (i in 1:ncol(taxa.species)){ # goes through each taxonomic level
  for (j in 1:nrow(seqtab.nochim)){ # goes through all samples
    count = 0
    for (k in 1:ncol(seqtab.nochim)){ # goes through all ASVs
      if(seqtab.nochim[j,k] != 0 & !is.na(taxa.species[k,i])) {
        count = count + seqtab.nochim[j,k]
      }
    }
    out.sample.reads[j,i+1] <- count
  }
}

overview.samples <- rbind(c("differentASVsAssigned",rep(NA,ncol(out.sample.ASVs)-1)),out.sample.ASVs,c("ReadsAssigned",rep(NA,ncol(out.sample.ASVs)-1)),out.sample.reads)

data.table::fwrite(as.data.frame(overview),file=file.path(PathToDadaOutputFiles,paste(ProjectName,"_assigned-overview.csv",sep="")),col.names = T,row.names = T)
data.table::fwrite(as.data.frame(overview.samples),file=file.path(PathToDadaOutputFiles,paste(ProjectName,"_assigned-samples-overview.csv",sep="")),col.names = T,row.names = T)
```
