## Supplementary material for "A global atlas of subsurface microbiomes reveals phylogenetic novelty, large scale biodiversity gradients, and a marine-terrestrial divide": Dataset S4 - VisuaR Analysis R script: DatasetS4_VisuaR_v40_script.R

```
basic.packages <- c("package:stats","package:graphics","package:grDevices","package:utils","package:datasets","package:methods","package:base")
package.list <- search()[ifelse(unlist(gregexpr("package:",search()))==1,TRUE,FALSE)]
package.list <- setdiff(package.list,basic.packages)
if (length(package.list)>0)  for (package in package.list) detach(package, character.only=TRUE)
rm(list=ls()); closeAllConnections(); gc()

# #=== VisuaR Info ========================================================================================================================================================================
# # Workflow by Emil Ruff & Isabella Hrabe de Angelis 09/2023
# # Copyright Emil Ruff
# # The authors acknowledge valuable input by Alban Ramette and Angelique Gobet
# # Please cite https://github.com/EmilRuff/VisuaR

printinfo <- function(df) {
  cat('(total=',sum(df),', min=',min(df),', 1st Qu.=',as.integer(summary(df))[2],', median=',median(df),', mean=',round(mean(df),0),', 3rd Qu.=',as.integer(summary(df))[5],', max=',max(df),', sd=',round(sd(df,na.rm = T),0),', se=',round(sd(df, na.rm=TRUE)/sqrt(length(na.omit(df))),0),')\n',file=(file.path(PathToVisuaRAnalysis,paste(VisuaRProjectName,'_Analysis_log.txt',sep=''))),sep='',append=TRUE)
}

fill.info <- function(df.reads,df.ASVs,df.seqtab,stepname,infotable) {
  infostep <- data.frame(matrix(nrow=1,ncol=ncol(infotable)))
  rownames(infostep)[1] <- stepname
  infostep[1,1] <- length(df.reads) ## number of samples
  infostep[1,2] <- sum(df.reads) ## total number of reads in all samples
  infostep[1,3] <- min(df.reads) ## minimum number of reads in all samples
  infostep[1,4] <- as.integer(summary(df.reads))[2] ## first Quartile of number of reads in all samples
  infostep[1,5] <- median(df.reads) ## median of number of reads in all samples
  infostep[1,6] <- round(mean(df.reads),0) ## mean of number of reads in all samples
  infostep[1,7] <- as.integer(summary(df.reads))[5] ## Third Quartile of number of reads in all samples
  infostep[1,8] <- max(df.reads) ## maxmum number of reads in all samples
  infostep[1,9] <- round(sd(df.reads,na.rm = T),0) ## standard deviation of number of reads in all samples
  infostep[1,10] <- round(sd(df.reads, na.rm=TRUE)/sqrt(length(na.omit(df.reads))),0) ## standard error of number of reads in all samples
  
  infostep[1,11] <- nrow(df.seqtab) ## total number of different ASVs
  infostep[1,12] <- sum(df.ASVs) ## total number of ASVs in all samples
  infostep[1,13] <- min(df.ASVs) ## minimum number of ASVs in all samples
  infostep[1,14] <- as.integer(summary(df.ASVs))[2] ## first Quartile of number of ASVs in all samples
  infostep[1,15] <- median(df.ASVs) ## median of number of ASVs in all samples
  infostep[1,16] <- round(mean(df.ASVs),0) ## mean of number of ASVs in all samples
  infostep[1,17] <- as.integer(summary(df.ASVs))[5] ## Third Quartile of number of ASVs in all samples
  infostep[1,18] <- max(df.ASVs) ## maxmum number of ASVs in all samples
  infostep[1,19] <- round(sd(df.ASVs,na.rm = T),0) ## standard deviation of number of ASVs in all samples
  infostep[1,20] <- round(sd(df.ASVs, na.rm=TRUE)/sqrt(length(na.omit(df.ASVs))),0) ## standard error of number of ASVs in all samples
  colnames(infostep) <- colnames(infotable)
  infotable <- rbind(infotable,infostep)
  return(infotable)
}

Sys.setenv(TZ = "UTC") # set to UTC or change to your time one

# #=== 1. Source external file with set parameters =========================================================================================================================================================
# # Provide the location of your input .R script.
# # The script will automatically be saved to your analysis folder

# UserInput <- file.path("/path/to/VisuaR_Input.R")
# UserInput <- file.path("/Users/emilruff/Dropbox/__Emil/_Beaver_Pond_Project/Amplicon/2023_Amplicon_analysis/VisuaR_v33_UserInput_BP_2023_Arch_SampleType_noBlanks.R") 
# UserInput <- file.path("C:/Users/i.hrabedeangelis/Dropbox/scripts/R/Eugen/Eugen_VisuaR_v30_UserInput.R")
# UserInput <- file.path("C:/Users/i.hrabedeangelis/Dropbox/scripts/R/VisuaR_v35_UserInput_CoDL.R")
# UserInput <- file.path("C:/Users/i.hrabedeangelis/Dropbox/_CoDL_2020_2022/_VisuaR_2022/20230925_CoDL_arc_mar_depthbiome/20230925_CoDL_arc_mar_depthbiome_UserInput.R")


# UserInput <- file.path("C:/Users/i.hrabedeangelis/Dropbox/scripts/R/Eugen/Eugen_VisuaR_v30_UserInput_16S.R")

UserInput <- file.path("C:/Users/i.hrabedeangelis/Dropbox/scripts/R/VisuaR_v39_UserInput_CoDL.R")

source(UserInput)

# #=== 2. Prologue =========================================================================================================================

# #=== 2.1. Creates file directories =========================================================================================================================
PathToVisuaRAnalysis=file.path(PathToVisuarOutput,'VisuaR',paste(VisuaRProjectName,sep='_')) # creates directories for the current VisuaR analysis

dir.create(file.path(PathToVisuarOutput,'VisuaR'),recursive = T) 

if (dir.exists(PathToVisuaRAnalysis)) {
  stop("The folder already exists. Script execution stopped.")
} else {
  dir.create(PathToVisuaRAnalysis)
}
dir.create(file.path(PathToVisuaRAnalysis,'Alpha_Diversity'))
dir.create(file.path(PathToVisuaRAnalysis,'Beta_Diversity'))
file.copy(UserInput,file.path(PathToVisuaRAnalysis,paste(VisuaRProjectName,'_UserInput.R',sep='')))

# #=== 2.2. Creates log file =========================================================================================================================
cat('VISUAR Analysis',file=(file.path(PathToVisuaRAnalysis,paste(VisuaRProjectName,'_Analysis_log.txt',sep=''))),sep='',append=TRUE)
cat('\n\n1. User INPUT',file=(file.path(PathToVisuaRAnalysis,paste(VisuaRProjectName,'_Analysis_log.txt',sep=''))),sep='',append=TRUE)
sink(file=(file.path(PathToVisuaRAnalysis,paste(VisuaRProjectName,'_Analysis_log.txt',sep=''))),append=TRUE,type='output')
cat('\n',sep='')
print(paste('AnalysisDate: ',Sys.Date()))
cat('VisuaRProjectName: ',VisuaRProjectName,'\n',sep='')
cat('Seqtab nochim originates from: ',PathToSeqtabNochim,'\n',sep='')
cat('Taxonomy originates from: ',PathToTaxonomy,'\n',sep='')
cat('contextdata originates from: ',contextdatafilepath,'\n\n',sep='')
cat('ASVs belonging to the KingdomOfInterest: ',KingdomOfInterest,' will be kept for the further analysis.','\n',sep='')
if (MinimumAllowedReadCount!=''|MinimumAllowedReadCount!=0) {
  cat('Samples with less then ',MinimumAllowedReadCount,' reads (in all kingdoms, without filtering) will be excluded from the analysis.','\n',sep='')
} else {
  cat('No samples will be excluded due to low read counts (in all kingdoms, without filtering) will be excluded as no MinimumAllowedReadCount was set.','\n',sep='')
}
if (CladeOfInterest!='') {
  cat('Only ASVs belonging to the CladeOfInterest ',CladeOfInterest,' will be kept for further analysis','\n',sep='')
} else {
  cat('No CladeOfInterest was chosen.','\n',sep='')
}
if (ExcludeClade!='') {
  cat('ASVs belonging to ExcludeClade ',ExcludeClade,' will be excluded from the analysis.','\n',sep='')
} else {
  cat('No ExcludeClade was chosen.','\n',sep='')
}
if (KeepSamplesbyName!='') {
  cat('Samples will be kept based on: ','\n',sep='')
  cat('KeepSamplesbyName: ',KeepSamplesbyName,'\n',sep='')
} else {
  cat('All samples will be kept based on the sample name as no KeepSamplesbyName was chosen.','\n',sep='')
}
if (excludeSamplesbyName!='') {
  cat('Samples will be excluded based on: ','\n',sep='')
  cat('excludeSamplesbyName: ',excludeSamplesbyName,'\n',sep='')
} else {
  cat('No samples will be excluded based on the sample name as no excludeSamplesbyName was chosen.','\n',sep='')
}
if (exists('MinimumAllowedReadCount.Analysis')==F) {
  cat('No MinimumAllowedReadCount.Analysis was set or it was set to 0. No sample will be excluded based on too low read counts after subsetting.')
} else {
  cat('Samples with less than ',MinimumAllowedReadCount.Analysis,' reads (in the subsetted data) will be excluded from the analysis.','\n',sep='')
}
if (contextdata=='N') {
  cat('\nNo contextdata was provided.','\n',sep='')
} else {
  cat('\ncontextdata was provided.','\n',sep='')
  cat('The contextdata originated from ',contextdatafilepath,'\n',sep='')
  cat('The samples will be grouped based on the category ',Grouping1,' (Grouping1).','\n',sep='')
  if (Grouping2!='') {
    cat('The samples will be sorted inside Grouping1 based on the category ',Grouping2,' (Grouping2).','\n',sep='')
  } else {
    cat('No Grouping2 was provided. The samples will not be sorted inside the main Grouping1.','\n',sep='')
  }
  if (VariableToExclude!='') {
    cat('Samples belonging to the VariableToExclude: ',VariableToExclude,' as found in the contextual data column ',ColumnOfVariableToExclude,' will be excluded from the analysis','\n',sep='')
  } else {
    cat('No samples belonging to a particular category in your contextdata sheet will be excluded as no VariableToExclude was chosen.','\n',sep='')
  }
  if (VariableToKeep!='') {
    cat('Samples not belonging to the VariableToKeep: ',VariableToKeep,' as found in the contextual data column ',ColumnOfVariableToKeep,' will be excluded from the analysis','\n',sep='')
  } else {
    cat('No samples belonging to a particular category in your contextdata sheet will be excluded as no VariableToKeep was chosen.','\n',sep='')
  }
}
sink()

cat('\n\n2. Prologue',file=(file.path(PathToVisuaRAnalysis,paste(VisuaRProjectName,'_Analysis_log.txt',sep=''))),sep='',append=TRUE)

# #=== 2.3. Loads required packages =======================================================================================================================================================

cat('\n2.3. Load required packages',file=(file.path(PathToVisuaRAnalysis,paste(VisuaRProjectName,'_Analysis_log.txt',sep=''))),sep='',append=TRUE)
cat('\nInformation on your used package Versions and R version can be found in the txt file VersionInformation.txt',file=(file.path(PathToVisuaRAnalysis,paste(VisuaRProjectName,'_Analysis_log.txt',sep=''))),sep='',append=TRUE)
sink(file=(file.path(PathToVisuaRAnalysis,'Session_Info.txt')),append=TRUE)
sink(stdout(),type='message')
if (!require('tidyverse')) {install.packages('tidyverse')}; library(tidyverse) # contains packages: ggplot2, dplyr, tidyr, readr, purrr, tibble, stringr, forcats
if (!require('plotrix')) {install.packages('plotrix')}; library(plotrix)
if (!require('plyr')) {install.packages('plyr')}; library(plyr) # shouold be fine with loading tidyverse only
if (!require('vegan')) {install.packages('vegan')}; library(vegan)
if (!require('reshape2')) {install.packages('reshape2')}; library(reshape2)
if (!require('ggsignif')) {install.packages('ggsignif')}; library(ggsignif)
if (!require('EnvStats')) {install.packages('EnvStats')}; library(EnvStats)
if (!require('ggpubr')) {install.packages('ggpubr')}; library(ggpubr)
if (!require('ape')) {install.packages('ape')}; library(ape)
if (!require('UpSetR')) {install.packages('UpSetR')}; library(UpSetR)
if (!require('venn')) {install.packages('venn')}; library(venn)
if (!require('eulerr')) {install.packages('eulerr')}; library(eulerr)
if (!require('indicspecies')) {install.packages('indicspecies')}; library(indicspecies)
if (!require('openxlsx')) {install.packages('openxlsx')}; library(openxlsx)
if (!require('RColorBrewer')) {install.packages('RColorBrewer')}; library(RColorBrewer)
if (!require('rstudioapi')) {install.packages('rstudioapi')}; library(rstudioapi)
if (!require('compositions')) {install.packages('compositions')}; library(compositions)
if (!require('gridGraphics')) {install.packages('gridGraphics')}; library(gridGraphics)
if(removeContaminants=="decontam"){
  if (!require('decontam')) {install.packages('decontam')}; library(decontam)
}
sessionInfo()
closeAllConnections() # closes all currently open connections.


# #=== 2.4. Reads and converts dada2 sequence table and taxonomy to match format used by VisuaR ===============================================================================================
cat('\n\n2.4. Read and convert dada2 sequence table and taxonomy to match format used by VisuaR.',file=(file.path(PathToVisuaRAnalysis,paste(VisuaRProjectName,'_Analysis_log.txt',sep=''))),sep='',append=T)

if (file.exists(file.path(paste(str_remove(PathToTaxonomy, '.rds'),'_noNAs.rds',sep='')))) {
  M.taxo.noNA=readRDS(file=file.path(paste(str_remove(PathToTaxonomy, '.rds'),'_noNAs.rds',sep='')))
} else {
  if (HappyBellydada2Input=='Y') {
    M.taxo.noNA=as.matrix(read.table(file = file.path(PathToTaxonomy)))
  } else if (QIIMEInput=='Y') {
    M.taxo.orig=read.table(file = file.path(PathToTaxonomy),sep='\t',header = T)
    rownames(M.taxo.orig)=M.taxo.orig[,1]
    M.taxo.orig=as.data.frame(M.taxo.orig[,-3])
    M.taxo.noNA=as.data.frame(str_split_fixed(M.taxo.orig[,2],';',n=Inf))
    colnames(M.taxo.noNA)=c('Kingdom','Phylum','Class','Order','Family','Genus','Species')
    M.taxo.noNA[M.taxo.noNA==""]=NA
    M.taxo.noNA=data.frame(lapply(M.taxo.noNA, function(x){gsub('D_0__','',x)}))      # replaces all brackets in the taxonomy file with a '-'. While this happens the ASVs obtain a running number instead of the ASV Sequence.
    M.taxo.noNA=data.frame(lapply(M.taxo.noNA, function(x){gsub('D_1__','',x)}))      # replaces all brackets in the taxonomy file with a '-'. While this happens the ASVs obtain a running number instead of the ASV Sequence.
    M.taxo.noNA=data.frame(lapply(M.taxo.noNA, function(x){gsub('D_2__','',x)}))      # replaces all brackets in the taxonomy file with a '-'. While this happens the ASVs obtain a running number instead of the ASV Sequence.
    M.taxo.noNA=data.frame(lapply(M.taxo.noNA, function(x){gsub('D_3__','',x)}))      # replaces all brackets in the taxonomy file with a '-'. While this happens the ASVs obtain a running number instead of the ASV Sequence.
    M.taxo.noNA=data.frame(lapply(M.taxo.noNA, function(x){gsub('D_4__','',x)}))      # replaces all brackets in the taxonomy file with a '-'. While this happens the ASVs obtain a running number instead of the ASV Sequence.
    M.taxo.noNA=data.frame(lapply(M.taxo.noNA, function(x){gsub('D_5__','',x)}))      # replaces all brackets in the taxonomy file with a '-'. While this happens the ASVs obtain a running number instead of the ASV Sequence.
    M.taxo.noNA=data.frame(lapply(M.taxo.noNA, function(x){gsub('D_6__','',x)}))      # replaces all brackets in the taxonomy file with a '-'. While this happens the ASVs obtain a running number instead of the ASV Sequence.
    rownames(M.taxo.noNA)=rownames(M.taxo.orig)
    rm(M.taxo.orig)
    gc()
    M.taxo.noNA=as.matrix(M.taxo.noNA)
  } else if (MetagenomeInput=='Y') {
    M.taxo.orig=read.table(file = file.path(PathToTaxonomy),sep='\t',header = T)
    rownames(M.taxo.orig)=M.taxo.orig[,1]
    M.taxo.orig=M.taxo.orig[,-1]
    M.taxo.noNA=M.taxo.orig
    M.taxo.noNA[M.taxo.noNA==""]=NA
    M.taxo.noNA=as.matrix(M.taxo.noNA)
  } else if (IsthisCoDLphyloSSU == "Y") {
    M.taxo.orig=as.matrix(read.csv(file=file.path(PathToTaxonomy))[,1])
    M.taxo.noNA=as.data.frame(str_split_fixed(M.taxo.orig[,1],';',n=Inf))
    colnames(M.taxo.noNA)=c('Kingdom','Phylum','Class','Order','Family','Genus','Species')
    
  } else if (IsthisCoDLrps3_kaiju == "Y") {
    M.taxo.orig=as.matrix(read.csv(file=file.path(PathToTaxonomy))[,2])
    M.taxo.noNA=as.data.frame(str_split_fixed(M.taxo.orig[,1],';',n=Inf))
    M.taxo.noNA <- M.taxo.noNA[,-c(ncol(M.taxo.noNA))]
    colnames(M.taxo.noNA)=c('Kingdom','Phylum','Class','Order','Family','Genus','Species')
    
  } else if (IsthisCoDLrps3_cat == "Y") {
    M.taxo.orig=as.matrix(read.csv(file=file.path(PathToTaxonomy))[,3])
    M.taxo.noNA=as.data.frame(str_split_fixed(M.taxo.orig[,1],';',n=Inf))
    colnames(M.taxo.noNA)=c('Kingdom','Phylum','Class','Order','Family','Genus','Species')
    
  } else if (IsthisCoDLrps3_uniprot == "Y") {
    M.taxo.orig=as.matrix(read.csv(file=file.path(PathToTaxonomy))[,4])
    M.taxo.noNA=as.data.frame(str_split_fixed(M.taxo.orig[,1],';',n=Inf))
    M.taxo.noNA <- M.taxo.noNA[,-1]
    M.taxo.noNA <- M.taxo.noNA[, !(colSums(is.na(M.taxo.noNA) | M.taxo.noNA == "") == nrow(M.taxo.noNA))]
    # colnames(M.taxo.noNA)=c('Kingdom','Phylum','Class','Order','Family','Genus','Species')
    
  
  } else {
    M.taxo.noNA=readRDS(file=file.path(PathToTaxonomy))
  }
}

short.save <- rownames(M.taxo.noNA)
M.taxo.noNA <- data.frame(M.taxo.noNA)
M.taxo.noNA <- data.frame(lapply(M.taxo.noNA, function(x){gsub('\\[|\\]','',x)}))      # replaces all [] in the taxonomy file with nothing. While this happens the ASVs obtain a running number instead of the ASV Sequence.
M.taxo.noNA <- data.frame(lapply(M.taxo.noNA, function(x){gsub('\\*','',x)}))      # replaces all * in the taxonomy file with nothing. While this happens the ASVs obtain a running number instead of the ASV Sequence.
rownames(M.taxo.noNA) <- short.save
M.taxo.noNA=as.data.frame(M.taxo.noNA)  # ASV by Taxonomy data frame

ncol.taxo.noNA=ncol(M.taxo.noNA) # Number of taxonomic levels. Should be 7.
nrow.taxo.noNA=nrow(M.taxo.noNA) # Number of rows in taxonomy file = number of observed ASVs

gc()
# #=== 2.5. Modifies taxonomy tables ===============================================================================================================

# # deletes double information in pr2 database and replaces it with NA
if (class.algorithm == "dada2" && tax.database == "pr2") {
  colnames(M.taxo.noNA) <- c("Kingdom","Supergroup","Phylum","rp2Class","Class","Order","Family","Genus","Species")
  M.taxo.noNA[] <- lapply(M.taxo.noNA, function(x) sub("_XXXXXXXXX", "", x, fixed = TRUE))
  M.taxo.noNA[] <- lapply(M.taxo.noNA, function(x) sub("_XXXXXXXX", "", x, fixed = TRUE))
  M.taxo.noNA[] <- lapply(M.taxo.noNA, function(x) sub("_XXXXXXX", "", x, fixed = TRUE))
  M.taxo.noNA[] <- lapply(M.taxo.noNA, function(x) sub("_XXXXXX", "", x, fixed = TRUE))
  M.taxo.noNA[] <- lapply(M.taxo.noNA, function(x) sub("_XXXXX", "", x, fixed = TRUE))
  M.taxo.noNA[] <- lapply(M.taxo.noNA, function(x) sub("_XXXX", "", x, fixed = TRUE))
  M.taxo.noNA[] <- lapply(M.taxo.noNA, function(x) sub("_XXX", "", x, fixed = TRUE))
  M.taxo.noNA[] <- lapply(M.taxo.noNA, function(x) sub("_XX", "", x, fixed = TRUE))
  M.taxo.noNA[] <- lapply(M.taxo.noNA, function(x) sub("_X", "", x, fixed = TRUE))
  M.taxo.noNA[,9] <- sub("_sp.","",M.taxo.noNA[,9]) 
  for (k in 9:2) {
    M.taxo.noNA[,k] <- ifelse(M.taxo.noNA[,k] == M.taxo.noNA[,k-1], NA, M.taxo.noNA[,k])
  }
}

# #=== 2.5. Replaces NAs in taxonomy with 'unc' and concatenates the taxonomy ===============================================================================================================

if (IsthisCoDLrps3_kaiju=="Y") {
  if(!file.exists(file.path(paste(str_remove(PathToTaxonomy, '.csv'),'_kaiju_noNAs.rds',sep='')))) {
    UnclassifiedTerm='unc'                                                                    
    M.taxo.noNA=as.matrix(M.taxo.noNA)
    M.taxo.noNA[is.na(M.taxo.noNA) | M.taxo.noNA == ""]=UnclassifiedTerm 
    M.taxo.noNA[,1]=(as.character(M.taxo.noNA[,1]))
    for(k in 1:nrow(M.taxo.noNA)){ # goes through all ASVs (rows) and combines names from different taxonomy levels with an _
      for (j in 2:ncol(M.taxo.noNA)) {
        M.taxo.noNA[k,j] <- paste(as.character(M.taxo.noNA[k,j-1]),as.character(M.taxo.noNA[k,j]),sep="_")
      }
    }
    M.taxo.noNA=as.data.frame(M.taxo.noNA,stringsAsFactors=FALSE)
    saveRDS(M.taxo.noNA,file.path(file.path(paste(str_remove(PathToTaxonomy, '.csv'),'_kaiju_noNAs.rds',sep=''))))
  } else {
    M.taxo.noNA <- readRDS(file.path(file.path(paste(str_remove(PathToTaxonomy, '.csv'),'_kaiju_noNAs.rds',sep=''))))
  }
} else if (IsthisCoDLrps3_cat=="Y") {
  if(!file.exists(file.path(paste(str_remove(PathToTaxonomy, '.csv'),'_cat_noNAs.rds',sep='')))) {
    UnclassifiedTerm='unc'                                                                    
    M.taxo.noNA=as.matrix(M.taxo.noNA)
    M.taxo.noNA[is.na(M.taxo.noNA) | M.taxo.noNA == ""]=UnclassifiedTerm 
    M.taxo.noNA[,1]=(as.character(M.taxo.noNA[,1]))
    for(k in 1:nrow(M.taxo.noNA)){ # goes through all ASVs (rows) and combines names from different taxonomy levels with an _
      for (j in 2:ncol(M.taxo.noNA)) {
        M.taxo.noNA[k,j] <- paste(as.character(M.taxo.noNA[k,j-1]),as.character(M.taxo.noNA[k,j]),sep="_")
      }
    }
    M.taxo.noNA=as.data.frame(M.taxo.noNA,stringsAsFactors=FALSE)
    saveRDS(M.taxo.noNA,file.path(file.path(paste(str_remove(PathToTaxonomy, '.csv'),'_cat_noNAs.rds',sep=''))))
  } else {
    M.taxo.noNA <- readRDS(file.path(file.path(paste(str_remove(PathToTaxonomy, '.csv'),'_cat_noNAs.rds',sep=''))))
  }
} else if (IsthisCoDLrps3_uniprot=="Y") {
  if(!file.exists(file.path(paste(str_remove(PathToTaxonomy, '.csv'),'_uniprot_noNAs.rds',sep='')))) {
    UnclassifiedTerm='unc'                                                                    
    M.taxo.noNA=as.matrix(M.taxo.noNA)
    M.taxo.noNA[is.na(M.taxo.noNA) | M.taxo.noNA == ""]=UnclassifiedTerm 
    M.taxo.noNA[,1]=(as.character(M.taxo.noNA[,1]))
    for(k in 1:nrow(M.taxo.noNA)){ # goes through all ASVs (rows) and combines names from different taxonomy levels with an _
      for (j in 2:ncol(M.taxo.noNA)) {
        M.taxo.noNA[k,j] <- paste(as.character(M.taxo.noNA[k,j-1]),as.character(M.taxo.noNA[k,j]),sep="_")
      }
    }
    M.taxo.noNA=as.data.frame(M.taxo.noNA,stringsAsFactors=FALSE)
    saveRDS(M.taxo.noNA,file.path(file.path(paste(str_remove(PathToTaxonomy, '.csv'),'_uniprot_noNAs.rds',sep=''))))
  } else {
    M.taxo.noNA <- readRDS(file.path(file.path(paste(str_remove(PathToTaxonomy, '.csv'),'_uniprot_noNAs.rds',sep=''))))
  }
} else {
  if (!file.exists(file.path(paste(str_remove(PathToTaxonomy, '.rds'),'_noNAs.rds',sep='')))) { # Replaces all 'NA' by 'unc' and concatenates taxonomic levels using '_' as separator
    UnclassifiedTerm='unc'                                                                    
    M.taxo.noNA=as.matrix(M.taxo.noNA)
    M.taxo.noNA[is.na(M.taxo.noNA)]=UnclassifiedTerm 
    M.taxo.noNA[,1]=(as.character(M.taxo.noNA[,1]))
    for(k in 1:nrow(M.taxo.noNA)){ # goes through all ASVs (rows) and combines names from different taxonomy levels with an _
      for (j in 2:ncol(M.taxo.noNA)) {
        M.taxo.noNA[k,j] <- paste(as.character(M.taxo.noNA[k,j-1]),as.character(M.taxo.noNA[k,j]),sep="_")
      }
    }
    M.taxo.noNA=as.data.frame(M.taxo.noNA,stringsAsFactors=FALSE)
    saveRDS(M.taxo.noNA,file.path(file.path(paste(str_remove(PathToTaxonomy, '.rds'),'_noNAs.rds',sep=''))))
  }
}


# 
# if (!file.exists(file.path(paste(str_remove(PathToTaxonomy, '.rds'),'_noNAs.rds',sep='')))) { # Replaces all 'NA' by 'unc' and concatenates taxonomic levels using '_' as separator
#   UnclassifiedTerm='unc'                                                                    
#   M.taxo.noNA=as.matrix(M.taxo.noNA)
#   M.taxo.noNA[is.na(M.taxo.noNA) | M.taxo.noNA == ""]=UnclassifiedTerm 
#   M.taxo.noNA[,1]=(as.character(M.taxo.noNA[,1]))
#   for(k in 1:nrow(M.taxo.noNA)){ # goes through all ASVs (rows) and combines names from different taxonomy levels with an _
#     for (j in 2:ncol(M.taxo.noNA)) {
#       M.taxo.noNA[k,j] <- paste(as.character(M.taxo.noNA[k,j-1]),as.character(M.taxo.noNA[k,j]),sep="_")
#     }
#   }
#   M.taxo.noNA=as.data.frame(M.taxo.noNA,stringsAsFactors=FALSE)
#   saveRDS(M.taxo.noNA,file.path(file.path(paste(str_remove(PathToTaxonomy, '.rds'),'_noNAs.rds',sep=''))))
# }

# #=== 2.6. Reads and prepares Seqtab_nochim =========================================================================================================================================================

if (HappyBellydada2Input=='Y') {
  M.seqtab.nochim=as.matrix(t(read.table(file.path(PathToSeqtabNochim))))
} else if (QIIMEInput=='Y') {
  QIIME_Table_orig = biom::read_biom(file.path(PathToSeqtabNochim))
  M.seqtab.nochim=as.data.frame(QIIME_Table_orig$data)
  names=as.character(QIIME_Table_orig$rows)
  names.split=as.data.frame(str_split_fixed(names,'"',n=Inf))
  rm(names)
  names.split=names.split[,2]
  colnames(M.seqtab.nochim)=names.split
  rm(QIIME_Table_orig,names.split)
  gc()
} else if (MetagenomeInput=='Y') {
  M.seqtab.nochim=as.matrix(t(read.table(file.path(PathToSeqtabNochim))))
  # # added for codl
} else if (IsthisCoDL_Metagenome=='Y') {
  M.seqtab.nochim=as.matrix(t(read.table(file.path(PathToSeqtabNochim))))
  rownames(M.seqtab.nochim)=M.seqtab.nochim[,1]
  colnames(M.seqtab.nochim)=M.seqtab.nochim[1,]
  M.seqtab.nochim=M.seqtab.nochim[-1,-1]
  M.seqtab.nochim=as.data.frame(M.seqtab.nochim)
  M.seqtab.nochim[]=lapply(M.seqtab.nochim,function(x) as.numeric(as.character(x)))
  M.seqtab.nochim=as.matrix(M.seqtab.nochim)
} else if (IsthisCoDLphyloSSU=='Y') {
  M.seqtab.nochim=as.matrix(read.csv(file=file.path(PathToTaxonomy)))
  M.seqtab.nochim<-M.seqtab.nochim[,2:ncol(M.seqtab.nochim)]
  M.seqtab.nochim<-as.data.frame(t(M.seqtab.nochim))
  M.seqtab.nochim[]=lapply(M.seqtab.nochim,function(x) as.numeric(as.character(x)))
  M.seqtab.nochim=as.matrix(M.seqtab.nochim)
} else if (IsthisCoDLrps3_cat=="Y"|IsthisCoDLrps3_kaiju=="Y"|IsthisCoDLrps3_uniprot=="Y") {
  M.seqtab.nochim=as.matrix(read.csv(file=file.path(PathToTaxonomy)))
  rownames(M.seqtab.nochim)=M.seqtab.nochim[,1]
  M.seqtab.nochim=M.seqtab.nochim[,-c(1:4)]
  M.seqtab.nochim<-as.data.frame(t(M.seqtab.nochim))
  M.seqtab.nochim[]=lapply(M.seqtab.nochim,function(x) as.numeric(as.character(x)))
  M.seqtab.nochim=as.matrix(M.seqtab.nochim)
} else if  (file.exists(file.path(paste(str_remove(PathToSeqtabNochim, '.rds'),'_rename.rds',sep='')))) {
  M.seqtab.nochim=readRDS(file.path(paste(str_remove(PathToSeqtabNochim, '.rds'),'_rename.rds',sep='')))
} else {
  M.seqtab.nochim=readRDS(file.path(PathToSeqtabNochim))
}

# if (HappyBellydada2Input=='Y') {
#   M.seqtab.nochim=as.matrix(t(read.table(file.path(PathToSeqtabNochim))))
# } else if (QIIMEInput=='Y') {
#   QIIME_Table_orig = biom::read_biom(file.path(PathToSeqtabNochim))
#   M.seqtab.nochim=as.data.frame(QIIME_Table_orig$data)
#   names=as.character(QIIME_Table_orig$rows)
#   names.split=as.data.frame(str_split_fixed(names,'"',n=Inf))
#   rm(names)
#   names.split=names.split[,2]
#   colnames(M.seqtab.nochim)=names.split
#   rm(QIIME_Table_orig,names.split)
#   gc()
# } else if (MetagenomeInput=='Y') {
#   M.seqtab.nochim=as.matrix(t(read.table(file.path(PathToSeqtabNochim))))
# } else if (exists("isthiscodl")) {
#   if (isthiscodl=="Y") {
#     if  (file.exists(file.path(paste(str_remove(PathToSeqtabNochim, '.rds'),'_rename.rds',sep='')))) {
#       M.seqtab.nochim=readRDS(file.path(paste(str_remove(PathToSeqtabNochim, '.rds'),'_rename.rds',sep='')))
#     }
#   }
# } else {
#   M.seqtab.nochim=readRDS(file.path(PathToSeqtabNochim))
# }

ncol.seqtab.nochim=ncol(M.seqtab.nochim) # Number of columns in Seqtab_nochim = number of observed ASVs

ASV.names=rownames(M.taxo.noNA)


if (HappyBellydada2Input=='Y') {
  if (nrow(M.taxo.noNA)==ncol(M.seqtab.nochim)) {
    M.taxo.noNA=data.frame(lapply(M.taxo.noNA, function(x){gsub('[()]','-',x)}))      # replaces all brackets in the taxonomy file with a '-'. While this happens the ASVs obtain a running number instead of the ASV Sequence.
    colnames(M.seqtab.nochim)=rownames(M.taxo.noNA)                                   # replaces ASV identifiers in M.seqtab.nochim by running number as well
    M.seqtab.nochim=as.matrix(as.data.frame(M.seqtab.nochim)) 
  } else {
    cat("\nError: You're Taxonomy file does not match your Seqtab.\n",file=(file.path(PathToVisuaRAnalysis,paste(VisuaRProjectName,'_Analysis_log.txt',sep=''))),sep='\t',append=TRUE)
  }
} else if (exists("isthiscodl")) {
  if (isthiscodl=="Y") {
    if (ncol(M.seqtab.nochim)==nrow(M.taxo.noNA)) {
      M.taxo.noNA=data.frame(lapply(M.taxo.noNA, function(x){gsub('[()]','-',x)}))      # replaces all brackets in the taxonomy file with a '-'. While this happens the ASVs obtain a running number instead of the ASV Sequence.
      colnames(M.seqtab.nochim)=rownames(M.taxo.noNA)                                   # replaces ASV identifiers in M.seqtab.nochim by running number as well
      M.seqtab.nochim=as.matrix(as.data.frame(M.seqtab.nochim)) 
    }else {
      cat("\nError: You're Taxonomy file does not match your Seqtab.\n",file=(file.path(PathToVisuaRAnalysis,paste(VisuaRProjectName,'_Analysis_log.txt',sep=''))),sep='\t',append=TRUE)
      stop("You're Taxonomy file does not match your Seqtab.")
    }
  }
} else {
  if (identical(rownames(M.taxo.noNA),colnames(M.seqtab.nochim))){                    # checks whether the ASVs in the taxonomy and in the seqtab_nochim are identical
    M.taxo.noNA=data.frame(lapply(M.taxo.noNA, function(x){gsub('[()]','-',x)}))      # replaces all brackets in the taxonomy file with a '-'. While this happens the ASVs obtain a running number instead of the ASV Sequence.
    colnames(M.seqtab.nochim)=rownames(M.taxo.noNA)                                   # replaces ASV identifiers in M.seqtab.nochim by running number as well
    M.seqtab.nochim=as.matrix(as.data.frame(M.seqtab.nochim)) 
  } else {
    cat("\nError: You're Taxonomy file does not match your Seqtab.\n",file=(file.path(PathToVisuaRAnalysis,paste(VisuaRProjectName,'_Analysis_log.txt',sep=''))),sep='\t',append=TRUE)
    stop("You're Taxonomy file does not match your Seqtab.")
  }
}

ASV.mapfile=cbind(ASV.names,M.taxo.noNA)                                            # prints a summary to archive which ASV identifier (running number) belongs to which ASV sequence
ASV.mapfile=cbind(row.names(ASV.mapfile),data.frame(ASV.mapfile,row.names = NULL))  # reformats the table to obtain excel readable output file
colnames(ASV.mapfile)[1]='ASV'
colnames(ASV.mapfile)[2]='ASV sequence'

data.table::fwrite(ASV.mapfile,file=file.path(PathToVisuaRAnalysis,paste(VisuaRProjectName,'_ASV_mapfile.csv',sep='')),col.names = T,row.names = T)

if (SaveWholeworkspace=='N') { # removes file for efficient use of resources
  rm(ASV.mapfile,ASV.names) 
}
gc()

# if(exists("isthiscodl")) {
#   if (isthiscodl=="Y") {
#     if  (file.exists(file.path(paste(str_remove(PathToSeqtabNochim, '.rds'),'_rename.rds',sep='')))) {
      # M.seqtab.nochim=readRDS(file.path(paste(str_remove(PathToSeqtabNochim, '.rds'),'_rename.rds',sep='')))
#     } else {  
#       rownames.seqtab.nochim=(rownames(M.seqtab.nochim))
#       rownames.seqtab.nochim.split=as.data.frame(str_split_fixed(rownames.seqtab.nochim,"__",n=Inf))
#       rownames.new.seqtab.nochim=as.data.frame(paste(rownames.seqtab.nochim.split$V2,rownames.seqtab.nochim.split$V3,sep="_"))
#       rownames(M.seqtab.nochim)=rownames.new.seqtab.nochim[,1]
#       saveRDS(M.seqtab.nochim,file.path(paste(str_remove(PathToSeqtabNochim, '.rds'),'_rename.rds',sep='')))  
#     }
#   }
# }

# if(exists("isthiscodl")) {
#   if (isthiscodl=="Y") {
#     result_vector <- sapply(strsplit(rownames(M.seqtab.nochim), "_"), function(x) paste(x[5:(length(x)-2)], collapse = "_"))
#     rownames(M.seqtab.nochim) <- result_vector
#   }
# }

# #=== 2.7. Read contextual data, if set, rename and match =====================================================================

cat('\n\n2.7. Load and prepare your contextual data','\n',file=(file.path(PathToVisuaRAnalysis,paste(VisuaRProjectName,'_Analysis_log.txt',sep=''))),sep='',append=TRUE)
cntxtdtfrmt <- as.data.frame(str_split(contextdatafilepath,"\\."))
cntxtdtfrmt <- cntxtdtfrmt[nrow(cntxtdtfrmt),]

if (contextdata =="N"){                                  # checks whether contextual data was provided and if not it creates a dummy contextdata sheet with "A" for Grouping1 and "1" for Grouping2
  M.contextdata <- base::as.data.frame(base::rownames(M.seqtab.nochim))
  base::row.names(M.contextdata) <- M.contextdata[,1]
  M.contextdata[,1] <- base::rep("A",base::nrow(M.contextdata))
  M.contextdata$VisuaRDummySubGroup <- base::rep(1,base::nrow(M.contextdata))
  base::colnames(M.contextdata)[1]="VisuaRDummyGroup"
  Grouping1="VisuaRDummyGroup"
  Grouping2="VisuaRDummySubGroup"
} else if (contextdata=="Y") {
  if (cntxtdtfrmt == "csv") {
    M.contextdata <- utils::read.delim(file.path(contextdatafilepath),sep=",",header = T,row.names = NULL,check.names = F) # check.names=F. otherwise "-" in columnnames are changed to "."
  } else if (cntxtdtfrmt=="xlsx") {
    M.contextdata <- openxlsx::read.xlsx(file.path(contextdatafilepath),sheet=sheetname,rowNames = F)
    if (excelstartrow > 1) {
      M.contextdata <- M.contextdata[-c(1:(excelstartrow-2)),]
    }
  } else if (cntxtdtfrmt=="txt") {
    M.contextdata <- utils::read.table(file.path(contextdatafilepath),header=T,row.names = NULL)
  }
  M.contextdata[M.contextdata=="NA"] <- NA
  M.contextdata[M.contextdata==""] <- NA
  if(ColumnForMerging=="") {
    M.contextdata <- subset(M.contextdata,!is.na(M.contextdata[,1])) # Creates contextdata df without NAs in merging column
    rownames(M.contextdata) <- M.contextdata[,1]
    M.contextdata=M.contextdata[,-1] 
  } else {
    M.contextdata=subset(M.contextdata,!is.na(M.contextdata[,as.numeric(which(names(M.contextdata)==ColumnForMerging))])) # Creates contextdata df without NAs in merging column
    rownames(M.contextdata) <- M.contextdata[,as.numeric(which(names(M.contextdata)==ColumnForMerging))]
  }
  if(Grouping1=="") {
    M.contextdata$VisuaRDummyGroup=rep("A",nrow(M.contextdata))
    Grouping1="VisuaRDummyGroup"
  }
  if(Grouping2=="") {
    M.contextdata$VisuaRDummySubGroup=rep(1,nrow(M.contextdata))
    Grouping2="VisuaRDummySubGroup"
  }
  # Subset and order M.contextdata to match the row order of M.seqtab.nochim
  M.contextdata <- M.contextdata[match(as.character(rownames(M.seqtab.nochim)),as.character(rownames(M.contextdata))),] # this subsets the contextdata to the samples we also have in the sequencing data, so we can rename them in the next step easily, if we want to
  
  if(any(rownames(M.contextdata)=="NA")) {stop("You did not provide contextual data for all your samples. Please update contextual data, otherwise sample assignment will be screwd.")}
  if(ColumnForAnalysisNames!=""){
    rownames(M.seqtab.nochim) <- M.contextdata[,as.numeric(which(names(M.contextdata)==ColumnForAnalysisNames))]
    rownames(M.contextdata)<-M.contextdata[,as.numeric(which(names(M.contextdata)==ColumnForAnalysisNames))]
  }
} 
M.contextdata[M.contextdata == ""] <- NA
gc()

if(is.numeric(M.contextdata[,Grouping1])){                           # If Grouping1 is a numeric value this adds an A in front of the values.
  M.contextdata[,Grouping1]=paste("A",M.contextdata[,Grouping1],sep="")
  M.contextdata[,Grouping1][M.contextdata[,Grouping1]=='ANA']=NA        # turns created 'ANA's back to 'NA's
}

cat("The following Paramters are available in your mapfile:\n",file=(file.path(PathToVisuaRAnalysis,paste(VisuaRProjectName,'_Analysis_log.txt',sep=''))),sep="",append=TRUE)
cat(colnames(M.contextdata),file=(file.path(PathToVisuaRAnalysis,paste(VisuaRProjectName,'_Analysis_log.txt',sep=''))),sep=", ",append=TRUE)

if (contextdata=='Y'& Grouping1!='VisuaRDummyGroup') { # checks whether your selected Groupings do occur in your contextdata file
  if(Grouping1 %in% colnames(M.contextdata)) {
    cat('\nYour chosen Grouping1 (',Grouping1,') occurs in your contextdata file. Yeay!',file=(file.path(PathToVisuaRAnalysis,paste(VisuaRProjectName,'_Analysis_log.txt',sep=''))),sep="",append=TRUE)
    if(Grouping2 != "VisuaRDummySubGroup") {
      if (Grouping2 %in% colnames(M.contextdata)) {
        cat('\nHea laps, your chosen Grouping2 (',Grouping2,') also occurs in your contextdata file.',file=(file.path(PathToVisuaRAnalysis,paste(VisuaRProjectName,'_Analysis_log.txt',sep=''))),sep="",append=TRUE)
        message('Hea laps, you chose two groupings and both occur in your context data (Grouping1: ',Grouping1,', Grouping2: ',Grouping2,').')
      } else {
        cat('\nOhno! Your Grouping2 (',Grouping2,') does not occur in your contextdata file. Please check spelling.\n',file=(file.path(PathToVisuaRAnalysis,paste(VisuaRProjectName,'_Analysis_log.txt',sep=''))),sep='\t',append=TRUE)
        stop('Ohno! Your Grouping2 (',Grouping2, ') does not occur in your contextdata file. Please check spelling.')
      }
    } else if (Grouping2 == "VisuaRDummySubGroup"){
      cat('\nYou did not choose a second Grouping (Grouping2).\n',file=(file.path(PathToVisuaRAnalysis,paste(VisuaRProjectName,'_Analysis_log.txt',sep=''))),sep='\t',append=TRUE)
    }
  } else {
    if(Grouping2 != "VisuaRDummySubGroup") {
      if(Grouping2 %in% colnames(M.contextdata)){
        cat('\nOhno!: Your Grouping1 (' ,Grouping1,') does not occur in your contextdata file. Please check spelling. Your Grouping 2 (',Grouping2,') is fine though. At least.\n',file=(file.path(PathToVisuaRAnalysis,paste(VisuaRProjectName,'_Analysis_log.txt',sep=''))),sep='\t',append=TRUE)
        stop('Ohno!: Your Grouping1 (' ,Grouping1,') does not occur in your contextdata file. Please check spelling. Your Grouping 2 (',Grouping2,') is fine though. At least.')
      } else {
        cat('\nPaha laps, neither your Grouping1 (' ,Grouping1,') nor your Grouping 2 (',Grouping2,') occur in your contextdata. Aiaiai.\n',file=(file.path(PathToVisuaRAnalysis,paste(VisuaRProjectName,'_Analysis_log.txt',sep=''))),sep='\t',append=TRUE)
        stop('Paha laps, neither your Grouping1 (' ,Grouping1,') nor your Grouping 2 (',Grouping2,') occur in your contextdata. Aiaiai.')
      }
    }
  }
} else if (contextdata=="Y" & Grouping1=="VisuaRDummyGroup") {
  cat('\nContext data was provided but no Grouping chosen, That does not make too much sense, but sure, go ahead.\n',file=(file.path(PathToVisuaRAnalysis,paste(VisuaRProjectName,'_Analysis_log.txt',sep=''))),sep='\t',append=TRUE)
  message('Context data was provided but no Grouping chosen, That does not make too much sense, but sure, go ahead.')
} else if (contextdata=="N") {
  cat('\nNo context data was provided, samples will be analysed but not grouped.\n',file=(file.path(PathToVisuaRAnalysis,paste(VisuaRProjectName,'_Analysis_log.txt',sep=''))),sep='\t',append=TRUE)
  message('No context data was provided, samples will be analysed but not grouped.')
}


gc()


# #=== 2.7. Merges seqtab_nochim and taxonomy ==============================================================================================================================================
infotable <- data.frame(matrix(nrow=0,ncol=20))
colnames(infotable) <- c("Sample number",
                         "Reads_total",
                         "Reads_min",
                         "Reads_1stQu.",
                         "Reads_median",
                         "Reads_mean",
                         "Reads_3rdQu.",
                         "Reads_max",
                         "Reads_sd",
                         "Reads_se",
                         "ASVs_total_dif",
                         "ASVs_total",
                         "ASVs_min",
                         "ASVs_1stQu.",
                         "ASVs_median",
                         "ASVs_mean",
                         "ASVs_3rdQu.",
                         "ASVs_max",
                         "ASVs_sd",
                         "ASVs_se")

M.seq.tax=cbind.data.frame(t(M.seqtab.nochim),M.taxo.noNA)  # creates dataframe with all samples, ASVs and taxonomy. 
if (SaveWholeworkspace=='N') {rm(M.taxo.noNA)}
gc()
ncol.seq.tax=ncol(M.seq.tax)                                # Number of samples + number of taxonomy classes (7)

M.reads=rowSums(M.seqtab.nochim[,])                         # Number of reads per sample

M.ASV.reads=colSums(M.seqtab.nochim[,]) # Number of reads per ASV
M.ASV.reads.df=as.data.frame(M.ASV.reads)
M.ASV.reads.df=cbind('ASV'=rownames(M.ASV.reads.df),M.ASV.reads.df)
rownames(M.ASV.reads.df)=NULL 
colnames(M.ASV.reads.df)[2]='Reads'
data.table::fwrite(M.ASV.reads.df,file=file.path(PathToVisuaRAnalysis,'Alpha_Diversity',paste(VisuaRProjectName,'_ReadsperASV_Original.csv',sep='')),col.names = T,row.names = T)
if (SaveWholeworkspace=='N') {rm(M.ASV.reads.df)}
gc()

sink(file=(file.path(PathToVisuaRAnalysis,paste(VisuaRProjectName,'_Analysis_log.txt',sep=''))),append=TRUE,type='output')
cat('\n\nGeneral Information on sequenced samples.\n',sep=' ')
cat('You started with a total of ',nrow(M.seqtab.nochim),' samples containing a total of ',sum(M.reads),' reads.\n',sep='')
sink()
printinfo(M.reads)

M.differentASVsperSample=rowSums(M.seqtab.nochim[,]!= 0) # creates a table with the number of different ASVs per sample
sink(file=(file.path(PathToVisuaRAnalysis,paste(VisuaRProjectName,'_Analysis_log.txt',sep=''))),append=TRUE,type='output')
cat("These (unfiltered) samples contain on average ",round(mean(M.differentASVsperSample),1),' different ASVs (ASV richness). Overall you have ', ncol(M.seqtab.nochim),' different ASVs.\n',sep='')
sink()
printinfo(M.differentASVsperSample)
sink(file=(file.path(PathToVisuaRAnalysis,paste(VisuaRProjectName,'_Analysis_log.txt',sep=''))),append=TRUE,type='output')
cat("Be aware that the total value reported above is summed up over all samples, several of the ASVs are hence counted multiple times. Overall, the (unfiltered) samples contain a total of ",ncol(M.seqtab.nochim),' different ASVs (ASV richness).\n',sep='')
sink()

infotable <- fill.info(df.reads=M.reads,df.ASVs =M.differentASVsperSample,df.seqtab=M.seq.tax,stepname="Start",infotable=infotable)

sink(file=(file.path(PathToVisuaRAnalysis,paste(VisuaRProjectName,'_Analysis_log.txt',sep=''))),append=TRUE,type='output')
cat("You're original ASVs occur on average ",round(mean(M.ASV.reads),0),' times.\n',sep='')
sink()  
printinfo(M.ASV.reads)
if (SaveWholeworkspace=='N') {rm(M.ASV.reads)}
gc()

M.reads.df=as.data.frame(M.reads) 
M.differentASVsperSample=as.data.frame(M.differentASVsperSample)
M.differentASVsperSample=cbind(row.names(M.seqtab.nochim),data.frame(M.reads.df,row.names = NULL),data.frame(M.differentASVsperSample,row.names = NULL))
colnames(M.differentASVsperSample)[1]='Sample Name'
colnames(M.differentASVsperSample)[2]='Total Reads'
colnames(M.differentASVsperSample)[3]='Observed ASVs - Richness'
data.table::fwrite(M.differentASVsperSample,file=file.path(PathToVisuaRAnalysis,'Alpha_Diversity',paste(VisuaRProjectName,'_ReadsandASVsperSample_Original.csv',sep='')),col.names = T,row.names = T)
if (SaveWholeworkspace=='N') {rm(M.differentASVsperSample,M.reads)}
gc()

# #=== 2.8. Contamination control ==========================================================================================================================
cat('\n2.8. Excludes samples based on chosen blanks.',file=(file.path(PathToVisuaRAnalysis,paste(VisuaRProjectName,'_Analysis_log.txt',sep=''))),sep='',append=TRUE)

if (removeContaminants != '') {
  dir.create(file.path(PathToVisuaRAnalysis,'ContaminationControl'))
  
  # creates table with only samples which have information in the chosen column (= blanks)
  blankSamplenames <- rownames(M.contextdata[!is.na(M.contextdata[,which(names(M.contextdata)==ColumnWithBlankGrouping)]),]) 
  
  # cat('\nASVs will be excluded or their mean subtracted or decontaminated using decontam::isNotContaminant() based on the provided blanks: ',paste0(blankSamplenames,sep=","),file=(file.path(PathToVisuaRAnalysis,paste(VisuaRProjectName,'_Analysis_log.txt',sep=''))),sep='',append=T)
  
  # # create list of unique blank groups as found in your contextual data
  blanksamplegroups.blanks <- sort(unique(na.omit(M.contextdata[,which(names(M.contextdata)==ColumnWithBlankGrouping)])))
  # # unique groupings in our contextdata
  blanksamplegroups.samples <- sort(unique(na.omit(M.contextdata[,which(names(M.contextdata)==ColumnWithBlankGroupingSamples)]))) 
  
  if(all(blanksamplegroups.blanks==blanksamplegroups.samples)) {} else {stop("You're blank groups and sample blank groups do not match. Please revise your contextual data.")}
  
  # #  create a list with sample names belonging to each blank group and also a one element with samples not belonging to any blank group
  blanksamplegroups.samplenames.list <- vector("list",length(blanksamplegroups.samples)+1)
  for (i in 1:(length(blanksamplegroups.samples))) {
    blanksamplegroups.samplenames.list[[i]] <-rownames(M.contextdata[(M.contextdata[, which(names(M.contextdata) == ColumnWithBlankGroupingSamples)] == blanksamplegroups.samples[i]) & !is.na(M.contextdata[, which(names(M.contextdata) == ColumnWithBlankGroupingSamples)]),])
  }
  if((length(blanksamplegroups.samples)+1)==length(sort(unique((M.contextdata[,which(names(M.contextdata)==ColumnWithBlankGroupingSamples)]))))) { # in case we have NA in the blank group column for the samples that would be true
    blanksamplegroups.samplenames.list[[length(blanksamplegroups.samplenames.list)]] <- rownames(M.contextdata[is.na(M.contextdata[, which(names(M.contextdata) == ColumnWithBlankGroupingSamples)]),]) 
    cat('\nThe samples ',paste0(as.character(blanksamplegroups.samplenames.list[[length(blanksamplegroups.samplenames.list)]]),sep=",")," have no associated blank and no ASVs will be removed or subtracted from them.",file=(file.path(PathToVisuaRAnalysis,paste(VisuaRProjectName,'_Analysis_log.txt',sep=''))),sep='',append=T)
  } else {
    cat('\nAll samples have blanks associated to them.',file=(file.path(PathToVisuaRAnalysis,paste(VisuaRProjectName,'_Analysis_log.txt',sep=''))),sep='',append=T)
  }
  # #  create a list with blank names belonging to each blank group
  blanksamplegroups.blanknames.list <- vector("list",length(blanksamplegroups.blanks))
  for (i in 1:length(blanksamplegroups.blanks)) {
    blanksamplegroups.blanknames.list[[i]] <-rownames(M.contextdata[(M.contextdata[, which(names(M.contextdata) == ColumnWithBlankGrouping)] == blanksamplegroups.blanks[i]) & !is.na(M.contextdata[, which(names(M.contextdata) == ColumnWithBlankGrouping)]),])
  }
  
  if(normalizeMseqtax=="Y" & removeContaminants != "decontam") {
    M.seq.tax.save <- M.seq.tax
    M.seq.tax.colsums <- colSums(M.seq.tax[,1:(ncol(M.seq.tax)-ncol.taxo.noNA)]) # number of reads per sample, this creates NA in case the number of reads is 0
    M.seq.tax[,1:(ncol(M.seq.tax)-ncol.taxo.noNA)]<- t(t(M.seq.tax[,1:(ncol(M.seq.tax)-ncol.taxo.noNA)])/M.seq.tax.colsums) # number of reads per asv devided by number of reads in the sample to normalize the read counts per sample to 1
    M.seq.tax[is.na(M.seq.tax)] <- 0     # remove introduced NAs
  }
}

if (removeContaminants=='') {
  cat('\nNo ASVs were excluded/subtracted as no blank samples were chosen.',file=(file.path(PathToVisuaRAnalysis,paste(VisuaRProjectName,'_Analysis_log.txt',sep=''))),sep='',append=T)
  M.seq.tax.subset=M.seq.tax
  
} else if (removeContaminants=="remove"|removeContaminants=="subtract-average"|removeContaminants=="subtract-max"|(removeContaminants=="decontam" & decontam_isnotcontaminant=="Y")){
 
  
  if (removeContaminants!="decontam") {
    # # create indices of ASVs which will be excluded/subtracted (TRUE: needs to be removed/subtracted. FALSE: can stay.)
    indices.blank.ASVs <- vector("list", length(blanksamplegroups.blanks))
    # # for ASVs being excluded/subtracted even if they only occur once in one of the blanks
    for (i in 1:length(blanksamplegroups.blanks)) {
      
      # #  in case we have only one blank
      if(!is.numeric(ncol(M.seq.tax[, blanksamplegroups.blanknames.list[[i]]]))){
        if (stat.test=="N") {
          indices.blank.ASVs[[i]] <- M.seq.tax[, blanksamplegroups.blanknames.list[[i]]] > ASVcountInBlanksToExclude
        } else if(stat.test=="Y") {
          blanks <- as.data.frame(M.seq.tax[, blanksamplegroups.blanknames.list[[i]]]);colnames(blanks) <- blanksamplegroups.blanknames.list[[i]] # extract all the blanks from blank group i from M.seq.tax
          samples <- M.seq.tax[, blanksamplegroups.samplenames.list[[i]]] # extract all the samples from sample group i (belongs to blank group i) from M.seq.tax
          samples <- samples[, !names(samples) %in% names(blanks)] # gets rid of the blanks in the sample list
          # # goes through each row (ASVs and tests whether samples are greater than blanks, if they are indices.blank.ASVs is set to FALSE, if not to TRUE) -> "FALSE"=no contaminant, "TRUE"=contaminant
          test_results <- logical(nrow(blanks)) # FALSE for ASVs that are not considered contaminants
          for (j in 1:nrow(blanks)) {
            if (testtouse == "w") {
              testresult<-stats::wilcox.test(x=as.numeric(samples[j, ]),y=as.numeric(blanks[j, ]), alternative = "greater") # same as t-test but not assuming normal distribution
            } 
            if (testresult$p.value <= signiflevel) {
              test_results[j] <- FALSE # read counts are statistically more in the samples compared to the blanks
            } else {
              test_results[j] <- TRUE # read counts are either similar or statistically more in the blanks compared to the samples
            }
          }
          indices.blank.ASVs[[i]] <- test_results
        }
        # # in case we have more than one blank 
      } else {
        if (stat.test=="N") {
          indices.blank.ASVs[[i]] <- rowSums(M.seq.tax[, blanksamplegroups.blanknames.list[[i]]])/length(blanksamplegroups.blanknames.list[[i]]) > ASVcountInBlanksToExclude
        } else if(stat.test=="Y") {
          blanks <- M.seq.tax[, blanksamplegroups.blanknames.list[[i]]] # extract all the blanks from blank group i from M.seq.tax
          samples <- M.seq.tax[, blanksamplegroups.samplenames.list[[i]]] # extract all the samples from sample group i (belongs to blank group i) from M.seq.tax
          samples <- samples[, !names(samples) %in% names(blanks)] # gets rid of the blanks in the sample list
          # # goes through each row (ASVs and tests whether samples are greater than blanks, if they are indices.blank.ASVs is set to FALSE, if not to TRUE) -> "FALSE"=no contaminant, "TRUE"=contaminant
          test_results <- logical(nrow(blanks)) # FALSE for ASVs that are not considered contaminants
          for (j in 1:nrow(blanks)) {
            if (testtouse == "w") {
              testresult<-stats::wilcox.test(x=as.numeric(samples[j, ]),y=as.numeric(blanks[j, ]), alternative = "greater") # same as t-test but not assuming normal distribution
            } 
            if (testresult$p.value <= signiflevel) {
              test_results[j] <- FALSE # read counts are statistically more in the samples compared to the blanks
            } else {
              test_results[j] <- TRUE # read counts are either similar or statistically more in the blanks compared to the samples
            }
          }
          indices.blank.ASVs[[i]] <- test_results
        }
      }
    }
  }
  gc()
  # # actual removal/subtraction of contaminants
  if(removeContaminants=="remove" & VisuaRContaminants == "N"){
    M.seq.tax.subset <- M.seq.tax
    for (i in 1:length(blanksamplegroups.blanks)) { # goes through all blank groups
      # # checks where in M.seq.tax.subset "indices.blank.ASVs" is TRUE (rows) and where blanksamplegroups.samplenames.list (columns) fits to the current group and sets the ASV count to zero 
      M.seq.tax.subset[as.logical(indices.blank.ASVs[[i]]), blanksamplegroups.samplenames.list[[i]]] <- 0
    }
  } else if (removeContaminants=="remove" & VisuaRContaminants == "Y") {
    M.seq.tax.subset <- M.seq.tax
    for (i in 1:length(blanksamplegroups.blanks)) { # goes through all blank groups
      # # checks where in M.seq.tax.subset "indices.blank.ASVs" is TRUE (rows) and where blanksamplegroups.samplenames.list (columns) fits to the current group and sets the ASV count to zero 
      M.seq.tax.subset[!as.logical(indices.blank.ASVs[[i]]), blanksamplegroups.samplenames.list[[i]]] <- 0
    }
  } else if ((removeContaminants=="subtract-average"|removeContaminants=="subtract-max") & VisuaRContaminants == "N"){
    M.seq.tax.subset <- M.seq.tax
    for (i in 1:length(blanksamplegroups.blanks)) {
      # #  in case we have only one blank
      if(!is.numeric(ncol(M.seq.tax[, blanksamplegroups.blanknames.list[[i]]]))){
        average <- (M.seq.tax[, blanksamplegroups.blanknames.list[[i]]])
      } else  if (removeContaminants=="subtract-average") {
        # Calculate the average of selected columns in the data frame
        average <- rowMeans(M.seq.tax[, blanksamplegroups.blanknames.list[[i]]])
      } else if (removeContaminants=="subtract-max") {
        average <- apply(M.seq.tax[, blanksamplegroups.blanknames.list[[i]]], 1, max) # its called average but actually gives the maximum value 
      }
      # Subtract the average/maximum from columns with samples and blanks belonging to the group based on the logical vector giving the rowindices
      M.seq.tax.subset[as.logical(indices.blank.ASVs[[i]]), c(blanksamplegroups.samplenames.list[[i]])] <- M.seq.tax.subset[as.logical(indices.blank.ASVs[[i]]), c(blanksamplegroups.samplenames.list[[i]])] - average[as.logical(indices.blank.ASVs[[i]])]
      
      if (normalizeMseqtax=="N") {
        # Round the updated values to whole digits
        M.seq.tax.subset[,1:(ncol(M.seq.tax)-ncol.taxo.noNA)] <- round(M.seq.tax.subset[,1:(ncol(M.seq.tax)-ncol.taxo.noNA)])
        # Set negative values to zero
        M.seq.tax.subset[M.seq.tax.subset < 0] <- 0
      }
    }
  } else if ((removeContaminants=="subtract-average"|removeContaminants=="subtract-max") & VisuaRContaminants == "Y"){
    M.seq.tax.subset <- M.seq.tax
    for (i in 1:length(blanksamplegroups.blanks)) {
      # #  in case we have only one blank
      if(!is.numeric(ncol(M.seq.tax[, blanksamplegroups.blanknames.list[[i]]]))){
        average <- (M.seq.tax[, blanksamplegroups.blanknames.list[[i]]])
      } else  if (removeContaminants=="subtract-average") {
        # Calculate the average of selected columns in the data frame
        average <- rowMeans(M.seq.tax[, blanksamplegroups.blanknames.list[[i]]])
      } else if (removeContaminants=="subtract-max") {
        average <- apply(M.seq.tax[, blanksamplegroups.blanknames.list[[i]]], 1, max) # its called average but actually gives the maximum value 
      }
      # # set all ASVs which do not occur in blanks to 0
      M.seq.tax.subset[!as.logical(indices.blank.ASVs[[i]]), blanksamplegroups.samplenames.list[[i]]] <- 0
      
      # Subtract the average/maximum from columns with samples and blanks belonging to the group based on the logical vector giving the rowindices
      # M.seq.tax.subset[as.logical(indices.blank.ASVs[[i]]), c(blanksamplegroups.samplenames.list[[i]])] <- M.seq.tax.subset[as.logical(indices.blank.ASVs[[i]]), c(blanksamplegroups.samplenames.list[[i]])] - average[as.logical(indices.blank.ASVs[[i]])]
      
      if (normalizeMseqtax=="N") {
        # Round the updated values to whole digits
        M.seq.tax.subset[,1:(ncol(M.seq.tax)-ncol.taxo.noNA)] <- round(M.seq.tax.subset[,1:(ncol(M.seq.tax)-ncol.taxo.noNA)])
        # Set negative values to zero
        M.seq.tax.subset[M.seq.tax.subset < 0] <- 0
      }
    }
  } 
  gc()
  if (normalizeMseqtax=="Y") {
    # # multiplies the values again with the total readcounts we had in our samples to get back to our read count table
    M.seq.tax.subset[,1:(ncol(M.seq.tax.subset)-ncol.taxo.noNA)] <- t(t(M.seq.tax.subset[,1:(ncol(M.seq.tax.subset)-ncol.taxo.noNA)])*M.seq.tax.colsums)
    M.seq.tax.subset[,1:(ncol(M.seq.tax)-ncol.taxo.noNA)] <- round(M.seq.tax.subset[,1:(ncol(M.seq.tax)-ncol.taxo.noNA)])
    # Set negative values to zero
    M.seq.tax.subset[M.seq.tax.subset < 0] <- 0
    
    # # transforms M.seq.tax back to original table
    M.seq.tax <- M.seq.tax.save
    rm(M.seq.tax.save)
  }
  
  
  if (removeContaminants!="decontam") {
    # # creates an ASV by Sample&Taxonomy df with all ASVs which now do not occur in any of the samples anymore
    M.seq.tax.subset.contam.ASVs.excluded=M.seq.tax.subset[rowSums(M.seq.tax.subset[,1:(ncol(M.seq.tax.subset)-ncol.taxo.noNA)])==0,]
    # creates an ASV by Sample&Taxonomy df with only ASVs which are still present in the samples
    M.seq.tax.subset<-M.seq.tax.subset[rowSums(M.seq.tax.subset[,1:(ncol(M.seq.tax.subset)-ncol.taxo.noNA)])>0,] 
  }
} 

if (removeContaminants=="decontam") {
  # # excludes ASVs based on the prevalence method in the package decontam
  # # creates sample by asv table without taxonomy
  M.seq.tax.temp<-as.data.frame(t(M.seq.tax[,1:(ncol(M.seq.tax)-ncol.taxo.noNA)]))
  
  # # selects the column with F/T, defining which samples are blanks and DNA concentration column if provided
  if(IncludeDNAconc=="Y") {
    M.contextdata.temp <- M.contextdata[, c(which(names(M.contextdata) == ColumnWithBlankGroupingSamples),which(names(M.contextdata) == columnwithDNAconcentrations), which(names(M.contextdata) == columnfordecontam))]
  } else if (IncludeDNAconc=="N") {
    M.contextdata.temp <- M.contextdata[, c(which(names(M.contextdata) == ColumnWithBlankGroupingSamples),which(names(M.contextdata) == columnfordecontam))]
  }
  
  # # merges seqtab and contextual data column
  M.seq.tax.temp <- merge(M.seq.tax.temp,M.contextdata.temp,by=0,all.x = T,all.y=F,sort=F)
  rownames(M.seq.tax.temp)<-M.seq.tax.temp[,1];M.seq.tax.temp<-M.seq.tax.temp[,-1]
  
  # # creates table with only samples which have information in the chosen columns (last three ones)
  M.seq.tax.temp.noNA=subset(M.seq.tax.temp,!is.na(M.seq.tax.temp[,ncol(M.seq.tax.temp)])&!is.na(M.seq.tax.temp[,ncol(M.seq.tax.temp)-1])&!is.na(M.seq.tax.temp[,ncol(M.seq.tax.temp)-2]))
  
  if (decontam_isnotcontaminant == "N") {
    # # extracts vector from this table with FALSE/TRUE (no blank/blank)
    vector_for_decontam <- M.seq.tax.temp.noNA[,which(names(M.seq.tax.temp.noNA)==columnfordecontam)]
    batches_for_decontam <- M.seq.tax.temp.noNA[,which(names(M.seq.tax.temp.noNA) == ColumnWithBlankGroupingSamples)]
    if (IncludeDNAconc=="Y") {
      # # dna vector for decontam if provided
      dna_for_decontam <- M.seq.tax.temp.noNA[,which(names(M.seq.tax.temp.noNA)==columnwithDNAconcentrations)]
      # # and removes it from the seqtab file (no taxonomy here)
      M.seq.tax.temp.noNA <- M.seq.tax.temp.noNA[,-c(which(names(M.seq.tax.temp.noNA) == ColumnWithBlankGroupingSamples),which(names(M.seq.tax.temp.noNA)==columnfordecontam),which(names(M.seq.tax.temp.noNA)==columnwithDNAconcentrations))]
    } else if (IncludeDNAconc=="N") {
      M.seq.tax.temp.noNA <- M.seq.tax.temp.noNA[,-c(which(names(M.seq.tax.temp.noNA) == ColumnWithBlankGroupingSamples),which(names(M.seq.tax.temp.noNA)==columnfordecontam))]
    }
    M.seq.tax.temp.noNA<-as.matrix(M.seq.tax.temp.noNA)
  }
  
  
  
  # # actual contaminant classification. this creates a table with frequency, prevalence, p.freq, p.prev, p and classification as contaminant
  if (decontam_isnotcontaminant=="N" & IncludeDNAconc == "Y") {
    contam_df <- isContaminant(M.seq.tax.temp.noNA,neg=vector_for_decontam,method=decontam.method,conc=dna_for_decontam,batch=batches_for_decontam)
    # # number of ASVs classified as contaminants (TRUE)
    table(contam_df$contaminant)
    # # indices of contaminant ASVs
    contam_asvs <- row.names(contam_df[contam_df$contaminant == TRUE, ])
  } else if (decontam_isnotcontaminant=="N" & IncludeDNAconc == "N") {
    contam_df <- isContaminant(M.seq.tax.temp.noNA,neg=vector_for_decontam,method=decontam.method)
    # # number of ASVs classified as contaminants (TRUE)
    table(contam_df$contaminant)
    # # indices of contaminant ASVs
    contam_asvs <- row.names(contam_df[contam_df$contaminant == TRUE, ])
  } else if (decontam_isnotcontaminant=="Y") {
    M.seq.tax.subset.groups <- vector("list",length(blanksamplegroups.samplenames.list))
    overview.contam.groups <- data.frame(ncol=3,nrow=0)
    colnames(overview.contam.groups) <- c("contaminant","notcontaminant")
    for (i in 1:length(blanksamplegroups.blanks)) {
      # # takes the samples belonging to the first blank group from the M.seq.tax.temp.noNA
      M.seq.tax.temp.noNA.batch <- M.seq.tax.temp.noNA[row.names(M.seq.tax.temp.noNA) %in% blanksamplegroups.samplenames.list[[i]], ]
      # # takes vector indicating which samples are blanks and which ones are samples from this df
      vector_for_decontam <- M.seq.tax.temp.noNA.batch[,which(names(M.seq.tax.temp.noNA.batch)==columnfordecontam)]
      # # removes columns with information on blank groups and blank identification
      M.seq.tax.temp.noNA.batch <- M.seq.tax.temp.noNA.batch[,-c(which(names(M.seq.tax.temp.noNA.batch) == ColumnWithBlankGroupingSamples),which(names(M.seq.tax.temp.noNA.batch)==columnfordecontam))]
      M.seq.tax.temp.noNA.batch<-as.matrix(M.seq.tax.temp.noNA.batch)
      # # calculates which ASVs are contaminants/not contaminants
      notcontam_df <- isNotContaminant(M.seq.tax.temp.noNA.batch,neg=vector_for_decontam,method="prevalence",detailed=TRUE)
      
      # # number of ASVs classified as contaminants (FALSE), TRUE indicates non-contaminants
      overview.contam.groups[i,1] <- table(notcontam_df$not.contaminant)[1]
      overview.contam.groups[i,2] <- table(notcontam_df$not.contaminant)[2]
      # # indices of contaminant ASVs
      contam_asvs <- row.names(notcontam_df[notcontam_df$not.contaminant == FALSE, ])
      
      # # extracts taxonomy of contaminant ASVs 
      contam_asvs_tax<-ASV.mapfile[ASV.mapfile[,1] %in% contam_asvs, ]
      data.table::fwrite(contam_asvs_tax,file=file.path(PathToVisuaRAnalysis,'ContaminationControl',paste(VisuaRProjectName,"_Group_",blanksamplegroups.blanks[i],"_contaminantASVs.csv",sep="")),col.names = T,row.names = T)
      
      if (VisuaRContaminants == "N") {
        # # sets ASVs categorized as contaminants to 0 
        M.seq.tax.temp.noNA.batch <- t(M.seq.tax.temp.noNA.batch)
        M.seq.tax.temp.noNA.batch[rownames(M.seq.tax.temp.noNA.batch) %in% contam_asvs, ] <- 0
        M.seq.tax.subset.groups[[i]] <- M.seq.tax.temp.noNA.batch
      } else if (VisuaRContaminants == "Y") {
        # # sets ASVs NOT categorized as contaminants to 0 
        M.seq.tax.temp.noNA.batch <- t(M.seq.tax.temp.noNA.batch)
        M.seq.tax.temp.noNA.batch[!(rownames(M.seq.tax.temp.noNA.batch) %in% contam_asvs), ] <- 0
        M.seq.tax.subset.groups[[i]] <- M.seq.tax.temp.noNA.batch
      }
      rm(M.seq.tax.temp.noNA.batch,contam_asvs_tax)
    }
    if (length(blanksamplegroups.samplenames.list[[length(blanksamplegroups.samplenames.list)]])!=0) { # # in case that there are samples with no blank associated
      M.seq.tax.subset.groups[[length(blanksamplegroups.samplenames.list)]]  <- M.seq.tax.temp.noNA[row.names(M.seq.tax.temp.noNA) %in% blanksamplegroups.samplenames.list[[length(blanksamplegroups.samplenames.list)]], ]
    } else if (length(M.seq.tax.subset.groups[[length(blanksamplegroups.samplenames.list)]])==0) { # just to doublecheck that the element really is empty
      M.seq.tax.subset.groups <- M.seq.tax.subset.groups[-length(M.seq.tax.subset.groups)]
    }
    
    # # prepares M.seq.tax.subset to be filled with the new, contamination corrected data
    M.seq.tax.subset <- data.frame(matrix(0, nrow = nrow(M.seq.tax)))
    for (i in 1:length(M.seq.tax.subset.groups)) {
      M.seq.tax.subset <- cbind(M.seq.tax.subset,M.seq.tax.subset.groups[[i]] )
    }
    M.seq.tax.subset <- M.seq.tax.subset[,-1]
    
    # # adds samples which had no blank grouping available and taxonomy back to df
    M.seq.tax.subset <- cbind(M.seq.tax.subset, M.seq.tax[setdiff(names(M.seq.tax),names(M.seq.tax.subset))])
    
    # # remove ASVs which are no longer present in any of the samples 
    M.seq.tax.subset <- M.seq.tax.subset[rowSums(M.seq.tax.subset[,1:(ncol(M.seq.tax.subset)-ncol.taxo.noNA)])!=0,] # excludes ASVs not longer present in the samples
    
    # # bring M.seq.tax.subset columns in the same order as M.seq.tax columns
    M.seq.tax.subset <- M.seq.tax.subset[names(M.seq.tax)]
  }
  
  if (decontam_isnotcontaminant != "Y") {
    # # extracts taxonomy of contaminant ASVs 
    contam_asvs_tax<-ASV.mapfile[ASV.mapfile[,1] %in% contam_asvs, ]
    data.table::fwrite(contam_asvs_tax,file=file.path(PathToVisuaRAnalysis,'ContaminationControl',paste(VisuaRProjectName,"_contaminantASVs.csv",sep="")),col.names = T,row.names = T)
    
    cat('\nThe following ',nrow(contam_asvs_tax),' contaminant ASVs, out of ',nrow(contam_df),' ASVs have been identified with decontam in your blanks and will be excluded from the analysis. Exact sequences can be found in the file ',paste(VisuaRProjectName,"_contaminantASVs.txt",sep=""),'\n',file=(file.path(PathToVisuaRAnalysis,paste(VisuaRProjectName,'_Analysis_log.txt',sep=''))),sep='',append=T)
    cat(contam_asvs_tax$Species,file=(file.path(PathToVisuaRAnalysis,paste(VisuaRProjectName,'_Analysis_log.txt',sep=''))),sep='\n',append=T)
    if(IncludeDNAconc=="Y") {
      cat('\nThe measured DNA concentrations of blanks and samples were included when calculating for contaminant ASVs.\n',file=(file.path(PathToVisuaRAnalysis,paste(VisuaRProjectName,'_Analysis_log.txt',sep=''))),sep='',append=T)
    }
    
    if(VisuaRContaminants =="N") {
      # # removes ASVs categorized as contaminants
      M.seq.tax.subset<-M.seq.tax[!row.names(M.seq.tax) %in% contam_asvs,]
    } else if (VisuaRContaminants =="Y") {
      # # removes ASVs categorized NOT as contaminants
      M.seq.tax.subset<-M.seq.tax[row.names(M.seq.tax) %in% contam_asvs,]
    }
    rm(M.seq.tax.temp,M.contextdata.temp,M.seq.tax.temp.noNA)
  }
}  
gc()
if (removeContaminants!='') {
  ##### to plot percentages of contamination
  # #  returns named num with percentages of kept reads per sample
  percentage.noncontams.reads <- (colSums(M.seq.tax.subset[,1:(ncol(M.seq.tax.subset)-ncol.taxo.noNA)])/colSums(M.seq.tax[,1:(ncol(M.seq.tax)-ncol.taxo.noNA)]))*100
  # # returns named num with percentages of removed ASVs per sample
  percentage.noncontam.ASVs <- ((apply(M.seq.tax.subset[,1:(ncol(M.seq.tax.subset)-ncol.taxo.noNA)] != 0, 2, sum))/(apply(M.seq.tax[,1:(ncol(M.seq.tax)-ncol.taxo.noNA)] != 0, 2, sum)))*100
  
  # # in case contextual data has more samples than 
  M.contextdata <- M.contextdata[rownames(M.contextdata) %in% colnames(M.seq.tax.subset),]
  # # to order the columns of M.contextdata in the same way as M.seq.tax.subset
  M.contextdata <- M.contextdata[colnames(M.seq.tax.subset[,1:(ncol(M.seq.tax.subset)-ncol.taxo.noNA)]), ]  
  
  # # df for plots with last column the blank groupings
  percentages.noncontams <- cbind(percentage.noncontam.ASVs,percentage.noncontams.reads,M.contextdata[,which(names(M.contextdata)==ColumnWithBlankGroupingSamples)])
  percentages.noncontams <- as.data.frame(percentages.noncontams)
  percentages.noncontams[,1] <- as.numeric(percentages.noncontams[,1])
  percentages.noncontams[,2] <- as.numeric(percentages.noncontams[,2])
  percentages.noncontams <- cbind(rownames(percentages.noncontams),percentages.noncontams)
  colnames(percentages.noncontams) <- c("Sample","PercentNoContamASVs","PercentNoContamReads","BlankGroup")
  data.table::fwrite(percentages.noncontams,file=file.path(PathToVisuaRAnalysis,'ContaminationControl',paste(VisuaRProjectName,"_ASVsAndReadsAfterDecontamination.csv",sep="")),col.names = T,row.names = T)
  
  # # only take blanks
  percentages.nocontams.blanks <- percentages.noncontams[(rownames(percentages.noncontams) %in% blankSamplenames), ]; percentages.nocontams.blanks <- percentages.nocontams.blanks[,-1]
  # # create grouped averages
  group_percASVAvg <- aggregate(percentages.nocontams.blanks$PercentNoContamASVs, list(percentages.nocontams.blanks$BlankGroup), FUN=mean) 
  group_percReadsAvg <- aggregate(percentages.nocontams.blanks$PercentNoContamReads, list(percentages.nocontams.blanks$BlankGroup), FUN=mean) 
  groupedaverages <- cbind(group_percASVAvg[,2],group_percReadsAvg[,2],group_percASVAvg[,1]); colnames(groupedaverages) <- colnames(percentages.nocontams.blanks)
  # # add row with average and grouped average
  percentages.nocontams.blanks[nrow(percentages.nocontams.blanks)+1,] <- c(colMeans(percentages.nocontams.blanks[,c(1,2)]),"All"); rownames(percentages.nocontams.blanks)[nrow(percentages.nocontams.blanks)] <- "Average"
  percentages.nocontams.blanks <- rbind(percentages.nocontams.blanks,groupedaverages)
  data.table::fwrite(percentages.nocontams.blanks,file=file.path(PathToVisuaRAnalysis,'ContaminationControl',paste(VisuaRProjectName,"_ASVsAndReadsAfterDecontamination-blanksonly.csv",sep="")),col.names = T,row.names = T)
  
  correctedsamples <- unique(unlist(blanksamplegroups.samplenames.list))
  # # only take corrected samples
  percentages.nocontams.samples <- percentages.noncontams[!(rownames(percentages.noncontams) %in% blankSamplenames) & rownames(percentages.noncontams) %in% correctedsamples, ]; percentages.nocontams.samples <- percentages.nocontams.samples[,-1]
  # # create grouped averages
  group_percASVAvg <- aggregate(percentages.nocontams.samples$PercentNoContamASVs, list(percentages.nocontams.samples$BlankGroup), FUN=mean) 
  group_percReadsAvg <- aggregate(percentages.nocontams.samples$PercentNoContamReads, list(percentages.nocontams.samples$BlankGroup), FUN=mean) 
  groupedaverages <- cbind(group_percASVAvg[,2],group_percReadsAvg[,2],group_percASVAvg[,1]); colnames(groupedaverages) <- colnames(percentages.nocontams.samples)
  # # add row with average and grouped average
  percentages.nocontams.samples[nrow(percentages.nocontams.samples)+1,] <- c(colMeans(percentages.nocontams.samples[,c(1,2)]),"All"); rownames(percentages.nocontams.samples)[nrow(percentages.nocontams.samples)] <- "Average"
  percentages.nocontams.samples <- rbind(percentages.nocontams.samples,groupedaverages)
  data.table::fwrite(percentages.nocontams.samples,file=file.path(PathToVisuaRAnalysis,'ContaminationControl',paste(VisuaRProjectName,"_ASVsAndReadsAfterDecontamination-samplesonly.csv",sep="")),col.names = T,row.names = T)
  
  # # only take uncorrected samples
  percentages.nocontams.samples.uncor <- percentages.noncontams[!(rownames(percentages.noncontams) %in% correctedsamples), ]; percentages.nocontams.samples.uncor <- percentages.nocontams.samples.uncor[,-1]
  data.table::fwrite(percentages.nocontams.samples.uncor,file=file.path(PathToVisuaRAnalysis,'ContaminationControl',paste(VisuaRProjectName,"_ASVsAndReadsAfterDecontamination-samplesonly-uncor.csv",sep="")),col.names = T,row.names = T)
  
  # Convert group column to factor
  percentages.noncontams$BlankGroup <- factor(percentages.noncontams$BlankGroup)
  # Reorder the levels of Sample based on Color
  percentages.noncontams$Sample <- factor(percentages.noncontams$Sample, levels = percentages.noncontams$Sample[order(percentages.noncontams$BlankGroup)])
  
  pdf(file.path(PathToVisuaRAnalysis,'ContaminationControl',paste(VisuaRProjectName,"_percentASVsAfterDecontamination.pdf",sep="")),height=7.5,width=nrow(percentages.noncontams)/5,useDingbats=F) 
  perc.asv.nocontam <- ggplot(percentages.noncontams, aes(x = Sample, y = PercentNoContamASVs, fill = BlankGroup)) +
    geom_bar(stat = "identity",width=0.7) +
    labs(x = "Sample", y = "% ASVs left after decontamination", title=paste(VisuaRProjectName," - decontamination - ",removeContaminants,sep="")) +
    scale_y_continuous(
      limits = c(0, 100),  # Set y-axis limits
      breaks = seq(0, 100, by = 10),  # Major tick marks every 10
      minor_breaks = seq(0, 100, by = 5),  # Minor tick marks every 5
      expand = c(0, 0)  # Remove extra space around the plot
    ) +
    theme(axis.text = element_text(colour="black"),
          legend.title = element_blank(),
          panel.grid.major.x = element_blank(),
          axis.text.x=element_text(angle=90,colour="black",size=14,vjust=0.5,hjust=1), #,vjust=-0.1
          axis.text.y=element_text(colour="black",size=14),
          axis.title = element_text(colour="black",size=16),
          axis.line=element_line(colour="black"),
          panel.background = element_blank(),
          panel.grid.major.y = element_line(colour="grey",size=0.5),
          panel.grid.minor.y = element_line(colour="grey",size = 0.25),
          panel.border = element_rect(fill=NA,colour = "black",size=0.5),
          legend.box.background = element_rect(colour="black"),
          legend.box.margin = margin(0.18,0.1,0.1,0.5),
          legend.key=element_rect(fill="white"),
          legend.text=element_text(size=14),
          axis.ticks=element_line(colour = "black"))
  print(perc.asv.nocontam)
  dev.off()
  
  pdf(file.path(PathToVisuaRAnalysis,'ContaminationControl',paste(VisuaRProjectName,"_percentReadsAfterDecontamination.pdf",sep="")),height=7.5,width=nrow(percentages.noncontams)/5,useDingbats=F) 
  perc.reads.noncontam <- ggplot(percentages.noncontams, aes(x = Sample, y = PercentNoContamReads, fill = BlankGroup)) +
    geom_bar(stat = "identity",width=0.7) +
    labs(x = "Sample", y = "% Reads left after decontamination", title=paste(VisuaRProjectName," - decontamination - ",removeContaminants,sep="")) +
    scale_y_continuous(
      limits = c(0, 100),  # Set y-axis limits
      breaks = seq(0, 100, by = 10),  # Major tick marks every 10
      minor_breaks = seq(0, 100, by = 5),  # Minor tick marks every 5
      expand = c(0, 0)  # Remove extra space around the plot
    ) +
    theme(axis.text = element_text(colour="black"),
          legend.title = element_blank(),
          panel.grid.major.x = element_blank(),
          axis.text.x=element_text(angle=90,colour="black",size=14,vjust=0.5,hjust=1), #,vjust=-0.1
          axis.text.y=element_text(colour="black",size=14),
          axis.title = element_text(colour="black",size=16),
          axis.line=element_line(colour="black"),
          panel.background = element_blank(),
          panel.grid.major.y = element_line(colour="grey",size=0.5),
          panel.grid.minor.y = element_line(colour="grey",size = 0.25),
          panel.border = element_rect(fill=NA,colour = "black",size=0.5),
          legend.box.background = element_rect(colour="black"),
          legend.box.margin = margin(0.18,0.1,0.1,0.5),
          legend.key=element_rect(fill="white"),
          legend.text=element_text(size=14),
          axis.ticks=element_line(colour = "black"))
  print(perc.reads.noncontam)
  dev.off()
  
  cat('\n\n',(ncol(M.seq.tax.subset)-ncol.taxo.noNA),' samples with a total of ',(nrow(M.seq.tax.subset)),' different ASVs out of originally ',nrow(M.seqtab.nochim),' samples with ',(ncol(M.seqtab.nochim)),' different ASVs remain for analysis.',file=(file.path(PathToVisuaRAnalysis,paste(VisuaRProjectName,'_Analysis_log.txt',sep=''))),sep='',append=T)
  gc()
}
gc()
# # save decontaminated seqtab nochim and taxonomy file to your original folders to continue working with them
if(removeContaminants!="" && SaveTables == "Y") {
  M.seq.tax.subset.save <- M.seq.tax.subset
  row_indices <-  match(rownames(M.seq.tax.subset),ASV.mapfile[,"ASV"])
  rownames(M.seq.tax.subset.save) <- ASV.mapfile[row_indices,"ASV sequence"]

  if(removeContaminants == "decontam") {
    # # save decontaminated taxonomy
    saveRDS((M.seq.tax.subset.save[,(ncol(M.seq.tax.subset.save)-ncol.taxo.noNA+1):ncol(M.seq.tax.subset.save)]),file.path(file.path(paste(str_remove(PathToTaxonomy, '.rds'),'_',removeContaminants,"-",decontam_isnotcontaminant,"-",IncludeDNAconc,'_noNAs.rds',sep=''))))
    # # save decontaminated seqtab
    saveRDS(t(M.seq.tax.subset.save[,1:(ncol(M.seq.tax.subset.save)-ncol.taxo.noNA)]),file.path(file.path(paste(str_remove(PathToSeqtabNochim,'.rds'),'_',removeContaminants,"-",decontam_isnotcontaminant,"-",IncludeDNAconc,'.rds',sep=''))))
  } else if(removeContaminants != "") {
    # # save decontaminated taxonomy
    saveRDS((M.seq.tax.subset.save[,(ncol(M.seq.tax.subset.save)-ncol.taxo.noNA+1):ncol(M.seq.tax.subset.save)]),file.path(file.path(paste(str_remove(PathToTaxonomy, '.rds'),'_',removeContaminants,"-",normalizeMseqtax,"-",stat.test,'_noNAs.rds',sep=''))))
    # # save decontaminated seqtab
    saveRDS(t(M.seq.tax.subset.save[,1:(ncol(M.seq.tax.subset.save)-ncol.taxo.noNA)]),file.path(file.path(paste(str_remove(PathToSeqtabNochim,'.rds'),'_',removeContaminants,"-",normalizeMseqtax,"-",stat.test,'.rds',sep=''))))
  }
  rm(M.seq.tax.subset.save,row_indices)  
}

M.sample.reads.left.temp=colSums(M.seq.tax.subset[,1:(ncol(M.seq.tax.subset)-ncol.taxo.noNA)])
M.ASV.per.sample.temp=colSums(M.seq.tax.subset[,1:(ncol(M.seq.tax.subset)-ncol.taxo.noNA)]!=0)


infotable <- fill.info(df.reads=M.sample.reads.left.temp,df.ASVs =M.ASV.per.sample.temp,df.seqtab=M.seq.tax.subset,stepname="AfterDecontamination",infotable=infotable)

rm(M.sample.reads.left.temp,M.ASV.per.sample.temp)

# #=== 2.8. MinimumAllowedReadCount Control ==========================================================================================================================
cat('\n2.8. Excludes samples based on read counts lower than ',MinimumAllowedReadCount,' (MinimumAllowedReadCount).',file=(file.path(PathToVisuaRAnalysis,paste(VisuaRProjectName,'_Analysis_log.txt',sep=''))),sep='',append=TRUE)

if (MinimumAllowedReadCount==''|MinimumAllowedReadCount==0) {
  cat('\nNo samples were excluded based on their read counts as no MinimumAllowedReadCount was chosen.',file=(file.path(PathToVisuaRAnalysis,paste(VisuaRProjectName,'_Analysis_log.txt',sep=''))),sep='',append=T)
} else {
  M.seq.tax.MinimumAllowedReadCount.samples.excluded=as.data.frame(M.seq.tax.subset[,-which(numcolwise(sum)(M.seq.tax.subset)>=MinimumAllowedReadCount)]) 
  if (ncol(M.seq.tax.MinimumAllowedReadCount.samples.excluded)==ncol.taxo.noNA) { # this is TRUE if all samples have read counts higher than MinimumAllowedReadCount
    cat('\nNo samples were excluded based on low read counts as all samples had more than ',MinimumAllowedReadCount,' reads.',file=(file.path(PathToVisuaRAnalysis,paste(VisuaRProjectName,'_Analysis_log.txt',sep=''))),sep='',append=T)
    if (SaveWholeworkspace=='N') {rm(M.seq.tax.MinimumAllowedReadCount.samples.excluded)
      gc()}
  } else {
    M.seq.tax.subset=M.seq.tax.subset[,-which(numcolwise(sum)(M.seq.tax.subset)<MinimumAllowedReadCount)] # only keeps samples with read counts higher than or equal to MinimumAllowedReadCount
    cat('\nThe following ',(ncol(M.seq.tax.MinimumAllowedReadCount.samples.excluded)-ncol.taxo.noNA),' samples were exclulded based on low read counts:\n',file=(file.path(PathToVisuaRAnalysis,paste(VisuaRProjectName,'_Analysis_log.txt',sep=''))),sep='',append=TRUE)
    cat(colnames(M.seq.tax.MinimumAllowedReadCount.samples.excluded[1:((ncol(M.seq.tax.MinimumAllowedReadCount.samples.excluded))-ncol.taxo.noNA)]),file=(file.path(PathToVisuaRAnalysis,paste(VisuaRProjectName,'_Analysis_log.txt',sep=''))),sep='\n',append=TRUE)
    M.seq.tax.MinimumAllowedReadCount.ASVs.excluded=M.seq.tax.subset[rowSums(M.seq.tax.subset[,1:(ncol(M.seq.tax.subset)-ncol.taxo.noNA)])==0,] # makes a df of ASVs which were removed from the dataset.
    if (SaveWholeworkspace=='N') {rm(M.seq.tax.MinimumAllowedReadCount.samples.excluded)
      gc()}
    if (nrow(M.seq.tax.MinimumAllowedReadCount.ASVs.excluded)!=0) {
      M.seq.tax.subset=M.seq.tax.subset[rowSums(M.seq.tax.subset[,1:(ncol(M.seq.tax.subset)-ncol.taxo.noNA)])!=0,] 
      cat('\nThe following ',nrow(M.seq.tax.MinimumAllowedReadCount.ASVs.excluded),' ASVs have been excluded because they are no longer present in any of the samples.\n',file=(file.path(PathToVisuaRAnalysis,paste(VisuaRProjectName,'_Analysis_log.txt',sep=''))),sep='',append=TRUE)
      cat(rownames(M.seq.tax.MinimumAllowedReadCount.ASVs.excluded),file=(file.path(PathToVisuaRAnalysis,paste(VisuaRProjectName,'_Analysis_log.txt',sep=''))),sep=',',append=T)
      if(SaveWholeworkspace=='N') {rm(M.seq.tax.MinimumAllowedReadCount.ASVs.excluded)
        gc()}
    } else {
      cat('No ASVs have been completely excluded.',file=(file.path(PathToVisuaRAnalysis,paste(VisuaRProjectName,'_Analysis_log.txt',sep=''))),sep='',append=T)
      if(SaveWholeworkspace=='N') {rm(M.seq.tax.MinimumAllowedReadCount.ASVs.excluded)
        gc()}
    }
  }
}
cat('\n\n',(ncol(M.seq.tax.subset)-ncol.taxo.noNA),' samples with a total of ',(nrow(M.seq.tax.subset)),' different ASVs out of originally ',nrow(M.seqtab.nochim),' samples with ',(ncol(M.seqtab.nochim)),' different ASVs remain for analysis.',file=(file.path(PathToVisuaRAnalysis,paste(VisuaRProjectName,'_Analysis_log.txt',sep=''))),sep='',append=T)

M.sample.reads.left.temp=colSums(M.seq.tax.subset[,1:(ncol(M.seq.tax.subset)-ncol.taxo.noNA)])
M.ASV.per.sample.temp=colSums(M.seq.tax.subset[,1:(ncol(M.seq.tax.subset)-ncol.taxo.noNA)]!=0)
infotable <- fill.info(df.reads=M.sample.reads.left.temp,df.ASVs =M.ASV.per.sample.temp,df.seqtab=M.seq.tax.subset,stepname=paste0("After Minimum Allowed Read Count Filtration (",MinimumAllowedReadCount,")",sep=""),infotable=infotable)
rm(M.sample.reads.left.temp,M.ASV.per.sample.temp)

gc()

# #=== 2.9. KingdomofInterest Selection ===========================================================================================================================
cat('\n\n2.9. Excludes ASVs not belonging to the chosen kingdom ',KingdomOfInterest,file=(file.path(PathToVisuaRAnalysis,paste(VisuaRProjectName,'_Analysis_log.txt',sep=''))),sep='',append=TRUE)

if (KingdomOfInterest=='') {
  cat('\nError: You did not choose a kingdom of interest.\n',file=(file.path(PathToVisuaRAnalysis,paste(VisuaRProjectName,'_Analysis_log.txt',sep=''))),sep='\t',append=TRUE)
  stop('You did not choose a kingdom of interest.')
} else if (KingdomOfInterest=="All") {
  cat('\nNo samples were excluded because you chose to include all kingdoms in your analysis.\n',file=(file.path(PathToVisuaRAnalysis,paste(VisuaRProjectName,'_Analysis_log.txt',sep=''))),sep='',append=T)
} else {
  M.seq.tax.kingdom.ASV.excluded.org=M.seq.tax[grep(KingdomOfInterest,M.seq.tax[,(ncol.seq.tax-ncol.taxo.noNA+1)],invert=T),]                                      # creates an ASV by Sample&Taxonomy df with ASVs not belonging to the KingdomOfInterest (based on the original dataset (M.seq.tax))
  M.seq.tax.kingdom.ASV.excluded=as.data.frame(M.seq.tax.subset[grep(KingdomOfInterest,M.seq.tax.subset[,(ncol(M.seq.tax.subset)-ncol.taxo.noNA+1)],invert=T),])   # creates an ASV by Sample&Taxonomy df with ASVs not belonging to the KingdomOfInterest (based on the subsetted M.seq.tax.subset)
  M.seq.tax.kingdom.ASV.kept.org=M.seq.tax[grep(KingdomOfInterest,M.seq.tax[,(ncol.seq.tax-ncol.taxo.noNA+1)],invert=F),]                                          # creates an ASV by Sample&Taxonomy df with ASVs and Samples belonging to the KingdomOfInterst (based on the original dataset (M.seq.tax))
  M.seq.tax.subset=M.seq.tax.subset[grep(KingdomOfInterest,M.seq.tax.subset[,(ncol(M.seq.tax.subset)-ncol.taxo.noNA+1)],invert=F),]                                # Further Subsets M.seq.tax.subset, such that only ASVs belonging to the KingdomOfInterest are present (based on previous subsetting)
  cat('\nThe following ',nrow(M.seq.tax.kingdom.ASV.excluded.org),' out of ',nrow(M.seq.tax),' ASVs have been excluded because they do not belong to the chosen kingdom ',KingdomOfInterest,'.\n',file=(file.path(PathToVisuaRAnalysis,paste(VisuaRProjectName,'_Analysis_log.txt',sep=''))),sep='',append=T)
  cat(rownames(M.seq.tax.kingdom.ASV.excluded.org),file=(file.path(PathToVisuaRAnalysis,paste(VisuaRProjectName,'_Analysis_log.txt',sep=''))),sep=', ',append=T)
  cat('\n',nrow(M.seq.tax.subset),' out of ',nrow(M.seq.tax),' ASVs remain belonging to the kingdom ',KingdomOfInterest,'.',file=(file.path(PathToVisuaRAnalysis,paste(VisuaRProjectName,'_Analysis_log.txt',sep=''))),sep='',append=T)
  M.seq.tax.kingdom.samples.excluded.org=M.seq.tax.kingdom.ASV.kept.org[,-which(numcolwise(sum)(M.seq.tax.kingdom.ASV.kept.org)!=0)]                # creates an ASV by Sample&Taxonomy df of samples which no longer have any ASV counts belonging to the chosen KingdomOfInterest (based on the original dataset (M.seq.tax))
  M.seq.tax.kingdom.samples.excluded=M.seq.tax.subset[,-which(numcolwise(sum)(M.seq.tax.subset)!=0)]                                                # creates an ASV by Sample&Taxonomy df of samples which no longer have any ASV counts belonging to the chosen KingdomOfInterest (based on the subsetted M.seq.tax.subset)
  if (SaveWholeworkspace=='N') {rm(M.seq.tax.kingdom.ASV.excluded,M.seq.tax.kingdom.ASV.excluded.org)
    gc()}
  if (ncol(M.seq.tax.kingdom.samples.excluded)!=ncol.taxo.noNA&ncol(M.seq.tax.kingdom.samples.excluded.org)!=ncol.taxo.noNA) {                      # only happens if at least one sample was excluded in both dfs (M.seq.tax and M.seq.tax.subset)
    M.seq.tax.kingdom.ASV.kept.org=M.seq.tax.kingdom.ASV.kept.org[,-which(numcolwise(sum)(M.seq.tax.kingdom.ASV.kept.org)==0)]                      # excludes samples which no longer have any ASV counts in original df
    M.seq.tax.subset=M.seq.tax.subset[,-which(numcolwise(sum)(M.seq.tax.subset)==0)]                                                                # excludes samples which no longer have any ASV counts in subsetted df
    cat('\n\nThe following ',(ncol(M.seq.tax.kingdom.samples.excluded.org)-ncol.taxo.noNA),' samples have been excluded due to no observed ASVs in the chosen kingdom ',KingdomOfInterest,'.\n',file=(file.path(PathToVisuaRAnalysis,paste(VisuaRProjectName,'_Analysis_log.txt',sep=''))),sep='',append=T)
    cat(colnames(M.seq.tax.kingdom.samples.excluded.org[c(1:(ncol(M.seq.tax.kingdom.samples.excluded.org)-ncol.taxo.noNA))]),file=(file.path(PathToVisuaRAnalysis,paste(VisuaRProjectName,'_Analysis_log.txt',sep=''))),sep='\n',append = T)
    if (SaveWholeworkspace=='N') {rm(M.seq.tax.kingdom.ASV.kept.org,M.seq.tax.kingdom.samples.excluded,M.seq.tax.kingdom.samples.excluded.org)
      gc()}
  } else if (ncol(M.seq.tax.kingdom.samples.excluded)!=ncol.taxo.noNA&ncol(M.seq.tax.kingdom.samples.excluded.org)==ncol.taxo.noNA) {               # only happens if at least one sample was excluded in the df M.seq.tax.subset but not in M.seq.tax
    M.seq.tax.subset=M.seq.tax.subset[,-which(numcolwise(sum)(M.seq.tax.subset)==0)]                                                                # excludes samples which no longer have any ASV counts in subsetted df
    cat('\n\nThe following ',(ncol(M.seq.tax.kingdom.samples.excluded)-ncol.taxo.noNA),' samples have been excluded due to no observed ASVs in the chosen kingdom ',KingdomOfInterest,'.\n',file=(file.path(PathToVisuaRAnalysis,paste(VisuaRProjectName,'_Analysis_log.txt',sep=''))),sep='',append=T)
    cat(colnames(M.seq.tax.kingdom.samples.excluded[c(1:(ncol(M.seq.tax.kingdom.samples.excluded)-ncol.taxo.noNA))]),file=(file.path(PathToVisuaRAnalysis,paste(VisuaRProjectName,'_Analysis_log.txt',sep=''))),sep='\n',append = T)
    if (SaveWholeworkspace=='N') {rm(M.seq.tax.kingdom.ASV.kept.org,M.seq.tax.kingdom.samples.excluded,M.seq.tax.kingdom.samples.excluded.org)
      gc()}
  } else if (ncol(M.seq.tax.kingdom.samples.excluded)==ncol.taxo.noNA&ncol(M.seq.tax.kingdom.samples.excluded.org)!=ncol.taxo.noNA) {               # only happens if at least one sample was excluded in the df M.seq.tax but not in M.seq.tax.subset
    M.seq.tax.kingdom.ASV.kept.org=M.seq.tax.kingdom.ASV.kept.org[,-which(numcolwise(sum)(M.seq.tax.kingdom.ASV.kept.org)==0)]                      # excludes samples which no longer have any ASV counts in original df
    cat('\n\nThe following ',(ncol(M.seq.tax.kingdom.samples.excluded.org)-ncol.taxo.noNA),' samples have been excluded due to no observed ASVs in the chosen kingdom ',KingdomOfInterest,'.\n',file=(file.path(PathToVisuaRAnalysis,paste(VisuaRProjectName,'_Analysis_log.txt',sep=''))),sep='',append=T)
    cat(colnames(M.seq.tax.kingdom.samples.excluded.org[c(1:(ncol(M.seq.tax.kingdom.samples.excluded.org)-ncol.taxo.noNA))]),file=(file.path(PathToVisuaRAnalysis,paste(VisuaRProjectName,'_Analysis_log.txt',sep=''))),sep='\n',append = T)
    if (SaveWholeworkspace=='N') {rm(M.seq.tax.kingdom.ASV.kept.org,M.seq.tax.kingdom.samples.excluded,M.seq.tax.kingdom.samples.excluded.org)
      gc()}
  } else if (ncol(M.seq.tax.kingdom.samples.excluded)==ncol.taxo.noNA&ncol(M.seq.tax.kingdom.samples.excluded.org)==ncol.taxo.noNA) {               # only happens if no sample was excluded
    cat('\nNo samples were excluded due to no observed ASVs in the chosen kingdom.\n',file=(file.path(PathToVisuaRAnalysis,paste(VisuaRProjectName,'_Analysis_log.txt',sep=''))),sep='',append=T)
    if (SaveWholeworkspace=='N') {rm(M.seq.tax.kingdom.ASV.kept.org,M.seq.tax.kingdom.samples.excluded,M.seq.tax.kingdom.samples.excluded.org)
      gc()}
  }
}
cat((ncol(M.seq.tax.subset)-ncol.taxo.noNA),' samples with a total of ',(nrow(M.seq.tax.subset)),' different ASVs out of originally ',nrow(M.seqtab.nochim),' samples with ',(ncol(M.seqtab.nochim)),' different ASVs remain for analysis.',file=(file.path(PathToVisuaRAnalysis,paste(VisuaRProjectName,'_Analysis_log.txt',sep=''))),sep='',append=T)

M.sample.reads.left.temp=colSums(M.seq.tax.subset[,1:(ncol(M.seq.tax.subset)-ncol.taxo.noNA)])
M.ASV.per.sample.temp=colSums(M.seq.tax.subset[,1:(ncol(M.seq.tax.subset)-ncol.taxo.noNA)]!=0)
infotable <- fill.info(df.reads=M.sample.reads.left.temp,df.ASVs =M.ASV.per.sample.temp,df.seqtab=M.seq.tax.subset,stepname=paste0("After Filtering for the selected Kingdom ",KingdomOfInterest,sep=""),infotable=infotable)
rm(M.sample.reads.left.temp,M.ASV.per.sample.temp)

gc()
# #=== 2.10. Clade of Interest Selection ========================================================================================================================
cat('\n\n2.10. Keeps particular ASVs based on the clade they belong to',file=(file.path(PathToVisuaRAnalysis,paste(VisuaRProjectName,'_Analysis_log.txt',sep=''))),sep='',append=TRUE)

if (CladeOfInterest=='') {
  cat('\nNo Clade of Interest was chosen.',file=(file.path(PathToVisuaRAnalysis,paste(VisuaRProjectName,'_Analysis_log.txt',sep=''))),sep='',append = T)
} else {
  # # Generate a list of column indices for filtering
  filterCols <- (ncol(M.seq.tax) - ncol.taxo.noNA+1):(ncol(M.seq.tax))
  filterCols.sub <- (ncol(M.seq.tax.subset)-ncol.taxo.noNA+1):(ncol(M.seq.tax.subset))
  
  # # generate a table based on the original data (unfiltered) with only the clades we are not interested in
  M.seq.tax.clade.ASV.excluded.org <- M.seq.tax[!apply(M.seq.tax[, filterCols], 1, function(row) any(grepl(CladeOfInterest, row, ignore.case = TRUE))), ] 
  # # generate a table based on the already pre-filtered data with only the clades we are not interested in
  M.seq.tax.clade.ASV.excluded <- M.seq.tax.subset[!apply(M.seq.tax.subset[, filterCols.sub], 1, function(row) any(grepl(CladeOfInterest, row, ignore.case = TRUE))), ] 
  
  # # generate a table based on the original data (unfiltered) with only the clade of interest
  M.seq.tax.clade.ASV.kept.org <- M.seq.tax[apply(M.seq.tax[, filterCols], 1, function(row) any(grepl(CladeOfInterest, row, ignore.case = TRUE))), ] 
  # # generate a table based on pre-filtered data with only the clade of interest
  M.seq.tax.subset <- M.seq.tax.subset[apply(M.seq.tax.subset[, filterCols.sub], 1, function(row) any(grepl(CladeOfInterest, row, ignore.case = TRUE))), ] 
  
  cat('\n',nrow(M.seq.tax.clade.ASV.excluded.org),' out of ',nrow(M.seq.tax),' ASVs have been excluded because they do not belong to the chosen clade.',file=(file.path(PathToVisuaRAnalysis,paste(VisuaRProjectName,'_Analysis_log.txt',sep=''))),sep='',append=T)
  cat('\n',nrow(M.seq.tax.subset),' out of ',nrow(M.seq.tax),' ASVs remain belonging to the clade ',CladeOfInterest,' or where this expression occurs in the taxonomy.',file=(file.path(PathToVisuaRAnalysis,paste(VisuaRProjectName,'_Analysis_log.txt',sep=''))),sep='',append=T)
  M.seq.tax.clade.samples.excluded.org=M.seq.tax.clade.ASV.kept.org[,-which(numcolwise(sum)(M.seq.tax.clade.ASV.kept.org)!=0)]      # creates an ASV by Sample&Taxonomy df of all samples which no longer have any ASV counts belonging to the chosen CladeOfInterest (based on the original dataset (M.seq.tax))
  M.seq.tax.clade.samples.excluded=M.seq.tax.subset[,-which(numcolwise(sum)(M.seq.tax.subset)!=0)]                                  # creates an ASV by Sample&Taxonomy df of all samples which no longer have any ASV counts belonging to the chosen CladeOfInterest (based on the subsetted M.seq.tax.subset)
  if (SaveWholeworkspace=='N') {rm(M.seq.tax.clade.ASV.excluded.org,M.seq.tax.clade.ASV.excluded)
    gc()}
  if (ncol(M.seq.tax.clade.samples.excluded)!=ncol.taxo.noNA&(ncol(M.seq.tax.clade.samples.excluded.org)!=ncol.taxo.noNA)) {        # only happens if at least one sample was excluded in both dfs
    M.seq.tax.clade.ASV.kept.org=M.seq.tax.clade.ASV.kept.org[,-which(numcolwise(sum)(M.seq.tax.clade.ASV.kept.org)==0)]            # excludes samples which no longer have any ASV counts
    M.seq.tax.subset=M.seq.tax.subset[,-which(numcolwise(sum)(M.seq.tax.subset)==0)]                                                # excludes samples which no longer have any ASV counts
    cat('\nThe following ',(ncol(M.seq.tax.clade.samples.excluded.org)-ncol.taxo.noNA),' samples have been excluded due to no observed ASVs in the chosen CladeOfInterest ',CladeOfInterest,'.\n',file=(file.path(PathToVisuaRAnalysis,paste(VisuaRProjectName,'_Analysis_log.txt',sep=''))),sep='',append=T)
    cat(colnames(M.seq.tax.clade.samples.excluded.org[c(1:(ncol(M.seq.tax.clade.samples.excluded.org)-ncol.taxo.noNA))]),file=(file.path(PathToVisuaRAnalysis,paste(VisuaRProjectName,'_Analysis_log.txt',sep=''))),sep='\n',append=T)
    if (SaveWholeworkspace=='N') {rm(M.seq.tax.clade.ASV.kept.org,M.seq.tax.clade.samples.excluded.org,M.seq.tax.clade.samples.excluded)
      gc()}
  } else if (ncol(M.seq.tax.clade.samples.excluded)!=ncol.taxo.noNA&(ncol(M.seq.tax.clade.samples.excluded.org)==ncol.taxo.noNA)) { # only happens if at least one sample was excluded in the df M.seq.tax.subset but not in M.seq.tax
    M.seq.tax.subset=M.seq.tax.subset[,-which(numcolwise(sum)(M.seq.tax.subset)==0)]                                                # excludes samples which no longer have any ASV counts
    cat('\nThe following ',(ncol(M.seq.tax.clade.samples.excluded)-ncol.taxo.noNA),' samples have been excluded due to no observed ASVs in the chosen CladeOfInterest ',CladeOfInterest,'.\n',file=(file.path(PathToVisuaRAnalysis,paste(VisuaRProjectName,'_Analysis_log.txt',sep=''))),sep='',append=T)
    cat(colnames(M.seq.tax.clade.samples.excluded[c(1:(ncol(M.seq.tax.clade.samples.excluded)-ncol.taxo.noNA))]),file=(file.path(PathToVisuaRAnalysis,paste(VisuaRProjectName,'_Analysis_log.txt',sep=''))),sep='\n',append=T)
    if (SaveWholeworkspace=='N') {rm(M.seq.tax.clade.ASV.kept.org,M.seq.tax.clade.samples.excluded.org,M.seq.tax.clade.samples.excluded)
      gc()}
  } else if (ncol(M.seq.tax.clade.samples.excluded)==ncol.taxo.noNA&(ncol(M.seq.tax.clade.samples.excluded.org)!=ncol.taxo.noNA)) { # only happens if at least one sample was excluded in the df M.seq.tax but not in M.seq.tax.subset
    M.seq.tax.clade.ASV.kept.org=M.seq.tax.clade.ASV.kept.org[,-which(numcolwise(sum)(M.seq.tax.clade.ASV.kept.org)==0)]            # excludes samples which no longer have any ASV counts
    cat('\nThe following ',(ncol(M.seq.tax.clade.samples.excluded.org)-ncol.taxo.noNA),' samples have been excluded due to no observed ASVs in the chosen CladeOfInterest ',CladeOfInterest,'.\n',file=(file.path(PathToVisuaRAnalysis,paste(VisuaRProjectName,'_Analysis_log.txt',sep=''))),sep='',append=T)
    cat(colnames(M.seq.tax.clade.samples.excluded.org[c(1:(ncol(M.seq.tax.clade.samples.excluded.org)-ncol.taxo.noNA))]),file=(file.path(PathToVisuaRAnalysis,paste(VisuaRProjectName,'_Analysis_log.txt',sep=''))),sep='\n',append=T)
    if (SaveWholeworkspace=='N') {rm(M.seq.tax.clade.ASV.kept.org,M.seq.tax.clade.samples.excluded.org,M.seq.tax.clade.samples.excluded)
      gc()}
  } else if (ncol(M.seq.tax.clade.samples.excluded)==ncol.taxo.noNA&(ncol(M.seq.tax.clade.samples.excluded.org)==ncol.taxo.noNA)) { # only happens if at no sample was excluded
    cat('\nNo samples were excluded due to no observed ASVs in the chosen clade.',file=(file.path(PathToVisuaRAnalysis,paste(VisuaRProjectName,'_Analysis_log.txt',sep=''))),sep='',append=T)
    if (SaveWholeworkspace=='N') {rm(M.seq.tax.clade.ASV.kept.org,M.seq.tax.clade.samples.excluded.org,M.seq.tax.clade.samples.excluded)
      gc()}
  }
}
cat('\n',(ncol(M.seq.tax.subset)-ncol.taxo.noNA),' samples with a total of ',(nrow(M.seq.tax.subset)),' different ASVs out of originally ',nrow(M.seqtab.nochim),' samples with ',(ncol(M.seqtab.nochim)),' different ASVs remain for analysis.',file=(file.path(PathToVisuaRAnalysis,paste(VisuaRProjectName,'_Analysis_log.txt',sep=''))),sep='',append=T)

M.sample.reads.left.temp=colSums(M.seq.tax.subset[,1:(ncol(M.seq.tax.subset)-ncol.taxo.noNA)])
M.ASV.per.sample.temp=colSums(M.seq.tax.subset[,1:(ncol(M.seq.tax.subset)-ncol.taxo.noNA)]!=0)
infotable <- fill.info(df.reads=M.sample.reads.left.temp,df.ASVs =M.ASV.per.sample.temp,df.seqtab=M.seq.tax.subset,stepname="After Selecting the clade of interest",infotable=infotable)
rm(M.sample.reads.left.temp,M.ASV.per.sample.temp)

gc()

# #=== 2.11. Clade to exclude selection =====================================================================================================================

cat('\n\n2.11. Excludes particular ASVs based on the clade they belong to.',file=(file.path(PathToVisuaRAnalysis,paste(VisuaRProjectName,'_Analysis_log.txt',sep=''))),sep='',append=TRUE)

if (ExcludeClade=='') {
  cat('\nNo Clade to exclude was chosen.',file=(file.path(PathToVisuaRAnalysis,paste(VisuaRProjectName,'_Analysis_log.txt',sep=''))),sep='',append = T)
} else {
  # # Generate a list of column indices for filtering
  filterCols <- (ncol(M.seq.tax) - ncol.taxo.noNA+1):(ncol(M.seq.tax))
  filterCols.sub <- (ncol(M.seq.tax.subset)-ncol.taxo.noNA+1):(ncol(M.seq.tax.subset))

  # # generate a table based on the original data (unfiltered) with only the clades we want to exclude
  M.seq.tax.clade2.ASV.excluded.org <- M.seq.tax[apply(M.seq.tax[, filterCols], 1, function(row) any(grepl(ExcludeClade, row, ignore.case = TRUE))), ] 
  # # generate a table based on the already pre-filtered data with only the clades we want to exclude
  M.seq.tax.clade2.ASV.excluded <- M.seq.tax.subset[apply(M.seq.tax.subset[, filterCols.sub], 1, function(row) any(grepl(ExcludeClade, row, ignore.case = TRUE))), ] 
  
  # # generate a table based on the original data (unfiltered) with only the remaining clades (without the one we want to exclude)
  M.seq.tax.clade2.ASV.kept.org <- M.seq.tax[!apply(M.seq.tax[, filterCols], 1, function(row) any(grepl(ExcludeClade, row, ignore.case = TRUE))), ] 
  # # generate a table based on pre-filtered data with only the remaining clades (without the one we want to exclude)
  M.seq.tax.subset <- M.seq.tax.subset[!apply(M.seq.tax.subset[, filterCols.sub], 1, function(row) any(grepl(ExcludeClade, row, ignore.case = TRUE))), ] 
  
  cat('\n',nrow(M.seq.tax.clade2.ASV.excluded.org),' out of ',nrow(M.seq.tax),' ASVs have been excluded because they do belong to the clade ',ExcludeClade,'.',file=(file.path(PathToVisuaRAnalysis,paste(VisuaRProjectName,'_Analysis_log.txt',sep=''))),sep='',append=T)
  cat('\n',nrow(M.seq.tax.subset),' out of ',nrow(M.seq.tax),' ASVs remain not belonging to the clade ',ExcludeClade,'.',file=(file.path(PathToVisuaRAnalysis,paste(VisuaRProjectName,'_Analysis_log.txt',sep=''))),sep='',append=T)
  M.seq.tax.clade2.samples.excluded.org=M.seq.tax.clade2.ASV.kept.org[,-which(numcolwise(sum)(M.seq.tax.clade2.ASV.kept.org)!=0)]       # creates an ASV by Sample&Taxonomy df of all samples which no longer have any ASV counts (based on the original dataset (M.seq.tax))
  M.seq.tax.clade2.samples.excluded=M.seq.tax.subset[,-which(numcolwise(sum)(M.seq.tax.subset)!=0)]                                     # creates an ASV by Sample&Taxonomy df of all samples which no longer have any ASV counts (based on the subsetted M.seq.tax.subset)
  if (SaveWholeworkspace=='N') {
    rm(M.seq.tax.clade2.ASV.excluded.org,M.seq.tax.clade2.ASV.excluded)
    gc()
  }
  if (ncol(M.seq.tax.clade2.samples.excluded)!=ncol.taxo.noNA&ncol(M.seq.tax.clade2.samples.excluded.org)!=ncol.taxo.noNA) {            # only happens if at least one sample was excluded in both dfs
    M.seq.tax.clade2.ASV.kept.org=M.seq.tax.clade2.ASV.kept.org[,-which(numcolwise(sum)(M.seq.tax.clade2.ASV.kept.org)==0)]             # excludes samples which no longer have any ASV counts
    M.seq.tax.subset=M.seq.tax.subset[,-which(numcolwise(sum)(M.seq.tax.subset)==0)]                                                    # excludes samples which no longer have any ASV counts
    cat('\nThe following ',(ncol(M.seq.tax.clade2.samples.excluded.org)-ncol.taxo.noNA),' samples have been excluded due to no observed ASVs in the remaining clades.\n',file=(file.path(PathToVisuaRAnalysis,paste(VisuaRProjectName,'_Analysis_log.txt',sep=''))),sep='',append=T)
    cat(colnames(M.seq.tax.clade2.samples.excluded.org[c(1:(ncol(M.seq.tax.clade2.samples.excluded.org)-ncol.taxo.noNA))]),file=(file.path(PathToVisuaRAnalysis,paste(VisuaRProjectName,'_Analysis_log.txt',sep=''))),sep='\n',append=T)
    if (SaveWholeworkspace=='N') {
      rm(M.seq.tax.clade2.ASV.kept.org,M.seq.tax.clade2.samples.excluded.org,M.seq.tax.clade2.samples.excluded)
      gc()}
  } else if (ncol(M.seq.tax.clade2.samples.excluded)!=ncol.taxo.noNA&ncol(M.seq.tax.clade2.samples.excluded.org)==ncol.taxo.noNA) {     # only happens if at least one sample was excluded in the df M.seq.tax.subset but not in M.seq.tax
    M.seq.tax.subset=M.seq.tax.subset[,-which(numcolwise(sum)(M.seq.tax.subset)==0)]                                                    # excludes samples which no longer have any ASV counts
    cat('\nThe following ',(ncol(M.seq.tax.clade2.samples.excluded)-ncol.taxo.noNA),' samples have been excluded due to no observed ASVs in the remaining clades.\n',file=(file.path(PathToVisuaRAnalysis,paste(VisuaRProjectName,'_Analysis_log.txt',sep=''))),sep='',append=T)
    cat(colnames(M.seq.tax.clade2.samples.excluded[c(1:(ncol(M.seq.tax.clade2.samples.excluded)-ncol.taxo.noNA))]),file=(file.path(PathToVisuaRAnalysis,paste(VisuaRProjectName,'_Analysis_log.txt',sep=''))),sep='\n',append=T)
    if (SaveWholeworkspace=='N') {
      rm(M.seq.tax.clade2.ASV.kept.org,M.seq.tax.clade2.samples.excluded.org,M.seq.tax.clade2.samples.excluded)
      gc()}
  } else if (ncol(M.seq.tax.clade2.samples.excluded)==ncol.taxo.noNA&ncol(M.seq.tax.clade2.samples.excluded.org)!=ncol.taxo.noNA) {     # only happens if at least one sample was excluded in the df M.seq.tax but not in M.seq.tax.subset
    M.seq.tax.clade2.ASV.kept.org=M.seq.tax.clade2.ASV.kept.org[,-which(numcolwise(sum)(M.seq.tax.clade2.ASV.kept.org)==0)]             # excludes samples which no longer have any ASV counts
    cat('\nThe following ',(ncol(M.seq.tax.clade2.samples.excluded.org)-ncol.taxo.noNA),' samples have been excluded due to no observed ASVs in the remaining clades.\n',file=(file.path(PathToVisuaRAnalysis,paste(VisuaRProjectName,'_Analysis_log.txt',sep=''))),sep='',append=T)
    cat(colnames(M.seq.tax.clade2.samples.excluded.org[c(1:(ncol(M.seq.tax.clade2.samples.excluded.org)-ncol.taxo.noNA))]),file=(file.path(PathToVisuaRAnalysis,paste(VisuaRProjectName,'_Analysis_log.txt',sep=''))),sep='\n',append=T)
    if (SaveWholeworkspace=='N') {
      rm(M.seq.tax.clade2.ASV.kept.org,M.seq.tax.clade2.samples.excluded.org,M.seq.tax.clade2.samples.excluded)
      gc()}
  } else if (ncol(M.seq.tax.clade2.samples.excluded)==ncol.taxo.noNA&ncol(M.seq.tax.clade2.samples.excluded.org)==ncol.taxo.noNA) {     # only happens if no sample was excluded
    cat('\nNo samples were excluded due to no observed ASVs in the remaining clades.',file=(file.path(PathToVisuaRAnalysis,paste(VisuaRProjectName,'_Analysis_log.txt',sep=''))),sep='',append=T)
    if (SaveWholeworkspace=='N') {
      rm(M.seq.tax.clade2.ASV.kept.org,M.seq.tax.clade2.samples.excluded.org,M.seq.tax.clade2.samples.excluded)
      gc()}
  }
}

cat('\n',(ncol(M.seq.tax.subset)-ncol.taxo.noNA),' samples with a total of ',(nrow(M.seq.tax.subset)),' different ASVs out of originally ',nrow(M.seqtab.nochim),' samples with ',(ncol(M.seqtab.nochim)),' different ASVs remain for analysis.',file=(file.path(PathToVisuaRAnalysis,paste(VisuaRProjectName,'_Analysis_log.txt',sep=''))),sep='',append=T)

M.sample.reads.left.temp=colSums(M.seq.tax.subset[,1:(ncol(M.seq.tax.subset)-ncol.taxo.noNA)])
M.ASV.per.sample.temp=colSums(M.seq.tax.subset[,1:(ncol(M.seq.tax.subset)-ncol.taxo.noNA)]!=0)
infotable <- fill.info(df.reads=M.sample.reads.left.temp,df.ASVs =M.ASV.per.sample.temp,df.seqtab=M.seq.tax.subset,stepname="After Excluding selected clades",infotable=infotable)
rm(M.sample.reads.left.temp,M.ASV.per.sample.temp)

gc()

# #=== 2.12. Sample selection (based on the sample name) ==============================================================================================================================

cat('\n\n2.12. Keeps particular samples based on the sample name.',file=(file.path(PathToVisuaRAnalysis,paste(VisuaRProjectName,'_Analysis_log.txt',sep=''))),sep='',append=TRUE)

if (KeepSamplesbyName=='') {
  cat('\nNo particular samples to keep were chosen.',file=(file.path(PathToVisuaRAnalysis,paste(VisuaRProjectName,'_Analysis_log.txt',sep=''))),sep='',append = T)
} else {
  M.seq.tax.keepsample.excluded.org=M.seq.tax[,grep(KeepSamplesbyName,colnames(M.seq.tax),invert=T)]                                    # creates an ASV by Sample&Taxonomy df with only those samples not belonging to the 'KeepSamplesbyName' (includes taxonomy, based on the original dataset (M.seq.tax))
  M.seq.tax.keepsample.excluded=as.data.frame(M.seq.tax.subset[,grep(KeepSamplesbyName,colnames(M.seq.tax.subset),invert=T)])           # creates an ASV by Sample&Taxonomy df with only those samples not belonging to the 'KeepSamplesbyName' (includes taxonomy, based on the subsetted dataset (M.seq.tax.subset))
  M.seq.tax.keepsample.kept.org=cbind(M.seq.tax[,grep(KeepSamplesbyName,colnames(M.seq.tax),invert=F)],M.seq.tax[,(ncol(M.seq.tax)-ncol.taxo.noNA+1):ncol(M.seq.tax)]) # creates an ASV by Sample&Taxonomy df with only those samples belonging to the 'KeepSamplesByName' (based on the original dataset (M.seq.tax)). The taxonomy is deleted by the grep function and therefore added with cbind
  M.seq.tax.subset=cbind(M.seq.tax.subset[,grep(KeepSamplesbyName,colnames(M.seq.tax.subset),invert=F)],M.seq.tax.subset[,(ncol(M.seq.tax.subset)-ncol.taxo.noNA+1):ncol(M.seq.tax.subset)]) # creates an ASV by Sample&Taxonomy df with only those samples belonging to the 'KeepSamplesByName' (based on the subsetted dataset (M.seq.tax.subset)). The taxonomy is deleted by the grep function and therefore added with cbind
  cat('\n','Based on the expressions chosen to keep:\n',file=(file.path(PathToVisuaRAnalysis,paste(VisuaRProjectName,'_Analysis_log.txt',sep=''))),sep='',append=T)
  cat(KeepSamplesbyName,file=(file.path(PathToVisuaRAnalysis,paste(VisuaRProjectName,'_Analysis_log.txt',sep=''))),sep='\n',append=T)
  cat('The following ',ncol(M.seq.tax.keepsample.excluded.org)-ncol.taxo.noNA,' samples were excluded.\n',file=(file.path(PathToVisuaRAnalysis,paste(VisuaRProjectName,'_Analysis_log.txt',sep=''))),sep='',append=T)
  cat(colnames(M.seq.tax.keepsample.excluded.org[,1:(ncol(M.seq.tax.keepsample.excluded.org)-ncol.taxo.noNA)]),file=(file.path(PathToVisuaRAnalysis,paste(VisuaRProjectName,'_Analysis_log.txt',sep=''))),sep='\n',append=T)
  M.seq.tax.keepsample.ASVs.excluded.org=M.seq.tax.keepsample.kept.org[rowSums(M.seq.tax.keepsample.kept.org[,1:(ncol(M.seq.tax.keepsample.kept.org)-ncol.taxo.noNA)])==0,] # creates an ASV by Sample&Taxonomy df with all ASVs which do not occur in the subsetted samples (based on the original dataset (M.seq.tax))
  M.seq.tax.keepsample.ASVs.excluded=M.seq.tax.subset[rowSums(M.seq.tax.subset[,1:(ncol(M.seq.tax.subset)-ncol.taxo.noNA)])==0,]        # creates an ASV by Sample&Taxonomy df with all ASVs which do not occur in the subsetted samples (based on the subsetted dataset (M.seq.tax.subset))
  if (SaveWholeworkspace=='N') {rm(M.seq.tax.keepsample.excluded.org,M.seq.tax.keepsample.excluded)
    gc()}
  if (nrow(M.seq.tax.keepsample.ASVs.excluded)!=0&nrow(M.seq.tax.keepsample.ASVs.excluded.org)!=0) {                                    # only happens if at least one ASV was excluded in both dfs
    M.seq.tax.keepsample.kept.org=M.seq.tax.keepsample.kept.org[rowSums(M.seq.tax.keepsample.kept.org[,1:(ncol(M.seq.tax.keepsample.kept.org)-ncol.taxo.noNA)])!=0,] # excludes ASVs not longer present in the samples
    M.seq.tax.subset=M.seq.tax.subset[rowSums(M.seq.tax.subset[,1:(ncol(M.seq.tax.subset)-ncol.taxo.noNA)])!=0,]                        # excludes ASVs not longer present in the samples
    cat('The following ',nrow(M.seq.tax.keepsample.ASVs.excluded.org),' ASVs have been excluded because they are no longer present in any of the samples.\n',file=(file.path(PathToVisuaRAnalysis,paste(VisuaRProjectName,'_Analysis_log.txt',sep=''))),sep='',append=T)
    cat(rownames(M.seq.tax.keepsample.ASVs.excluded.org),file=(file.path(PathToVisuaRAnalysis,paste(VisuaRProjectName,'_Analysis_log.txt',sep=''))),sep=', ',append=T)
    if (SaveWholeworkspace=='N') {rm(M.seq.tax.keepsample.kept.org,M.seq.tax.keepsample.ASVs.excluded.org,M.seq.tax.keepsample.ASVs.excluded)
      gc()}
  } else if (nrow(M.seq.tax.keepsample.ASVs.excluded)!=0&nrow(M.seq.tax.keepsample.ASVs.excluded.org)==0) {                             # only happens if at least one ASv was excluded in the df M.seq.tax.subset but not in M.seq.tax
    M.seq.tax.subset=M.seq.tax.subset[rowSums(M.seq.tax.subset[,1:(ncol(M.seq.tax.subset)-ncol.taxo.noNA)])!=0,]                        # excludes ASVs not longer present in the samples
    cat('The following ',nrow(M.seq.tax.keepsample.ASVs.excluded),' ASVs have been excluded because they are no longer present in any of the samples.\n',file=(file.path(PathToVisuaRAnalysis,paste(VisuaRProjectName,'_Analysis_log.txt',sep=''))),sep='',append=T)
    cat(rownames(M.seq.tax.keepsample.ASVs.excluded),file=(file.path(PathToVisuaRAnalysis,paste(VisuaRProjectName,'_Analysis_log.txt',sep=''))),sep=', ',append=T)
    if (SaveWholeworkspace=='N') {rm(M.seq.tax.keepsample.kept.org,M.seq.tax.keepsample.ASVs.excluded.org,M.seq.tax.keepsample.ASVs.excluded)
      gc()}
  } else if (nrow(M.seq.tax.keepsample.ASVs.excluded)==0&nrow(M.seq.tax.keepsample.ASVs.excluded.org)!=0) {                             # only happens if at least one sample was excluded in the df M.seq.tax but not in M.seq.tax.subset
    M.seq.tax.keepsample.kept.org=M.seq.tax.keepsample.kept.org[rowSums(M.seq.tax.keepsample.kept.org[,1:(ncol(M.seq.tax.keepsample.kept.org)-ncol.taxo.noNA)])!=0,] # excludes ASVs not longer present in the samples
    cat('The following ',nrow(M.seq.tax.keepsample.ASVs.excluded.org),' ASVs have been excluded because they are no longer present in any of the samples.\n',file=(file.path(PathToVisuaRAnalysis,paste(VisuaRProjectName,'_Analysis_log.txt',sep=''))),sep='',append=T)
    cat(rownames(M.seq.tax.keepsample.ASVs.excluded.org),file=(file.path(PathToVisuaRAnalysis,paste(VisuaRProjectName,'_Analysis_log.txt',sep=''))),sep=', ',append=T)
    if (SaveWholeworkspace=='N') {rm(M.seq.tax.keepsample.kept.org,M.seq.tax.keepsample.ASVs.excluded.org,M.seq.tax.keepsample.ASVs.excluded)
      gc()}
  } else if (nrow(M.seq.tax.keepsample.ASVs.excluded)==0&nrow(M.seq.tax.keepsample.ASVs.excluded.org)==0) {                             # only happens if no ASV was excluded
    cat('\nNo ASVs were excluded due to no observed ASVs in the remaining samples.',file=(file.path(PathToVisuaRAnalysis,paste(VisuaRProjectName,'_Analysis_log.txt',sep=''))),sep='',append=T)
    if (SaveWholeworkspace=='N') {rm(M.seq.tax.keepsample.kept.org,M.seq.tax.keepsample.ASVs.excluded.org,M.seq.tax.keepsample.ASVs.excluded)
      gc()}
  }
}
cat('\n',(ncol(M.seq.tax.subset)-ncol.taxo.noNA),' samples with a total of ',(nrow(M.seq.tax.subset)),' different ASVs out of originally ',nrow(M.seqtab.nochim),' samples with ',(ncol(M.seqtab.nochim)),' different ASVs remain for analysis.',file=(file.path(PathToVisuaRAnalysis,paste(VisuaRProjectName,'_Analysis_log.txt',sep=''))),sep='',append=T)

M.sample.reads.left.temp=colSums(M.seq.tax.subset[,1:(ncol(M.seq.tax.subset)-ncol.taxo.noNA)])
M.ASV.per.sample.temp=colSums(M.seq.tax.subset[,1:(ncol(M.seq.tax.subset)-ncol.taxo.noNA)]!=0)
infotable <- fill.info(df.reads=M.sample.reads.left.temp,df.ASVs =M.ASV.per.sample.temp,df.seqtab=M.seq.tax.subset,stepname="After Selecting certain samples",infotable=infotable)
rm(M.sample.reads.left.temp,M.ASV.per.sample.temp)

gc()

# #=== 2.13. Excludes particular samples based on the sample name ===========================================================================================================================
cat('\n\n2.13. Excludes particular samples based on the sample name.',file=(file.path(PathToVisuaRAnalysis,paste(VisuaRProjectName,'_Analysis_log.txt',sep=''))),sep='',append=T)

if (excludeSamplesbyName=='') {
  cat('\nNo particular samples to exclude were chosen.',file=(file.path(PathToVisuaRAnalysis,paste(VisuaRProjectName,'_Analysis_log.txt',sep=''))),sep='',append = T)
} else {
  M.seq.tax.excludesample.excluded.org=M.seq.tax[,grep(excludeSamplesbyName,colnames(M.seq.tax),invert=F)]                                  # creates an ASV by Sample&Taxonomy df with only those samples belonging to the 'ExcludeSamplesbyName' (based on the original dataset (M.seq.tax))
  M.seq.tax.excludesample.excluded=as.data.frame(M.seq.tax.subset[,grep(excludeSamplesbyName,colnames(M.seq.tax.subset),invert=F)])         # creates an ASV by Sample&Taxonomy df with only those samples belonging to the 'ExcludeSamplebyName' (based on the subsetted dataset (M.seq.tax.subset))
  M.seq.tax.excludesample.kept.org=M.seq.tax[,grep(excludeSamplesbyName,colnames(M.seq.tax),invert=T)]                                      # creates an ASV by Sample&Taxonomy df with only those samples not belonging to the 'ExcludeSamplebyName' (based on the original dataset (M.seq.tax)). 
  M.seq.tax.subset=M.seq.tax.subset[,grep(excludeSamplesbyName,colnames(M.seq.tax.subset),invert=T)]                                        # creates an ASV by Sample&Taxonomy df with only those samples not belonging to the 'ExcludeSamplebyName' (based on the subsetted dataset (M.seq.tax.subset)).
  cat('\n','Based on the expressions chosen to exclude:\n',file=(file.path(PathToVisuaRAnalysis,paste(VisuaRProjectName,'_Analysis_log.txt',sep=''))),sep='',append=T)
  cat(excludeSamplesbyName,file=(file.path(PathToVisuaRAnalysis,paste(VisuaRProjectName,'_Analysis_log.txt',sep=''))),sep='\n',append=T)
  cat('The following ',ncol(M.seq.tax.excludesample.excluded.org),' samples were excluded.\n',file=(file.path(PathToVisuaRAnalysis,paste(VisuaRProjectName,'_Analysis_log.txt',sep=''))),sep='',append=T)
  cat(colnames(M.seq.tax.excludesample.excluded.org),file=(file.path(PathToVisuaRAnalysis,paste(VisuaRProjectName,'_Analysis_log.txt',sep=''))),sep='\n',append=T)
  M.seq.tax.excludesample.ASVs.excluded.org=M.seq.tax.excludesample.kept.org[rowSums(M.seq.tax.excludesample.kept.org[,1:(ncol(M.seq.tax.excludesample.kept.org)-ncol.taxo.noNA)])==0,] # creates an ASV by Sample&Taxonomy df with all ASVs which do not occur in the subsetted samples (based on the original dataset (M.seq.tax))
  M.seq.tax.excludesample.ASVs.excluded=M.seq.tax.subset[rowSums(M.seq.tax.subset[,1:(ncol(M.seq.tax.subset)-ncol.taxo.noNA)])==0,]         # creates an ASV by Sample&Taxonomy df with all ASVs which do not occur in the subsetted samples (based on the subsetted dataset (M.seq.tax.subset))
  if (SaveWholeworkspace=='N') {rm(M.seq.tax.excludesample.excluded.org,M.seq.tax.excludesample.excluded)
    gc()}
  if (nrow(M.seq.tax.excludesample.ASVs.excluded)!=0&nrow(M.seq.tax.excludesample.ASVs.excluded.org)!=0) {                                  # only happens if at least one ASV was excluded in both dfs
    M.seq.tax.excludesample.kept.org=M.seq.tax.excludesample.kept.org[rowSums(M.seq.tax.excludesample.kept.org[,1:(ncol(M.seq.tax.excludesample.kept.org)-ncol.taxo.noNA)])!=0,] # excludes ASVs not longer present in the samples
    M.seq.tax.subset=M.seq.tax.subset[rowSums(M.seq.tax.subset[,1:(ncol(M.seq.tax.subset)-ncol.taxo.noNA)])!=0,]                            # excludes ASVs not longer present in the samples
    cat('\nThe following ',nrow(M.seq.tax.excludesample.ASVs.excluded.org),' ASVs have been excluded because they are no longer present in any of the samples.\n',file=(file.path(PathToVisuaRAnalysis,paste(VisuaRProjectName,'_Analysis_log.txt',sep=''))),sep='',append=T)
    cat(rownames(M.seq.tax.excludesample.ASVs.excluded.org),file=(file.path(PathToVisuaRAnalysis,paste(VisuaRProjectName,'_Analysis_log.txt',sep=''))),sep=', ',append=T)
    if (SaveWholeworkspace=='N') {rm(M.seq.tax.excludesample.kept.org,M.seq.tax.excludesample.ASVs.excluded.org,M.seq.tax.excludesample.ASVs.excluded)
      gc()}
  } else if (nrow(M.seq.tax.excludesample.ASVs.excluded)!=0&nrow(M.seq.tax.excludesample.ASVs.excluded.org)==0) {                           # only happens if at least one ASv was excluded in the df M.seq.tax.subset but not in M.seq.tax
    M.seq.tax.subset=M.seq.tax.subset[rowSums(M.seq.tax.subset[,1:(ncol(M.seq.tax.subset)-ncol.taxo.noNA)])!=0,]                            # excludes ASVs not longer present in the samples
    cat('\nThe following ',nrow(M.seq.tax.excludesample.ASVs.excluded.org),' ASVs have been excluded because they are no longer present in any of the samples.\n',file=(file.path(PathToVisuaRAnalysis,paste(VisuaRProjectName,'_Analysis_log.txt',sep=''))),sep='',append=T)
    cat(rownames(M.seq.tax.excludesample.ASVs.excluded.org),file=(file.path(PathToVisuaRAnalysis,paste(VisuaRProjectName,'_Analysis_log.txt',sep=''))),sep=', ',append=T)
    if (SaveWholeworkspace=='N') {rm(M.seq.tax.excludesample.kept.org,M.seq.tax.excludesample.ASVs.excluded.org,M.seq.tax.excludesample.ASVs.excluded)
      gc()}
  } else if (nrow(M.seq.tax.excludesample.ASVs.excluded)==0&nrow(M.seq.tax.excludesample.ASVs.excluded.org)!=0) {                           # only happens if at least one sample was excluded in the df M.seq.tax but not in M.seq.tax.subset
    M.seq.tax.excludesample.kept.org=M.seq.tax.excludesample.kept.org[rowSums(M.seq.tax.excludesample.kept.org[,1:(ncol(M.seq.tax.excludesample.kept.org)-ncol.taxo.noNA)])!=0,] # excludes ASVs not longer present in the samples
    cat('\nThe following ',nrow(M.seq.tax.excludesample.ASVs.excluded.org),' ASVs have been excluded because they are no longer present in any of the samples.\n',file=(file.path(PathToVisuaRAnalysis,paste(VisuaRProjectName,'_Analysis_log.txt',sep=''))),sep='',append=T)
    cat(rownames(M.seq.tax.excludesample.ASVs.excluded.org),file=(file.path(PathToVisuaRAnalysis,paste(VisuaRProjectName,'_Analysis_log.txt',sep=''))),sep=', ',append=T)
    if (SaveWholeworkspace=='N') {rm(M.seq.tax.excludesample.kept.org,M.seq.tax.excludesample.ASVs.excluded.org,M.seq.tax.excludesample.ASVs.excluded)
      gc()}
  } else if (nrow(M.seq.tax.excludesample.ASVs.excluded)==0&nrow(M.seq.tax.excludesample.ASVs.excluded.org)==0) {                           # only happens if no ASV was excluded
    cat('\nNo ASVs were excluded due to no observed ASVs in the remaining samples.',file=(file.path(PathToVisuaRAnalysis,paste(VisuaRProjectName,'_Analysis_log.txt',sep=''))),sep='',append=T)
    if (SaveWholeworkspace=='N') {rm(M.seq.tax.excludesample.kept.org,M.seq.tax.excludesample.ASVs.excluded.org,M.seq.tax.excludesample.ASVs.excluded)
      gc()}
  }
}  
cat('\n',(ncol(M.seq.tax.subset)-ncol.taxo.noNA),' samples with a total of ',(nrow(M.seq.tax.subset)),' different ASVs out of originally ',nrow(M.seqtab.nochim),' samples with ',(ncol(M.seqtab.nochim)),' different ASVs remain for analysis.',file=(file.path(PathToVisuaRAnalysis,paste(VisuaRProjectName,'_Analysis_log.txt',sep=''))),sep='',append=T)

M.sample.reads.left.temp=colSums(M.seq.tax.subset[,1:(ncol(M.seq.tax.subset)-ncol.taxo.noNA)])
M.ASV.per.sample.temp=colSums(M.seq.tax.subset[,1:(ncol(M.seq.tax.subset)-ncol.taxo.noNA)]!=0)
infotable <- fill.info(df.reads=M.sample.reads.left.temp,df.ASVs =M.ASV.per.sample.temp,df.seqtab=M.seq.tax.subset,stepname="After excluding certain samples",infotable=infotable)
rm(M.sample.reads.left.temp,M.ASV.per.sample.temp)

gc()
# #=== 2.14. Excludes subsetted samples based on the remaining reads ========================================================================================================================
# ?? print ausgabe von wenn ein sample rausf?llt funktioniert nicht.???

cat('\n\n2.14. Excludes remaining samples based on read counts lower than ',MinimumAllowedReadCount.Analysis,' (MinimumAllowedReadCount.Analysis).',file=(file.path(PathToVisuaRAnalysis,paste(VisuaRProjectName,'_Analysis_log.txt',sep=''))),sep='',append=TRUE)

if (MinimumAllowedReadCount.Analysis==''|MinimumAllowedReadCount.Analysis==0) {
  cat('\nNo samples were excluded based on their read counts as no MinimumAllowedReadCount.Analysis was chosen or it was set to 0.',file=(file.path(PathToVisuaRAnalysis,paste(VisuaRProjectName,'_Analysis_log.txt',sep=''))),sep='',append=T)
} else {
  M.seq.tax.MinimumAllowedReadCount.Analysis.samples.excluded.org=M.seq.tax[,-which(numcolwise(sum)(M.seq.tax)>=MinimumAllowedReadCount.Analysis)]                                  # goes through all numeric columns and excludes those samples (columns) which have more or equal reads (sum) than MinimumAllowedReadCount.Analysis (based on not subsetted df(M.seq.tax))
  M.seq.tax.MinimumAllowedReadCount.Analysis.samples.excluded=as.data.frame(M.seq.tax.subset[,-which(numcolwise(sum)(M.seq.tax.subset)>=MinimumAllowedReadCount.Analysis)])         # goes through all numeric columns and excludes those samples (columns) which have more or equal reads (sum) than MinimumAllowedReadCount.Analysis (based on  subsetted df(M.seq.tax.subset))
  if (ncol(M.seq.tax.MinimumAllowedReadCount.Analysis.samples.excluded)!=ncol.taxo.noNA&ncol(M.seq.tax.MinimumAllowedReadCount.Analysis.samples.excluded.org)!=ncol.taxo.noNA) {    # only if at least one sample was excluded in both dfs
    M.seq.tax.MinimumAllowedReadCount.Analysis.samples.kept.org=M.seq.tax[,-which(numcolwise(sum)(M.seq.tax.subset)<MinimumAllowedReadCount.Analysis)]                              # only keeps samples with more or equal to MinimumAllowedReadCount reads (based on original df (M.seq.tax))
    M.seq.tax.subset=M.seq.tax.subset[,-which(numcolwise(sum)(M.seq.tax.subset)<MinimumAllowedReadCount.Analysis)]                                                                  # only keeps samples with more or equal to MinimumAllowedReadCount reads (based on subsetted df (M.seq.tax.subset))
    cat('\nThe following ',(ncol(M.seq.tax.MinimumAllowedReadCount.Analysis.samples.excluded)-ncol.taxo.noNA),' samples were exclulded based on low read counts:\n',file=(file.path(PathToVisuaRAnalysis,paste(VisuaRProjectName,'_Analysis_log.txt',sep=''))),sep='',append=T)
    cat(colnames(M.seq.tax.MinimumAllowedReadCount.Analysis.samples.excluded[1:((ncol(M.seq.tax.MinimumAllowedReadCount.Analysis.samples.excluded))-ncol.taxo.noNA)]),file=(file.path(PathToVisuaRAnalysis,paste(VisuaRProjectName,'_Analysis_log.txt',sep=''))),sep='\n',append=T)
    M.seq.tax.MinimumAllowedReadCount.Analysis.ASVs.excluded.org=M.seq.tax.MinimumAllowedReadCount.Analysis.samples.kept.org[rowSums(M.seq.tax.MinimumAllowedReadCount.Analysis.samples.kept.org[,1:(ncol(M.seq.tax.MinimumAllowedReadCount.Analysis.samples.kept.org)-ncol.taxo.noNA)])==0,] # creates an ASV by Sample&Taxonomy df with all ASVs which do not occur in the  samples (based on the original dataset (M.seq.tax)
    M.seq.tax.MinimumAllowedReadCount.Analysis.ASVs.excluded=M.seq.tax.subset[rowSums(M.seq.tax.subset[,1:(ncol(M.seq.tax.subset)-ncol.taxo.noNA)])==0,]                            # creates an ASV by Sample&Taxonomy df with all ASVs which do not occur in the subsetted samples (based on the subsetted dataset (M.seq.tax.subset))
    if (SaveWholeworkspace=='N') {rm(M.seq.tax.MinimumAllowedReadCount.Analysis.samples.excluded.org,M.seq.tax.MinimumAllowedReadCount.Analysis.samples.excluded)
      gc()}
    if (nrow(M.seq.tax.MinimumAllowedReadCount.Analysis.ASVs.excluded)!=0&nrow(M.seq.tax.MinimumAllowedReadCount.Analysis.ASVs.excluded.org)!=0) {                                  # only happens if at least one ASV was excluded in both dfs
      M.seq.tax.MinimumAllowedReadCount.Analysis.samples.kept.org=M.seq.tax.MinimumAllowedReadCount.Analysis.samples.kept.org[rowSums(M.seq.tax.MinimumAllowedReadCount.Analysis.samples.kept.org[,1:(ncol(M.seq.tax.MinimumAllowedReadCount.Analysis.samples.kept.org)-ncol.taxo.noNA)])!=0,] #excludes ASVs which do not occur anymore in the original df
      M.seq.tax.subset=M.seq.tax.subset[rowSums(M.seq.tax.subset[,1:(ncol(M.seq.tax.subset)-ncol.taxo.noNA)])!=0,]                                                                  # excludes ASVs which do not occur anymore in the subsetted df
      cat('The following ',nrow(M.seq.tax.MinimumAllowedReadCount.Analysis.ASVs.excluded),' ASVs have been exclulded because they are no longer present in any of the samples:\n',file=(file.path(PathToVisuaRAnalysis,paste(VisuaRProjectName,'_Analysis_log.txt',sep=''))),sep='',append=T)
      cat(rownames(M.seq.tax.MinimumAllowedReadCount.Analysis.ASVs.excluded),file=(file.path(PathToVisuaRAnalysis,paste(VisuaRProjectName,'_Analysis_log.txt',sep=''))),sep=', ',append=T)
      if (SaveWholeworkspace=='N') {rm(M.seq.tax.MinimumAllowedReadCount.Analysis.samples.kept.org,M.seq.tax.MinimumAllowedReadCount.Analysis.ASVs.excluded.org,M.seq.tax.MinimumAllowedReadCount.Analysis.ASVs.excluded)
        gc()}
    } else if (nrow(M.seq.tax.MinimumAllowedReadCount.Analysis.ASVs.excluded)!=0&nrow(M.seq.tax.MinimumAllowedReadCount.Analysis.ASVs.excluded.org)==0) {                           # only happens if at least one ASV was excluded in M.seq.tax.subset but not in M.seq.tax
      M.seq.tax.subset=M.seq.tax.subset[rowSums(M.seq.tax.subset[,1:(ncol(M.seq.tax.subset)-ncol.taxo.noNA)])!=0,]                                                                  # excludes ASVs which do not occur anymore in the subsetted df
      cat('The following ',nrow(M.seq.tax.MinimumAllowedReadCount.Analysis.ASVs.excluded),' ASVs have been exclulded because they are no longer present in any of the samples:\n',file=(file.path(PathToVisuaRAnalysis,paste(VisuaRProjectName,'_Analysis_log.txt',sep=''))),sep='',append=T)
      cat(rownames(M.seq.tax.MinimumAllowedReadCount.Analysis.ASVs.excluded),file=(file.path(PathToVisuaRAnalysis,paste(VisuaRProjectName,'_Analysis_log.txt',sep=''))),sep=', ',append=T)
      if (SaveWholeworkspace=='N') {rm(M.seq.tax.MinimumAllowedReadCount.Analysis.samples.kept.org,M.seq.tax.MinimumAllowedReadCount.Analysis.ASVs.excluded.org,M.seq.tax.MinimumAllowedReadCount.Analysis.ASVs.excluded)
        gc()}
    } else if (nrow(M.seq.tax.MinimumAllowedReadCount.Analysis.ASVs.excluded)==0&nrow(M.seq.tax.MinimumAllowedReadCount.Analysis.ASVs.excluded.org)!=0) {                           # only happens if at least one ASV was excluded in M.seq.tax but not in M.seq.tax.subset
      M.seq.tax.MinimumAllowedReadCount.Analysis.samples.kept.org=M.seq.tax.MinimumAllowedReadCount.Analysis.samples.kept.org[rowSums(M.seq.tax.MinimumAllowedReadCount.Analysis.samples.kept.org[,1:(ncol(M.seq.tax.MinimumAllowedReadCount.Analysis.samples.kept.org)-ncol.taxo.noNA)])!=0,] #excludes ASVs which do not occur anymore in the original df
      cat('The following ',nrow(M.seq.tax.MinimumAllowedReadCount.Analysis.ASVs.excluded.org),' ASVs have been exclulded because they are no longer present in any of the samples:\n',file=(file.path(PathToVisuaRAnalysis,paste(VisuaRProjectName,'_Analysis_log.txt',sep=''))),sep='',append=T)
      cat(rownames(M.seq.tax.MinimumAllowedReadCount.Analysis.ASVs.excluded.org),file=(file.path(PathToVisuaRAnalysis,paste(VisuaRProjectName,'_Analysis_log.txt',sep=''))),sep=', ',append=T)
      if (SaveWholeworkspace=='N') {rm(M.seq.tax.MinimumAllowedReadCount.Analysis.samples.kept.org,M.seq.tax.MinimumAllowedReadCount.Analysis.ASVs.excluded.org,M.seq.tax.MinimumAllowedReadCount.Analysis.ASVs.excluded)
        gc()}
    } else if (nrow(M.seq.tax.MinimumAllowedReadCount.Analysis.ASVs.excluded)==0&nrow(M.seq.tax.MinimumAllowedReadCount.Analysis.ASVs.excluded.org)==0) {                           # only happens if no ASV was excluded
      cat('\nNo ASVs were excluded because they do not occur in any of the samples anymore.',file=(file.path(PathToVisuaRAnalysis,paste(VisuaRProjectName,'_Analysis_log.txt',sep=''))),sep='',append=T)
      if (SaveWholeworkspace=='N') {rm(M.seq.tax.MinimumAllowedReadCount.Analysis.samples.kept.org,M.seq.tax.MinimumAllowedReadCount.Analysis.ASVs.excluded.org,M.seq.tax.MinimumAllowedReadCount.Analysis.ASVs.excluded)
        gc()}
    }
  } else if (ncol(M.seq.tax.MinimumAllowedReadCount.Analysis.samples.excluded)!=ncol.taxo.noNA&ncol(M.seq.tax.MinimumAllowedReadCount.Analysis.samples.excluded.org)==ncol.taxo.noNA) {   # only happens if at least one sample was excluded in the df M.seq.tax.subset but not in M.seq.tax
    M.seq.tax.subset=M.seq.tax.subset[,-which(numcolwise(sum)(M.seq.tax.subset)<MinimumAllowedReadCount.Analysis)]                                                                        # only keeps samples with more or equal to MinimumAllowedReadCount reads (based on subsetted df (M.seq.tax.subset))
    cat('\nThe following ',(ncol(M.seq.tax.MinimumAllowedReadCount.Analysis.samples.excluded)-ncol.taxo.noNA),' samples were exclulded based on low read counts:\n',file=(file.path(PathToVisuaRAnalysis,paste(VisuaRProjectName,'_Analysis_log.txt',sep=''))),sep='',append=T)
    cat(colnames(M.seq.tax.MinimumAllowedReadCount.Analysis.samples.excluded[1:((ncol(M.seq.tax.MinimumAllowedReadCount.Analysis.samples.excluded))-ncol.taxo.noNA)]),file=(file.path(PathToVisuaRAnalysis,paste(VisuaRProjectName,'_Analysis_log.txt',sep=''))),sep='\n',append=T)
    M.seq.tax.MinimumAllowedReadCount.Analysis.ASVs.excluded=M.seq.tax.subset[rowSums(M.seq.tax.subset[,1:(ncol(M.seq.tax.subset)-ncol.taxo.noNA)])==0,]                                  # creates an ASV by Sample&Taxonomy df with all ASVs which do not occur in the subsetted samples (based on the subsetted dataset (M.seq.tax.subset))
    if (SaveWholeworkspace=='N') {rm(M.seq.tax.MinimumAllowedReadCount.Analysis.samples.excluded.org,M.seq.tax.MinimumAllowedReadCount.Analysis.samples.excluded)
      gc()}
    if (nrow(M.seq.tax.MinimumAllowedReadCount.Analysis.ASVs.excluded)!=0) {                                                                                                              # only happens if at least one ASV was excluded
      M.seq.tax.subset=M.seq.tax.subset[rowSums(M.seq.tax.subset[,1:(ncol(M.seq.tax.subset)-ncol.taxo.noNA)])!=0,]                                                                        # excludes ASVs which do not occur anymore in the subsetted df
      cat('The following ',nrow(M.seq.tax.MinimumAllowedReadCount.Analysis.ASVs.excluded),' ASVs have been exclulded because they are no longer present in any of the samples:\n',file=(file.path(PathToVisuaRAnalysis,paste(VisuaRProjectName,'_Analysis_log.txt',sep=''))),sep='',append=T)
      cat(rownames(M.seq.tax.MinimumAllowedReadCount.Analysis.ASVs.excluded),file=(file.path(PathToVisuaRAnalysis,paste(VisuaRProjectName,'_Analysis_log.txt',sep=''))),sep=', ',append=T)
      if (SaveWholeworkspace=='N') {rm(M.seq.tax.MinimumAllowedReadCount.Analysis.ASVs.excluded)
        gc()}
    } else { # only happens if no ASV was excluded
      cat('\nNo ASVs were excluded because they do not occur in any of the samples anymore.',file=(file.path(PathToVisuaRAnalysis,paste(VisuaRProjectName,'_Analysis_log.txt',sep=''))),sep='',append=T)
      if (SaveWholeworkspace=='N') {rm(M.seq.tax.MinimumAllowedReadCount.Analysis.ASVs.excluded)
        gc()}
    } 
  } else if (ncol(M.seq.tax.MinimumAllowedReadCount.Analysis.samples.excluded)==ncol.taxo.noNA&ncol(M.seq.tax.MinimumAllowedReadCount.Analysis.samples.excluded.org)!=ncol.taxo.noNA) {   # only happens if at least one sample was excluded in the df M.seq.tax but not in M.seq.tax.subset
    M.seq.tax.MinimumAllowedReadCount.Analysis.samples.kept.org=M.seq.tax[,-which(numcolwise(sum)(M.seq.tax)<MinimumAllowedReadCount.Analysis)]                                           # only keeps samples with more or equal to MinimumAllowedReadCount reads (based on original df (M.seq.tax))
    cat('\nThe following ',(ncol(M.seq.tax.MinimumAllowedReadCount.Analysis.samples.excluded.org)-ncol.taxo.noNA),' samples were exclulded based on low read counts:\n',file=(file.path(PathToVisuaRAnalysis,paste(VisuaRProjectName,'_Analysis_log.txt',sep=''))),sep='',append=T)
    cat(colnames(M.seq.tax.MinimumAllowedReadCount.Analysis.samples.excluded.org[1:((ncol(M.seq.tax.MinimumAllowedReadCount.Analysis.samples.excluded.org))-ncol.taxo.noNA)]),file=(file.path(PathToVisuaRAnalysis,paste(VisuaRProjectName,'_Analysis_log.txt',sep=''))),sep='\n',append=T)
    M.seq.tax.MinimumAllowedReadCount.Analysis.ASVs.excluded.org=M.seq.tax.MinimumAllowedReadCount.Analysis.samples.kept.org[rowSums(M.seq.tax.MinimumAllowedReadCount.Analysis.samples.kept.org[,1:(ncol(M.seq.tax.MinimumAllowedReadCount.Analysis.samples.kept.org)-ncol.taxo.noNA)])==0,] # creates an ASV by Sample&Taxonomy df with all ASVs which do not occur in the  samples (based on the original dataset (M.seq.tax)
    if (SaveWholeworkspace=='N') {rm(M.seq.tax.MinimumAllowedReadCount.Analysis.samples.excluded.org,M.seq.tax.MinimumAllowedReadCount.Analysis.samples.excluded)
      gc()}
    if (nrow(M.seq.tax.MinimumAllowedReadCount.Analysis.ASVs.excluded.org)!=0) {                                                                                                          # only happens if at least one ASV was excluded
      M.seq.tax.MinimumAllowedReadCount.Analysis.samples.kept.org=M.seq.tax.MinimumAllowedReadCount.Analysis.samples.kept.org[rowSums(M.seq.tax.MinimumAllowedReadCount.Analysis.samples.kept.org[,1:(ncol(M.seq.tax.MinimumAllowedReadCount.Analysis.samples.kept.org)-ncol.taxo.noNA)])!=0,] #excludes ASVs which do not occur anymore in the original df
      cat('The following ',nrow(M.seq.tax.MinimumAllowedReadCount.Analysis.ASVs.excluded.org),' ASVs have been exclulded because they are no longer present in any of the samples:\n',file=(file.path(PathToVisuaRAnalysis,paste(VisuaRProjectName,'_Analysis_log.txt',sep=''))),sep='',append=T)
      cat(rownames(M.seq.tax.MinimumAllowedReadCount.Analysis.ASVs.excluded.org),file=(file.path(PathToVisuaRAnalysis,paste(VisuaRProjectName,'_Analysis_log.txt',sep=''))),sep=', ',append=T)
      if (SaveWholeworkspace=='N') {rm(M.seq.tax.MinimumAllowedReadCount.Analysis.samples.kept.org,M.seq.tax.MinimumAllowedReadCount.Analysis.ASVs.excluded.org)
        gc()}
    } else { # only happens if no ASV was excluded
      cat('\nNo ASVs were excluded because they do not occur in any of the samples anymore.',file=(file.path(PathToVisuaRAnalysis,paste(VisuaRProjectName,'_Analysis_log.txt',sep=''))),sep='',append=T)
      if (SaveWholeworkspace=='N') {rm(M.seq.tax.MinimumAllowedReadCount.Analysis.samples.kept.org,M.seq.tax.MinimumAllowedReadCount.Analysis.ASVs.excluded.org)
        gc()}
    }
  } else if (ncol(M.seq.tax.MinimumAllowedReadCount.Analysis.samples.excluded.org)==ncol.taxo.noNA&ncol(M.seq.tax.MinimumAllowedReadCount.Analysis.samples.excluded.org)==ncol.taxo.noNA) { # only happens if no sample was excluded in both df.
    cat('\nNo samples were excluded based on low read counts as all samples had more than ',MinimumAllowedReadCount.Analysis,' reads.',file=(file.path(PathToVisuaRAnalysis,paste(VisuaRProjectName,'_Analysis_log.txt',sep=''))),sep='',append=T)
    if (SaveWholeworkspace=='N') {rm(M.seq.tax.MinimumAllowedReadCount.Analysis.samples.excluded.org,M.seq.tax.MinimumAllowedReadCount.Analysis.samples.excluded)
      gc()}
  }
}
cat('\n',(ncol(M.seq.tax.subset)-ncol.taxo.noNA),' samples with a total of ',(nrow(M.seq.tax.subset)),' different ASVs out of originally ',nrow(M.seqtab.nochim),' samples with ',(ncol(M.seqtab.nochim)),' different ASVs remain for analysis.',file=(file.path(PathToVisuaRAnalysis,paste(VisuaRProjectName,'_Analysis_log.txt',sep=''))),sep='',append=T)

M.sample.reads.left.temp=colSums(M.seq.tax.subset[,1:(ncol(M.seq.tax.subset)-ncol.taxo.noNA)])
M.ASV.per.sample.temp=colSums(M.seq.tax.subset[,1:(ncol(M.seq.tax.subset)-ncol.taxo.noNA)]!=0)
infotable <- fill.info(df.reads=M.sample.reads.left.temp,df.ASVs =M.ASV.per.sample.temp,df.seqtab=M.seq.tax.subset,stepname=paste0("After excluding filtered samples with read counts lower than ",MinimumAllowedReadCount.Analysis,paste=""),infotable=infotable)
rm(M.sample.reads.left.temp,M.ASV.per.sample.temp)

gc()


# #=== 2.16. Excludes Samples based on a category in your map file ==========================================================================================================================

cat('\n\n2.16. Excludes Samples from your contextdata sheet based on a category in your map file.',file=(file.path(PathToVisuaRAnalysis,paste(VisuaRProjectName,'_Analysis_log.txt',sep=''))),sep='',append=TRUE)

if (ColumnOfVariableToExclude!='') {
  M.contextdata.CategoryToExcluded.samples.excluded=M.contextdata[grep(VariableToExclude,M.contextdata[,ColumnOfVariableToExclude],invert=F),]   # only takes samples belonging to the VariableToExclude
  M.contextdata=M.contextdata[grep(VariableToExclude,M.contextdata[,ColumnOfVariableToExclude],invert=TRUE),]                                    # Only takes samples not belonging to the VariableToExclude
  if (nrow(M.contextdata.CategoryToExcluded.samples.excluded)!=0) {                                                                        # Only happens if at least 1 sample was excluded
    cat('\nThe following ',nrow(M.contextdata.CategoryToExcluded.samples.excluded),' samples were excluded because they belong to the Category to exclude (',VariableToExclude,') found in the column ',ColumnOfVariableToExclude,' in your contextdata.\n',file=(file.path(PathToVisuaRAnalysis,paste(VisuaRProjectName,'_Analysis_log.txt',sep=''))),sep='',append=TRUE)
    cat(rownames(M.contextdata.CategoryToExcluded.samples.excluded),file=(file.path(PathToVisuaRAnalysis,paste(VisuaRProjectName,'_Analysis_log.txt',sep=''))),sep="\n",append=TRUE)
    if (SaveWholeworkspace=='N') {rm(M.contextdata.CategoryToExcluded.samples.excluded)
      gc()}
  } else {
    cat('\nNo samples were excluded because none of the sample in your contextdata belong to the Category to exclude (',VariableToExclude,') found in the column ',ColumnOfVariableToExclude,' in your contextdata.\n',file=(file.path(PathToVisuaRAnalysis,paste(VisuaRProjectName,'_Analysis_log.txt',sep=''))),sep='',append=TRUE)
    if (SaveWholeworkspace=='N') {rm(M.contextdata.CategoryToExcluded.samples.excluded)
      gc()}
  }
} else {
  cat('\nNo samples will be excluded as no VariableToExclude was chosen. \n',file=(file.path(PathToVisuaRAnalysis,paste(VisuaRProjectName,'_Analysis_log.txt',sep=''))),sep='',append=TRUE)
}

M.sample.reads.left.temp=colSums(M.seq.tax.subset[,1:(ncol(M.seq.tax.subset)-ncol.taxo.noNA)])
M.ASV.per.sample.temp=colSums(M.seq.tax.subset[,1:(ncol(M.seq.tax.subset)-ncol.taxo.noNA)]!=0)
infotable <- fill.info(df.reads=M.sample.reads.left.temp,df.ASVs =M.ASV.per.sample.temp,df.seqtab=M.seq.tax.subset,stepname=paste0("After excluding samples based on a category in your contextual data",sep=""),infotable=infotable)
rm(M.sample.reads.left.temp,M.ASV.per.sample.temp)

gc()

# #=== 2.16.2. Subsets Samples based on a category in your map file ==========================================================================================================================

cat('\n\n2.16. Keep Samples from your contextdata sheet based on a category in your map file (VariableToKeep).',file=(file.path(PathToVisuaRAnalysis,paste(VisuaRProjectName,'_Analysis_log.txt',sep=''))),sep='',append=TRUE)

if (ColumnOfVariableToKeep!='') {
  M.contextdata.CategoryToKeep.samples.excluded=M.contextdata[grep(VariableToKeep,M.contextdata[,ColumnOfVariableToKeep],invert=T),]   # only takes samples belonging to the VariableToKeep
  M.contextdata=M.contextdata[grep(VariableToKeep,M.contextdata[,ColumnOfVariableToKeep],invert=F),]                                    # Only takes samples not belonging to the VariableToKeep
  if (nrow(M.contextdata.CategoryToKeep.samples.excluded)!=0) {                                                                        # Only happens if at least 1 sample was excluded
    cat('\nThe following ',nrow(M.contextdata.CategoryToKeep.samples.excluded),' samples were excluded because they do not belong to the Category to keep (',VariableToKeep,') found in the column ',ColumnOfVariableToKeep,' in your contextdata.\n',file=(file.path(PathToVisuaRAnalysis,paste(VisuaRProjectName,'_Analysis_log.txt',sep=''))),sep='',append=TRUE)
    cat(rownames(M.contextdata.CategoryToKeep.samples.excluded),file=(file.path(PathToVisuaRAnalysis,paste(VisuaRProjectName,'_Analysis_log.txt',sep=''))),sep="\n",append=TRUE)
    if (SaveWholeworkspace=='N') {rm(M.contextdata.CategoryToKeep.samples.excluded)
      gc()}
  } else {
    cat('\nNo samples were excluded because none of the sample in your contextdata do not belong to the Category to keep (',VariableToKeep,') found in the column ',ColumnOfVariableToKeep,' in your contextdata.\n',file=(file.path(PathToVisuaRAnalysis,paste(VisuaRProjectName,'_Analysis_log.txt',sep=''))),sep='',append=TRUE)
    if (SaveWholeworkspace=='N') {rm(M.contextdata.CategoryToKeep.samples.excluded)
      gc()}
  }
} else {
  cat('\nNo samples will be excluded as no VariableToKeep was chosen. \n',file=(file.path(PathToVisuaRAnalysis,paste(VisuaRProjectName,'_Analysis_log.txt',sep=''))),sep='',append=TRUE)
}

M.sample.reads.left.temp=colSums(M.seq.tax.subset[,1:(ncol(M.seq.tax.subset)-ncol.taxo.noNA)])
M.ASV.per.sample.temp=colSums(M.seq.tax.subset[,1:(ncol(M.seq.tax.subset)-ncol.taxo.noNA)]!=0)
infotable <- fill.info(df.reads=M.sample.reads.left.temp,df.ASVs =M.ASV.per.sample.temp,df.seqtab=M.seq.tax.subset,stepname=paste0("After only keeping samples based on a category in your contextual data",sep=""),infotable=infotable)
rm(M.sample.reads.left.temp,M.ASV.per.sample.temp)

gc()

# #=== 2.17. Excludes Samples based on selected grouping parameters ===========================================================================================

cat('\n2.17. Excludes Samples based on selected grouping parameters.',file=(file.path(PathToVisuaRAnalysis,paste(VisuaRProjectName,'_Analysis_log.txt',sep=''))),sep='',append=TRUE)
M.contextdata.subset=M.contextdata
if (SaveWholeworkspace=='N') {rm(M.contextdata)
  gc()}
M.contextdata.subset[M.contextdata.subset=="NA"]=NA
MapCol1=which(names(M.contextdata.subset)==Grouping1) # finds the number of the column of Grouping1 in your contextdata
MapCol2=which(names(M.contextdata.subset)==Grouping2) # finds the number of the column of Grouping2 in your contextdata

cat('\nYour samples will be grouped  based on values in the parameter ',Grouping1,' (Grouping1) and sorted inside these groups based on the parameter ',Grouping2,' (Grouping2).',file=(file.path(PathToVisuaRAnalysis,paste(VisuaRProjectName,'_Analysis_log.txt',sep=''))),sep='',append=TRUE)

M.contextdata.subset.NA=subset(M.contextdata.subset,is.na(M.contextdata.subset[,MapCol1])|is.na(M.contextdata.subset[,MapCol2])) # creates a Sample by contextdata file of samples which have NAs in one of the selected Groupings
if (nrow(M.contextdata.subset.NA)==0){ 
  cat('\nNo sample was excluded from your contextual data file due to missing contextual data in the chosen groupings.',file=(file.path(PathToVisuaRAnalysis,paste(VisuaRProjectName,'_Analysis_log.txt',sep=''))),sep='',append=TRUE)
} else {
  cat('\nThe following ',nrow(M.contextdata.subset.NA),' samples were excluded from your contextual data sheet because they do not have contextual data in the selected groupings: \n',file=(file.path(PathToVisuaRAnalysis,paste(VisuaRProjectName,'_Analysis_log.txt',sep=''))),sep='',append=TRUE)
  cat(rownames(M.contextdata.subset.NA),file=(file.path(PathToVisuaRAnalysis,paste(VisuaRProjectName,'_Analysis_log.txt',sep=''))),sep="\n",append=TRUE)
  cat('No contextdata in Grouping1: ',file=(file.path(PathToVisuaRAnalysis,paste(VisuaRProjectName,'_Analysis_log.txt',sep=''))),sep='',append=TRUE)
  cat(rownames(subset(M.contextdata.subset,is.na(M.contextdata.subset[,MapCol1]))),file=(file.path(PathToVisuaRAnalysis,paste(VisuaRProjectName,'_Analysis_log.txt',sep=''))),sep=", ",append=TRUE)
  cat('\nNo contextdata in Grouping2: ',file=(file.path(PathToVisuaRAnalysis,paste(VisuaRProjectName,'_Analysis_log.txt',sep=''))),sep='',append=TRUE)
  cat(rownames(subset(M.contextdata.subset,is.na(M.contextdata.subset[,MapCol2]))),file=(file.path(PathToVisuaRAnalysis,paste(VisuaRProjectName,'_Analysis_log.txt',sep=''))),sep=", ",append=TRUE)
}
if (SaveWholeworkspace=='N') {rm(M.contextdata.subset.NA);gc()}

M.contextdata.subset.noNA=subset(M.contextdata.subset,!is.na(M.contextdata.subset[,MapCol1])&!is.na(M.contextdata.subset[,MapCol2])) # Creates contextdata df without NAs in either of the Grouping columns

gc()

# #=== 2.18. Subsets contextdata to fit to your subsetted sequencing data (M.seq.tax.subset) ===================================================================================================

cat('\n\n2.18. Subsets contextdata to fit to your subsetted sequencing data (M.seq.tax.subset).',file=(file.path(PathToVisuaRAnalysis,paste(VisuaRProjectName,'_Analysis_log.txt',sep=''))),sep='',append=TRUE)

M.contextdata.subset=M.contextdata.subset.noNA[match((as.character(colnames(M.seq.tax.subset[1:(ncol(M.seq.tax.subset)-ncol.taxo.noNA)]))),rownames(M.contextdata.subset.noNA)),] # Only keeps samples with the same names as those occuring in the M.seq.tax.subset. This creates empty rows for those samples which do occur in the M.seq.tax.subset but do not occur in M.contextdata.subset.noNAs. Does not take the taxonomy in M.seq.tax.subset into account.
M.contextdata.subset=subset(M.contextdata.subset,!is.na(M.contextdata.subset[,MapCol1]),)  # deletes newly introduced NA columns

if (nrow(M.contextdata.subset.noNA)!=nrow(M.contextdata.subset)) {                      # only happens if at least one sample does not occur in the contextdata which does occur in M.seq.tax.subset and vice versa
  KeepRows=!match(rownames(M.contextdata.subset.noNA),(as.character(colnames(M.seq.tax.subset[1:(ncol(M.seq.tax.subset)-ncol.taxo.noNA)])))) # this is a detour to only keep the samples which do not match to the samples in the M.seq.tax.subset because using !match does result in an empty df
  M.contextdata.subset.noseq=cbind(M.contextdata.subset.noNA,KeepRows)
  M.contextdata.subset.noseq=subset(M.contextdata.subset.noseq,is.na(M.contextdata.subset.noseq[,ncol(M.contextdata.subset.noseq)]))
  M.contextdata.subset.noseq=M.contextdata.subset.noseq[,-ncol(M.contextdata.subset.noseq)]
  cat('\nThe following ',nrow(M.contextdata.subset.noseq),' samples have been excluded from your contextdata because they do not occur in your subsetted M.seq.tax.subset: \n',file=(file.path(PathToVisuaRAnalysis,paste(VisuaRProjectName,'_Analysis_log.txt',sep=''))),sep='',append=TRUE)
  cat(rownames(M.contextdata.subset.noseq),file=(file.path(PathToVisuaRAnalysis,paste(VisuaRProjectName,'_Analysis_log.txt',sep=''))),sep="\n",append=TRUE)
  if (SaveWholeworkspace=='N') {rm(M.contextdata.subset.noseq,KeepRows)
    gc()}
} else {
  cat('\nNo samples have been excluded from your contextdata because all samples in your contextdata occur in your M.seq.tax.subset.',file=(file.path(PathToVisuaRAnalysis,paste(VisuaRProjectName,'_Analysis_log.txt',sep=''))),sep='',append=TRUE)
}

M.contextdata.subset.noNA=M.contextdata.subset
if (SaveWholeworkspace=='N') {rm(M.contextdata.subset)}
gc()

# #=== 2.19. Subsets M.seq.tax.subset to samples where contextdata is available for the selected grouping parameters. ==========================================================================

cat('\n\n2.19. Subsets M.seq.tax.subset to samples where contextdata is available for the selected grouping parameters.',file=(file.path(PathToVisuaRAnalysis,paste(VisuaRProjectName,'_Analysis_log.txt',sep=''))),sep='',append=TRUE)
M.seq.tax.subset.nocontextdata=M.seq.tax.subset[,-match(rownames(M.contextdata.subset.noNA),(colnames(M.seq.tax.subset)))] # creates table from M.seq.tax.subset only containing samples which do not have contextdata available

if(ncol(M.seq.tax.subset.nocontextdata)==0){  # only happens if no samples was excluded (all samples occur in the subsetted contextdatafile.)
  cat('\nNo samples have been excluded from your sequencing data because they do not occur in your contextdata file (after subsetting).',file=(file.path(PathToVisuaRAnalysis,paste(VisuaRProjectName,'_Analysis_log.txt',sep=''))),sep='',append=TRUE)
} else {                                                # happens if at least one sample was excluded
  M.seq.tax.subset=cbind(M.seq.tax.subset[,match(rownames(M.contextdata.subset.noNA),(colnames(M.seq.tax.subset)))],M.seq.tax.subset[,(ncol(M.seq.tax.subset)-ncol.taxo.noNA+1):ncol(M.seq.tax.subset)]) # creates table from M.seq.tax.subset only containing samples which do have contextdata available
  cat('\nThe following ',(ncol(M.seq.tax.subset.nocontextdata)-ncol.taxo.noNA) ,' samples have been excluded from your sequencing data because they do not occur in your contextdatafile or have no data for your selected groupings (after subsetting).\n',file=(file.path(PathToVisuaRAnalysis,paste(VisuaRProjectName,'_Analysis_log.txt',sep=''))),sep='',append=TRUE)
  cat(colnames(M.seq.tax.subset.nocontextdata[,1:(ncol(M.seq.tax.subset.nocontextdata)-ncol.taxo.noNA)]),file=(file.path(PathToVisuaRAnalysis,paste(VisuaRProjectName,'_Analysis_log.txt',sep=''))),sep="\n",append=TRUE)
  cat('\nThe following ',(ncol(M.seq.tax.subset)-ncol.taxo.noNA) ,' samples out of originally ' ,(ncol(M.seq.tax)-ncol.taxo.noNA),'samples remain for further analysis\n',file=(file.path(PathToVisuaRAnalysis,paste(VisuaRProjectName,'_Analysis_log.txt',sep=''))),sep='',append=TRUE)
  cat(colnames(M.seq.tax.subset[,1:(ncol(M.seq.tax.subset)-ncol.taxo.noNA)]),file=(file.path(PathToVisuaRAnalysis,paste(VisuaRProjectName,'_Analysis_log.txt',sep=''))),sep='\n',append=TRUE)
}
if (SaveWholeworkspace=='N') {rm(M.seq.tax.subset.nocontextdata)}

M.seq.tax.contextdata.ASVs.excluded=M.seq.tax.subset[rowSums(M.seq.tax.subset[,1:(ncol(M.seq.tax.subset)-ncol.taxo.noNA)])==0,] # creates ASV by Sample&Taxonomy df with all the ASVs which do not occur anymore in any of the samples

# # give out overview of excluded samples
exc.samples <- setdiff(colnames(M.seq.tax),colnames(M.seq.tax.subset))
cat('\nFrom your original sequencing data, the following ',length(exc.samples),' samples out of originally ',ncol(M.seq.tax)-ncol.taxo.noNA,' samples were excluded due to several reasons (see above for details):\n',file=(file.path(PathToVisuaRAnalysis,paste(VisuaRProjectName,'_Analysis_log.txt',sep=''))),sep='',append=TRUE)
cat(exc.samples,file=(file.path(PathToVisuaRAnalysis,paste(VisuaRProjectName,'_Analysis_log.txt',sep=''))),sep='\n',append=TRUE)

M.sample.reads.left.temp=colSums(M.seq.tax.subset[,1:(ncol(M.seq.tax.subset)-ncol.taxo.noNA)])
M.ASV.per.sample.temp=colSums(M.seq.tax.subset[,1:(ncol(M.seq.tax.subset)-ncol.taxo.noNA)]!=0)
infotable <- fill.info(df.reads=M.sample.reads.left.temp,df.ASVs =M.ASV.per.sample.temp,df.seqtab=M.seq.tax.subset,stepname=paste0("After keeping only samples with contextual data available in the selected grouping",sep=""),infotable=infotable)
rm(M.sample.reads.left.temp,M.ASV.per.sample.temp)

gc()

# #=== 2.20 Creates Group and Color vectors to represent the project/samples based on projects/treatments ===================================================================================

cat('\n\n2.20. Creates Group and Color vectors for the visualization.',file=(file.path(PathToVisuaRAnalysis,paste(VisuaRProjectName,'_Analysis_log.txt',sep=''))),sep='',append=TRUE)

M.projects=as.vector(M.contextdata.subset.noNA[,MapCol1],mode = "character")      # vector defining groups/projects based on MapCol1. In original order
M.projects.unique=unique(M.projects)                        # vector/list of individual projects, in original order
M.projects.unique.ord=sort(M.projects.unique, decreasing=F) # vector/list of individual projects sorted alphabetically
saveRDS(M.projects.unique.ord,file.path(PathToVisuaRAnalysis,paste(VisuaRProjectName,'_Mprojectsuniqueord.rds',sep='')))

if (exists("M.col")==FALSE && length(M.projects.unique.ord)<9 || (exists("M.col")==TRUE && length(M.projects.unique.ord)!=length(M.col))) {
  M.palette=c("#56B4E9","#E69F00", "#009E73", "#CC79A7","#999999", "#0072B2", "#D55E00","#F0E442") 
  M.col=M.palette[1:length(M.projects.unique)]              # This palette is matched to Grouping1 in alphabetical order
  rm(M.palette)
} else if (exists("M.col")==FALSE && length(M.projects.unique.ord)>8) {
  M.col=rainbow(length(M.projects.unique.ord))
} 

saveRDS(M.col,file.path(PathToVisuaRAnalysis,paste(VisuaRProjectName,'_Mcol.rds',sep='')))

M.col.names=(sapply(M.col,color.id))                        # gives names for the colors
if (!is.matrix(M.col.names)) {
  M.col.names=ldply (M.col.names, data.frame)               # if more than one name for at least one color a list will be created, will be transformed to a data.frame here
  M.col.names=M.col.names[!duplicated((M.col.names$.id)),]  # this removes duplicate values
} else {
  M.col.names=t(M.col.names)
}
colnames(M.col.names)=c('HexCode','Name') 

M.match.col=cbind(M.projects.unique.ord,M.col)              # table that assigns a specific color to a sample. In alphabetical order
M.match.col.names=cbind(M.projects.unique.ord,M.col.names)  # table that assigns a specific color to a sample and shows the name of the color (alphabetical order)

M.groups=match(M.projects, M.match.col)                                       # converts project names to values (in alphabetical order (A=1,B=2....)) This vector is in the original order
M.colvec=mapvalues(M.projects, from = M.match.col[,1], to = M.match.col[,2])  # function in plyr, recodes sample vector. Takes values in M.project and transforms them to the corresponding colors as found in M.match col. This vector is in the original order
saveRDS(M.colvec,file.path(PathToVisuaRAnalysis,paste(VisuaRProjectName,'_Mcolvec.rds',sep='')))
saveRDS(M.groups,file.path(PathToVisuaRAnalysis,paste(VisuaRProjectName,'_Mgroups.rds',sep='')))

M.group.count=matrix(NA,nrow=length(M.projects.unique),ncol=2)  # matrix to be filled by the function with as many rows as unique projects and 2 columns
M.projects.unique <- as.character(M.projects.unique)
for (i in 1:length(M.projects.unique)) {                        # Counts how many samples belong to each Group 
  M.group.count[i,1]=M.projects.unique[i]
  M.group.count[i,2]=sum(str_count(M.projects,pattern=M.projects.unique[i])) 
}
M.group.count=as.data.frame(M.group.count)                  # needs to be transformed to a dataframe to order
M.group.count.ord=M.group.count[order(M.group.count$V1),]   # data frame of individual projects together with number of how often this group appears in the subsorted data sorted alphabetically
M.group.count.ord=as.matrix(M.group.count.ord)              # needs to be transformed back to a matrix to work for the txt output.
cat('\nYour chosen groups subsetted for VisuaR contain in total ',length(M.projects),' samples.',file=(file.path(PathToVisuaRAnalysis,paste(VisuaRProjectName,'_Analysis_log.txt',sep=''))),sep='',append=TRUE)
for (i in 1:length(M.projects.unique)) {
  cat('\nGroup ',M.group.count.ord[i,1],' contains ',M.group.count.ord[i,2],' samples and will appear colored in ',as.character(M.match.col.names[i,3]),' (',M.match.col.names[i,2],').',file=(file.path(PathToVisuaRAnalysis,paste(VisuaRProjectName,'_Analysis_log.txt',sep=''))),sep='',append=T)
}
if (SaveWholeworkspace=='N') {rm(M.match.col.names)}
gc()
saveRDS(M.group.count.ord,file.path(PathToVisuaRAnalysis,paste(VisuaRProjectName,"_Mgroupcountord.rds",sep="")))

# #=== 2.22. Subsampling finished ==========================================================================================================================================================

cat('\n\n2.22. The subsampling is finished.',file=(file.path(PathToVisuaRAnalysis,paste(VisuaRProjectName,'_Analysis_log.txt',sep=''))),sep='',append=TRUE)
if (nrow(M.seq.tax.contextdata.ASVs.excluded)!=0) { # Only happens if a sample was excluded and some of the ASVs now do not occur anymore
  M.seq.tax.subset=M.seq.tax.subset[rowSums(M.seq.tax.subset[,1:(ncol(M.seq.tax.subset)-ncol.taxo.noNA)])!=0,]
  cat('\nThe following ',nrow(M.seq.tax.contextdata.ASVs.excluded),' ASVs have been excluded because they are no longer present in any of the samples.\n',file=(file.path(PathToVisuaRAnalysis,paste(VisuaRProjectName,'_Analysis_log.txt',sep=''))),sep='',append=TRUE)
  cat(rownames(M.seq.tax.contextdata.ASVs.excluded),file=(file.path(PathToVisuaRAnalysis,paste(VisuaRProjectName,'_Analysis_log.txt',sep=''))),sep=",",append=TRUE)
} else {
  cat('\nNo ASVs have been excluded because they are no longer present in any of the samples.',file=(file.path(PathToVisuaRAnalysis,paste(VisuaRProjectName,'_Analysis_log.txt',sep=''))),sep='',append=TRUE)
}
if (SaveWholeworkspace=='N') {rm(M.seq.tax.contextdata.ASVs.excluded)}
gc()
cat('\n\nThe following ',(ncol(M.seq.tax.subset)-ncol.taxo.noNA),' samples with a total of ',(nrow(M.seq.tax.subset)),' different ASVs out of originally ',nrow(M.seqtab.nochim),' samples with ',(ncol(M.seqtab.nochim)),' different ASVs remain for analysis.',file=(file.path(PathToVisuaRAnalysis,paste(VisuaRProjectName,'_Analysis_log.txt',sep=''))),sep='',append=T)
cat('\n',colnames(M.seq.tax.subset[,1:(ncol(M.seq.tax.subset)-ncol.taxo.noNA)]),file=(file.path(PathToVisuaRAnalysis,paste(VisuaRProjectName,'_Analysis_log.txt',sep=''))),sep=',',append=T)
if (SaveWholeworkspace=='N') {rm(M.seqtab.nochim)}
gc()

# # Reads of the remaining samples
M.sample.reads.left=colSums(M.seq.tax.subset[,1:(ncol(M.seq.tax.subset)-ncol.taxo.noNA)]) # creates a named numeric with the reads per sample
cat('\n\nThe remaining ',(ncol(M.seq.tax.subset)-ncol.taxo.noNA),' samples contain in total ',sum(M.sample.reads.left),' reads.\n',file=(file.path(PathToVisuaRAnalysis,paste(VisuaRProjectName,'_Analysis_log.txt',sep=''))),sep='',append=T)
printinfo(M.sample.reads.left)

# # ASVs per sample
M.ASV.per.sample=colSums(M.seq.tax.subset[,1:(ncol(M.seq.tax.subset)-ncol.taxo.noNA)]!=0) # creates a named numeric with the number of different ASVs per sample
cat('\nThe remaining ',(ncol(M.seq.tax.subset)-ncol.taxo.noNA),' samples contain in total ',nrow(M.seq.tax.subset),' different ASVs,\n',file=(file.path(PathToVisuaRAnalysis,paste(VisuaRProjectName,'_Analysis_log.txt',sep=''))),sep='',append=T)
printinfo(M.ASV.per.sample)


infotable <- fill.info(df.reads=M.sample.reads.left,df.ASVs =M.ASV.per.sample,df.seqtab=M.seq.tax.subset,stepname="End",infotable=infotable)
data.table::fwrite(infotable,file=file.path(PathToVisuaRAnalysis,paste(VisuaRProjectName,'_ReadsandASVs_Track.csv',sep='')),col.names = T,row.names = T)

# # Reads of the remaining ASVs
M.ASV.reads.left=rowSums(M.seq.tax.subset[,1:(ncol(M.seq.tax.subset)-ncol.taxo.noNA)]) # creates a named numeric with the reads per ASV
cat('\nThe remaining ',length(M.ASV.reads.left),' ASVs occur on average ',round(mean(M.ASV.reads.left),0),' times.\n',file=(file.path(PathToVisuaRAnalysis,paste(VisuaRProjectName,'_Analysis_log.txt',sep=''))),sep='',append=T)
printinfo(M.ASV.reads.left) 

closeAllConnections() # closes all currently open connections.

gc()

# #=== 2b. Transformation of input data==========================================================================================================================================================

# if(TransformInput=="clr"){
#   M.seq.tax.subset.clr1<-cbind(t(compositions::clr(t(M.seq.tax.subset[,1:(ncol(M.seq.tax.subset)-ncol.taxo.noNA)])+1)),M.seq.tax.subset[,((ncol(M.seq.tax.subset)-ncol.taxo.noNA)+1):ncol(M.seq.tax.subset)])
#   # test2<-cbind(t(compositions::clr(t(M.seq.tax.subset[,1:(ncol(M.seq.tax.subset)-ncol.taxo.noNA)]))+1),M.seq.tax.subset[,((ncol(M.seq.tax.subset)-ncol.taxo.noNA)+1):ncol(M.seq.tax.subset)])
#   # cat('\nNo samples have been excluded from your sequencing data because they do not occur in your contextdata file (after subsetting).',file=(file.path(PathToVisuaRAnalysis,paste(VisuaRProjectName,'_Analysis_log.txt',sep=''))),sep='',append=TRUE)
# } else {                                                
#   # cat(colnames(M.seq.tax.subset[,1:(ncol(M.seq.tax.subset)-ncol.taxo.noNA)]),file=(file.path(PathToVisuaRAnalysis,paste(VisuaRProjectName,'_Analysis_log.txt',sep=''))),sep='\n',append=TRUE)
# }

# #=== 3. Community Composition ============================================================================================================================

cat('\n\n\n3. Community Composition',file=(file.path(PathToVisuaRAnalysis,paste(VisuaRProjectName,'_Analysis_log.txt',sep=''))),sep='',append=TRUE)

# #=== 3.1. Creates relative abundance, presence absence tables ============================================================================================================================

cat('\n3.1. Creates presence/absence, relative abundance and other tables.',file=(file.path(PathToVisuaRAnalysis,paste(VisuaRProjectName,'_Analysis_log.txt',sep=''))),sep='',append=TRUE)

# # This file includes ASV identities in the first column, followed by respective ASV frequencies, and finally respective ASV taxonomy in the last column
M.seq.tax.subset.print=cbind(rownames(M.seq.tax.subset),data.frame(M.seq.tax.subset,row.names = NULL))
colnames(M.seq.tax.subset.print)[1]="ASV"
data.table::fwrite(M.seq.tax.subset.print,file=file.path(PathToVisuaRAnalysis,"Alpha_Diversity",paste(VisuaRProjectName,"_ASVbySample_abund_final.csv",sep="")),col.names = T,row.names = T)
if (SaveWholeworkspace=='N') {rm(M.seq.tax.subset.print)}
gc()

# # Split in two dataframes (Taxonomy and ASVs)
M.seq.tax.subset.ASVs=M.seq.tax.subset[,1:(ncol(M.seq.tax.subset)-ncol.taxo.noNA)]                          # creates ASV by sample table without taxonomy
M.seq.tax.subset.ASVs=as.data.frame(t(M.seq.tax.subset.ASVs)) 
M.seq.tax.subset.tax=M.seq.tax.subset[,c((ncol(M.seq.tax.subset)-ncol.taxo.noNA+1):ncol(M.seq.tax.subset))] # creates ASV by Taxonomy table

# # merge into one table with relative abundances and Taxonomy
M.ASV.rel=decostand(M.seq.tax.subset.ASVs, method="total")                    # vegan package, calculates relative abundance
# M.ASV.rel.clr=decostand(M.seq.tax.subset.ASVs, method = "rclr")                    # vegan package, calculates relative abundance
M.seq.tax.subset.rel=cbind(as.data.frame(t(M.ASV.rel)),M.seq.tax.subset.tax)  # transforms table and adds taxonomy back to the table
M.seq.tax.subset.rel=cbind(rownames(M.seq.tax.subset.rel),data.frame(M.seq.tax.subset.rel,row.names = NULL)) # modifies table for txt output
colnames(M.seq.tax.subset.rel)[1]="ASV"
data.table::fwrite(M.seq.tax.subset.rel,file=file.path(PathToVisuaRAnalysis,"Alpha_Diversity",paste(VisuaRProjectName,"_ASVbySample_relabund_final.csv",sep="")),col.names = T,row.names = T) # _Mgroups.rds is in similar order regarding samples
if (SaveWholeworkspace=='N') {rm(M.ASV.rel,M.seq.tax.subset.rel)}
gc()

# # merge into one table with presence absence and taxonomy
M.ASV.pa=decostand(M.seq.tax.subset.ASVs, method="pa")                                                    # vegan package, calculates presence absence table
M.seq.tax.subset.pa=cbind(as.data.frame(t(M.ASV.pa)),M.seq.tax.subset.tax)                                # transforms table and adds taxonomy back to the table
M.seq.tax.subset.pa=cbind(rownames(M.seq.tax.subset.pa),data.frame(M.seq.tax.subset.pa,row.names = NULL)) # modifies table for txt output
colnames(M.seq.tax.subset.pa)[1]="ASV"
data.table::fwrite(M.seq.tax.subset.pa,file=file.path(PathToVisuaRAnalysis,"Alpha_Diversity",paste(VisuaRProjectName,"_ASVbySample_pa_final.csv",sep="")),col.names = T,row.names = T)
if (SaveWholeworkspace=='N') {rm(M.ASV.pa,M.seq.tax.subset.ASVs,M.seq.tax.subset.pa,M.seq.tax.subset.tax)}
gc()

# # Calculates Reads per ASV after subsetting.
M.ASV.reads.left=as.data.frame(M.ASV.reads.left)
M.ASV.reads.left=cbind('ASV'=rownames(M.ASV.reads.left),data.frame(M.ASV.reads.left,row.names = NULL))
colnames(M.ASV.reads.left)[2]="Reads"
data.table::fwrite(M.ASV.reads.left,file=file.path(PathToVisuaRAnalysis,"Alpha_Diversity",paste(VisuaRProjectName,"_ReadsperASV_final.csv",sep="")),col.names = T,row.names = T)
if (SaveWholeworkspace=='N') {rm(M.ASV.reads.left)}

gc()
# #=== 3.2. Creates M.seq.tax.subset.ord: Ordered based on taxonomic levels and based on Grouping ==========================================================================================

# # Orders rows based on taxonomy (A-Z)
ncol.M.seq.tax.subset=ncol(M.seq.tax.subset) # counts how many samples plus taxonomy are in the subsetted M.seq.tax.subset
# Generate a list of column indices for ordering
orderCols <- (ncol(M.seq.tax.subset) - ncol.taxo.noNA + 1):(ncol(M.seq.tax.subset) - ncol.taxo.noNA + ncol.taxo.noNA)
# Order the table based on the specified columns (alphabetically with Phylum, etc.)
M.seq.tax.subset.ord <- M.seq.tax.subset[do.call(order, M.seq.tax.subset[, orderCols]), ]

if (SaveWholeworkspace=='N') {rm(M.seq.tax.subset)}
gc()

# #Orders M.seq.tax columns based on projects/groups/contextual data (A-Z or 1-n) 
M.seq.tax.subset.ord.ASV=M.seq.tax.subset.ord[,1:(ncol.M.seq.tax.subset-ncol.taxo.noNA)]                                      # makes ASV table without taxonomy, ordered based on taxonomy
M.seq.tax.subset.ord.tax=M.seq.tax.subset.ord[,c((ncol.M.seq.tax.subset-ncol.taxo.noNA+1):ncol.M.seq.tax.subset)]             # make taxonomy table, alphabetical ordered
M.contextdata.subset.noNA.ord=M.contextdata.subset.noNA[order(M.contextdata.subset.noNA[,MapCol1],M.contextdata.subset.noNA[,MapCol2]),]  # orders contextdata alphabetically based on the provided Groupings
M.seq.tax.subset.ord.ASV.ord=M.seq.tax.subset.ord.ASV[,rownames(M.contextdata.subset.noNA.ord)]                                  # Orders ASV table based on ordered contextdata (columns are ordered as they are ordered as rows in M.contextdata.subset.noNA.ord)
M.seq.tax.subset.ord=cbind(M.seq.tax.subset.ord.ASV.ord,M.seq.tax.subset.ord.tax)  # Rows=ASVs, ordered alphabetically for the taxonomy. Columns=Samples, ordered alphabetically depending on the group they belong to.
saveRDS(M.seq.tax.subset.ord,file.path(PathToVisuaRAnalysis,'Beta_Diversity',paste(VisuaRProjectName,"_Mseqtaxsubsetord.rds",sep="")))
if (SaveWholeworkspace=='N') {rm(M.seq.tax.subset.ord.ASV.ord,M.contextdata.subset.noNA.ord)}
gc()

# # This workspace can be used in combination with VisuaR_CladeTables.R
save.image(file.path(PathToVisuaRAnalysis,paste(VisuaRProjectName,'_VisuaR_inc_3_2','.RData',sep='')))

# #=== 3.3. Creates tables using individual taxonomic levels ===============================================================================================================================

cat('\n\n3.3. Creates tables using individual taxonomic levels',file=(file.path(PathToVisuaRAnalysis,paste(VisuaRProjectName,'_Analysis_log.txt',sep=''))),sep='',append=TRUE)
CompositionTablePath=file.path(PathToVisuaRAnalysis,"Alpha_Diversity","CompositionTables")
dir.create(file.path(PathToVisuaRAnalysis,"Alpha_Diversity","CompositionTables"))
cat('\nThe tables will be saved to ',CompositionTablePath,'.\nFor each taxonomic level 4 tables will be created.\n1. ..TaxonomicLevelBySamples_abund.txt: A Taxonomy by sample table with all reads belonging to the clades summed up and the average of the clades in the last column.\n2. ..TaxonomicLevelBySamples_abund.sort.txt: A Taxonomy by sample table with all reads belonging to the clades summed up and the average of the clades in the last column. Depending on the value set in NumberOfTOPClades the rare taxa will be summed up in the last row and named as others. The Clades are ordered based on descending means.\n3. ..TaxonomicLevelBySamples_relabund_sort.txt: Same as 2. but wiht relative abundances instead of Reads.\n4. ..TaxonomicLevelBySamples_pa.sort.txt: Same as 2. but wiht presences/absences instead of Reads.',file=(file.path(PathToVisuaRAnalysis,paste(VisuaRProjectName,'_Analysis_log.txt',sep=''))),sep='',append=TRUE)

# #--- 3.3.1 Taxonomic level tables ----------------------------------------------------------------------------------------------------------------------------------------------------------
# #  create lists to be filled with information on the different taxonomic levels
tax.levels <- colnames(M.seq.tax)[(ncol(M.seq.tax)-ncol.taxo.noNA+1):ncol(M.seq.tax)] # names of the taxonomic levels
tax.levels <- lapply(tax.levels, tolower)    # makes list of the names
tax.levels.tables.list <- vector("list",length(tax.levels)); names(tax.levels.tables.list) <- paste0("M.seq.tax.subset.",tax.levels)
tax.levels.tables.list.group <- vector("list",length(tax.levels)); names(tax.levels.tables.list) <- paste0("M.seq.tax.subset.group",tax.levels)

tax.levels.tables.sums.list <- vector("list",length(tax.levels)); names(tax.levels.tables.sums.list) <- paste0("M.seq.tax.subset.",tax.levels,".sums")
tax.levels.tables.indicators.list <- vector("list",length(tax.levels)); names(tax.levels.tables.indicators.list) <- paste0("Indicator",tax.levels) #tax.levels.tables.indicators <- paste0("Indicator",tax.levels)
tax.levels.tables.sums.ord.list <- vector("list",length(tax.levels)); names(tax.levels.tables.sums.ord.list) <- paste0("M.seq.tax.subset.",tax.levels,".sums.ord.") #tax.levels.tables.sums.ord <- paste0("M.seq.tax.subset.",tax.levels,".sums.ord")

tax.levels.tables.sums.ord.box.list <- vector("list",length(tax.levels)); names(tax.levels.tables.sums.ord.box.list) <- paste0("M.seq.tax.subset.",tax.levels,".sums.ord.box") #tax.levels.tables.sums.ord.box <- paste0("M.seq.tax.subset.",tax.levels,".sums.ord.box")
tax.levels.tables.sums.ord.box.rel.list <- vector("list",length(tax.levels)); names(tax.levels.tables.sums.ord.box.rel.list) <- paste0("M.seq.tax.subset.",tax.levels,".sums.ord.box.rel") #tax.levels.tables.sums.ord.box.rel <- paste0("M.seq.tax.subset.",tax.levels,".sums.ord.box.rel")

tax.levels.tables.sums.ord.rel.list <- vector("list",length(tax.levels)); names(tax.levels.tables.sums.ord.rel.list) <- paste0("M.seq.tax.subset.",tax.levels,".sums.ord.rel") #tax.levels.tables.sums.ord.rel <- paste0("M.seq.tax.subset.",tax.levels,".sums.ord.box.rel")
tax.levels.tables.sums.ord.pa.list <- vector("list",length(tax.levels)); names(tax.levels.tables.sums.ord.pa.list) <- paste0("M.seq.tax.subset.",tax.levels,".sums.ord.pa") #tax.levels.tables.sums.ord.pa <- paste0("M.seq.tax.subset.",tax.levels,".sums.ord.box.rel")
tax.levels.tables.sums.ord.rel.all.list <- vector("list",length(tax.levels)); names(tax.levels.tables.sums.ord.rel.all.list) <- paste0("M.seq.tax.subset.",tax.levels,".sums.ord.all.rel") #tax.levels.tables.sums.ord.rel <- paste0("M.seq.tax.subset.",tax.levels,".sums.ord.box.rel")

tax.levels.tables.asv.sums <- vector("list",length(tax.levels)); names(tax.levels.tables.asv.sums) <- paste0("M.seq.tax.subset.",tax.levels,".asv.sums")
tax.levels.tables.asv.sums.ord <- vector("list",length(tax.levels)); names(tax.levels.tables.asv.sums) <- paste0("M.seq.tax.subset.",tax.levels,".asv.sums.ord")
tax.levels.tables.asv.sums.ord.box.list <- vector("list",length(tax.levels)); names(tax.levels.tables.asv.sums) <- paste0("M.seq.tax.subset.",tax.levels,".asv.sums.ord.box")

tax.levels.tables.asv.sums.grouped <- vector("list",length(tax.levels)); names(tax.levels.tables.asv.sums) <- paste0("M.seq.tax.subset.",tax.levels,".asv.sums.grouped")
tax.levels.tables.asv.sums.grouped.ord <- vector("list",length(tax.levels)); names(tax.levels.tables.asv.sums) <- paste0("M.seq.tax.subset.",tax.levels,".asv.sums.ord.grouped")
tax.levels.tables.asv.sums.grouped.ord.box.list <- vector("list",length(tax.levels)); names(tax.levels.tables.asv.sums) <- paste0("M.seq.tax.subset.",tax.levels,".asv.sums.ord.box.grouped")


countNonZero <- function(x) {
  sum(x != 0)
}

# # create grouped M.seq.tax.subset.ord so we have a table which we can use to assign the number of different ASVs per selected grouping
M.seq.tax.subset.ord.grouped <- M.seq.tax.subset.ord[,c(1:(ncol(M.seq.tax.subset.ord)-ncol.taxo.noNA))] # remove taxonomy
M.seq.tax.subset.ord.grouped <- t(M.seq.tax.subset.ord.grouped) # now sample by ASVs table
gc()
M.seq.tax.subset.ord.grouped <- as.data.frame(M.seq.tax.subset.ord.grouped)
projects.sort.temp <- sort(M.projects)
M.seq.tax.subset.ord.grouped <- cbind(M.seq.tax.subset.ord.grouped,projects.sort.temp) # adds current grouping vector
rm(projects.sort.temp)
colnames(M.seq.tax.subset.ord.grouped)[ncol(M.seq.tax.subset.ord.grouped)] <- "Group"
M.seq.tax.subset.ord.grouped <- aggregate(M.seq.tax.subset.ord.grouped[1:(ncol(M.seq.tax.subset.ord.grouped)-1)],list(group=M.seq.tax.subset.ord.grouped[,ncol(M.seq.tax.subset.ord.grouped)]),FUN=countNonZero) # group by ASV table with number of samples the ASV was present in as fill
rownames(M.seq.tax.subset.ord.grouped) <- M.seq.tax.subset.ord.grouped[,1]
M.seq.tax.subset.ord.grouped <- M.seq.tax.subset.ord.grouped[,-1]

M.seq.tax.subset.ord.grouped <- t(M.seq.tax.subset.ord.grouped)
gc()
M.seq.tax.subset.ord.grouped <- as.data.frame(M.seq.tax.subset.ord.grouped)
M.seq.tax.subset.ord.grouped <- cbind(M.seq.tax.subset.ord.grouped,M.seq.tax.subset.ord[,c((ncol(M.seq.tax.subset.ord)-ncol.taxo.noNA+1):ncol(M.seq.tax.subset.ord))]) # add taxonomy again

for (i in 1:length(tax.levels)) {
  # # creates tables with only taxonomic information for one taxonomic level, to be analysed separately
  tax.levels.tables.list[[i]] <- M.seq.tax.subset.ord[,c(1:(ncol(M.seq.tax.subset.ord)-ncol.taxo.noNA),(ncol(M.seq.tax.subset.ord)-ncol.taxo.noNA+i))]
  tax.levels.tables.list.group[[i]] <- M.seq.tax.subset.ord.grouped[,c(1:(ncol(M.seq.tax.subset.ord.grouped)-ncol.taxo.noNA),(ncol(M.seq.tax.subset.ord.grouped)-ncol.taxo.noNA+i))]
  # # Taxonomic level tables with ASVs summed up within each clade
  # # as a side effect this orders the list alphabetically, this is still count data
  tax.levels.tables.sums.list[[i]] <- aggregate(tax.levels.tables.list[[i]][1:(ncol(tax.levels.tables.list[[i]])-1)],list(taxcur=tax.levels.tables.list[[i]][,ncol(tax.levels.tables.list[[i]])]),FUN=sum)
  # # calculate the number of different ASVs per clade
  tax.levels.tables.asv.sums[[i]] <- aggregate(tax.levels.tables.list[[i]][1:(ncol(tax.levels.tables.list[[i]])-1)],list(taxcur=tax.levels.tables.list[[i]][,ncol(tax.levels.tables.list[[i]])]),FUN=countNonZero)
  
  # # calculate the number of different ASVs per clade but group-wise
  tax.levels.tables.asv.sums.grouped[[i]] <- aggregate(tax.levels.tables.list.group[[i]][1:(ncol(tax.levels.tables.list.group[[i]])-1)],list(taxcur=tax.levels.tables.list.group[[i]][,ncol(tax.levels.tables.list.group[[i]])]),FUN=countNonZero)
  
  
  # tax.levels.tables.asv.sums.grouped.temp <- cbind(as.data.frame(t(tax.levels.tables.list[[i]][,-ncol(tax.levels.tables.list[[i]])])),sort(M.projects)) # attaches grouping vector to tavle of sample by ASV, filled with ASV richness
  # colnames(tax.levels.tables.asv.sums.grouped.temp)[ncol(tax.levels.tables.asv.sums.grouped.temp)] <- "Group"
  # tax.levels.tables.asv.sums.grouped.temp <- tax.levels.tables.asv.sums.grouped.temp %>% # calculates mean but basically we just want presence absence on ASV level per group
  #     dplyr::group_by(Group) %>%
  #     dplyr::summarise(across(where(is.numeric), mean, na.rm = TRUE))
  # 
  # 
  # # # bring it back to the original format
  # tax.levels.tables.asv.sums.grouped.temp <- as.data.frame(tax.levels.tables.asv.sums.grouped.temp)
  # rownames(tax.levels.tables.asv.sums.grouped.temp) <- tax.levels.tables.asv.sums.grouped.temp[,1]
  # tax.levels.tables.asv.sums.grouped.temp <- tax.levels.tables.asv.sums.grouped.temp[,-1]
  # tax.levels.tables.asv.sums.grouped.temp <- t(tax.levels.tables.asv.sums.grouped.temp)
  # tax.levels.tables.asv.sums.grouped.temp <- as.data.frame(tax.levels.tables.asv.sums.grouped.temp)
  # tax.levels.tables.asv.sums.grouped.temp <- cbind(tax.levels.tables.asv.sums.grouped.temp,tax.levels.tables.list[[i]][,ncol(tax.levels.tables.list[[i]])]) # add clade info again
  # colnames(tax.levels.tables.asv.sums.grouped.temp)[ncol(tax.levels.tables.asv.sums.grouped.temp)] <- colnames(tax.levels.tables.list[[i]][ncol(tax.levels.tables.list[[i]])])
  # # # calculate the number of different ASVs per clade, per grouping
  # tax.levels.tables.asv.sums.grouped[[i]] <- aggregate(tax.levels.tables.asv.sums.grouped.temp[1:(ncol(tax.levels.tables.asv.sums.grouped.temp)-1)],list(taxcur=tax.levels.tables.asv.sums.grouped.temp[,ncol(tax.levels.tables.asv.sums.grouped.temp)]),FUN=countNonZero)
  # rm(tax.levels.tables.asv.sums.grouped.temp)
  
  rownames(tax.levels.tables.sums.list[[i]]) <- tax.levels.tables.sums.list[[i]][,1] # assign rownames
  rownames(tax.levels.tables.asv.sums[[i]]) <- tax.levels.tables.asv.sums[[i]][,1] # assign rownames
  rownames(tax.levels.tables.asv.sums.grouped[[i]]) <- tax.levels.tables.asv.sums.grouped[[i]][,1] # assign rownames
  
  tax.levels.tables.sums.list[[i]] <- tax.levels.tables.sums.list[[i]][,-1] # remove first column, taxonomic level names are now the rownames, and first column deleted
  tax.levels.tables.asv.sums[[i]] <- tax.levels.tables.asv.sums[[i]][,-1] # remove first column, taxonomic level names are now the rownames, and first column deleted
  tax.levels.tables.asv.sums.grouped[[i]] <- tax.levels.tables.asv.sums.grouped[[i]][,-1] # remove first column, taxonomic level names are now the rownames, and first column deleted
  
  # # Create table for indicator species analysis
  if(CalculateIndicators=='Y'){
    tax.levels.tables.indicators.list[[i]]=t(tax.levels.tables.sums.list[[i]])
  }
  
  # # calculates relative abundance for ALL clades, add column with average and sort by descending average
  tax.levels.tables.sums.ord.rel.all.list[[i]] <- t(decostand(t(tax.levels.tables.sums.list[[i]]), method="total"))
  tax.levels.tables.sums.ord.rel.all.list[[i]] <- cbind(tax.levels.tables.sums.ord.rel.all.list[[i]],rowMeans(tax.levels.tables.sums.ord.rel.all.list[[i]]))
  colnames(tax.levels.tables.sums.ord.rel.all.list[[i]]) [ncol(tax.levels.tables.sums.ord.rel.all.list[[i]])]="Average"
  tax.levels.tables.sums.ord.rel.all.list[[i]] <- as.data.frame(tax.levels.tables.sums.ord.rel.all.list[[i]])
  tax.levels.tables.sums.ord.rel.all.list[[i]]=tax.levels.tables.sums.ord.rel.all.list[[i]][order(-tax.levels.tables.sums.ord.rel.all.list[[i]]$Average),] # Order based on decreasing average
  
  # # add column with average and sort by descending average to ASV sum table
  tax.levels.tables.asv.sums[[i]] <- cbind(tax.levels.tables.asv.sums[[i]],rowMeans(tax.levels.tables.asv.sums[[i]]))
  colnames(tax.levels.tables.asv.sums[[i]]) [ncol(tax.levels.tables.asv.sums[[i]])]="Average"
  tax.levels.tables.asv.sums[[i]] <- as.data.frame(tax.levels.tables.asv.sums[[i]])
  tax.levels.tables.asv.sums.ord[[i]]=tax.levels.tables.asv.sums[[i]][order(-tax.levels.tables.asv.sums[[i]]$Average),] # Order based on decreasing average
  
  tax.levels.tables.asv.sums.grouped[[i]] <- cbind(tax.levels.tables.asv.sums.grouped[[i]],rowMeans(tax.levels.tables.asv.sums.grouped[[i]]))
  colnames(tax.levels.tables.asv.sums.grouped[[i]]) [ncol(tax.levels.tables.asv.sums.grouped[[i]])]="Average"
  tax.levels.tables.asv.sums.grouped[[i]] <- as.data.frame(tax.levels.tables.asv.sums.grouped[[i]])
  tax.levels.tables.asv.sums.grouped.ord[[i]]=tax.levels.tables.asv.sums.grouped[[i]][order(-tax.levels.tables.asv.sums.grouped[[i]]$Average),] # Order based on decreasing average
  
  # # Add column with row average to counts data
  tax.levels.tables.sums.list[[i]] <- cbind(tax.levels.tables.sums.list[[i]],rowMeans(tax.levels.tables.sums.list[[i]]))
  colnames(tax.levels.tables.sums.list[[i]]) [ncol(tax.levels.tables.sums.list[[i]])]="Average"
  # # Sort table based on decreasing average, show top 20 taxa and combine other rare taxa to "Other"
  tax.levels.tables.sums.ord.list[[i]]=tax.levels.tables.sums.list[[i]][order(-tax.levels.tables.sums.list[[i]]$Average),] # Order based on decreasing average
  
  NRP=nrow(tax.levels.tables.sums.ord.list[[i]]) # Number of rows in table
    # # for counts table: Takes the first 'NumberOfTOPClades.box' rows; sums other rows with rare taxa
  if(NRP>NumberOfTOPClades.box) {
    tax.levels.tables.sums.ord.box.list[[i]]=rbind(tax.levels.tables.sums.ord.list[[i]][c(1:NumberOfTOPClades.box),],colSums(tax.levels.tables.sums.ord.list[[i]][c((NumberOfTOPClades.box+1):NRP),])); 
    rownames(tax.levels.tables.sums.ord.box.list[[i]]) [nrow(tax.levels.tables.sums.ord.box.list[[i]])]="Other" 
  } else {
    tax.levels.tables.sums.ord.box.list[[i]]=tax.levels.tables.sums.ord.list[[i]]
  }
    # # for ASV richness table: Takes the first 'NumberOfTOPClades.box' rows; sums other rows with rare taxa
  if(NRP>NumberOfTOPClades.box) {
    tax.levels.tables.asv.sums.ord.box.list[[i]] =rbind(tax.levels.tables.asv.sums.ord[[i]][c(1:NumberOfTOPClades.box),],colSums(tax.levels.tables.asv.sums.ord[[i]][c((NumberOfTOPClades.box+1):NRP),]));
    rownames(tax.levels.tables.asv.sums.ord.box.list[[i]]) [nrow(tax.levels.tables.asv.sums.ord.box.list[[i]])]="Other" 
  } else {
    tax.levels.tables.asv.sums.ord.box.list[[i]]=tax.levels.tables.asv.sums.ord[[i]]
  }
  # # for ASV richness table: Takes the first 'NumberOfTOPClades.box' rows; sums other rows with rare taxa
  if(NRP>NumberOfTOPClades.box) {
    tax.levels.tables.asv.sums.grouped.ord.box.list[[i]] =rbind(tax.levels.tables.asv.sums.grouped.ord[[i]][c(1:NumberOfTOPClades.box),],colSums(tax.levels.tables.asv.sums.grouped.ord[[i]][c((NumberOfTOPClades.box+1):NRP),]));
    rownames(tax.levels.tables.asv.sums.grouped.ord.box.list[[i]]) [nrow(tax.levels.tables.asv.sums.grouped.ord.box.list[[i]])]="Other" 
  } else {
    tax.levels.tables.asv.sums.grouped.ord.box.list[[i]]=tax.levels.tables.asv.sums.grouped.ord[[i]]
  }
  
  # # for rel abund table: Takes the first 'NumberOfTOPClades.box' rows; sums other rows with rare taxa
  if(NRP>NumberOfTOPClades.box) {
    tax.levels.tables.sums.ord.box.rel.list[[i]]=rbind(tax.levels.tables.sums.ord.rel.all.list[[i]][c(1:NumberOfTOPClades.box),],colSums(tax.levels.tables.sums.ord.rel.all.list[[i]][c((NumberOfTOPClades.box+1):NRP),])); 
    rownames(tax.levels.tables.sums.ord.box.rel.list[[i]]) [nrow(tax.levels.tables.sums.ord.box.rel.list[[i]])]="Other" 
  } else {
    tax.levels.tables.sums.ord.box.rel.list[[i]]=tax.levels.tables.sums.ord.rel.all.list[[i]]
  }
  
  # # for counts table:Takes the first 'NumberOfTOPClades' rows; sums other rows with rare taxa
  if(NRP>NumberOfTOPClades) {
    tax.levels.tables.sums.ord.list[[i]]=rbind(tax.levels.tables.sums.ord.list[[i]][c(1:NumberOfTOPClades),],colSums(tax.levels.tables.sums.ord.list[[i]][c((NumberOfTOPClades+1):NRP),])); 
    rownames(tax.levels.tables.sums.ord.list[[i]]) [nrow(tax.levels.tables.sums.ord.list[[i]])]="Other" 
  }
  
  if(NRP>NumberOfTOPClades) {
    tax.levels.tables.asv.sums.ord[[i]] =rbind(tax.levels.tables.asv.sums.ord[[i]][c(1:NumberOfTOPClades),],colSums(tax.levels.tables.asv.sums.ord[[i]][c((NumberOfTOPClades+1):NRP),]));
    rownames(tax.levels.tables.asv.sums.ord[[i]]) [nrow(tax.levels.tables.asv.sums.ord[[i]])]="Other" 
  }
  
  if(NRP>NumberOfTOPClades) {
    tax.levels.tables.asv.sums.grouped.ord[[i]] =rbind(tax.levels.tables.asv.sums.grouped.ord[[i]][c(1:NumberOfTOPClades),],colSums(tax.levels.tables.asv.sums.grouped.ord[[i]][c((NumberOfTOPClades+1):NRP),]));
    rownames(tax.levels.tables.asv.sums.grouped.ord[[i]]) [nrow(tax.levels.tables.asv.sums.grouped.ord[[i]])]="Other" 
  }
  
  if(NRP>NumberOfTOPClades) {
    tax.levels.tables.sums.ord.rel.list[[i]]=rbind(tax.levels.tables.sums.ord.rel.all.list[[i]][c(1:NumberOfTOPClades),],colSums(tax.levels.tables.sums.ord.rel.all.list[[i]][c((NumberOfTOPClades+1):NRP),])); 
    rownames(tax.levels.tables.sums.ord.rel.list[[i]]) [nrow(tax.levels.tables.sums.ord.rel.list[[i]])]="Other" 
  } else {
    tax.levels.tables.sums.ord.rel.list[[i]]=tax.levels.tables.sums.ord.rel.all.list[[i]]
  }
  
  # # Calculate presence/absence tables
  # tax.levels.tables.sums.ord.rel.list[[i]]=t(decostand(t(tax.levels.tables.sums.ord.list[[i]]), method="total"))
  tax.levels.tables.sums.ord.pa.list[[i]]=t(decostand(t(tax.levels.tables.sums.ord.list[[i]]), method="pa"))
  # # remove average column again and calculate actual average of presence absence
  tax.levels.tables.sums.ord.pa.list[[i]] <- tax.levels.tables.sums.ord.pa.list[[i]][,-ncol(tax.levels.tables.sums.ord.pa.list[[i]])]
  tax.levels.tables.sums.ord.pa.list[[i]] <- as.data.frame(tax.levels.tables.sums.ord.pa.list[[i]])
  tax.levels.tables.sums.ord.pa.list[[i]] <- cbind(tax.levels.tables.sums.ord.pa.list[[i]],apply(tax.levels.tables.sums.ord.pa.list[[i]][1:(ncol(tax.levels.tables.sums.ord.pa.list[[i]]))],1,mean))
  colnames(tax.levels.tables.sums.ord.pa.list[[i]])[ncol(tax.levels.tables.sums.ord.pa.list[[i]])] <- "Average"
  
  
  # # add standard deviation and print out
  a <- cbind(tax.levels.tables.sums.list[[i]],apply(tax.levels.tables.sums.list[[i]][1:(ncol(tax.levels.tables.sums.list[[i]])-1)],1,sd))
  colnames(a)[ncol(a)] <- "sd"
  data.table::fwrite(a,file=file.path(CompositionTablePath,paste(VisuaRProjectName,"_",tax.levels[i],"_abund.csv",sep="")),col.names = T,row.names = T)
  a <- cbind(tax.levels.tables.sums.ord.list[[i]],apply(tax.levels.tables.sums.ord.list[[i]][1:(ncol(tax.levels.tables.sums.ord.list[[i]])-1)],1,sd))
  colnames(a)[ncol(a)] <- "sd"
  data.table::fwrite(a,file=file.path(CompositionTablePath,paste(VisuaRProjectName,"_",tax.levels[i],"_abund_TOP",NumberOfTOPClades,".csv",sep="")),col.names = T,row.names = T)
  a <- cbind(tax.levels.tables.sums.ord.rel.all.list[[i]],apply(tax.levels.tables.sums.ord.rel.all.list[[i]][1:(ncol(tax.levels.tables.sums.ord.rel.all.list[[i]])-1)],1,sd))
  colnames(a)[ncol(a)] <- "sd"
  data.table::fwrite(as.data.frame(a),file=file.path(CompositionTablePath,paste(VisuaRProjectName,"_",tax.levels[i],"_relabund.csv",sep="")),col.names = T,row.names = T)
  a <- cbind(tax.levels.tables.sums.ord.rel.list[[i]],apply(tax.levels.tables.sums.ord.rel.list[[i]][1:(ncol(tax.levels.tables.sums.ord.rel.list[[i]])-1)],1,sd))
  colnames(a)[ncol(a)] <- "sd"
  data.table::fwrite(as.data.frame(a),file=file.path(CompositionTablePath,paste(VisuaRProjectName,"_",tax.levels[i],"_relabund_TOP",NumberOfTOPClades,".csv",sep="")),col.names = T,row.names = T)
  a <- cbind(tax.levels.tables.sums.ord.pa.list[[i]],apply(tax.levels.tables.sums.ord.pa.list[[i]][1:(ncol(tax.levels.tables.sums.ord.pa.list[[i]])-1)],1,sd))
  colnames(a)[ncol(a)] <- "sd"
  data.table::fwrite(as.data.frame(a),file=file.path(CompositionTablePath,paste(VisuaRProjectName,"_",tax.levels[i],"_abund_pa_TOP",NumberOfTOPClades,".csv",sep="")),col.names = T,row.names = T)
  
  if (SaveWholeworkspace=='N') {
    tax.levels.tables.sums.list[[i]]<- NULL
    tax.levels.tables.sums.ord.list[[i]] <- NULL
    tax.levels.tables.sums.ord.box.list[[i]] <- NULL
    tax.levels.tables.sums.ord.pa.list[[i]] <- NULL
    # tax.levels.tables.sums.ord.rel.all.list[[i]] <- NULL
  }
  gc()
}
# # from here only these lists will be used for the further analysis
# tax.levels.tables.sums.ord.rel.all.list
# tax.levels.tables.sums.ord.box.rel.list
# tax.levels.tables.sums.ord.rel.list


# #=== 3.4. Creates Bubble Plots and BoxPlots of relative sequence abundances for all taxonomic levels ==============================================================================================
CompositionPlotsPath=file.path(PathToVisuaRAnalysis,"Alpha_Diversity","CompositionPlots") 
dir.create(file.path(PathToVisuaRAnalysis,"Alpha_Diversity","CompositionPlots"))          # Creates Folder for composition plots

cat('\n\n3.4. Create Bubble and BoxPlots of relative abundances for all taxonomic levels',file=(file.path(PathToVisuaRAnalysis,paste(VisuaRProjectName,'_Analysis_log.txt',sep=''))),sep='',append=TRUE)
cat('\nThe plots will be saved to ',CompositionPlotsPath,'.\nFor each taxonomic level a seperated folder will be created inside this folder.',file=(file.path(PathToVisuaRAnalysis,paste(VisuaRProjectName,'_Analysis_log.txt',sep=''))),sep='',append=TRUE)
cat('\n1. The bubbleplot will show the samples on the x-axis (grouped and alphabetically sorted by ',Grouping1,' (Grouping1) and inside the groups sorted on increasing ',Grouping2,' (Grouping2)). The TOP',NumberOfTOPClades,' clades will be shown on the y axis. Relative Abundances will be presented as dot size.',file=(file.path(PathToVisuaRAnalysis,paste(VisuaRProjectName,'_Analysis_log.txt',sep=''))),sep='',append=TRUE)
cat('\n2. The boxplot BoxByClade_RA will show the TOP',NumberOfTOPClades.box,' clades on the x-axis and the relative abundance on the y axis. For each Clade ',length(M.projects.unique),' boxplots will be shown (Number of your Groups provided as Grouping1).',file=(file.path(PathToVisuaRAnalysis,paste(VisuaRProjectName,'_Analysis_log.txt',sep=''))),sep='',append=TRUE)
cat('\n3. The boxplot BoxByGroup_RA will show the ',length(M.projects.unique),' Groups provided as Grouping1 on the x axis and the relative abundance on the y axis. For each Clade 1 boxplot will be shown per Group.',file=(file.path(PathToVisuaRAnalysis,paste(VisuaRProjectName,'_Analysis_log.txt',sep=''))),sep='',append=TRUE)
cat('\n4. ',NumberOfTOPClades,' Boxplots will be produced of the TOP',NumberOfTOPClades ,' clades. Clade_1 is the Clade with the highest abundance, Clade_2 the one with the second highest abundance and so on.',file=(file.path(PathToVisuaRAnalysis,paste(VisuaRProjectName,'_Analysis_log.txt',sep=''))),sep='',append=TRUE)


tax.levels.tables.tv <- vector("list",length(tax.levels)); names(tax.levels.tables.tv) <- paste0("TV",tax.levels) #old list "levels with TVP,TVC etc. inside
tax.levels.tables.tv.box <- vector("list",length(tax.levels)); names(tax.levels.tables.tv.box) <-   paste0("TV",tax.levels,".box")# old list "levels.b" with TVP.b, TVC.b etc. inside
tax.levels.tables.tv.all <- vector("list",length(tax.levels)); names(tax.levels.tables.tv.all) <-   paste0("TV",tax.levels,".all")# old list "levels.b" with TVP.b, TVC.b etc. inside
tax.levels.tables.tv.asvs <- vector("list",length(tax.levels)); names(tax.levels.tables.tv.all) <-   paste0("TV",tax.levels,".asvs")# old list "levels.b" with TVP.b, TVC.b etc. inside
tax.levels.tables.tv.asvs.group <- vector("list",length(tax.levels)); names(tax.levels.tables.tv.all) <-   paste0("TV",tax.levels,".asvs.group")# old list "levels.b" with TVP.b, TVC.b etc. inside

# # removal of average in tables
for (i in 1:length(tax.levels)) {
  tax.levels.tables.tv[[i]] <- tax.levels.tables.sums.ord.rel.list[[i]][,c(1:(ncol(tax.levels.tables.sums.ord.rel.list[[i]])-1))] # takes a ASV x Sample Table (without the average). In this table the samples are ordered by the Groupings (1. Grouping1, 2. Grouping2)
  tax.levels.tables.tv.box[[i]] <- tax.levels.tables.sums.ord.box.rel.list[[i]][,c(1:(ncol(tax.levels.tables.sums.ord.box.rel.list[[i]])-1))] #takes a ASV x Sample Table, removes last column (average)
  tax.levels.tables.tv.all[[i]] <- tax.levels.tables.sums.ord.rel.all.list[[i]][,c(1:(ncol(tax.levels.tables.sums.ord.rel.all.list[[i]])-1))] #takes a ASV x Sample Table, removes last column (average)
  tax.levels.tables.tv.asvs[[i]] <- tax.levels.tables.asv.sums.ord[[i]][,c(1:(ncol(tax.levels.tables.asv.sums.ord[[i]])-1))] #takes a ASV x Sample Table, removes last column (average)
  tax.levels.tables.tv.asvs.group[[i]] <- tax.levels.tables.asv.sums.grouped.ord[[i]][,c(1:(ncol(tax.levels.tables.asv.sums.grouped.ord[[i]])-1))]
}

tax.levels <- as.character(tax.levels)

if (SaveWholeworkspace=='N') {rm(tax.levels.tables.sums.ord.rel.list,tax.levels.tables.sums.ord.box.rel.list,tax.levels.tables.sums.ord.rel.all.list)}

M.projects.ord=sort(M.projects) # sorts the color vector alphabetically to fit TVP, TVC etc.
M.colvec.ord=mapvalues(M.projects.ord, from = M.match.col[,1], to = M.match.col[,2]) # function in library (plyr), recodes sample vector, creates a color vector for the alphabetically by Grouping sorted data

# # # only taxonomic levels with more than 1 lineage will be visualized. 
for (i in length(tax.levels):1) {
  if (nrow(tax.levels.tables.tv[[i]])>1 & nrow(tax.levels.tables.tv.box[[i]])>1){
    startclade <- i
  } 
}

cat('\nAfter subsetting, the highest taxonomic level with more than one clade is ',tax.levels[startclade],'. Plots will only be created for this level and lower taxonomic levels.',file=(file.path(PathToVisuaRAnalysis,paste(VisuaRProjectName,'_Analysis_log.txt',sep=''))),sep='',append=TRUE)


if (SaveWholeworkspace=='N') {rm(tax.levels.tables.list)}

gc()

# # we are using the Bonferroni correction (rather conversative) by dividing the significance levels by the number of pairwise comparisons to obtain adjusted thresholds
numpairs<-length(M.projects.unique.ord)*(length(M.projects.unique.ord)-1)/2

wilcox.results <- as.data.frame(matrix(nrow=0,ncol=12))
# # calculation of wilcoxon ranked sums and output of statistics
for (k in startclade:(ncol.taxo.noNA)){ # goes through each taxonomix level
  TV=as.matrix(as.data.frame(tax.levels.tables.tv.all[k]))      # takes the respective relative abundance matrix for the bubble plots and the single-clade box plots (Clade by Sample)#
  if(nrow(TV)>1) {
    colnames(TV) <- colnames(tax.levels.tables.tv.all[[k]])
  }
  # number of rows = number of different clades
  if (NumberOfWilcoxTtestClades != "all") {
    # # Takes the first 'NumberOfTOPClades' rows; sums other rows with rare taxa
    if(nrow(TV)>NumberOfWilcoxTtestClades) {
      TV <- rbind(TV[c(1:NumberOfWilcoxTtestClades),],colSums(TV[c((NumberOfWilcoxTtestClades+1):nrow(TV)),]))
      rownames(TV) [nrow(TV)]="Other" 
    } 
  }
  
  nrow.TV=nrow(TV) 
  # # Creates group file for coloring of the plot.
  TV.col.group=c(rep(M.projects,nrow.TV))       # group file: fits to concatenated sample column, because the order of Groups (colnames) is multiplied by nrow.TV
  TV.col.group=sort(TV.col.group, decreasing=F) # sorts new group file alphabetically/increasing
  # # prepares relative sequence abundance tables for ggplot
  TV.levels=rownames(TV)                                                          # Level IDs (clade names) used for bubbleplot
  TVX <- melt(TV, id.vars = "Clade", variable.name="Sample", value.name = "Size") # turns multiple "Sample" columns (x axis) into one concatenated "Sample" column 
  colnames(TVX)=c("Clade","Sample","Size")
  # #---Creates boxplot of top clades. Main Group=Clade, SubGroup=Grouping1
  TVX=cbind(TVX,TV.col.group) # attaches the group column to the concatenated table
  # TVX$Size[is.na(TVX$Size)]=0
  if (length(M.projects.unique)>1){               # only if more than 1 group in Grouping1
    BoxClades=unique(TVX[,1])                     # list of unique clades in TVX
    for(f in seq_along(BoxClades)) {              # for-loop goes through every single clade
    # for(f in 1:20) {              # for-loop goes through every single clade
      BoxClade=as.character(BoxClades[f])         # this clade will be plotted
      TVX.single.box=TVX[grep(BoxClade,TVX[,1]),] # takes all relative abundance values belonging to the selected clade, adds a column with Grouping1, replaces NAs with 0
      TVX.single.box.groups=cbind(TVX.single.box,M.projects.ord)
      TVX.single.box.groups[is.na(TVX.single.box.groups)]=0
      TVX.single.box.groups.sums <- aggregate(TVX.single.box.groups$Size,by=list(Category=TVX.single.box.groups$M.projects.ord),FUN=sum)
      TVX.single.box.groups.avg <-aggregate(TVX.single.box.groups$Size,by=list(Category=TVX.single.box.groups$M.projects.ord),FUN=mean)
      TVX.single.box.groups.sd <-aggregate(TVX.single.box.groups$Size,by=list(Category=TVX.single.box.groups$M.projects.ord),FUN=sd)
      
      zero_x_categories <- TVX.single.box.groups.avg$Category[TVX.single.box.groups.avg$x ==0 |TVX.single.box.groups.avg$x ==1]
      # zero_x_categories <- TVX.single.box.groups.sums$Category[TVX.single.box.groups.sums$x == 0|TVX.single.box.groups.sums$x == 1]
      notzero_x_categories <- TVX.single.box.groups.avg$Category[TVX.single.box.groups.avg$x !=0 & TVX.single.box.groups.avg$x !=1]
      # notzero_x_categories <- TVX.single.box.groups.sums$Category[TVX.single.box.groups.sums$x != 0]
      
      if (length(zero_x_categories) <= 1) {
        # Perform pairwise Wilcoxon rank-sum tests
        test <- rstatix::pairwise_wilcox_test(
          data = TVX.single.box.groups,  # Your data frame
          formula = Size  ~ M.projects.ord,  # Adjust the formula to match your data
          p.adjust.method = "bonferroni"  # You can choose an appropriate adjustment method
        )
        test$Clade <- BoxClade
        test$group1avg <- NA
        test$group1sd <- NA
        test$group2avg <- NA
        test$group2sd <- NA
        group1avg.str <- NULL
        group1sd.str <- NULL
        for (numgroups in 1:(length(M.projects.unique)-1)) {
          group1avg.str <- append(group1avg.str,rep(TVX.single.box.groups.avg[numgroups,2],(length(M.projects.unique)-numgroups)))
          group1sd.str <- append(group1sd.str,rep(TVX.single.box.groups.sd[numgroups,2],(length(M.projects.unique)-numgroups)))
        }
        group2avg.str <- NULL
        group2sd.str <- NULL
        for (numgroups in 2:(length(M.projects.unique))) {
          group2avg.str <- append(group2avg.str,TVX.single.box.groups.avg[(numgroups:nrow(TVX.single.box.groups.avg)),2])
          group2sd.str <- append(group2sd.str,TVX.single.box.groups.sd[(numgroups:nrow(TVX.single.box.groups.sd)),2])
        }
        test$group1avg <- group1avg.str
        test$group1sd <- group1sd.str
        test$group2avg <- group2avg.str
        test$group2sd <- group2sd.str
        wilcox.results <- rbind(wilcox.results,test)
      } else if (length(zero_x_categories) > 1) {
        for (zerogroup in zero_x_categories) { # goes through all groups where we have a zero and only takes this group and the other groups where we dont have a zero
          # # to subset to one group where we only have zeroes and all non-zero groups
          subset_condition <- TVX.single.box.groups$M.projects.ord == zerogroup |
            TVX.single.box.groups$M.projects.ord %in% notzero_x_categories
          TVX.single.box.groups.sub <- TVX.single.box.groups[subset_condition,]
          
          TVX.single.box.groups.avg.sub <-aggregate(TVX.single.box.groups.sub$Size,by=list(Category=TVX.single.box.groups.sub$M.projects.ord),FUN=mean)
          TVX.single.box.groups.sd.sub <-aggregate(TVX.single.box.groups.sub$Size,by=list(Category=TVX.single.box.groups.sub$M.projects.ord),FUN=sd)
          
          M.projects.unique.temp <- length(unique(TVX.single.box.groups.sub$M.projects.ord))
          
          
          test <- rstatix::pairwise_wilcox_test(
            data = TVX.single.box.groups.sub,  # Your data frame
            formula = Size  ~ M.projects.ord,  # Adjust the formula to match your data
            p.adjust.method = "bonferroni"  # You can choose an appropriate adjustment method
          )
          test$Clade <- BoxClade
          test$group1avg <- NA
          test$group1sd <- NA
          test$group2avg <- NA
          test$group2sd <- NA
          group1avg.str <- NULL
          group1sd.str <- NULL
          
          for (numgroups in 1:((M.projects.unique.temp)-1)) {
            group1avg.str <- append(group1avg.str,rep(TVX.single.box.groups.avg.sub[numgroups,2],((M.projects.unique.temp)-numgroups)))
            group1sd.str <- append(group1sd.str,rep(TVX.single.box.groups.sd.sub[numgroups,2],((M.projects.unique.temp)-numgroups)))
          }
          group2avg.str <- NULL
          group2sd.str <- NULL
          for (numgroups in 2:((M.projects.unique.temp))) {
            group2avg.str <- append(group2avg.str,TVX.single.box.groups.avg.sub[(numgroups:nrow(TVX.single.box.groups.avg.sub)),2])
            group2sd.str <- append(group2sd.str,TVX.single.box.groups.sd.sub[(numgroups:nrow(TVX.single.box.groups.sd.sub)),2])
          }
          test$group1avg <- group1avg.str
          test$group1sd <- group1sd.str
          test$group2avg <- group2avg.str
          test$group2sd <- group2sd.str
          
          wilcox.results <- rbind(wilcox.results,test)
        }
        zero_x_combinations <- t(utils::combn(zero_x_categories, 2))
        zero_x_combinations_df <- as.data.frame(matrix(nrow=nrow(zero_x_combinations),ncol=1))
        zero_x_combinations_df <- cbind(zero_x_combinations_df,zero_x_combinations)
        zero_x_combinations_df$n1 <- 0
        zero_x_combinations_df$n2 <- 0
        zero_x_combinations_df$statistic <- NA
        zero_x_combinations_df$p <- NA
        zero_x_combinations_df$p.adj <- NA
        zero_x_combinations_df$p.adj.signif <- NA
        zero_x_combinations_df$Clade <- BoxClade
        zero_x_combinations_df$group1avg <- NA
        zero_x_combinations_df$group1sd <- NA
        zero_x_combinations_df$group2avg <- NA
        zero_x_combinations_df$group2sd <- NA
        
        zerogroup1avg <- NULL
        zerogroup1sd <- NULL
        for (zerocomp in (1:length(zero_x_categories)-1)) {
          zerogroup1avg <- append(zerogroup1avg, rep(TVX.single.box.groups.avg[which(TVX.single.box.groups.avg[,1]==zero_x_categories[zerocomp]),2],(length(zero_x_categories)-zerocomp)))
          zerogroup1sd <- append(zerogroup1sd, rep(TVX.single.box.groups.sd[which(TVX.single.box.groups.sd[,1]==zero_x_categories[zerocomp]),2],(length(zero_x_categories)-zerocomp)))
        }
        zerogroup2avg <- NULL
        zerogroup2sd <- NULL
        for (numgroups in 2:(length(zero_x_categories))) {
          zerogroup2avg <- append(zerogroup2avg,as.numeric(TVX.single.box.groups.avg[TVX.single.box.groups.avg$Category %in% zero_x_categories[numgroups:length(zero_x_categories)],2]))
          zerogroup2sd <- append(zerogroup2sd,as.numeric(TVX.single.box.groups.sd[TVX.single.box.groups.sd$Category %in% zero_x_categories[numgroups:length(zero_x_categories)],2]))
        }
        
        zero_x_combinations_df$group1avg <- zerogroup1avg
        zero_x_combinations_df$group1sd <- zerogroup1sd
        zero_x_combinations_df$group2avg <- zerogroup2avg
        zero_x_combinations_df$group2sd <- zerogroup2sd
        
        colnames(zero_x_combinations_df)[1:3] <- c(".y.","group1","group2")
        wilcox.results <- rbind(wilcox.results,zero_x_combinations_df)
      }
    }
    data.table::fwrite(wilcox.results,file=paste0(PathToVisuaRAnalysis,"/Alpha_Diversity/CompositionTables/",VisuaRProjectName,"_",tax.levels[k],"_wilcox_results.csv"),col.names = T,row.names = F)
    wilcox.results <- as.data.frame(matrix(nrow=0,ncol=12))
  }
}
rm(TV,nrow.TV,TV.col.group,TV.levels,TVX,wilcox.results, test,zero_x_categories,notzero_x_categories)

Sys.sleep(2) # this stops the execution for 10 seconds. If this is not set, the for loop is not always correctly executed

# # PLOT generation
for (k in startclade:(ncol.taxo.noNA)){ 
  # # Reads the matrix (1 for the bubble plots and 1 for the box plots)
  TV=as.matrix(as.data.frame(tax.levels.tables.tv[k]))      # takes the respective relative abundance matrix for the bubble plots and the single-clade box plots (Clade by Sample)
  colnames(TV) <- colnames(tax.levels.tables.tv[[k]])
  TV.b=as.matrix(as.data.frame(tax.levels.tables.tv.box[k]))  # takes the respective relative abundance matrix for the grouped box plots (Clade by Sample)
  colnames(TV.b) <- colnames(tax.levels.tables.tv.box[[k]])
  TV.asv <- as.matrix(as.data.frame(tax.levels.tables.tv.asvs[k]))  # takes the respective ASV richness matrix for the bubble plots and the single-clade box plots (Clade by Sample)
  colnames(TV.asv) <- colnames(tax.levels.tables.tv.asvs[[k]])
  TV.asv.group <- as.matrix(as.data.frame(tax.levels.tables.tv.asvs.group[k]))  # takes the respective ASV richness matrix for the bubble plots and the single-clade box plots (Clade by Sample)
  colnames(TV.asv.group) <- colnames(tax.levels.tables.tv.asvs.group[[k]])
  
  nrow.TV=nrow(TV)                            # number of rows = number of different clades
  nrow.TV.b=nrow(TV.b)                        # number of rows = number of different clades
  nrow.TV.asv <- nrow(TV.asv)
  nrow.TV.asv.group <- nrow(TV.asv.group)
  
  # # Creates group file for coloring of the plot.
  TV.col.group=c(rep(M.projects,nrow.TV))       # group file: fits to concatenated sample column, because the order of Groups (colnames) is multiplied by nrow.TV
  TV.col.group.b=c(rep(M.projects,nrow.TV.b))
  TV.col.group.asv <- c(rep(M.projects,nrow.TV.asv))
  
  TV.col.group=sort(TV.col.group, decreasing=F) # sorts new group file alphabetically/increasing
  TV.col.group.b=sort(TV.col.group.b,decreasing = F)
  TV.col.group.asv=sort(TV.col.group.asv,decreasing = F)
  
  TV[TV==0]=NA # turns zeros in relative abundance file into NAs, elsewise zeros are shown as dots in the bubble plot
  TV.b[TV.b==0]=NA
  TV.asv[TV.asv==0]=NA
  TV.asv.group[TV.asv.group==0]=NA
  
  # # prepares relative sequence abundance tables for ggplot
  TV.levels=rownames(TV)                                                          # Level IDs (clade names) used for bubbleplot
  TVX <- melt(TV, id.vars = "Clade", variable.name="Sample", value.name = "Size") # turns multiple "Sample" columns (x axis) into one concatenated "Sample" column 
  TVX.b=melt(TV.b,id.vars="Clade", variable.name="Sample", value.name = "Size")
  TVX.asv=melt(TV.asv,id.vars="Clade", variable.name="Sample", value.name = "Size")
  TVX.asv.group=melt(TV.asv.group,id.vars="Clade", variable.name="Sample", value.name = "Size")
  
  colnames(TVX)=c("Clade","Sample","Size")
  colnames(TVX.b)=c("Clade","Sample","Size")
  colnames(TVX.asv)=c("Clade","Sample","Size")
  colnames(TVX.asv.group)=c("Clade","Sample","Size")
  
  dir.create(file.path(PathToVisuaRAnalysis,"Alpha_Diversity","CompositionPlots",tax.levels[k])) 
  
  # #---Create bubble plots 
  # pdf(file.path(PathToVisuaRAnalysis,"Alpha_Diversity","CompositionPlots",tax.levels[k],paste(VisuaRProjectName,"_",tax.levels[k],"_Bubble_RA_area.pdf",sep="")),height=(1.8+(length(unique(TVX$Clade))*0.12)),width=((ncol(TV)*0.2)+((k+1)*1.7)),useDingbats=F) # useDingbats=F is needed to avoid font display problems with downstream graphics software
  pdf(file.path(PathToVisuaRAnalysis,"Alpha_Diversity","CompositionPlots",tax.levels[k],paste(VisuaRProjectName,"_",tax.levels[k],"_Bubble_RA_area.pdf",sep="")),height=(NumberOfTOPClades*0.2)+4,width=((ncol(TV)*0.2)+((k+1)*1.7)),useDingbats=F) # useDingbats=F is needed to avoid font display problems with downstream graphics software
  M.bubble=ggplot(TVX, aes(x = Sample, y = factor(Clade, levels=rev(TV.levels)), col=factor(TV.col.group))) +   # plots samples vs Clades using colors according to group file (TV.col group is ordered alphabetically/increasing))
    geom_point(aes(size = Size*100)) +                                                                              # adjust size of circles for visualization reasons
    scale_size() +
    scale_colour_manual(values=M.col) +                                                                             # the same colors for projects throughout workflow. M.col has as many colors as groups and the first color in M.col is assigned to the first group in alphabetical/increasing order
    labs(x="Samples", y="Clades", col=Grouping1,size="Relative\nSequence\nAbundance\n(in %)",title=paste("Relative Sequence Abundance, ",tax.levels[k]," level ","(",VisuaRProjectName,")",sep="")) + #axis labels
    theme_bw() +
    theme(axis.text.x = element_text(face="plain", color="Black", size=10, angle=90,hjust=1), 
          axis.text.y = element_text(face="plain", color="Black", size=10, angle=0),
          plot.title = element_text(size=8))
  print(M.bubble) # the plot needs to be printed if the script is run at once. Elsewise an empty plot will be created.
  dev.off()
  # same but instead of RA now with Richness (number of differente ASVs in the respective clade)
  pdf(file.path(PathToVisuaRAnalysis,"Alpha_Diversity","CompositionPlots",tax.levels[k],paste(VisuaRProjectName,"_",tax.levels[k],"_Bubble_ASV_area.pdf",sep="")),height=(NumberOfTOPClades*0.2)+4,width=((ncol(TV)*0.2)+((k+1)*1.7)),useDingbats=F) # useDingbats=F is needed to avoid font display problems with downstream graphics software
  M.bubble=ggplot(TVX.asv, aes(x = Sample, y = factor(Clade, levels=rev(TV.levels)), col=factor(TV.col.group))) +   # plots samples vs Clades using colors according to group file (TV.col group is ordered alphabetically/increasing))
    geom_point(aes(size = Size)) +                                                                              # adjust size of circles for visualization reasons
    scale_size() +
    scale_colour_manual(values=M.col) +                                                                             # the same colors for projects throughout workflow. M.col has as many colors as groups and the first color in M.col is assigned to the first group in alphabetical/increasing order
    labs(x="Samples", y="Clades", col=Grouping1,size="Number\nof unique\nASVs",title=paste("Number of unique ASVs, ",tax.levels[k]," level ","(",VisuaRProjectName,")",sep="")) + #axis labels
    theme_bw() +
    theme(axis.text.x = element_text(face="plain", color="Black", size=10, angle=90,hjust=1), 
          axis.text.y = element_text(face="plain", color="Black", size=10, angle=0),
          plot.title = element_text(size=8))
  print(M.bubble) # the plot needs to be printed if the script is run at once. Elsewise an empty plot will be created.
  dev.off()
  
  
  pdf(file.path(PathToVisuaRAnalysis,"Alpha_Diversity","CompositionPlots",tax.levels[k],paste(VisuaRProjectName,"_",tax.levels[k],"_stackedbar_RA.pdf",sep="")),height=(NumberOfTOPClades*0.6),width=((ncol(TV)*0.15)+((k+1)*1.7)),useDingbats=F) # useDingbats=F is needed to avoid font display problems with downstream graphics software
  M.stacked_bar <- ggplot(TVX, aes(x = Sample, y = Size, fill = Clade)) +
    geom_bar(stat = "identity", position = "stack") +
    labs(x = "Samples", y = "Relative Sequence Abundance (in %)", fill = "Clades",
         title = paste("Relative Sequence Abundance, ", tax.levels[k], " level ", "(", VisuaRProjectName, ")", sep = "")) +
    theme_bw() +
    theme(axis.text.x = element_text(face = "plain", color = "Black", size = 10, angle = 90, hjust = 1),
          axis.text.y = element_text(face = "plain", color = "Black", size = 10, angle = 0),
          plot.title = element_text(size = 8))
  
  print(M.stacked_bar)
  dev.off()
  
  pdf(file.path(PathToVisuaRAnalysis,"Alpha_Diversity","CompositionPlots",tax.levels[k],paste(VisuaRProjectName,"_",tax.levels[k],"_stackedbar_ASV.pdf",sep="")),height=(NumberOfTOPClades*0.6),width=((ncol(TV)*0.15)+((k+1)*1.7)),useDingbats=F) # useDingbats=F is needed to avoid font display problems with downstream graphics software
  M.stacked_bar.asv <- ggplot(TVX.asv, aes(x = Sample, y = Size, fill = Clade)) +
    geom_bar(stat = "identity", position = "stack") +
    labs(x = "Samples", y = "Number of unique ASVs", fill = "Clades",
         title = paste("Number of unique ASVs within each, ", tax.levels[k], " level ", "(", VisuaRProjectName, ")", sep = "")) +
    theme_bw() +
    theme(axis.text.x = element_text(face = "plain", color = "Black", size = 10, angle = 90, hjust = 1),
          axis.text.y = element_text(face = "plain", color = "Black", size = 10, angle = 0),
          plot.title = element_text(size = 8))
  
  print(M.stacked_bar.asv)
  dev.off()
  
  TVXbar <- TVX
  TVXbar[is.na(TVXbar)]=0 # change NAs back to zeroes for stacked barplot
  TVX.asv.bar <- TVX.asv
  TVX.asv.bar[is.na(TVX.asv.bar)]=0 # change NAs back to zeroes for stacked barplot
  TVX.asv.bar.group <- TVX.asv.group
  TVX.asv.bar.group[is.na(TVX.asv.bar.group)]=0 # change NAs back to zeroes for stacked barplot
  
  # pdf(file.path(PathToVisuaRAnalysis,"Alpha_Diversity","CompositionPlots",tax.levels[k],paste(VisuaRProjectName,"_",tax.levels[k],"_Bubble_RA_radius.pdf",sep="")),height=(1.8+(length(unique(TVX$Clade))*0.12)),width=((ncol(TV)*0.2)+((k+1)*1.7)),useDingbats=F) # useDingbats=F is needed to avoid font display problems with downstream graphics software
  pdf(file.path(PathToVisuaRAnalysis,"Alpha_Diversity","CompositionPlots",tax.levels[k],paste(VisuaRProjectName,"_",tax.levels[k],"_Bubble_RA_radius.pdf",sep="")),height=(NumberOfTOPClades*0.2)+4,width=((ncol(TV)*0.2)+((k+1)*1.7)),useDingbats=F) # useDingbats=F is needed to avoid font display problems with downstream graphics software
  M.bubble=ggplot(TVX, aes(x = Sample, y = factor(Clade, levels=rev(TV.levels)), col=factor(TV.col.group))) +   # plots samples vs Clades using colors according to group file (TV.col group is ordered alphabetically/increasing))
    geom_point(aes(size = Size*100)) +                                                                              # adjust size of circles for visualization reasons
    scale_radius()+
    scale_colour_manual(values=M.col) +                                                                             # the same colors for projects throughout workflow. M.col has as many colors as groups and the first color in M.col is assigned to the first group in alphabetical/increasing order
    labs(x="Samples", y="Clades", col=Grouping1,size="Relative\nSequence\nAbundance\n(in %)",title=paste("Relative Sequence Abundance, ",tax.levels[k]," level ","(",VisuaRProjectName,")",sep="")) + #axis labels
    theme_bw() +
    theme(axis.text.x = element_text(face="plain", color="Black", size=10, angle=90,hjust=1), 
          axis.text.y = element_text(face="plain", color="Black", size=10, angle=0),
          plot.title = element_text(size=8))
  print(M.bubble) # the plot needs to be printed if the script is run at once. Elsewise an empty plot will be created.
  dev.off()
  # # same for ASV richness per clade
  pdf(file.path(PathToVisuaRAnalysis,"Alpha_Diversity","CompositionPlots",tax.levels[k],paste(VisuaRProjectName,"_",tax.levels[k],"_Bubble_ASV_radius.pdf",sep="")),height=(NumberOfTOPClades*0.2)+4,width=((ncol(TV)*0.2)+((k+1)*1.7)),useDingbats=F) # useDingbats=F is needed to avoid font display problems with downstream graphics software
  M.bubble=ggplot(TVX.asv, aes(x = Sample, y = factor(Clade, levels=rev(TV.levels)), col=factor(TV.col.group))) +   # plots samples vs Clades using colors according to group file (TV.col group is ordered alphabetically/increasing))
    geom_point(aes(size = Size)) +                                                                              # adjust size of circles for visualization reasons
    scale_radius()+
    scale_colour_manual(values=M.col) +                                                                             # the same colors for projects throughout workflow. M.col has as many colors as groups and the first color in M.col is assigned to the first group in alphabetical/increasing order
    labs(x="Samples", y="Clades", col=Grouping1,size="Number\nof unique\nASVs",title=paste("Number of unique ASVs, ",tax.levels[k]," level ","(",VisuaRProjectName,")",sep="")) + #axis labels
    theme_bw() +
    theme(axis.text.x = element_text(face="plain", color="Black", size=10, angle=90,hjust=1), 
          axis.text.y = element_text(face="plain", color="Black", size=10, angle=0),
          plot.title = element_text(size=8))
  print(M.bubble) # the plot needs to be printed if the script is run at once. Elsewise an empty plot will be created.
  dev.off()
  
  # # create averages of relative abundances within each group and clade to plot it as stacked bar chart
  TVXg <- cbind(TVXbar,TV.col.group) # TVXbar has zeroes where appropriate (not NAs)
  TVXg.asv <- cbind(TVX.asv.bar,TV.col.group)
  TVXg <- TVXg %>%
    dplyr::group_by(Clade,TV.col.group) %>% 
    dplyr::summarize(Average.Size =mean(Size)) 
  TVXg.asv.mean <- TVXg.asv %>%
    dplyr::group_by(Clade,TV.col.group) %>% 
    dplyr::summarize(Average.Size =mean(Size)) 
  
  TVXg.asv.group <- TVX.asv.bar.group

  pdf(file.path(PathToVisuaRAnalysis,"Alpha_Diversity","CompositionPlots",tax.levels[k],paste(VisuaRProjectName,"_",tax.levels[k],"_stackedbar_grouped_RA.pdf",sep="")),height=(NumberOfTOPClades*0.4),width=(k*1.1+(0.5+(length(unique(factor(TV.col.group)))))),useDingbats=F) # useDingbats=F is needed to avoid font display problems with downstream graphics software
  M.stacked_bar <- ggplot(TVXg, aes(x = factor(TV.col.group), y = Average.Size, fill = Clade)) +
    geom_bar(stat = "identity", position = "stack") +
    labs(x = "Samples", y = "Relative Sequence Abundance (in %)", fill = "Clades",
         title = paste("Relative Sequence Abundance, ", tax.levels[k], " level ", "(", VisuaRProjectName, ")", sep = "")) +
    theme_bw() +
    theme(axis.text.x = element_text(face = "plain", color = "Black", size = 10, angle = 90, hjust = 1),
          axis.text.y = element_text(face = "plain", color = "Black", size = 10, angle = 0),
          plot.title = element_text(size = 8))

  print(M.stacked_bar)
  dev.off()
  
  pdf(file.path(PathToVisuaRAnalysis,"Alpha_Diversity","CompositionPlots",tax.levels[k],paste(VisuaRProjectName,"_",tax.levels[k],"_stackedbar_grouped_ASV_mean.pdf",sep="")),height=(NumberOfTOPClades*0.4),width=(k*1.1+(0.5+(length(unique(factor(TV.col.group)))))),useDingbats=F) # useDingbats=F is needed to avoid font display problems with downstream graphics software
  M.stacked_bar <- ggplot(TVXg.asv.mean, aes(x = factor(TV.col.group), y = Average.Size, fill = Clade)) +
    geom_bar(stat = "identity", position = "stack") +
    labs(x = "Samples", y = "Number of unique ASVs (averaged)", fill = "Clades",
         title = paste("Number of unique ASVs (averaged), ", tax.levels[k], " level ", "(", VisuaRProjectName, ")", sep = "")) +
    theme_bw() +
    theme(axis.text.x = element_text(face = "plain", color = "Black", size = 10, angle = 90, hjust = 1),
          axis.text.y = element_text(face = "plain", color = "Black", size = 10, angle = 0),
          plot.title = element_text(size = 8))
  
  print(M.stacked_bar)
  dev.off()
  
  pdf(file.path(PathToVisuaRAnalysis,"Alpha_Diversity","CompositionPlots",tax.levels[k],paste(VisuaRProjectName,"_",tax.levels[k],"_stackedbar_grouped_ASV_sum.pdf",sep="")),height=(NumberOfTOPClades*0.4),width=(k*1.1+(0.5+(length(unique(factor(TV.col.group)))))),useDingbats=F) # useDingbats=F is needed to avoid font display problems with downstream graphics software
  M.stacked_bar <- ggplot(TVXg.asv.group, aes(x = factor(Sample), y = Size, fill = Clade)) +
    geom_bar(stat = "identity", position = "stack") +
    labs(x = "Samples", y = "Number of unique ASVs (summed)", fill = "Clades",
         title = paste("Number of unique ASVs (summed), ", tax.levels[k], " level ", "(", VisuaRProjectName, ")", sep = "")) +
    theme_bw() +
    theme(axis.text.x = element_text(face = "plain", color = "Black", size = 10, angle = 90, hjust = 1),
          axis.text.y = element_text(face = "plain", color = "Black", size = 10, angle = 0),
          plot.title = element_text(size = 8))
  
  print(M.stacked_bar)
  dev.off()
  
  TVXg[TVXg==0] <- NA
  TVXg.asv.mean[TVXg.asv.mean==0] <- NA
  TVXg.asv.group[TVXg.asv.group==0] <- NA
  
  # #---Create grouped bubble plots 
  pdf(file.path(PathToVisuaRAnalysis,"Alpha_Diversity","CompositionPlots",tax.levels[k],paste(VisuaRProjectName,"_",tax.levels[k],"_Bubble_grouped_RA_area.pdf",sep="")),height=(1.8+(length(unique(TVX$Clade))*0.1)),width=(k*1.1+(2+(length(unique(factor(TV.col.group)))*1.2))),useDingbats=F) # useDingbats=F is needed to avoid font display problems with downstream graphics software
  M.bubbleg=ggplot(TVXg, aes(x = factor(TV.col.group), y = factor(Clade, levels=rev(TV.levels)), col=factor(TV.col.group))) +   # plots samples vs Clades using colors according to group file (TV.col group is ordered alphabetically/increasing))
    geom_point(aes(size = Average.Size*100)) +                                                                              # adjust size of circles for visualization reasons
    scale_colour_manual(values=M.col) +                                                                             # the same colors for projects throughout workflow. M.col has as many colors as groups and the first color in M.col is assigned to the first group in alphabetical/increasing order
    labs(x=Grouping1, y="Clades", col=Grouping1,size="Relative\nSequence\nAbundance\n(in %)",title=paste("Relative Sequence Abundance, ",tax.levels[k]," level ","(",VisuaRProjectName,")",sep="")) + #axis labels
    theme_bw() +
    theme(axis.text.x = element_text(face="plain", color="Black", size=10, angle=90,hjust=1), 
          axis.text.y = element_text(face="plain", color="Black", size=10, angle=0),
          plot.title = element_text(size=8),
          legend.key.size = ggplot2::unit(0.01,"cm"))
  print(M.bubbleg) 
  dev.off()
  # # same for ASV richness for scaling
  pdf(file.path(PathToVisuaRAnalysis,"Alpha_Diversity","CompositionPlots",tax.levels[k],paste(VisuaRProjectName,"_",tax.levels[k],"_Bubble_grouped_ASV_area_mean.pdf",sep="")),height=(1.8+(length(unique(TVX$Clade))*0.1)),width=(k*1.1+(2+(length(unique(factor(TV.col.group)))*1.2))),useDingbats=F) # useDingbats=F is needed to avoid font display problems with downstream graphics software
  M.bubbleg=ggplot(TVXg.asv.mean, aes(x = factor(TV.col.group), y = factor(Clade, levels=rev(TV.levels)), col=factor(TV.col.group))) +   # plots samples vs Clades using colors according to group file (TV.col group is ordered alphabetically/increasing))
    geom_point(aes(size = Average.Size)) +                                                                              # adjust size of circles for visualization reasons
    scale_colour_manual(values=M.col) +                                                                             # the same colors for projects throughout workflow. M.col has as many colors as groups and the first color in M.col is assigned to the first group in alphabetical/increasing order
    labs(x=Grouping1, y="Clades", col=Grouping1,size="Number\nof unique\nASVs\n(averaged)",title=paste("Number of unique ASVs averaged per biome, ",tax.levels[k]," level ","(",VisuaRProjectName,")",sep="")) + #axis labels
    theme_bw() +
    theme(axis.text.x = element_text(face="plain", color="Black", size=10, angle=90,hjust=1), 
          axis.text.y = element_text(face="plain", color="Black", size=10, angle=0),
          plot.title = element_text(size=8),
          legend.key.size = ggplot2::unit(0.01,"cm"))
  print(M.bubbleg) 
  dev.off()
  
  pdf(file.path(PathToVisuaRAnalysis,"Alpha_Diversity","CompositionPlots",tax.levels[k],paste(VisuaRProjectName,"_",tax.levels[k],"_Bubble_grouped_ASV_area_sum.pdf",sep="")),height=(1.8+(length(unique(TVX$Clade))*0.1)),width=(k*1.1+(2+(length(unique(factor(TV.col.group)))*1.2))),useDingbats=F) # useDingbats=F is needed to avoid font display problems with downstream graphics software
  M.bubbleg=ggplot(TVXg.asv.group, aes(x = factor(Sample), y = factor(Clade, levels=rev(TV.levels)), col=factor(Sample))) +   # plots samples vs Clades using colors according to group file (TV.col group is ordered alphabetically/increasing))
    geom_point(aes(size = Size)) +                                                                              # adjust size of circles for visualization reasons
    scale_colour_manual(values=M.col) +                                                                             # the same colors for projects throughout workflow. M.col has as many colors as groups and the first color in M.col is assigned to the first group in alphabetical/increasing order
    labs(x=Grouping1, y="Clades", col=Grouping1,size="Number\nof unique\nASVs\n(summed)",title=paste("Number of unique ASVs summed per biome, ",tax.levels[k]," level ","(",VisuaRProjectName,")",sep="")) + #axis labels
    theme_bw() +
    theme(axis.text.x = element_text(face="plain", color="Black", size=10, angle=90,hjust=1),
          axis.text.y = element_text(face="plain", color="Black", size=10, angle=0),
          plot.title = element_text(size=8),
          legend.key.size = ggplot2::unit(0.01,"cm"))
  print(M.bubbleg)
  dev.off()
  
  
  # 
  pdf(file.path(PathToVisuaRAnalysis,"Alpha_Diversity","CompositionPlots",tax.levels[k],paste(VisuaRProjectName,"_",tax.levels[k],"_Bubble_grouped_RA_radius.pdf",sep="")),height=(1.8+(length(unique(TVX$Clade))*0.1)),width=(k*1.1+(2+(length(unique(factor(TV.col.group)))*1.2))),useDingbats=F) # useDingbats=F is needed to avoid font display problems with downstream graphics software
  M.bubbleg=ggplot(TVXg, aes(x = factor(TV.col.group), y = factor(Clade, levels=rev(TV.levels)), col=factor(TV.col.group))) +   # plots samples vs Clades using colors according to group file (TV.col group is ordered alphabetically/increasing))
    geom_point(aes(size = Average.Size*100)) + 
    scale_radius() + # adjust size of circles for visualization reasons
    scale_colour_manual(values=M.col) +                                                                             # the same colors for projects throughout workflow. M.col has as many colors as groups and the first color in M.col is assigned to the first group in alphabetical/increasing order
    labs(x="Samples", y="Clades", col=Grouping1,size="Relative\nSequence\nAbundance\n(in %)",title=paste("Relative Sequence Abundance, ",tax.levels[k]," level ","(",VisuaRProjectName,")",sep="")) + #axis labels
    theme_bw() +
    theme(axis.text.x = element_text(face="plain", color="Black", size=10, angle=90,hjust=1), 
          axis.text.y = element_text(face="plain", color="Black", size=10, angle=0),
          plot.title = element_text(size=8),
          legend.key.size = ggplot2::unit(0.01,"cm"))
  print(M.bubbleg)
  dev.off()
  rm(TVXg,M.bubbleg)
  
  pdf(file.path(PathToVisuaRAnalysis,"Alpha_Diversity","CompositionPlots",tax.levels[k],paste(VisuaRProjectName,"_",tax.levels[k],"_Bubble_grouped_ASV_radius_mean.pdf",sep="")),height=(1.8+(length(unique(TVX$Clade))*0.1)),width=(k*1.1+(2+(length(unique(factor(TV.col.group)))*1.2))),useDingbats=F) # useDingbats=F is needed to avoid font display problems with downstream graphics software
  M.bubbleg=ggplot(TVXg.asv.mean, aes(x = factor(TV.col.group), y = factor(Clade, levels=rev(TV.levels)), col=factor(TV.col.group))) +   # plots samples vs Clades using colors according to group file (TV.col group is ordered alphabetically/increasing))
    geom_point(aes(size = Average.Size)) + 
    scale_radius() + # adjust size of circles for visualization reasons
    scale_colour_manual(values=M.col) +                                                                             # the same colors for projects throughout workflow. M.col has as many colors as groups and the first color in M.col is assigned to the first group in alphabetical/increasing order
    labs(x="Samples", y="Clades", col=Grouping1,size="Number\nof unique\nASVs\n(averaged)",title=paste("Number of unique ASVs averaged per biome, ",tax.levels[k]," level ","(",VisuaRProjectName,")",sep="")) + #axis labels
    theme_bw() +
    theme(axis.text.x = element_text(face="plain", color="Black", size=10, angle=90,hjust=1), 
          axis.text.y = element_text(face="plain", color="Black", size=10, angle=0),
          plot.title = element_text(size=8),
          legend.key.size = ggplot2::unit(0.01,"cm"))
  print(M.bubbleg)
  dev.off()
  rm(TVXg.asv.mean,M.bubbleg)
  
  pdf(file.path(PathToVisuaRAnalysis,"Alpha_Diversity","CompositionPlots",tax.levels[k],paste(VisuaRProjectName,"_",tax.levels[k],"_Bubble_grouped_ASV_radius_sum.pdf",sep="")),height=(1.8+(length(unique(TVX$Clade))*0.1)),width=(k*1.1+(2+(length(unique(factor(TV.col.group)))*1.2))),useDingbats=F) # useDingbats=F is needed to avoid font display problems with downstream graphics software
  M.bubbleg=ggplot(TVXg.asv.group, aes(x = factor(Sample), y = factor(Clade, levels=rev(TV.levels)), col=factor(Sample))) +   # plots samples vs Clades using colors according to group file (TV.col group is ordered alphabetically/increasing))
    geom_point(aes(size = Size)) +
    scale_radius() + # adjust size of circles for visualization reasons
    scale_colour_manual(values=M.col) +                                                                             # the same colors for projects throughout workflow. M.col has as many colors as groups and the first color in M.col is assigned to the first group in alphabetical/increasing order
    labs(x="Samples", y="Clades", col=Grouping1,size="Number\nof unique\nASVs\n(summed)",title=paste("Number of unique ASVs summed per biome, ",tax.levels[k]," level ","(",VisuaRProjectName,")",sep="")) + #axis labels
    theme_bw() +
    theme(axis.text.x = element_text(face="plain", color="Black", size=10, angle=90,hjust=1),
          axis.text.y = element_text(face="plain", color="Black", size=10, angle=0),
          plot.title = element_text(size=8),
          legend.key.size = ggplot2::unit(0.01,"cm"))
  print(M.bubbleg)
  dev.off()
  rm(TVXg.asv.sum,M.bubbleg)
  
  # #---Creates boxplot of top clades. Main Group=Clade, SubGroup=Grouping1
  TVX.box=cbind(TVX.b,TV.col.group.b) # attaches the group column to the concatenated table
  TVX.box$Size[is.na(TVX.box$Size)]=0
  pdf(file.path(PathToVisuaRAnalysis,"Alpha_Diversity","CompositionPlots",tax.levels[k],paste(VisuaRProjectName,"_",tax.levels[k],"_TOP",NumberOfTOPClades.box,"_BoxByClade_RA.pdf",sep="")),height=(4+(k*2)),width=(4.2+(length(unique(TVX$Clade))*0.1)),useDingbats=F) 
  M.box=ggplot(TVX.box,aes(x=Clade,y=Size*100,fill=factor(TV.col.group.b),colour="black"))+ # Transforms relative sequence Abundance to % values
    stat_boxplot(geom = "errorbar",position =position_dodge(preserve='single'))+            
    geom_boxplot(varwidth=F, notch=F, outlier.size=1,outlier.colour = "black",position=position_dodge(preserve = "single")) + 
    labs(x="Clades",y=paste("Relative Sequence Abundance [%]"),title=paste("Relative Sequence Abundance, ",tax.levels[k]," level ","(",VisuaRProjectName,")",sep="")) +
    scale_color_manual(values = "black",guide=FALSE) +
    scale_fill_manual(drop=F,values=M.col) + 
    theme(axis.text = element_text(colour="black"),
          legend.title = element_blank(),
          panel.grid.major.x = element_blank(),
          axis.text.x=element_text(angle=90,colour="black",size=14), #,vjust=-0.1
          axis.text.y=element_text(colour="black",size=14),
          axis.title = element_text(colour="black",size=16),
          axis.line=element_line(colour="black"),
          panel.background = element_blank(),
          panel.grid.major.y = element_line(colour="grey",size=0.5),
          panel.grid.minor.y = element_line(colour="grey",size = 0.25),
          panel.border = element_rect(fill=NA,colour = "black",size=0.5),
          legend.box.background = element_rect(colour="black"),
          legend.box.margin = margin(0.18,0.1,0.1,0.5),
          legend.key=element_rect(fill="white"),
          legend.text=element_text(size=14),
          axis.ticks=element_line(colour = "black"))
  print(M.box)
  dev.off()
  
  # #---Create boxplots of top clades Main Group=Grouping1, SubGroup=Clade, Colors:Rainbow
  pdf(file.path(PathToVisuaRAnalysis,"Alpha_Diversity","CompositionPlots",tax.levels[k],paste(VisuaRProjectName,"_",tax.levels[k],"_BoxByGroup_RA.pdf",sep="")),height=(6),width=(4.6+(length(unique(TVX$Clade))*0.2)+((k-1)*2.5)),useDingbats=F) #height=(8),width=(25)
  M.box.2=ggplot(TVX.box,aes(x=TV.col.group.b,y=Size*100,fill=(Clade),colour='black')) +
    stat_boxplot(geom = "errorbar",position =position_dodge(preserve='single')) + 
    geom_boxplot(varwidth=F, notch=F, outlier.size=1,position=position_dodge(preserve = "single")) + 
    labs(x=Grouping1,y=paste("Relative Sequence Abundance [%]"),title=paste("Relative Sequence Abundance, ",tax.levels[k]," level ","(",VisuaRProjectName,")",sep="")) +
    scale_x_discrete()+ # Orders the groups in the plot
    scale_color_manual(values = "black",guide=FALSE)+
    scale_fill_brewer(palette = 'Paired') + 
    guides(fill=guide_legend(ncol=1))+ 
    theme(axis.text = element_text(colour="black"),
          legend.title = element_blank(),
          panel.grid.major.x = element_blank(),
          axis.text.x=element_text(angle=0,colour="black",size=14),
          axis.text.y=element_text(colour="black",size=14),
          axis.title = element_text(colour="black",size=16),
          axis.line=element_line(colour="black"),
          panel.background = element_blank(),
          panel.grid.major.y = element_line(colour="grey",size=0.5),
          panel.grid.minor.y = element_line(colour="grey",size = 0.25),
          panel.border = element_rect(fill=NA,colour = "black",size=0.5),
          legend.box.background = element_rect(colour="black"),
          legend.box.margin = margin(0.18,0.1,0.1,0.5),
          legend.key=element_rect(fill="white"),
          legend.text=element_text(size=14),
          axis.ticks=element_line(colour = "black"))
  print(M.box.2)
  dev.off()
  
  # #---Makes boxplots of all Top Clades seperately
  if (length(M.projects.unique)>1){               # only if more than 1 group in Grouping1
    BoxClades=unique(TVX[,1])                     # list of unique clades in TVX
    for(f in seq_along(BoxClades)) {              # for-loop goes through every single clade
      BoxClade=as.character(BoxClades[f])         # this clade will be plotted
      TVX.single.box=TVX[grep(BoxClade,TVX[,1]),] # takes all relative abundance values belonging to the selected clade, adds a column with Grouping1, replaces NAs with 0
      TVX.single.box.groups=cbind(TVX.single.box,M.projects.ord)
      TVX.single.box.groups[is.na(TVX.single.box.groups)]=0
      pdf(file.path(PathToVisuaRAnalysis,"Alpha_Diversity","CompositionPlots",tax.levels[k],paste("Clade_",f,".pdf",sep="")),height = 4,width=length(M.projects.unique),useDingbats=F) # height=3,width=((2)*length(M.projects.unique))
      M.single.box=ggplot(TVX.single.box.groups, aes(x=M.projects.ord,y=Size*100,group=M.projects.ord))
      M.single.box= M.single.box +
        stat_boxplot(geom = "errorbar", width = 0.1) +                  
        geom_boxplot(varwidth=F, notch=F, fill=M.col,outlier.size=1) +  
        geom_signif(comparisons=combn(sort(as.character(unique(TVX.single.box.groups$M.projects.ord))),2,simplify=F),
                    step_increase=0.1,
                    size=0.3,
                    textsize=3,
                    tip_length=0.01,
                    map_signif_level=c("****"=0.0001,"***"=0.001,"**"=0.01,"*"=0.05," "=2))+
        stat_n_text(size=4)+
        labs(y="Relative Sequence Abundance [%]",title=BoxClade) +
        stat_summary(fun.y=mean,colour="black",geom="point",shape=17) +
        theme(axis.text = element_text(colour="black"),
              legend.title = element_blank(),
              panel.grid.major.x = element_blank(),
              axis.text.x=element_text(colour="black",size=10),
              axis.text.y=element_text(colour="black",size=12),
              # axis.title.x = element_text(colour="black",size=14),
              axis.title.x=element_blank(),
              axis.title.y = element_text(colour='black',size=12),
              plot.title = element_text(colour='black',size = 3),
              axis.line=element_line(colour="black"),
              panel.background = element_blank(),
              panel.grid.major.y = element_line(colour="grey",size=0.25),
              panel.grid.minor.y = element_blank(),
              panel.border = element_rect(fill=NA,colour = "black",size=0.5),
              legend.box.background = element_rect(colour="black"),
              legend.box.margin = margin(0.18,0.1,0.1,0.5),
              legend.key=element_rect(fill="white"),
              legend.text=element_text(size=12),
              axis.ticks=element_line(colour = "black"))
      print(M.single.box)
      dev.off()
      pdf(file.path(PathToVisuaRAnalysis,"Alpha_Diversity","CompositionPlots",tax.levels[k],paste("Clade_",f,"_cor.pdf",sep="")),height = 4,width=length(M.projects.unique),useDingbats=F) # height=3,width=((2)*length(M.projects.unique))
      M.single.box=ggplot(TVX.single.box.groups, aes(x=M.projects.ord,y=Size*100,group=M.projects.ord))
      M.single.box= M.single.box +
        stat_boxplot(geom = "errorbar", width = 0.1) +                  
        geom_boxplot(varwidth=F, notch=F, fill=M.col,outlier.size=1) +  
        geom_signif(comparisons=combn(sort(as.character(unique(TVX.single.box.groups$M.projects.ord))),2,simplify=F),
                    step_increase=0.1,
                    size=0.3,
                    textsize=3,
                    tip_length=0.01,
                    map_signif_level=c("****"=0.0001/numpairs,"***"=0.001/numpairs,"**"=0.01/numpairs,"*"=0.05/numpairs," "=2))+
        stat_n_text(size=4)+
        labs(y="Relative Sequence Abundance [%]",title=BoxClade) +
        stat_summary(fun.y=mean,colour="black",geom="point",shape=17) +
        theme(axis.text = element_text(colour="black"),
              legend.title = element_blank(),
              panel.grid.major.x = element_blank(),
              axis.text.x=element_text(colour="black",size=10),
              axis.text.y=element_text(colour="black",size=12),
              # axis.title.x = element_text(colour="black",size=14),
              axis.title.x=element_blank(),
              axis.title.y = element_text(colour='black',size=12),
              plot.title = element_text(colour='black',size = 3),
              axis.line=element_line(colour="black"),
              panel.background = element_blank(),
              panel.grid.major.y = element_line(colour="grey",size=0.25),
              panel.grid.minor.y = element_blank(),
              panel.border = element_rect(fill=NA,colour = "black",size=0.5),
              legend.box.background = element_rect(colour="black"),
              legend.box.margin = margin(0.18,0.1,0.1,0.5),
              legend.key=element_rect(fill="white"),
              legend.text=element_text(size=12),
              axis.ticks=element_line(colour = "black"))
      print(M.single.box)
      dev.off()
    }
    # # 202309: stopped using this. I think that is just a repetition of the one from above
    # if (length(M.projects.unique)>1){               # only if more than 1 group in Grouping1
    #   BoxClades=unique(TVX[,1])                     # list of unique clades in TVX
    #   for(f in seq_along(BoxClades)) {              # for-loop goes through every single clade
    #     BoxClade=as.character(BoxClades[f])         # this clade will be plotted
    #     TVX.single.box=TVX[grep(BoxClade,TVX[,1]),] # takes all relative abundance values belonging to the selected clade, adds a column with Grouping1, replaces NAs with 0
    #     TVX.single.box.groups=cbind(TVX.single.box,M.projects.ord)
    #     TVX.single.box.groups[is.na(TVX.single.box.groups)]=0
    #     pdf(file.path(PathToVisuaRAnalysis,"Alpha_Diversity","CompositionPlots",tax.levels[k],paste("Clade_",f,".pdf",sep="")),height = 4,width=length(M.projects.unique),useDingbats=F) # height=3,width=((2)*length(M.projects.unique))
    #     M.single.box=ggplot(TVX.single.box.groups, aes(x=M.projects.ord,y=Size*100,group=M.projects.ord))
    #     M.single.box= M.single.box +
    #       stat_boxplot(geom = "errorbar", width = 0.1) +                  
    #       geom_boxplot(varwidth=F, notch=F, fill=M.col,outlier.size=1) +  
    #       geom_signif(comparisons=combn(sort(as.character(unique(TVX.single.box.groups$M.projects.ord))),2,simplify=F),
    #                   step_increase=0.1,
    #                   size=0.3,
    #                   textsize=3,
    #                   tip_length=0.01,
    #                   map_signif_level=c("***"=0.001,"**"=0.01,"*"=0.05," "=2))+
    #       stat_n_text(size=4)+
    #       labs(y="Relative Sequence Abundance [%]",title=BoxClade) +
    #       stat_summary(fun.y=mean,colour="black",geom="point",shape=17) +
    #       theme(axis.text = element_text(colour="black"),
    #             legend.title = element_blank(),
    #             panel.grid.major.x = element_blank(),
    #             axis.text.x=element_text(colour="black",size=10),
    #             axis.text.y=element_text(colour="black",size=12),
    #             # axis.title.x = element_text(colour="black",size=14),
    #             axis.title.x=element_blank(),
    #             axis.title.y = element_text(colour='black',size=12),
    #             plot.title = element_text(colour='black',size = 3),
    #             axis.line=element_line(colour="black"),
    #             panel.background = element_blank(),
    #             panel.grid.major.y = element_line(colour="grey",size=0.25),
    #             panel.grid.minor.y = element_blank(),
    #             panel.border = element_rect(fill=NA,colour = "black",size=0.5),
    #             legend.box.background = element_rect(colour="black"),
    #             legend.box.margin = margin(0.18,0.1,0.1,0.5),
    #             legend.key=element_rect(fill="white"),
    #             legend.text=element_text(size=12),
    #             axis.ticks=element_line(colour = "black"))
    #     print(M.single.box)
    #     dev.off()
    #     # # bonferroni corrected significances
    #     pdf(file.path(PathToVisuaRAnalysis,"Alpha_Diversity","CompositionPlots",tax.levels[k],paste("Clade_",f,"_cor.pdf",sep="")),height = 4,width=length(M.projects.unique),useDingbats=F) # height=3,width=((2)*length(M.projects.unique))
    #     M.single.box=ggplot(TVX.single.box.groups, aes(x=M.projects.ord,y=Size*100,group=M.projects.ord))
    #     M.single.box= M.single.box +
    #       stat_boxplot(geom = "errorbar", width = 0.1) +                  
    #       geom_boxplot(varwidth=F, notch=F, fill=M.col,outlier.size=1) +  
    #       geom_signif(comparisons=combn(sort(as.character(unique(TVX.single.box.groups$M.projects.ord))),2,simplify=F),
    #                   step_increase=0.1,
    #                   size=0.3,
    #                   textsize=3,
    #                   tip_length=0.01,
    #                   map_signif_level=c("***"=0.001/numpairs,"**"=0.01/numpairs,"*"=0.05/numpairs," "=2))+
    #       stat_n_text(size=4)+
    #       labs(y="Relative Sequence Abundance [%]",title=BoxClade) +
    #       stat_summary(fun.y=mean,colour="black",geom="point",shape=17) +
    #       theme(axis.text = element_text(colour="black"),
    #             legend.title = element_blank(),
    #             panel.grid.major.x = element_blank(),
    #             axis.text.x=element_text(colour="black",size=10),
    #             axis.text.y=element_text(colour="black",size=12),
    #             # axis.title.x = element_text(colour="black",size=14),
    #             axis.title.x=element_blank(),
    #             axis.title.y = element_text(colour='black',size=12),
    #             plot.title = element_text(colour='black',size = 3),
    #             axis.line=element_line(colour="black"),
    #             panel.background = element_blank(),
    #             panel.grid.major.y = element_line(colour="grey",size=0.25),
    #             panel.grid.minor.y = element_blank(),
    #             panel.border = element_rect(fill=NA,colour = "black",size=0.5),
    #             legend.box.background = element_rect(colour="black"),
    #             legend.box.margin = margin(0.18,0.1,0.1,0.5),
    #             legend.key=element_rect(fill="white"),
    #             legend.text=element_text(size=12),
    #             axis.ticks=element_line(colour = "black"))
    #     print(M.single.box)
    #     dev.off()
    #   }
    # }
  }
}

if (SaveWholeworkspace=='N') {rm(tax.levels.tables.tv,tax.levels.tables.tv.box,TVX,TVX.b,TVX.box,TVX.single.box,TVX.single.box.groups,M.box.2,M.box)}

gc()
save.image(file.path(PathToVisuaRAnalysis,paste(VisuaRProjectName,'_VisuaR','.RData',sep='')))

closeAllConnections() # closes all currently open connections.

# #================== 4. Alpha Diversity ================================================================================================================================================
cat('\n\n4. Alpha Diversity',file=(file.path(PathToVisuaRAnalysis,paste(VisuaRProjectName,'_Analysis_log.txt',sep=''))),sep='',append=TRUE)

M=t(M.seq.tax.subset.ord.ASV)       # Sample by ASV table ordered by taxonomy. Not ordered by Grouping1.

if (SaveWholeworkspace=='N') {rm(M.seq.tax.subset.ord.ASV)}
gc()

data.table::fwrite(as.data.frame(M),file=file.path(PathToVisuaRAnalysis,'Beta_Diversity',paste(VisuaRProjectName,"_ASVbySample_abund.csv",sep="")),col.names = T,row.names = T)

M.rel=decostand(M, method="total")  # Calculates relative sequence abundances
M.pa=decostand(M, method="pa")      # Calculates presence/absence

M.sobs=rowSums(M.pa)  # Number of different observed ASVs per sample
M.ASVs=rowSums(M)     # Number of reads per sample, not ordered based on Grouping1

gc()
# #=======  4.1. Calculates diversity indices ============================================================================================================================================
dir.create(file.path(PathToVisuaRAnalysis,"Alpha_Diversity","Diversity_Metrics"))

cat('\n4.1. Calculates diversity indices',file=(file.path(PathToVisuaRAnalysis,paste(VisuaRProjectName,'_Analysis_log.txt',sep=''))),sep='',append=TRUE)

M.minreads=min(M.ASVs) # minimum number of reads in any given sample. Will be used to subsample in alpha diversity indices calculation. This allows comparison of samples of different sequencing depth.

cat('\nThe alpha diversity indices will be calculated with a subsample of ',M.minreads,' reads (minimum number of reads occuring in any sample).',file=(file.path(PathToVisuaRAnalysis,paste(VisuaRProjectName,'_Analysis_log.txt',sep=''))),sep='',append=TRUE)
cat('\nObserved ASV richness, Chao1 richness estimate, Shannon Entropy, Inverse Simpson Diversity, absolute and relative Singletons can be found in the file ',file.path(PathToVisuaRAnalysis,"Alpha_Diversity","Diversity_Metrics",paste(VisuaRProjectName,"_DiversityIndices_",Grouping1,".txt",sep="")),'.',file=(file.path(PathToVisuaRAnalysis,paste(VisuaRProjectName,'_Analysis_log.txt',sep=''))),sep='',append=TRUE)
cat('\nNote: diversity indices may be based on iterative subsampling, whereas SSOabs and SSOrel are based on the whole dataset.\nThis can result in the odd case that there seem to be more observed relative ASVs than observed ASVs.',file=(file.path(PathToVisuaRAnalysis,paste(VisuaRProjectName,'_Analysis_log.txt',sep=''))),sep='',append=TRUE)

NR=nrow(M) # number of rows of matrix M = number of Samples

# #--- 4.1.1. Observed ASV richness subsampled -------------------------------------------------------------------------------------------------------------------------------------------

M.sobs.r=matrix(NA,nrow=NR,ncol=NI)                                                   # matrix to be filled by the function

for (j in 1:NR) {                                                                     # calculates iterated observed ASV richness, subsampled using M.minreads
  for (i in 1:NI) {
    M.sobs.r[j,i]=estimateR(rrarefy(M[j,],sample=M.minreads),replace=F)[1,1]          # replace=T, if sample-parameter is larger than the smallest sample (not recommended)
  } 
}

M.sobs.r.mean=rowMeans(M.sobs.r)                                                      # calculation of average observed ASV richness from (NI reps)

if (SaveWholeworkspace=='N') {rm(M.sobs.r)}
gc()

# #--- 4.1.2. Chao1 Richness Estimate ----------------------------------------------------------------------------------------------------------------------------------------------------

M.chao1.r=matrix(NA, nrow=NR,ncol=NI)                                                 # matrix to be filled by the function

for (j in 1:NR) {                                                                     # calculates iterated Chao1 richness estimate, subsampled using M.minreads
  for (i in 1:NI) {
    M.chao1.r[j,i]=estimateR(rrarefy(M[j,],sample=M.minreads),replace=F)[2,1] 
  }
}

M.chao1.r.mean=rowMeans(M.chao1.r)                                                    #calculation of average Chao richness estimate from (NI reps)

# #--- 4.1.3. Shannon Entropy ------------------------------------------------------------------------------------------------------------------------------------------------------------

M.shan.r=matrix(NA,nrow=NR,ncol=NI)                                                   # matrix to be filled by the function

for (j in 1:NR) {                                                                     # calculate iterated Shannon entropy index, subsampled using M.minreads
  for (i in 1:NI) {                                                                   # happens once for each iteration
    M.shan.r[j,i]=diversity(rrarefy(M[j,],sample=M.minreads), index="shannon") 
  }
}      

M.shan.r.mean=rowMeans(M.shan.r)                                                      # calculation of average Shannon Entropy from (NI reps)

if (SaveWholeworkspace=='N') {rm(M.shan.r)}
gc()

# #--- 4.1.4. Inverse Simpson Diversity --------------------------------------------------------------------------------------------------------------------------------------------------

M.invs.r=matrix(NA,nrow=NR,ncol=NI)                                                   # matrix to be filled by the function

for (j in 1:NR) {                                                                     # calculates iterated inverse simpson diversity index, subsampled using M.minreads
  RE=rep(NA,NI) 
  for (i in 1:NI) {
    RE[i]=diversity(rrarefy(M[j,],sample=M.minreads), index="invsimpson") 
  }
  M.invs.r[j,]= RE;
}     

M.invs.r.mean=rowMeans(M.invs.r)                                                      # calculation of average Inverse Simpson Diversity from (NI reps)

if (SaveWholeworkspace=='N') {rm(M.invs.r)}
gc()

# #--- 4.1.5. Number of Absolute Singletons ----------------------------------------------------------------------------------------------------------------------------------------------

M.SSASV.abs=as.matrix(M[,apply(M,2,sum)==1])                                          # Table of only absolute singletons, 2 for columns

if (ncol(M.SSASV.abs)==0) {                                                           # only happens if data set has no absolute Single Sequence ASV (SSASVabs), i.e. absolute singletons
  M.SSASV.abs.nr=rep(0,NR); names(M.SSASV.abs.nr)=names(M.sobs)                       # creates a string of 0's (length=NR=number of samples)
  M.SSASV.abs.pc=rep(0,NR); names(M.SSASV.abs.pc)=names(M.sobs)
  M.SSASV.abs.nr.t=t(M.SSASV.abs.nr); row.names(M.SSASV.abs.nr.t)='M.SSASV.abs.nr'    # transposed to fit combined table below
  M.SSASV.abs.pc.t=t(M.SSASV.abs.pc); row.names(M.SSASV.abs.pc.t)="M.SSASV.abs.pc"    # transposed to fit combined table below
  if (SaveWholeworkspace=='N') {rm(M.SSASV.abs,M.SSASV.abs.nr,M.SSASV.abs.pc)}
  gc()
} else if (ncol(M.SSASV.abs)==1) {
  M.SSASV.abs.name=which(colSums(M)==1)                                               # finds name and position of absolute singleton in M
  colnames(M.SSASV.abs)=as.character(names(M.SSASV.abs.name))                         # renames the ASV to actual ASV name
  M.SSASV.abs.nr=rowSums(M.SSASV.abs)                                                 # Number of SSASVabs per sample
  M.SSASV.abs.pc=round(M.SSASV.abs.nr/M.sobs*100,1)                                   # % SSASVabs
  M.SSASV.abs.nr.t=t(M.SSASV.abs.nr); row.names(M.SSASV.abs.nr.t)="M.SSASV.abs.nr"    # transposed to fit combined table below
  M.SSASV.abs.pc.t=t(M.SSASV.abs.pc);row.names(M.SSASV.abs.pc.t)="M.SSASV.abs.pc"     # transposed to fit combined table below
  if (SaveWholeworkspace=='N') {rm(M.SSASV.abs,M.SSASV.abs.nr,M.SSASV.abs.name,M.SSASV.abs.pc)}
  gc()
} else if (ncol(M.SSASV.abs>1)) {
  M.SSASV.abs.nr=rowSums(M.SSASV.abs)                                                 # Number of SSASVabs per sample
  M.SSASV.abs.pc=round(M.SSASV.abs.nr/M.sobs*100,1)                                   # % SSASVabs
  M.SSASV.abs.nr.t=t(M.SSASV.abs.nr); row.names(M.SSASV.abs.nr.t)="M.SSASV.abs.nr"    # transposed to fit combined table below
  M.SSASV.abs.pc.t=t(M.SSASV.abs.pc);row.names(M.SSASV.abs.pc.t)="M.SSASV.abs.pc"     # transposed to fit combined table below
  if (SaveWholeworkspace=='N') {rm(M.SSASV.abs,M.SSASV.abs.nr,M.SSASV.abs.pc)}
  gc()
}

# #--- 4.1.6. Number of Relative Singletons ----------------------------------------------------------------------------------------------------------------------------------------------

M.no.SSASVs.abs=M[,-(which(apply(M,2,sum)==1))]                                                       # removes absolute singletons
M.SSASVs.rel=as.matrix(M.no.SSASVs.abs[,which(apply(M.no.SSASVs.abs,2,function(x) any(x==1))==TRUE)]) # calculates relative Single Sequence ASVs (SSASVrel), i.e. relative singletons. (ASVs that occur once in one sample and at least once in at least one other sample)

if (ncol(M.SSASVs.rel)==0){                                                       # if dataset has no SSASVrel
  M.SSASVs.rel.nr=rep(0,NR); names(M.SSASVs.rel.nr)=names(M.sobs)
  M.SSASVs.rel.pc=rep(0,NR); names(M.SSASVs.rel.pc)=names(M.sobs)
  M.SSASVs.rel.nr.t=t(M.SSASVs.rel.nr); row.names(M.SSASVs.rel.nr.t)="M.SSASVs.rel.nr"  # transposed to fit combined table below
  M.SSASVs.rel.pc.t=t(M.SSASVs.rel.pc); row.names(M.SSASVs.rel.pc.t)="M.SSASVs.rel.pc"  # transposed to fit combined table below
  if (SaveWholeworkspace=='N') {rm(M.SSASVs.rel.nr,M.SSASVs.rel.pc)}
  gc()
} else if (ncol(M.SSASVs.rel)==1) {                                                     # if dataset has exactly 1 SSASVrel
  M.SSASVs.rel.name=apply(M.no.SSASVs.abs, 2, function(x) any(x==1)==TRUE)              # Gives the names of all ASVs. TRUE or FALSE indicate if they are singeltons or not.
  M.SSASVs.rel.name=which(M.SSASVs.rel.name, arr.ind = FALSE, useNames = TRUE)          # only keeps SSASVrel
  colnames(M.SSASVs.rel)=as.character(names(M.SSASVs.rel.name))                         # renames the ASVs
  M.SSASVs.rel.nr = matrix(NA, nrow=NR, ncol=1)                                         # matrix to be filled by the function, NR:number of samples
  for (j in 1:NR) {                                                                     # counts the number of SSASVrel per sample and fills the matrix
    M.SSASVs.rel.nr[j,]= sum(M.SSASVs.rel[j,]==1)
  }  
  M.SSASVs.rel.pc=round(M.SSASVs.rel.nr/M.sobs*100,1)                                   # % SSASVrel
  M.SSASVs.rel.nr.t=t(M.SSASVs.rel.nr); row.names(M.SSASVs.rel.nr.t)="M.SSASVs.rel.nr"  
  M.SSASVs.rel.pc.t=t(M.SSASVs.rel.pc); row.names(M.SSASVs.rel.pc.t)="M.SSASVs.rel.pc"  
  if (SaveWholeworkspace=='N') {rm(M.SSASVs.rel.nr,M.SSASVs.rel.pc,M.SSASVs.rel.name)}
  gc()
} else if (ncol(M.SSASVs.rel)>1) {                                                      # if dataset has more than 1 SSASVrel
  M.SSASVs.rel.nr = matrix(NA, nrow=NR, ncol=1)                                         
  for (j in 1:NR) {                                                                    
    M.SSASVs.rel.nr[j,]= sum(M.SSASVs.rel[j,]==1)                                       
  }    
  M.SSASVs.rel.pc=round(M.SSASVs.rel.nr/M.sobs*100,1)                                   
  M.SSASVs.rel.nr.t=t(M.SSASVs.rel.nr); row.names(M.SSASVs.rel.nr.t)="M.SSASVs.rel.nr"  
  M.SSASVs.rel.pc.t=t(M.SSASVs.rel.pc); row.names(M.SSASVs.rel.pc.t)="M.SSASVs.rel.pc"  
  if (SaveWholeworkspace=='N') {rm(M.SSASVs.rel.nr,M.SSASVs.rel.pc)}
  gc()
}

# #--- 4.1.7. Combines diversity indices in one table ---------------------------------------------------------------------------------------------------------------------------------

# # creates table of averaged indices, rounds and reformats table
M.diversity=rbind(M.ASVs, #1
                  M.sobs, #2
                  M.sobs.r.mean, #3 
                  M.chao1.r.mean, #4
                  M.invs.r.mean, #5
                  M.SSASV.abs.nr.t, #6
                  M.SSASVs.rel.nr.t, #7
                  M.shan.r.mean, #8
                  M.SSASV.abs.pc.t, #9 
                  M.SSASVs.rel.pc.t)  # 10 comes from M.seq.tax.subset.ord.ASV; is not ordered based on Grouping1. 
# # M.ASVs = Total reads (unsubsampled)
# # M.sobs = Observed ASVs (unsubsampled)
# # M.sobs.r.mean = Observed ASVs - Richness (subsampled) 
# # M.chao1.r.mean = Estimated ASVs - Chao1 (subsampled)
# # M.invs.r.mean = Inverse Simpson Diversity (subsampled)
# # M.SSASV.abs.nr.t = Absolute Single Sequence ASVs (unsubsampled)
# # M.SSASVs.rel.nr.t = Relative Single Sequence ASVs (unsubsampled)
# # M.shan.r.mean = Shannon Entropy (subsampled)
# # M.SSASV.abs.pc.t = Absolute Single Sequence ASVs in % (unsubsampled)
# # M.SSASVs.rel.pc.t = Relative Single Sequence ASVs in % (unsubsampled)
# # M.minreads = Total Reads (subsampled)

M.diversity.reord=t(rbind(M.ASVs, # total reads unsubsampled
                          M.sobs, # observed asvs unsubsampled
                          # M.sobs, # observed ASVs
                          M.SSASV.abs.nr.t,
                          M.SSASV.abs.pc.t,
                          M.SSASVs.rel.nr.t,
                          M.SSASVs.rel.pc.t,
                          rep(M.minreads,ncol(M.diversity)),
                          M.sobs.r.mean,
                          M.chao1.r.mean,
                          M.invs.r.mean,
                          M.shan.r.mean))
# # old
# M.diversity.reord=t(rbind(M.diversity[1,], # total reads unsubsampled
#                           M.diversity[2,], # observed asvs unsubsampled
#                           M.sobs, # observed ASVs
#                           M.diversity[5,], # inverse simpson diversity
#                           M.diversity[8,],
#                           M.diversity[6,],
#                           M.diversity[9,],
#                           rep(M.minreads,ncol(M.diversity)),
#                           M.diversity[2:4,],
#                           M.diversity[7,]))
# M.diversity.reord[,c(1:3,5,7)]=round(M.diversity.reord[,c(1:3,5,7)],digits=0)                                                                                   # rounds indices
# M.diversity.reord[,c(4,6,8,11)]=round(M.diversity.reord[,c(4,6,10,11)],digits=1)                                                                                       # rounds indices
colnames(M.diversity.reord)=c("Total Reads (unsubsampled)",
                              "Observed ASVs (unsubsampled)",
                              "Absolute Single Sequence ASVs (unsubsampled)",
                              "Absolute Single Sequence ASVs in % (unsubsampled)",
                              "Relative Single Sequence ASVs (unsubsampled)",
                              "Relative Single Sequence ASVs in % (unsubsampled)",
                              'Total Reads (subsampled)',
                              "Observed ASVs - Richness (subsampled)",
                              "Estimated ASVs - Chao1 (subsampled)",
                              "Inverse Simpson Diversity (subsampled)",
                              "Shannon Entropy (subsampled)")

data.table::fwrite(as.data.frame(M.diversity.reord),file=file.path(PathToVisuaRAnalysis,"Alpha_Diversity",paste(VisuaRProjectName,"_DiversityIndices.csv",sep="")),col.names = T,row.names = T)

if (SaveWholeworkspace=='N') {rm(M.SSASV.abs.nr.t,M.SSASV.abs.pc.t,M.SSASVs.rel.nr.t,M.SSASVs.rel.pc.t,M.diversity.reord)}
gc()
# #=== 4.2. Visualizes diversity indices ================================================================================================================================================

cat('\n\n4.2. Visualizes diversity indices',file=(file.path(PathToVisuaRAnalysis,paste(VisuaRProjectName,'_Analysis_log.txt',sep=''))),sep='',append=TRUE)


t.M.diversity=t(M.diversity)                                                                                    # Transposes diversity table
if (SaveWholeworkspace=='N') {rm(M.diversity)}
t.M.diversity.groups=cbind(M.contextdata.subset.noNA,t.M.diversity)                                                # adds group vectors to diversity index table. 
if (SaveWholeworkspace=='N') {rm(t.M.diversity)}
t.M.diversity.ord=t.M.diversity.groups[order(t.M.diversity.groups[,MapCol1], t.M.diversity.groups[,MapCol2]),]  # orders table using Grouping1 and Grouping2
if (SaveWholeworkspace=='N') {rm(t.M.diversity.groups)}
t.M.diversity.ord <- cbind(t.M.diversity.ord,M.colvec.ord)

gc()

# #=== 4.2.0. Creates boxplot with total number of observed reads per group and total number of ASVs per group (unsubsamped) ==============================================================================================
cat('\n\n4.2.0. Boxplots of total number of observed reads',file=(file.path(PathToVisuaRAnalysis,paste(VisuaRProjectName,'_Analysis_log.txt',sep=''))),sep='',append=TRUE)
# # we are using the Bonferroni correction (rather conversative) by dividing the significance levels by the number of pairwise comparisons to obtain adjusted thresholds

reads.plot=ggplot(t.M.diversity.ord, aes(x=t.M.diversity.ord[,MapCol1], y=M.ASVs))      # plots number of reads, using MapCol1 (Grouping1) as grouping
reads.plot=reads.plot+
  stat_boxplot(geom="errorbar",width=0.1)+ 
  geom_boxplot(fill=M.col,outlier.size=1) +                                                                     # M.col is a list of as many colours as present in Grouping1. The first colour is assigned to the first Grouping in alphabetical/increasing order. 
  labs(x=NULL,y="Number of reads (unsubsampled)") + 
  stat_summary(fun=mean,colour="black",shape=17,geom="point") +   
  EnvStats::stat_n_text()+                                                                                                # adds the number of samples per group at the bottom of the plot
  theme(axis.title = element_text(colour="black",size=16),
        axis.text = element_text(colour="black"),
        axis.text.x=element_text(angle=45,hjust=1,colour="black",size=14),
        axis.text.y=element_text(colour="black",size=14),
        axis.line=element_line(colour="black"),
        axis.ticks=element_line(colour = "black"),
        panel.grid.major.x = element_blank(),
        panel.grid.major.y = element_line(colour="grey",size=0.25),
        panel.grid.minor.y = element_line(),
        panel.border = element_rect(fill=NA,colour = "black",size=0.5),                             
        panel.background = element_blank())
reads.plot.dot<-reads.plot+
  geom_dotplot(binaxis='y',
               stackdir='center',
               dotsize =0.5,
               fill="black",
               method='histodot',
               binwidth = ((as.numeric(range(t.M.diversity.ord$M.ASVs)[2]-range(t.M.diversity.ord$M.ASVs)[1]))/40),
  )
if (length(M.projects.unique)>1){                                                                               # adds significance bars above the boxplots for each combination of Grouping1
  reads.plot.sig=reads.plot+
    geom_signif(comparisons=combn(sort(as.character(unique(t.M.diversity.ord[,MapCol1]))),2,simplify=F, test="wilcox.test"), # default: wilcox.test()
                vjust = 0.6,
                step_increase=0.06,
                size=0.2,
                textsize=3,
                tip_length=0.01,
                map_signif_level=c("****"=0.0001,"***"=0.001,"**"=0.01,"*"=0.05," "=2))}
if (length(M.projects.unique)>1){                                                                               # adds significance bars above the boxplots for each combination of Grouping1
  reads.plot.sig.cor=reads.plot+
    geom_signif(comparisons=combn(sort(as.character(unique(t.M.diversity.ord[,MapCol1]))),2,simplify=F, test="wilcox.test"), # default: wilcox.test()
                vjust = 0.6,
                step_increase=0.06,
                size=0.2,
                textsize=3,
                tip_length=0.01,
                map_signif_level=c("****"=0.0001/numpairs,"***"=0.001/numpairs,"**"=0.01/numpairs,"*"=0.05/numpairs," "=2))}
print(reads.plot)
print(reads.plot.dot)
if (length(M.projects.unique)>1){                                                                               # adds significance bars above the boxplots for each combination of Grouping1
  print(reads.plot.sig)
  print(reads.plot.sig.cor)
  pdf(file.path(PathToVisuaRAnalysis,"Alpha_Diversity","Diversity_Metrics",paste(VisuaRProjectName,"_NumberOfReads_unsub.pdf",sep = '')),height=5,width=length(M.projects.unique)*3,useDingbats=F)
  reads.Multiplot=ggarrange(reads.plot,reads.plot.dot,reads.plot.sig,reads.plot.sig.cor,ncol=4,nrow=1,align='hv')                                                 # creates a multiplot of the diversity indices
  reads.Multiplot <- annotate_figure(reads.Multiplot,top = text_grob(paste(VisuaRProjectName," Number of Reads, right: Bonferroni-corrected significance"), color = "black", size = 10))
  print(reads.Multiplot)
  dev.off()
  
  test.df <- t.M.diversity.ord[,c(Grouping1,"M.ASVs")]
  test.avgs <- aggregate(t.M.diversity.ord$M.ASVs,by=list(Category=t.M.diversity.ord[[Grouping1]]),FUN=mean)
  test.sds <- aggregate(t.M.diversity.ord$M.ASVs,by=list(Category=t.M.diversity.ord[[Grouping1]]),FUN=sd)

  colnames(test.df)[1] <- "Grouping1"
  test <- rstatix::pairwise_wilcox_test(
    data = test.df,  # Your data frame
    formula = M.ASVs  ~ Grouping1,  # Adjust the formula to match your data
    p.adjust.method = "bonferroni"  # You can choose an appropriate adjustment method
  )
  test$group1avg <- NA
  test$group1sd <- NA
  test$group2avg <- NA
  test$group2sd <- NA
  group1avg.str <- NULL
  group1sd.str <- NULL
  for (numgroups in 1:(length(M.projects.unique)-1)) {
    group1avg.str <- append(group1avg.str,rep(test.avgs[numgroups,2],(length(M.projects.unique)-numgroups)))
    group1sd.str <- append(group1sd.str,rep(test.sds[numgroups,2],(length(M.projects.unique)-numgroups)))
  }
  group2avg.str <- NULL
  group2sd.str <- NULL
  for (numgroups in 2:(length(M.projects.unique))) {
    group2avg.str <- append(group2avg.str,test.avgs[(numgroups:nrow(test.avgs)),2])
    group2sd.str <- append(group2sd.str,test.sds[(numgroups:nrow(test.sds)),2])
  }
  test$group1avg <- group1avg.str
  test$group1sd <- group1sd.str
  test$group2avg <- group2avg.str
  test$group2sd <- group2sd.str
  
  data.table::fwrite(test,file=paste0(PathToVisuaRAnalysis,"/Alpha_Diversity/Diversity_Metrics/",VisuaRProjectName,"_NumberOfReads_unsub_wilcox_results.csv"),col.names = T,row.names = F)
  rm(test.df,test,group2avg.str,group2sd.str,group1avg.str,group1sd.str,test.avgs,test.sds)
} else {
  pdf(file.path(PathToVisuaRAnalysis,"Alpha_Diversity","Diversity_Metrics",paste(VisuaRProjectName,"_NumberOfReads_unsub.pdf",sep = '')),height=5,width=length(M.projects.unique)*3.5,useDingbats=F)
  reads.Multiplot=ggarrange(reads.plot,reads.plot.dot,ncol=2,nrow=1,align='hv')                                                 # creates a multiplot of the diversity indices
  reads.Multiplot <- annotate_figure(reads.Multiplot,top = text_grob(paste(VisuaRProjectName," Number of Reads, right: Bonferroni-corrected significance"), color = "black", size = 10))
  print(reads.Multiplot)
  dev.off()
}

asvs.plot=ggplot(t.M.diversity.ord, aes(x=t.M.diversity.ord[,MapCol1], y=M.sobs))      # plots number of reads, using MapCol1 (Grouping1) as grouping
asvs.plot=asvs.plot+
  stat_boxplot(geom="errorbar",width=0.1)+ 
  geom_boxplot(fill=M.col,outlier.size=1) +                                                                     # M.col is a list of as many colours as present in Grouping1. The first colour is assigned to the first Grouping in alphabetical/increasing order. 
  labs(x=NULL,y="Number of ASVs (unsubsampled)") + 
  stat_summary(fun=mean,colour="black",shape=17,geom="point") +                                               # adds a mean triangle to each boxplot
  stat_n_text()+                                                                                                # adds the number of samples per group at the bottom of the plot
  theme(axis.title = element_text(colour="black",size=16),
        axis.text = element_text(colour="black"),
        axis.text.x=element_text(angle=45,hjust=1,colour="black",size=14),
        axis.text.y=element_text(colour="black",size=14),
        axis.line=element_line(colour="black"),
        axis.ticks=element_line(colour = "black"),
        panel.grid.major.x = element_blank(),
        panel.grid.major.y = element_line(colour="grey",size=0.25),
        panel.grid.minor.y = element_line(),
        panel.border = element_rect(fill=NA,colour = "black",size=0.5),                             
        panel.background = element_blank())
asvs.plot.dot<-asvs.plot+
  geom_dotplot(binaxis='y',
               stackdir='center',
               dotsize =0.5,
               fill="black",
               method='histodot',
               binwidth = ((as.numeric(range(t.M.diversity.ord$M.sobs)[2]-range(t.M.diversity.ord$M.sobs)[1]))/40),
  )
if (length(M.projects.unique)>1){                                                                               # adds significance bars above the boxplots for each combination of Grouping1
  asvs.plot.sig=asvs.plot+
    geom_signif(comparisons=combn(sort(as.character(unique(t.M.diversity.ord[,MapCol1]))),2,simplify=F, test="wilcox.test"), # default: wilcox.test()
                vjust = 0.6,
                step_increase=0.06,
                size=0.2,
                textsize=3,
                tip_length=0.01,
                map_signif_level=c("****"=0.0001,"***"=0.001,"**"=0.01,"*"=0.05," "=2))}
if (length(M.projects.unique)>1){                                                                               # adds significance bars above the boxplots for each combination of Grouping1
  asvs.plot.sig.cor=asvs.plot+
    geom_signif(comparisons=combn(sort(as.character(unique(t.M.diversity.ord[,MapCol1]))),2,simplify=F, test="wilcox.test"), # default: wilcox.test()
                vjust = 0.6,
                step_increase=0.06,
                size=0.2,
                textsize=3,
                tip_length=0.01,
                map_signif_level=c("****"=0.0001/numpairs,"***"=0.001/numpairs,"**"=0.01/numpairs,"*"=0.05/numpairs," "=2))}
print(asvs.plot)
print(asvs.plot.dot)
if (length(M.projects.unique)>1){                                                                               # adds significance bars above the boxplots for each combination of Grouping1
  print(asvs.plot.sig)
  print(asvs.plot.sig.cor)
  pdf(file.path(PathToVisuaRAnalysis,"Alpha_Diversity","Diversity_Metrics",paste(VisuaRProjectName,"_NumberOfASVs_unsub.pdf",sep = '')),height=5,width=length(M.projects.unique)*3,useDingbats=F)
  reads.Multiplot=ggarrange(asvs.plot,asvs.plot.dot,asvs.plot.sig,asvs.plot.sig.cor,ncol=4,nrow=1,align='hv')                                                 # creates a multiplot of the diversity indices
  reads.Multiplot <- annotate_figure(reads.Multiplot,top = text_grob(paste(VisuaRProjectName," Number of ASVs, right: Bonferroni-corrected significance"), color = "black", size = 10))
  print(reads.Multiplot)
  dev.off()
  
  test.df <- t.M.diversity.ord[,c(Grouping1,"M.sobs")]
  test.avgs <- aggregate(t.M.diversity.ord$M.sobs,by=list(Category=t.M.diversity.ord[[Grouping1]]),FUN=mean)
  test.sds <- aggregate(t.M.diversity.ord$M.sobs,by=list(Category=t.M.diversity.ord[[Grouping1]]),FUN=sd)
  
  colnames(test.df)[1] <- "Grouping1"
  test <- rstatix::pairwise_wilcox_test(
    data = test.df,  # Your data frame
    formula = M.sobs  ~ Grouping1,  # Adjust the formula to match your data
    p.adjust.method = "bonferroni"  # You can choose an appropriate adjustment method
  )
  test$group1avg <- NA
  test$group1sd <- NA
  test$group2avg <- NA
  test$group2sd <- NA
  group1avg.str <- NULL
  group1sd.str <- NULL
  for (numgroups in 1:(length(M.projects.unique)-1)) {
    group1avg.str <- append(group1avg.str,rep(test.avgs[numgroups,2],(length(M.projects.unique)-numgroups)))
    group1sd.str <- append(group1sd.str,rep(test.sds[numgroups,2],(length(M.projects.unique)-numgroups)))
  }
  group2avg.str <- NULL
  group2sd.str <- NULL
  for (numgroups in 2:(length(M.projects.unique))) {
    group2avg.str <- append(group2avg.str,test.avgs[(numgroups:nrow(test.avgs)),2])
    group2sd.str <- append(group2sd.str,test.sds[(numgroups:nrow(test.sds)),2])
  }
  test$group1avg <- group1avg.str
  test$group1sd <- group1sd.str
  test$group2avg <- group2avg.str
  test$group2sd <- group2sd.str
  
  data.table::fwrite(test,file=paste0(PathToVisuaRAnalysis,"/Alpha_Diversity/Diversity_Metrics/",VisuaRProjectName,"_NumberOfASVs_unsub_wilcox_results.csv"),col.names = T,row.names = F)
  rm(test.df,test,test.avgs,test.sds,group1avg.str,group1sd.str,group2avg.str,group2sd.str)
} else {
  pdf(file.path(PathToVisuaRAnalysis,"Alpha_Diversity","Diversity_Metrics",paste(VisuaRProjectName,"_NumberOfASVs_unsub.pdf",sep = '')),height=5,width=length(M.projects.unique)*3.5,useDingbats=F)
  reads.Multiplot=ggarrange(asvs.plot,asvs.plot.dot,ncol=2,nrow=1,align='hv')                                                 # creates a multiplot of the diversity indices
  reads.Multiplot <- annotate_figure(reads.Multiplot,top = text_grob(paste(VisuaRProjectName," Number of ASVs, right: Bonferroni-corrected significance"), color = "black", size = 10))
  print(reads.Multiplot)
  dev.off()
}

# #=== 4.2.1. Barplots of diversity indices - per sample ================================================================================================================================================
# Convert group column to factor
t.M.diversity.ord.temp <- t.M.diversity.ord
t.M.diversity.ord.temp[,MapCol1] <- factor(t.M.diversity.ord.temp[,MapCol1])
# # creates column with sample names from row names and reorder the levels of Sample based on Color
t.M.diversity.ord.temp$asdf1234 <- rownames(t.M.diversity.ord.temp)
t.M.diversity.ord.temp$asdf1234 <- factor(t.M.diversity.ord.temp$asdf1234, levels = t.M.diversity.ord.temp$asdf1234[order(t.M.diversity.ord.temp[,MapCol1])])


# Specify the colors and labels for the groups
group_colors <- unique(M.colvec.ord)
group_labels <- levels(t.M.diversity.ord.temp[, MapCol1])

reads.sample.unsub <- ggplot(t.M.diversity.ord.temp, aes(x = asdf1234, y = M.ASVs, fill = t.M.diversity.ord.temp[, MapCol1])) +
  geom_bar(stat = "identity", width = 0.7) +
  labs(x = "", y = "Number of Reads (unsubsampled)") +
  scale_fill_manual(values = group_colors, guide = FALSE) +
  theme(axis.text = element_text(colour = "black"),
        legend.title = element_blank(),
        panel.grid.major.x = element_blank(),
        axis.text.x = element_text(angle = 90, colour = "black", size = 14, vjust = 0.5, hjust = 1),
        axis.text.y = element_text(colour = "black", size = 14),
        axis.title = element_text(colour = "black", size = 16),
        axis.line = element_line(colour = "black"),
        panel.background = element_blank(),
        panel.grid.major.y = element_line(colour = "grey", size = 0.5),
        panel.grid.minor.y = element_line(colour = "grey", size = 0.25),
        panel.border = element_rect(fill = NA, colour = "black", size = 0.5),
        legend.box.background = element_rect(colour = "black"),
        legend.box.margin = margin(0.18, 0.1, 0.1, 0.5),
        legend.key = element_rect(fill = "white"),
        legend.text = element_text(size = 14),
        axis.ticks = element_line(colour = "black")) +
  guides(fill = guide_legend(title = "Groups", override.aes = list(fill = group_colors, labels = group_labels)))
print(reads.sample.unsub)

asvs.sample.unsub <- ggplot(t.M.diversity.ord.temp, aes(x = asdf1234, y = M.sobs, fill = t.M.diversity.ord.temp[, MapCol1])) +
  geom_bar(stat = "identity", width = 0.7) +
  labs(x = "", y = "Number of ASVs (unsubsampled)") +
  scale_fill_manual(values = group_colors, guide = FALSE) +
  theme(axis.text = element_text(colour = "black"),
        legend.title = element_blank(),
        panel.grid.major.x = element_blank(),
        axis.text.x = element_text(angle = 90, colour = "black", size = 14, vjust = 0.5, hjust = 1),
        axis.text.y = element_text(colour = "black", size = 14),
        axis.title = element_text(colour = "black", size = 16),
        axis.line = element_line(colour = "black"),
        panel.background = element_blank(),
        panel.grid.major.y = element_line(colour = "grey", size = 0.5),
        panel.grid.minor.y = element_line(colour = "grey", size = 0.25),
        panel.border = element_rect(fill = NA, colour = "black", size = 0.5),
        legend.box.background = element_rect(colour = "black"),
        legend.box.margin = margin(0.18, 0.1, 0.1, 0.5),
        legend.key = element_rect(fill = "white"),
        legend.text = element_text(size = 14),
        axis.ticks = element_line(colour = "black")) +
  guides(fill = guide_legend(title = "Groups", override.aes = list(fill = group_colors, labels = group_labels)))
print(asvs.sample.unsub)

sobs.sample <- ggplot(t.M.diversity.ord.temp, aes(x = asdf1234, y = M.sobs.r.mean, fill = t.M.diversity.ord.temp[, MapCol1])) +
  geom_bar(stat = "identity", width = 0.7) +
  labs(x = "", y = "Observed ASVs (subsampled)") +
  scale_fill_manual(values = group_colors, guide = FALSE) +
  theme(axis.text = element_text(colour = "black"),
        legend.title = element_blank(),
        panel.grid.major.x = element_blank(),
        axis.text.x = element_text(angle = 90, colour = "black", size = 14, vjust = 0.5, hjust = 1),
        axis.text.y = element_text(colour = "black", size = 14),
        axis.title = element_text(colour = "black", size = 16),
        axis.line = element_line(colour = "black"),
        panel.background = element_blank(),
        panel.grid.major.y = element_line(colour = "grey", size = 0.5),
        panel.grid.minor.y = element_line(colour = "grey", size = 0.25),
        panel.border = element_rect(fill = NA, colour = "black", size = 0.5),
        legend.box.background = element_rect(colour = "black"),
        legend.box.margin = margin(0.18, 0.1, 0.1, 0.5),
        legend.key = element_rect(fill = "white"),
        legend.text = element_text(size = 14),
        axis.ticks = element_line(colour = "black")) +
  guides(fill = guide_legend(title = "Groups", override.aes = list(fill = group_colors, labels = group_labels)))
print(sobs.sample)

shan.sample <- ggplot(t.M.diversity.ord.temp, aes(x = asdf1234, y = M.shan.r.mean, fill = t.M.diversity.ord.temp[, MapCol1])) +
  geom_bar(stat = "identity", width = 0.7) +
  labs(x = "", y = "Shannon Entropy") +
  scale_fill_manual(values = group_colors, guide = FALSE) +
  theme(axis.text = element_text(colour = "black"),
        legend.title = element_blank(),
        panel.grid.major.x = element_blank(),
        axis.text.x = element_text(angle = 90, colour = "black", size = 14, vjust = 0.5, hjust = 1),
        axis.text.y = element_text(colour = "black", size = 14),
        axis.title = element_text(colour = "black", size = 16),
        axis.line = element_line(colour = "black"),
        panel.background = element_blank(),
        panel.grid.major.y = element_line(colour = "grey", size = 0.5),
        panel.grid.minor.y = element_line(colour = "grey", size = 0.25),
        panel.border = element_rect(fill = NA, colour = "black", size = 0.5),
        legend.box.background = element_rect(colour = "black"),
        legend.box.margin = margin(0.18, 0.1, 0.1, 0.5),
        legend.key = element_rect(fill = "white"),
        legend.text = element_text(size = 14),
        axis.ticks = element_line(colour = "black")) +
  guides(fill = guide_legend(title = "Groups", override.aes = list(fill = group_colors, labels = group_labels)))
print(shan.sample)

simp.sample <- ggplot(t.M.diversity.ord.temp, aes(x = asdf1234, y = M.invs.r.mean, fill = t.M.diversity.ord.temp[, MapCol1])) +
  geom_bar(stat = "identity", width = 0.7) +
  labs(x = "", y = "Inverse Simpson Diversity") +
  scale_fill_manual(values = group_colors, guide = FALSE) +
  theme(axis.text = element_text(colour = "black"),
        legend.title = element_blank(),
        panel.grid.major.x = element_blank(),
        axis.text.x = element_text(angle = 90, colour = "black", size = 14, vjust = 0.5, hjust = 1),
        axis.text.y = element_text(colour = "black", size = 14),
        axis.title = element_text(colour = "black", size = 16),
        axis.line = element_line(colour = "black"),
        panel.background = element_blank(),
        panel.grid.major.y = element_line(colour = "grey", size = 0.5),
        panel.grid.minor.y = element_line(colour = "grey", size = 0.25),
        panel.border = element_rect(fill = NA, colour = "black", size = 0.5),
        legend.box.background = element_rect(colour = "black"),
        legend.box.margin = margin(0.18, 0.1, 0.1, 0.5),
        legend.key = element_rect(fill = "white"),
        legend.text = element_text(size = 14),
        axis.ticks = element_line(colour = "black")) +
  guides(fill = guide_legend(title = "Groups", override.aes = list(fill = group_colors, labels = group_labels)))
print(simp.sample)

chao.sample <- ggplot(t.M.diversity.ord.temp, aes(x = asdf1234, y = M.chao1.r.mean, fill = t.M.diversity.ord.temp[, MapCol1])) +
  geom_bar(stat = "identity", width = 0.7) +
  labs(x = "", y = "Chao1 Richness") +
  scale_fill_manual(values = group_colors, guide = FALSE) +
  theme(axis.text = element_text(colour = "black"),
        legend.title = element_blank(),
        panel.grid.major.x = element_blank(),
        axis.text.x = element_text(angle = 90, colour = "black", size = 14, vjust = 0.5, hjust = 1),
        axis.text.y = element_text(colour = "black", size = 14),
        axis.title = element_text(colour = "black", size = 16),
        axis.line = element_line(colour = "black"),
        panel.background = element_blank(),
        panel.grid.major.y = element_line(colour = "grey", size = 0.5),
        panel.grid.minor.y = element_line(colour = "grey", size = 0.25),
        panel.border = element_rect(fill = NA, colour = "black", size = 0.5),
        legend.box.background = element_rect(colour = "black"),
        legend.box.margin = margin(0.18, 0.1, 0.1, 0.5),
        legend.key = element_rect(fill = "white"),
        legend.text = element_text(size = 14),
        axis.ticks = element_line(colour = "black")) +
  guides(fill = guide_legend(title = "Groups", override.aes = list(fill = group_colors, labels = group_labels)))
print(chao.sample)

pdf(file.path(PathToVisuaRAnalysis,"Alpha_Diversity","Diversity_Metrics",paste(VisuaRProjectName,"_DiversityIndices_bySample.pdf",sep = '')),height=30,width=nrow(t.M.diversity.ord.temp)/2.5,useDingbats=F)
Diversity.Multiplot=ggarrange(reads.sample.unsub,asvs.sample.unsub,sobs.sample,shan.sample,simp.sample,chao.sample,ncol=1,nrow=6,align='hv')                                                 # creates a multiplot of the diversity indices
annotate_figure(Diversity.Multiplot,top = text_grob(paste(VisuaRProjectName," Alpha Diversity"), color = "black", size = 14))
dev.off()

rm(t.M.diversity.ord.temp)

closeAllConnections() # closes all currently open connections.
gc()

# #=== 4.2.1. Boxplots of diversity indices - grouped ================================================================================================================================================

cat('\n\n4.2.1. Boxplots of diversity indices',file=(file.path(PathToVisuaRAnalysis,paste(VisuaRProjectName,'_Analysis_log.txt',sep=''))),sep='',append=TRUE)


SOBS.plot=ggplot(t.M.diversity.ord, aes(x=t.M.diversity.ord[,MapCol1], y=M.sobs.r.mean))      # plots Observed ASVs, using MapCol1 (Grouping1) as grouping
SOBS.plot=SOBS.plot+
  stat_boxplot(geom="errorbar",width=0.1)+ 
  geom_boxplot(fill=M.col,outlier.size=1) +                                                                     # M.col is a list of as many colours as present in Grouping1. The first colour is assigned to the first Grouping in alphabetical/increasing order. 
  labs(x=NULL,y="Observed ASVs") + 
  stat_summary(fun=mean,colour="black",shape=17,geom="point") +                                               # adds a mean triangle to each boxplot
  stat_n_text()+                                                                                                # adds the number of samples per group at the bottom of the plot
  theme(axis.title = element_text(colour="black",size=16),
        axis.text = element_text(colour="black"),
        axis.text.x=element_text(angle=45,hjust=1,colour="black",size=14),
        axis.text.y=element_text(colour="black",size=14),
        axis.line=element_line(colour="black"),
        axis.ticks=element_line(colour = "black"),
        panel.grid.major.x = element_blank(),
        panel.grid.major.y = element_line(colour="grey",size=0.25),
        panel.grid.minor.y = element_line(),
        panel.border = element_rect(fill=NA,colour = "black",size=0.5),                             
        panel.background = element_blank())
SOBS.plot.dot<-SOBS.plot+
  geom_dotplot(binaxis='y',
               stackdir='center',
               dotsize =0.5,
               fill="black",
               method='histodot',
               binwidth = ((as.numeric(range(t.M.diversity.ord$M.sobs.r.mean)[2]-range(t.M.diversity.ord$M.sobs.r.mean)[1]))/40),
  )
if (length(M.projects.unique)>1){                                                                               # adds significance bars above the boxplots for each combination of Grouping1
  SOBS.plot.sig=SOBS.plot+
    geom_signif(comparisons=combn(sort(as.character(unique(t.M.diversity.ord[,MapCol1]))),2,simplify=F),
                vjust = 0.6,
                step_increase=0.06,
                size=0.2,
                textsize=3,
                tip_length=0.01,
                map_signif_level=c("****"=0.0001,"***"=0.001,"**"=0.01,"*"=0.05," "=2))}
if (length(M.projects.unique)>1){                                                                               # adds significance bars above the boxplots for each combination of Grouping1
  SOBS.plot.sig.cor=SOBS.plot+
    geom_signif(comparisons=combn(sort(as.character(unique(t.M.diversity.ord[,MapCol1]))),2,simplify=F),
                vjust = 0.6,
                step_increase=0.06,
                size=0.2,
                textsize=3,
                tip_length=0.01,
                map_signif_level=c("****"=0.0001/numpairs,"***"=0.001/numpairs,"**"=0.01/numpairs,"*"=0.05/numpairs," "=2))}
print(SOBS.plot)
print(SOBS.plot.dot)
if (length(M.projects.unique)>1){                                                                               # adds significance bars above the boxplots for each combination of Grouping1
  print(SOBS.plot.sig)
  print(SOBS.plot.sig.cor)
}

Shan.plot=ggplot(t.M.diversity.ord, aes(x=t.M.diversity.ord[,MapCol1], y=(M.shan.r.mean)))    # plots Shannon
Shan.plot=Shan.plot+
  stat_boxplot(geom="errorbar",width=0.1)+ 
  geom_boxplot(fill=M.col,outlier.size=1) +
  labs(x=NULL, y="Shannon Entropy") + 
  stat_summary(fun=mean,colour="black",geom="point",shape=17)+
  stat_n_text()+
  theme(axis.title = element_text(colour="black",size=16),
        axis.text = element_text(colour="black"),
        axis.text.x=element_text(angle=45,hjust=1,colour="black",size=14),
        axis.text.y=element_text(colour="black",size=14),
        axis.line=element_line(colour="black"),
        axis.ticks=element_line(colour = "black"),
        panel.grid.major.x = element_blank(),
        panel.grid.major.y = element_line(colour="grey",size=0.25),
        panel.grid.minor.y = element_line(),
        panel.border = element_rect(fill=NA,colour = "black",size=0.5),
        panel.background = element_blank())
Shan.plot.dot<-Shan.plot+
  geom_dotplot(binaxis='y',
               stackdir='center',
               dotsize =0.5,
               fill="black",
               method='histodot',
               binwidth = ((as.numeric(range(t.M.diversity.ord$M.shan.r.mean)[2]-range(t.M.diversity.ord$M.shan.r.mean)[1]))/40),
  ) 
if (length(M.projects.unique)>1){ 
  Shan.plot.sig=Shan.plot+ 
    geom_signif(comparisons=combn(sort(as.character(unique(t.M.diversity.ord[,MapCol1]))),2,simplify=F),
                vjust = 0.6,
                step_increase=0.06,
                size=0.2,
                textsize=3,
                tip_length=0.01,
                map_signif_level=c("****"=0.0001,"***"=0.001,"**"=0.01,"*"=0.05," "=2))}
if (length(M.projects.unique)>1){ 
  Shan.plot.sig.cor=Shan.plot+ 
    geom_signif(comparisons=combn(sort(as.character(unique(t.M.diversity.ord[,MapCol1]))),2,simplify=F),
                vjust = 0.6,
                step_increase=0.06,
                size=0.2,
                textsize=3,
                tip_length=0.01,
                map_signif_level=c("****"=0.0001/numpairs,"***"=0.001/numpairs,"**"=0.01/numpairs,"*"=0.05/numpairs," "=2))}
print(Shan.plot)
print(Shan.plot.dot)
if (length(M.projects.unique)>1){ 
  print(Shan.plot.sig)
  print(Shan.plot.sig.cor)
}

Simp.plot=ggplot(t.M.diversity.ord, aes(x=t.M.diversity.ord[,MapCol1], y=(M.invs.r.mean)))    # plots Inverse Simpson
Simp.plot=Simp.plot+
  stat_boxplot(geom="errorbar",width=0.1)+ 
  geom_boxplot(fill=M.col,outlier.size=1) +
  labs(x=NULL, y="Inverse Simpson Diversity", col=Grouping1) + 
  stat_summary(fun=mean,colour="black",geom="point",shape=17) +
  stat_n_text()+
  theme(axis.title = element_text(colour="black",size=16),
        axis.text = element_text(colour="black"),
        axis.text.x=element_text(angle=45,hjust=1,colour="black",size=14),
        axis.text.y=element_text(colour="black",size=14),
        axis.line=element_line(colour="black"),
        axis.ticks=element_line(colour = "black"),
        panel.grid.major.x = element_blank(),
        panel.grid.major.y = element_line(colour="grey",size=0.25),
        panel.grid.minor.y = element_line(),
        panel.border = element_rect(fill=NA,colour = "black",size=0.5), 
        panel.background = element_blank())
Simp.plot.dot<-Simp.plot+
  geom_dotplot(binaxis='y',
               stackdir='center',
               dotsize =0.5,
               fill="black",
               method='histodot',
               binwidth = ((as.numeric(range(t.M.diversity.ord$M.invs.r.mean)[2]-range(t.M.diversity.ord$M.invs.r.mean)[1]))/40),
  ) 
if (length(M.projects.unique)>1){
  Simp.plot.sig=Simp.plot+ 
    geom_signif(comparisons=combn(sort(as.character(unique(t.M.diversity.ord[,MapCol1]))),2,simplify=F),
                vjust = 0.6,
                step_increase=0.06,
                size=0.2,
                textsize=3,
                tip_length=0.01,
                map_signif_level=c("****"=0.0001,"***"=0.001,"**"=0.01,"*"=0.05," "=2))}
if (length(M.projects.unique)>1){
  Simp.plot.sig.cor=Simp.plot+ 
    geom_signif(comparisons=combn(sort(as.character(unique(t.M.diversity.ord[,MapCol1]))),2,simplify=F),
                vjust = 0.6,
                step_increase=0.06,
                size=0.2,
                textsize=3,
                tip_length=0.01,
                map_signif_level=c("****"=0.0001/numpairs,"***"=0.001/numpairs,"**"=0.01/numpairs,"*"=0.05/numpairs," "=2))}
print(Simp.plot)
print(Simp.plot.dot)
if (length(M.projects.unique)>1){
  print(Simp.plot.sig)
  print(Simp.plot.sig.cor)
}


Chao1.plot=ggplot(t.M.diversity.ord, aes(x=t.M.diversity.ord[,MapCol1], y=(M.chao1.r.mean)))    # plots Chao1 richness
Chao1.plot=Chao1.plot+
  stat_boxplot(geom="errorbar",width=0.1)+ 
  geom_boxplot(fill=M.col,outlier.size=1) +
  labs(x=NULL, y="Chao1 Richness", col=Grouping1) + 
  stat_summary(fun=mean,colour="black",geom="point",shape=17) +
  stat_n_text()+
  theme(axis.title = element_text(colour="black",size=16),
        axis.text = element_text(colour="black"),
        axis.text.x=element_text(angle=45,hjust=1,colour="black",size=14),
        axis.text.y=element_text(colour="black",size=14),
        axis.line=element_line(colour="black"),
        axis.ticks=element_line(colour = "black"),
        panel.grid.major.x = element_blank(),
        panel.grid.major.y = element_line(colour="grey",size=0.25),
        panel.grid.minor.y = element_line(),
        panel.border = element_rect(fill=NA,colour = "black",size=0.5), 
        panel.background = element_blank())
Chao1.plot.dot<-Chao1.plot+
  geom_dotplot(binaxis='y',
               stackdir='center',
               dotsize =0.5,
               fill="black",
               method='histodot',
               binwidth = ((as.numeric(range(t.M.diversity.ord$M.chao1.r.mean)[2]-range(t.M.diversity.ord$M.chao1.r.mean)[1]))/40),
  ) 
if (length(M.projects.unique)>1){
  Chao1.plot.sig=Chao1.plot+ 
    geom_signif(comparisons=combn(sort(as.character(unique(t.M.diversity.ord[,MapCol1]))),2,simplify=F),
                vjust = 0.6,
                step_increase=0.06,
                size=0.2,
                textsize=3,
                tip_length=0.01,
                map_signif_level=c("****"=0.0001,"***"=0.001,"**"=0.01,"*"=0.05," "=2))}
if (length(M.projects.unique)>1){
  Chao1.plot.sig.cor=Chao1.plot+ 
    geom_signif(comparisons=combn(sort(as.character(unique(t.M.diversity.ord[,MapCol1]))),2,simplify=F),
                vjust = 0.6,
                step_increase=0.06,
                size=0.2,
                textsize=3,
                tip_length=0.01,
                map_signif_level=c("****"=0.0001/numpairs,"***"=0.001/numpairs,"**"=0.01/numpairs,"*"=0.05/numpairs," "=2))}
print(Chao1.plot)
print(Chao1.plot.dot)
if (length(M.projects.unique)>1){
  print(Chao1.plot.sig)
  print(Chao1.plot.sig.cor)
}

pdf(file.path(PathToVisuaRAnalysis,"Alpha_Diversity","Diversity_Metrics",paste(VisuaRProjectName,"_DiversityIndices.pdf",sep = '')),height=4,width=length(M.projects.unique)*3,useDingbats=F)
Diversity.Multiplot=ggarrange(SOBS.plot,Shan.plot,Simp.plot,Chao1.plot,ncol=4,nrow=1,align='hv')                                                 # creates a multiplot of the diversity indices
annotate_figure(Diversity.Multiplot,top = text_grob(paste(VisuaRProjectName," Alpha Diversity"), color = "black", size = 14))
dev.off()

pdf(file.path(PathToVisuaRAnalysis,"Alpha_Diversity","Diversity_Metrics",paste(VisuaRProjectName,"_DiversityIndices_dots.pdf",sep = '')),height=4,width=length(M.projects.unique)*3,useDingbats=F)
Diversity.Multiplot=ggarrange(SOBS.plot.dot,Shan.plot.dot,Simp.plot.dot,Chao1.plot.dot,ncol=4,nrow=1,align='hv')                                                 # creates a multiplot of the diversity indices
annotate_figure(Diversity.Multiplot,top = text_grob(paste(VisuaRProjectName," Alpha Diversity"), color = "black", size = 14))
dev.off()

if (length(M.projects.unique)>1){
  pdf(file.path(PathToVisuaRAnalysis,"Alpha_Diversity","Diversity_Metrics",paste(VisuaRProjectName,"_DiversityIndices_sig.pdf",sep = '')),height=4,width=length(M.projects.unique)*3,useDingbats=F)
  Diversity.Multiplot=ggarrange(SOBS.plot.sig,Shan.plot.sig,Simp.plot.sig,Chao1.plot.sig,ncol=4,nrow=1,align='hv')    
  Diversity.Multiplot <- annotate_figure(Diversity.Multiplot,top = text_grob(paste(VisuaRProjectName," Alpha Diversity - Significances uncorrected"), color = "black", size = 10))
  print(Diversity.Multiplot)
  dev.off()
  rm(Diversity.Multiplot)
 
  pdf(file.path(PathToVisuaRAnalysis,"Alpha_Diversity","Diversity_Metrics",paste(VisuaRProjectName,"_DiversityIndices_sig_cor.pdf",sep = '')),height=4,width=length(M.projects.unique)*3,useDingbats=F)
  Diversity.Multiplot=ggarrange(SOBS.plot.sig.cor,Shan.plot.sig.cor,Simp.plot.sig.cor,Chao1.plot.sig.cor,ncol=4,nrow=1,align='hv')                                                 # creates a multiplot of the diversity indices
  Diversity.Multiplot <- annotate_figure(Diversity.Multiplot,top = text_grob(paste(VisuaRProjectName," Alpha Diversity - Significances Bonferroni corrected"), color = "black", size = 10))
  print(Diversity.Multiplot)
  dev.off()
  rm(Diversity.Multiplot)
  
  test.df <- t.M.diversity.ord[,c(Grouping1,"M.sobs.r.mean")]
  test.avgs <- aggregate(t.M.diversity.ord$M.sobs.r.mean,by=list(Category=t.M.diversity.ord[[Grouping1]]),FUN=mean)
  test.sds <- aggregate(t.M.diversity.ord$M.sobs.r.mean,by=list(Category=t.M.diversity.ord[[Grouping1]]),FUN=sd)
  
  colnames(test.df)[1] <- "Grouping1"
  test <- rstatix::pairwise_wilcox_test(
    data = test.df,  # Your data frame
    formula = M.sobs.r.mean  ~ Grouping1,  # Adjust the formula to match your data
    p.adjust.method = "bonferroni"  # You can choose an appropriate adjustment method
  )
  test$group1avg <- NA
  test$group1sd <- NA
  test$group2avg <- NA
  test$group2sd <- NA
  group1avg.str <- NULL
  group1sd.str <- NULL
  for (numgroups in 1:(length(M.projects.unique)-1)) {
    group1avg.str <- append(group1avg.str,rep(test.avgs[numgroups,2],(length(M.projects.unique)-numgroups)))
    group1sd.str <- append(group1sd.str,rep(test.sds[numgroups,2],(length(M.projects.unique)-numgroups)))
  }
  group2avg.str <- NULL
  group2sd.str <- NULL
  for (numgroups in 2:(length(M.projects.unique))) {
    group2avg.str <- append(group2avg.str,test.avgs[(numgroups:nrow(test.avgs)),2])
    group2sd.str <- append(group2sd.str,test.sds[(numgroups:nrow(test.sds)),2])
  }
  test$group1avg <- group1avg.str
  test$group1sd <- group1sd.str
  test$group2avg <- group2avg.str
  test$group2sd <- group2sd.str
  
  data.table::fwrite(test,file=paste0(PathToVisuaRAnalysis,"/Alpha_Diversity/Diversity_Metrics/",VisuaRProjectName,"_Richness_wilcox_results.csv"),col.names = T,row.names = F)
  rm(test.df,test)
  
  test.df <- t.M.diversity.ord[,c(Grouping1,"M.shan.r.mean")]
  test.avgs <- aggregate(t.M.diversity.ord$M.shan.r.mean,by=list(Category=t.M.diversity.ord[[Grouping1]]),FUN=mean)
  test.sds <- aggregate(t.M.diversity.ord$M.shan.r.mean,by=list(Category=t.M.diversity.ord[[Grouping1]]),FUN=sd)
  
  colnames(test.df)[1] <- "Grouping1"
  test <- rstatix::pairwise_wilcox_test(
    data = test.df,  # Your data frame
    formula = M.shan.r.mean  ~ Grouping1,  # Adjust the formula to match your data
    p.adjust.method = "bonferroni"  # You can choose an appropriate adjustment method
  )
  test$group1avg <- NA
  test$group1sd <- NA
  test$group2avg <- NA
  test$group2sd <- NA
  group1avg.str <- NULL
  group1sd.str <- NULL
  for (numgroups in 1:(length(M.projects.unique)-1)) {
    group1avg.str <- append(group1avg.str,rep(test.avgs[numgroups,2],(length(M.projects.unique)-numgroups)))
    group1sd.str <- append(group1sd.str,rep(test.sds[numgroups,2],(length(M.projects.unique)-numgroups)))
  }
  group2avg.str <- NULL
  group2sd.str <- NULL
  for (numgroups in 2:(length(M.projects.unique))) {
    group2avg.str <- append(group2avg.str,test.avgs[(numgroups:nrow(test.avgs)),2])
    group2sd.str <- append(group2sd.str,test.sds[(numgroups:nrow(test.sds)),2])
  }
  test$group1avg <- group1avg.str
  test$group1sd <- group1sd.str
  test$group2avg <- group2avg.str
  test$group2sd <- group2sd.str
  
  data.table::fwrite(test,file=paste0(PathToVisuaRAnalysis,"/Alpha_Diversity/Diversity_Metrics/",VisuaRProjectName,"_Shannon_wilcox_results.csv"),col.names = T,row.names = F)
  rm(test.df,test)
  
  test.df <- t.M.diversity.ord[,c(Grouping1,"M.invs.r.mean")]
  test.avgs <- aggregate(t.M.diversity.ord$M.invs.r.mean,by=list(Category=t.M.diversity.ord[[Grouping1]]),FUN=mean)
  test.sds <- aggregate(t.M.diversity.ord$M.invs.r.mean,by=list(Category=t.M.diversity.ord[[Grouping1]]),FUN=sd)
  
  colnames(test.df)[1] <- "Grouping1"
  test <- rstatix::pairwise_wilcox_test(
    data = test.df,  # Your data frame
    formula = M.invs.r.mean  ~ Grouping1,  # Adjust the formula to match your data
    p.adjust.method = "bonferroni"  # You can choose an appropriate adjustment method
  )
  test$group1avg <- NA
  test$group1sd <- NA
  test$group2avg <- NA
  test$group2sd <- NA
  group1avg.str <- NULL
  group1sd.str <- NULL
  for (numgroups in 1:(length(M.projects.unique)-1)) {
    group1avg.str <- append(group1avg.str,rep(test.avgs[numgroups,2],(length(M.projects.unique)-numgroups)))
    group1sd.str <- append(group1sd.str,rep(test.sds[numgroups,2],(length(M.projects.unique)-numgroups)))
  }
  group2avg.str <- NULL
  group2sd.str <- NULL
  for (numgroups in 2:(length(M.projects.unique))) {
    group2avg.str <- append(group2avg.str,test.avgs[(numgroups:nrow(test.avgs)),2])
    group2sd.str <- append(group2sd.str,test.sds[(numgroups:nrow(test.sds)),2])
  }
  test$group1avg <- group1avg.str
  test$group1sd <- group1sd.str
  test$group2avg <- group2avg.str
  test$group2sd <- group2sd.str
  
  data.table::fwrite(test,file=paste0(PathToVisuaRAnalysis,"/Alpha_Diversity/Diversity_Metrics/",VisuaRProjectName,"_InvSimpson_wilcox_results.csv"),col.names = T,row.names = F)
  rm(test.df,test)
  
  test.df <- t.M.diversity.ord[,c(Grouping1,"M.chao1.r.mean")]
  test.avgs <- aggregate(t.M.diversity.ord$M.chao1.r.mean,by=list(Category=t.M.diversity.ord[[Grouping1]]),FUN=mean)
  test.sds <- aggregate(t.M.diversity.ord$M.chao1.r.mean,by=list(Category=t.M.diversity.ord[[Grouping1]]),FUN=sd)
  
  colnames(test.df)[1] <- "Grouping1"
  test <- rstatix::pairwise_wilcox_test(
    data = test.df,  # Your data frame
    formula = M.chao1.r.mean  ~ Grouping1,  # Adjust the formula to match your data
    p.adjust.method = "bonferroni"  # You can choose an appropriate adjustment method
  )
  test$group1avg <- NA
  test$group1sd <- NA
  test$group2avg <- NA
  test$group2sd <- NA
  group1avg.str <- NULL
  group1sd.str <- NULL
  for (numgroups in 1:(length(M.projects.unique)-1)) {
    group1avg.str <- append(group1avg.str,rep(test.avgs[numgroups,2],(length(M.projects.unique)-numgroups)))
    group1sd.str <- append(group1sd.str,rep(test.sds[numgroups,2],(length(M.projects.unique)-numgroups)))
  }
  group2avg.str <- NULL
  group2sd.str <- NULL
  for (numgroups in 2:(length(M.projects.unique))) {
    group2avg.str <- append(group2avg.str,test.avgs[(numgroups:nrow(test.avgs)),2])
    group2sd.str <- append(group2sd.str,test.sds[(numgroups:nrow(test.sds)),2])
  }
  test$group1avg <- group1avg.str
  test$group1sd <- group1sd.str
  test$group2avg <- group2avg.str
  test$group2sd <- group2sd.str
  
  
  data.table::fwrite(test,file=paste0(PathToVisuaRAnalysis,"/Alpha_Diversity/Diversity_Metrics/",VisuaRProjectName,"_Chao1_wilcox_results.csv"),col.names = T,row.names = F)
  rm(test.df,test)
}
closeAllConnections() # closes all currently open connections.
gc()

# #===  4.2.2. Violinplots of diversity indices ================================================================================================================================================

cat('\n\n4.2.2. Violinplots of diversity indices',file=(file.path(PathToVisuaRAnalysis,paste(VisuaRProjectName,'_Analysis_log.txt',sep=''))),sep='',append=TRUE)

SOBS.plot.vio = ggplot(t.M.diversity.ord, aes(x=t.M.diversity.ord[,MapCol1],y=M.sobs.r.mean,fill=t.M.diversity.ord[,MapCol1])) 
SOBS.plot.vio=SOBS.plot.vio +
  geom_violin(trim=T, aes(alpha=0.5)) + 
  # scale_fill_manual(values=M.col, alpha=0.5) +
  # geom_violin(trim=T, draw_quantiles = c(0.25,0.5,0.75)) + 
  scale_fill_manual(values=M.col)+
  geom_boxplot(notch = T,width=0.05) +
  stat_summary(fun=mean,colour="black",geom="point",shape=17)+
  stat_n_text()+
  theme(legend.position = "none")+
  labs(x=NULL,y="Observed ASVs") +
  theme(axis.title = element_text(colour="black",size=16),
        axis.text = element_text(colour="black"),
        axis.text.x=element_text(angle=45,hjust=1,colour="black",size=14),
        axis.text.y=element_text(colour="black",size=14),
        axis.line=element_line(colour="black"),
        axis.ticks=element_line(colour = "black"),
        panel.grid.major.x = element_blank(),
        panel.grid.major.y = element_line(colour="grey",size=0.25),
        panel.grid.minor.y = element_line(),
        panel.border = element_rect(fill=NA,colour = "black",size=0.5),
        panel.background = element_blank())
SOBS.plot.vio.dot <- SOBS.plot.vio +
  geom_dotplot(binaxis='y',
               stackdir='center',
               dotsize =0.5,
               fill="black",
               method='histodot',
               binwidth = ((as.numeric(range(t.M.diversity.ord$M.sobs.r.mean)[2]-range(t.M.diversity.ord$M.sobs.r.mean)[1]))/40),
  ) 
if (length(M.projects.unique)>1){ 
  SOBS.plot.vio.sig=SOBS.plot.vio+ 
    geom_signif(comparisons=combn(sort(as.character(unique(t.M.diversity.ord[,MapCol1]))),2,simplify=F),
                vjust = 0.1,
                step_increase=0.06,
                size=0.2,
                textsize=3,
                tip_length=0.01,
                map_signif_level=c("****"=0.0001,"***"=0.001,"**"=0.01,"*"=0.05," "=2))}
if (length(M.projects.unique)>1){ 
  SOBS.plot.vio.sig.cor=SOBS.plot.vio+ 
    geom_signif(comparisons=combn(sort(as.character(unique(t.M.diversity.ord[,MapCol1]))),2,simplify=F),
                vjust = 0.1,
                step_increase=0.06,
                size=0.2,
                textsize=3,
                tip_length=0.01,
                map_signif_level=c("****"=0.0001/numpairs,"***"=0.001/numpairs,"**"=0.01/numpairs,"*"=0.05/numpairs," "=2))}
print(SOBS.plot.vio)
print(SOBS.plot.vio.dot)
if (length(M.projects.unique)>1){ 
  print(SOBS.plot.vio.sig)
  print(SOBS.plot.vio.sig.cor)
}

Shan.plot.vio = ggplot(t.M.diversity.ord, aes(x=t.M.diversity.ord[,MapCol1],y=M.shan.r.mean,fill=t.M.diversity.ord[,MapCol1])) 
Shan.plot.vio=Shan.plot.vio +
  geom_violin(trim=T, aes(alpha=0.5)) + 
  # scale_fill_manual(values=M.col, alpha=0.5) +
  # geom_violin(trim=T, draw_quantiles = c(0.25,0.5,0.75)) + 
  scale_fill_manual(values=M.col)+
  geom_boxplot(notch = T,width=0.05) +
  stat_summary(fun=mean,colour="black",geom="point",shape=17)+
  stat_n_text()+
  theme(legend.position = "none")+
  labs(x=NULL,y="Shannon Entropy") + 
  theme(axis.title = element_text(colour="black",size=16),
        axis.text = element_text(colour="black"),
        axis.text.x=element_text(angle=45,hjust=1,colour="black",size=14),
        axis.text.y=element_text(colour="black",size=14),
        axis.line=element_line(colour="black"),
        axis.ticks=element_line(colour = "black"),
        panel.grid.major.x = element_blank(),
        panel.grid.major.y = element_line(colour="grey",size=0.25),
        panel.grid.minor.y = element_line(),
        panel.border = element_rect(fill=NA,colour = "black",size=0.5), 
        panel.background = element_blank())
Shan.plot.vio.dot <- Shan.plot.vio +
  geom_dotplot(binaxis='y',
               stackdir='center',
               dotsize =0.5,
               fill="black",
               method='histodot',
               binwidth = ((as.numeric(range(t.M.diversity.ord$M.shan.r.mean)[2]-range(t.M.diversity.ord$M.shan.r.mean)[1]))/40),
  ) 
if (length(M.projects.unique)>1){
  Shan.plot.vio.sig=Shan.plot.vio+
    geom_signif(comparisons=combn(sort(as.character(unique(t.M.diversity.ord[,MapCol1]))),2,simplify=F),
                vjust = 0.1,
                step_increase=0.06,
                size=0.2,
                textsize=3,
                tip_length=0.01,
                map_signif_level=c("****"=0.0001,"***"=0.001,"**"=0.01,"*"=0.05," "=2))}
if (length(M.projects.unique)>1){
  Shan.plot.vio.sig.cor=Shan.plot.vio+
    geom_signif(comparisons=combn(sort(as.character(unique(t.M.diversity.ord[,MapCol1]))),2,simplify=F),
                vjust = 0.1,
                step_increase=0.06,
                size=0.2,
                textsize=3,
                tip_length=0.01,
                map_signif_level=c("****"=0.0001/numpairs,"***"=0.001/numpairs,"**"=0.01/numpairs,"*"=0.05/numpairs," "=2))}
print(Shan.plot.vio)
print(Shan.plot.vio.dot)
if (length(M.projects.unique)>1){ 
  print(Shan.plot.vio.sig)
  print(Shan.plot.vio.sig.cor)
}

Simp.plot.vio = ggplot(t.M.diversity.ord, aes(x=t.M.diversity.ord[,MapCol1],y=M.invs.r.mean,fill=t.M.diversity.ord[,MapCol1])) 
Simp.plot.vio=Simp.plot.vio + 
  geom_violin(trim=T, aes(alpha=0.5)) + 
  # scale_fill_manual(values=M.col, alpha=0.5) +
  # geom_violin(trim=T, draw_quantiles = c(0.25,0.5,0.75)) + 
  scale_fill_manual(values=M.col)+
  geom_boxplot(notch = T,width=0.05) +
  stat_summary(fun=mean,colour="black",geom="point",shape=17)+
  stat_n_text()+
  theme(legend.position = "none")+
  labs(x=NULL,y="Inverse Simpson Diversity") + 
  theme(axis.title = element_text(colour="black",size=16),
        axis.text = element_text(colour="black"),
        axis.text.x=element_text(angle=45,hjust=1,colour="black",size=14),
        axis.text.y=element_text(colour="black",size=14),
        axis.line=element_line(colour="black"),
        axis.ticks=element_line(colour = "black"),
        panel.grid.major.x = element_blank(),
        panel.grid.major.y = element_line(colour="grey",size=0.25),
        panel.grid.minor.y = element_line(),
        panel.border = element_rect(fill=NA,colour = "black",size=0.5), 
        panel.background = element_blank())
Simp.plot.vio.dot <- Simp.plot.vio +
  geom_dotplot(binaxis='y',
               stackdir='center',
               dotsize =0.5,
               fill="black",
               method='histodot',
               binwidth = ((as.numeric(range(t.M.diversity.ord$M.invs.r.mean)[2]-range(t.M.diversity.ord$M.invs.r.mean)[1]))/40),
  ) 
if (length(M.projects.unique)>1){
  Simp.plot.vio.sig=Simp.plot.vio+
    geom_signif(comparisons=combn(sort(as.character(unique(t.M.diversity.ord[,MapCol1]))),2,simplify=F),
                vjust = 0.1,
                step_increase=0.06,
                size=0.2,
                textsize=3,
                tip_length=0.01,
                map_signif_level=c("****"=0.0001,"***"=0.001,"**"=0.01,"*"=0.05," "=2))}
if (length(M.projects.unique)>1){
  Simp.plot.vio.sig.cor=Simp.plot.vio+
    geom_signif(comparisons=combn(sort(as.character(unique(t.M.diversity.ord[,MapCol1]))),2,simplify=F),
                vjust = 0.1,
                step_increase=0.06,
                size=0.2,
                textsize=3,
                tip_length=0.01,
                map_signif_level=c("****"=0.0001/numpairs,"***"=0.001/numpairs,"**"=0.01/numpairs,"*"=0.05/numpairs," "=2))}
print(Simp.plot.vio)
print(Simp.plot.vio.dot)
if (length(M.projects.unique)>1){ 
  print(Simp.plot.vio.sig)
  print(Simp.plot.vio.sig.cor)
}
Chao1.plot.vio = ggplot(t.M.diversity.ord, aes(x=t.M.diversity.ord[,MapCol1],y=M.chao1.r.mean,fill=t.M.diversity.ord[,MapCol1])) 
Chao1.plot.vio=Chao1.plot.vio + 
  geom_violin(trim=T, aes(alpha=0.5)) + 
  # scale_fill_manual(values=M.col, alpha=0.5) +
  # geom_violin(trim=T, draw_quantiles = c(0.25,0.5,0.75)) + 
  scale_fill_manual(values=M.col)+
  geom_boxplot(notch = T,width=0.05) +
  stat_summary(fun=mean,colour="black",geom="point",shape=17)+
  stat_n_text()+
  theme(legend.position = "none")+
  labs(x=NULL,y="Chao1 Richness") + 
  theme(axis.title = element_text(colour="black",size=16),
        axis.text = element_text(colour="black"),
        axis.text.x=element_text(angle=45,hjust=1,colour="black",size=14),
        axis.text.y=element_text(colour="black",size=14),
        axis.line=element_line(colour="black"),
        axis.ticks=element_line(colour = "black"),
        panel.grid.major.x = element_blank(),
        panel.grid.major.y = element_line(colour="grey",size=0.25),
        panel.grid.minor.y = element_line(),
        panel.border = element_rect(fill=NA,colour = "black",size=0.5), 
        panel.background = element_blank())
Chao1.plot.vio.dot <- Chao1.plot.vio +
  geom_dotplot(binaxis='y',
               stackdir='center',
               dotsize =0.5,
               fill="black",
               method='histodot',
               binwidth = ((as.numeric(range(t.M.diversity.ord$M.chao1.r.mean)[2]-range(t.M.diversity.ord$M.chao1.r.mean)[1]))/40),
  ) 
if (length(M.projects.unique)>1){
  Chao1.plot.vio.sig=Chao1.plot.vio+
    geom_signif(comparisons=combn(sort(as.character(unique(t.M.diversity.ord[,MapCol1]))),2,simplify=F),
                vjust = 0.1,
                step_increase=0.06,
                size=0.2,
                textsize=3,
                tip_length=0.01,
                map_signif_level=c("****"=0.0001,"***"=0.001,"**"=0.01,"*"=0.05," "=2))}
if (length(M.projects.unique)>1){
  Chao1.plot.vio.sig.cor=Chao1.plot.vio+
    geom_signif(comparisons=combn(sort(as.character(unique(t.M.diversity.ord[,MapCol1]))),2,simplify=F),
                vjust = 0.1,
                step_increase=0.06,
                size=0.2,
                textsize=3,
                tip_length=0.01,
                map_signif_level=c("****"=0.0001/numpairs,"***"=0.001/numpairs,"**"=0.01/numpairs,"*"=0.05/numpairs," "=2))}
print(Chao1.plot.vio)
print(Chao1.plot.vio.dot)
if (length(M.projects.unique)>1){ 
  print(Chao1.plot.vio.sig)
  print(Chao1.plot.vio.sig.cor)
}

pdf(file.path(PathToVisuaRAnalysis,"Alpha_Diversity","Diversity_Metrics",paste(VisuaRProjectName,"_DiversityIndices_Violin.pdf",sep = '')),height=4,width=length(M.projects.unique)*3,useDingbats=F)
Diversity.Multiplot.vio=ggarrange(SOBS.plot.vio,Shan.plot.vio,Simp.plot.vio,Chao1.plot.vio,ncol=4,nrow=1,align='hv')                                  # creates a multiplot of the diversity indices
annotate_figure(Diversity.Multiplot.vio,top = text_grob(paste(VisuaRProjectName," Alpha Diversity"), color = "black", size = 14))
dev.off()

pdf(file.path(PathToVisuaRAnalysis,"Alpha_Diversity","Diversity_Metrics",paste(VisuaRProjectName,"_DiversityIndices_Violin_dots.pdf",sep = '')),height=4,width=length(M.projects.unique)*3,useDingbats=F)
Diversity.Multiplot.vio=ggarrange(SOBS.plot.vio.dot,Shan.plot.vio.dot,Simp.plot.vio.dot,Chao1.plot.vio.dot,ncol=4,nrow=1,align='hv')                                  # creates a multiplot of the diversity indices
annotate_figure(Diversity.Multiplot.vio,top = text_grob(paste(VisuaRProjectName," Alpha Diversity"), color = "black", size = 14))
dev.off()

if (length(M.projects.unique)>1){ 
  pdf(file.path(PathToVisuaRAnalysis,"Alpha_Diversity","Diversity_Metrics",paste(VisuaRProjectName,"_DiversityIndices_Violin_sig.pdf",sep = '')),height=4,width=length(M.projects.unique)*3,useDingbats=F)
  Diversity.Multiplot.vio=ggarrange(SOBS.plot.vio.sig,Shan.plot.vio.sig,Simp.plot.vio.sig,Chao1.plot.vio.sig,ncol=4,nrow=1,align='hv')                                  # creates a multiplot of the diversity indices
  Diversity.Multiplot.vio <- annotate_figure(Diversity.Multiplot.vio,top = text_grob(paste(VisuaRProjectName," Alpha Diversity - Significances uncorrected"), color = "black", size = 10))
  print(Diversity.Multiplot.vio)
  dev.off()
  
  rm(Diversity.Multiplot.vio)
  pdf(file.path(PathToVisuaRAnalysis,"Alpha_Diversity","Diversity_Metrics",paste(VisuaRProjectName,"_DiversityIndices_Violin_sig_cor.pdf",sep = '')),height=4,width=length(M.projects.unique)*3,useDingbats=F)
  Diversity.Multiplot.vio=ggarrange(SOBS.plot.vio.sig.cor,Shan.plot.vio.sig.cor,Simp.plot.vio.sig.cor,Chao1.plot.vio.sig.cor,ncol=4,nrow=1,align='hv')                                  # creates a multiplot of the diversity indices
  Diversity.Multiplot.vio <- annotate_figure(Diversity.Multiplot.vio,top = text_grob(paste(VisuaRProjectName," Alpha Diversity - Significances Bonferroni corrected"), color = "black", size = 10))
  print(Diversity.Multiplot.vio)
  dev.off()
  rm(Diversity.Multiplot.vio)
}

if (SaveWholeworkspace=='N') {rm(SOBS.plot.vio,Shan.plot.vio,Simp.plot.vio,Chao1.plot.vio)}
saveRDS(t.M.diversity.ord,file=file.path(PathToVisuaRAnalysis,"Alpha_Diversity",paste(VisuaRProjectName,"_DiversityIndices-t.rds",sep="")))

closeAllConnections() # closes all currently open connections.
gc()
# #======= 4.3. Species accumulation curves ====================================================================================================================================================

cat('\n\n4.3. Species accumulation curves',file=(file.path(PathToVisuaRAnalysis,paste(VisuaRProjectName,'_Analysis_log.txt',sep=''))),sep='',append=TRUE)

# #------- 4.3.1. Calculates Species accumulation curve using samples -----------------------------------------------------------------------------------------------------------------
#?Species richness not shown in plot (y acis label)
cat('\n4.3.1. Calculates Species accumulation curve using samples',file=(file.path(PathToVisuaRAnalysis,paste(VisuaRProjectName,'_Analysis_log.txt',sep=''))),sep='',append=TRUE)

M.spec=specaccum(M, method="random", permutations=NI, conditioned=T, gamma="Chao")
pdf(file.path(PathToVisuaRAnalysis,"Alpha_Diversity","Diversity_Metrics",paste(VisuaRProjectName,"_Species_Accumulation.pdf",sep = '')),useDingbats=F)
par(mar=c(5,2,5,7), xpd=TRUE)                                                                             # mar: margin sizes in the following order: bottom, left, top, and right.
plot(M.spec,ci.type="poly", col="grey", lwd=3, ci.lty=0, ci.col=alpha('grey',alpha=0.5),main=c(VisuaRProjectName,'Species Accumulation_Total'),ylab='Species richness',xlab='Samples')
boxplot(M.spec,col="grey", add=TRUE, outpch=19,outcex=0.35,boxwex=0.25,medlwd=1.8,whisklty=1)             # medlwd: width of the median line, boxwex:a scale factor applied to all boxes (to make them narrower or wider), whisklty=1 sets whisker line type to line (not dashed as default), outcex: size of outlier points, outpch=19: outlier points are filled dots
par(new=T)
plot.new()
legend(x=1.05,y=1.0,legend=c('All samples'),fill=c('grey'),bty ='n',cex=0.75,text.font = 2)
dev.off()

# #------- 4.3.2. Calculate Species accumulation curve using Grouping1 -----------------------------------------------------------------------------------------------------------------------
if (all(as.integer(M.group.count[,2])>1)){
  cat('\n4.3.2. Calculate Species accumulation curve using ',Grouping1,' (Grouping1).',file=(file.path(PathToVisuaRAnalysis,paste(VisuaRProjectName,'_Analysis_log.txt',sep=''))),sep='',append=TRUE)
  
  M.plus.contextdata=cbind(as.data.frame(M),M.contextdata.subset.noNA[,Grouping1]) # adds Grouping1 column to M
  
  pdf(file.path(PathToVisuaRAnalysis,"Alpha_Diversity","Diversity_Metrics",paste(VisuaRProjectName,"_Species_Accumulation_Groups.pdf",sep='')),useDingbats=F)
  par(mar=c(5,2,5,7), xpd=TRUE) 
  plot(M.spec,ci.type="poly", col='grey', lwd=3, ci.lty=0, ci.col=alpha('grey',alpha=0.2),main=c(VisuaRProjectName,'Species Accumulation_Total_and_Grouped'),ylab='Species richness',xlab='Samples')  # The accumulation curve by sample is plotted first
  for (i in 1:length(M.projects.unique)) {                                                                                                                                                            # adds accumulation curves by Group as in Grouping1 to the above curve
    SubsetDataframe=M.plus.contextdata[grep(M.projects.unique.ord[i],M.plus.contextdata[,ncol(M.plus.contextdata)],invert=F),]                                                                                 # only takes samples belonging to the first grouping (alphabetically)
    SubsetDataframe=SubsetDataframe[,-ncol(SubsetDataframe)]                                                                                                                                          # deletes the column with the Grouping1 information
    loopvariable=paste('Specaccum.Group.',M.projects.unique.ord[i],sep='')                                                                                                                            # creates a new 'loopvariable' for each round
    to.specaccum.plot=assign(loopvariable,specaccum(SubsetDataframe, method="random", permutations=NI, conditioned=T, gamma="Chao"))                                                                  # assign: assigns the value calculated in specaccum (the respective species accumulation of the group calculating at this moment) to the variable saved in loopvariable (not to loopvariable itself). In addition the result is saved in the variable 'to.specaccum.plot'. 
    plot(to.specaccum.plot,ci.type="poly", col=M.col[i], lwd=3, ci.lty=0, ci.col=alpha(M.col[i],alpha=0.25),add=T)                                                                                    # adds the specaccum plot of each group to the plot
    rm(to.specaccum.plot)
  }
  par(new=T)
  plot.new()
  legend(x=1.05,y=1.0,legend=c('All',M.projects.unique.ord),fill=c('grey',M.col),bty ='n',cex=0.75,text.font = 2) # adds a legend to the above created plot
  dev.off()
  
  if (SaveWholeworkspace=='N') {rm(M.spec,M.plus.contextdata,SubsetDataframe,loopvariable)}
  
  closeAllConnections() # closes all currently open connections.
  gc()
}
# #------- 4.4. Calculate Indicator Species -----------------------------------------------------------------------------------------------------------------------

IndicatorPhylum.exp.list <- vector("list",length(tax.levels)); names(IndicatorPhylum.exp.list) <- paste0("Indicator",tax.levels) #tax.levels.tables.indicators <- paste0("Indicator",tax.levels)


if (contextdata=='Y' & CalculateIndicators=='Y' & length(M.projects.unique)>1) {
  cat('\n4.4. Calculate Indicator Species for the provided ',Grouping1,' (Grouping1).',file=(file.path(PathToVisuaRAnalysis,paste(VisuaRProjectName,'_Analysis_log.txt',sep=''))),sep='',append=TRUE)
  for (k in 1:length(tax.levels)) {
    if (dim(tax.levels.tables.indicators.list[[k]])[2] > 1) {
      tax.levels.tables.indicators.list[[k]] <- indicspecies::multipatt(tax.levels.tables.indicators.list[[k]],cluster=M.projects.ord,func='r.g',control=how(nperm=9999)) # will only return statistically significant species (p<0.05). Set alpha=1 to see all species
      IndicatorPhylum.exp.list[[k]]=tax.levels.tables.indicators.list[[k]]$sign
      IndicatorPhylum.exp.list[[k]]=IndicatorPhylum.exp.list[[k]][order(IndicatorPhylum.exp.list[[k]]$index,-IndicatorPhylum.exp.list[[k]]$stat,IndicatorPhylum.exp.list[[k]]$p.value),]
    }
  }
  indicators <- createWorkbook()
  for (k in 1:length(tax.levels)) {
    addWorksheet(indicators,as.character(tax.levels[[k]]));writeData(indicators,as.character(tax.levels[[k]]),IndicatorPhylum.exp.list[[k]],row.names = T)
  }
  saveWorkbook(indicators,file=file.path(PathToVisuaRAnalysis,'Alpha_Diversity',paste(VisuaRProjectName,"_IndicatorSpecies.xlsx",sep="")))
  
  if (SaveWholeworkspace=='N') {rm(indicators,tax.levels.tables.indicators.list,M.seq.tax.subset.ord,IndicatorPhylum.exp.list)}
  gc()
}

save.image(file.path(PathToVisuaRAnalysis,paste(VisuaRProjectName,'_VisuaR','.RData',sep='')))

# #================== 5. Beta Diversity =================================================================================================================================================

cat('\n\n5. Beta Diversity',file=(file.path(PathToVisuaRAnalysis,paste(VisuaRProjectName,'_Analysis_log.txt',sep=''))),sep='',append=TRUE)

M.grouped=rowsum(M, group=M.groups, reorder = TRUE)   # sums up ASV abundances of a condition using Grouping1 (columns:ASVs, rows:groups,ordered alphabetically/increasing, fill: ASV reads)
saveRDS(M.grouped,file.path(PathToVisuaRAnalysis,'Beta_Diversity',paste(VisuaRProjectName,"_Mgrouped.rds",sep="")))
M.grouped.pa=decostand(M.grouped, method='pa')        # creates presence absence table of the Group  by ASV table


# #========== 5.1. Anaylsis of Similarities (ANOSIM) and calculation of Non-metric Multi-Dimensional Scaling (NMDS) ordinations. =====================================================

cat('\n5.1.  Anaylsis of Similarities (ANOSIM) and calculation of Non-metric Multi-Dimensional Scaling (NMDS) ordinations.',file=(file.path(PathToVisuaRAnalysis,paste(VisuaRProjectName,'_Analysis_log.txt',sep=''))),sep='',append=TRUE)

NMDSPlotsPath=file.path(PathToVisuaRAnalysis,"Beta_Diversity","NMDSPlots")
dir.create(file.path(PathToVisuaRAnalysis,"Beta_Diversity","NMDSPlots"))

# #------- 5.1.1. ANOSIM ----------------------------------------------------------------------------------------------------------------------------------------------------------------
if(length(M.projects.unique)>1){
  M.ano=anosim(M.rel, permutations=999, grouping=M.groups, distance="bray")                                                           # Anosim calculation of different groups . M.rel is a sample by ASV table filled with the relative abundances. The samples are not in the order of the sorted Grouping1. M.groups is in the same 'unordered' format
  pdf(file.path(PathToVisuaRAnalysis,'Beta_Diversity',paste(VisuaRProjectName,"_Anosim.pdf")),useDingbats=F)
  plot(M.ano,col=c("black",M.col),xlab=NULL,ylab='Dissimilarity Rank',main=paste('Anosim\n',VisuaRProjectName),cex.main=0.8,xaxt='n') # plots the result of ANOSIM
  axis(1, at=1:(length(M.projects.unique)+1),labels=c('Between',M.projects.unique.ord))                                               # renames the axis with the Grouping1 names
  dev.off()
  cat('\n5.1.1. Analysis of Similarities (ANOSIM)',file=(file.path(PathToVisuaRAnalysis,paste(VisuaRProjectName,'_Analysis_log.txt',sep=''))),sep='',append=TRUE)
  cat("\nAnosim_Dissimilarity: ",M.ano$dissimilarity,file=(file.path(PathToVisuaRAnalysis,paste(VisuaRProjectName,'_Analysis_log.txt',sep=''))),sep='',append=TRUE)
  cat("\nAnosim_statistic_R: ",M.ano$statistic,file=(file.path(PathToVisuaRAnalysis,paste(VisuaRProjectName,'_Analysis_log.txt',sep=''))),sep='',append=TRUE)
  cat("\nAnosim_Significance: ",M.ano$signif,file=(file.path(PathToVisuaRAnalysis,paste(VisuaRProjectName,'_Analysis_log.txt',sep=''))),sep='',append=TRUE)
  cat("\nAnosim_Permutations: ",M.ano$permutations,file=(file.path(PathToVisuaRAnalysis,paste(VisuaRProjectName,'_Analysis_log.txt',sep=''))),sep='',append=TRUE)
}

if(length(M.projects.unique)>1){
  # # write Anosim output in seperate file
  summaryAno <- as.data.frame(capture.output(summary(M.ano)))
  summaryAno <- summaryAno[17:(nrow(summaryAno)-1),]
  split_data <- lapply(summaryAno, function(x) unlist(strsplit(x, "\\s+")))
  summary_df <- as.data.frame(do.call(rbind, split_data), stringsAsFactors = FALSE)
  colnames(summary_df) <- summary_df[1,]
  summary_df[3:nrow(summary_df),1] <- M.projects.unique.ord
  rownames(summary_df) <- summary_df[,1]
  summary_df <- summary_df[-1,-1]
  data.table::fwrite(summary_df,file=file.path(PathToVisuaRAnalysis,'Beta_Diversity',paste(VisuaRProjectName,"_Anosim.csv",sep="")),col.names = T,row.names = T)
  rm(summary_df,summaryAno)
}

# paired.permanova <- vegan::adonis2(M.dist~M.projects,data=M.contextdata.subset.noNA,strata = M.contextdata.subset.noNA$VisuaR_samplegroups)


if (SaveWholeworkspace=='N') {rm(M.rel)}
gc()

cat('\n\n5.1.2. NMDS calculation and Visualization',file=(file.path(PathToVisuaRAnalysis,paste(VisuaRProjectName,'_Analysis_log.txt',sep=''))),sep='',append=TRUE)
cat('\nNote: Using metaMDS with a distance matrix is the same as using monoMDS without a distance matrix. You create a distance matrix using vegdist(method=bray) and apply metaMDS to this distrance matrix.' ,file=(file.path(PathToVisuaRAnalysis,paste(VisuaRProjectName,'_Analysis_log.txt',sep=''))),sep='',append=TRUE)

M.dist=vegdist(M, method="bray") # Calculates disimilarity indices. Samples in M are not ordered based on Grouping1

# # Community Similarity in %
Max.Com.Sim=(1-(min(M.dist)))*100 
Min.Com.Sim=(1-(max(M.dist)))*100
Mean.Com.Sim=(1-(mean(M.dist)))*100
cat('\nNote: Maximum, Mean and Minimal Community Similarity in % (as shown in the NMDS plots) refers to the comparison of any two samples using the bray-curtis distance matrix.' ,file=(file.path(PathToVisuaRAnalysis,paste(VisuaRProjectName,'_Analysis_log.txt',sep=''))),sep='',append=TRUE)

M.mMDS=metaMDS(M.dist) # takes distances from n-dimensional M.dist (n=number of samples-1) and creates a 2-dimensional ordination (by default). 
# # metaMDS: if provided with a distance matrix (our case) it calls monoMDS()
# # monoMDS() generates a starting configuration (PCoA) repeatedly (max 20 default) each time with a new random starting configuration, evaluates stress and adjusts the points
# # also internally does procrustes to see if configurations have converged and then configuration with best stress level chosen. 
# # Two final configurations are considered to have converged (arrived at essentially the same solution) when the rmse (root mean square error) is less than 0.01, and no single residual value exceeds 0.005. (https://www.flutterbys.com.au/stats/tut/tut15.1.html)
# # M.mMDS$points points in the ordination space


saveRDS(M.dist,file.path(PathToVisuaRAnalysis,'Beta_Diversity',paste(VisuaRProjectName,"_distanceMatrix.rds",sep="")))
saveRDS(M.mMDS,file.path(PathToVisuaRAnalysis,'Beta_Diversity',paste(VisuaRProjectName,"_distanceMatrix2d.rds",sep="")))
cat("\nNote: In NMDS ordinations the distance between two samples represents the distance of their underlying communities.\nNMDS distances are relative measures and thus, do not need an axis.\nAxes in NMDS ordinations are only used when NMDS is combined (i.e. superimposed on the NMDS ordination) with another method.",file=(file.path(PathToVisuaRAnalysis,paste(VisuaRProjectName,'_Analysis_log.txt',sep=''))),sep="\t",append=TRUE)

M.mMDS.stress=M.mMDS$stress 
cat('\nNMDS_Stress_Level: ',M.mMDS.stress,file=(file.path(PathToVisuaRAnalysis,paste(VisuaRProjectName,'_Analysis_log.txt',sep=''))),sep='',append=TRUE)
cat('\nNote: Stress values are a measure for the goodness of the NMDS (i.e. how realistic is the n-dimensional space represented in the 2-dimensional plot).',file=(file.path(PathToVisuaRAnalysis,paste(VisuaRProjectName,'_Analysis_log.txt',sep=''))),sep="\t",append=TRUE)
cat('\nNote: R and p values shown in the NMDS plots are calculated using ANOSIM. Please consider limitations and interpretation of the ANOSIM test.',file=(file.path(PathToVisuaRAnalysis,paste(VisuaRProjectName,'_Analysis_log.txt',sep=''))),sep="\t",append=TRUE)
cat('\nNote: NMDS featuring text of the samples is meant for reference and to look up specific samples yet is not optimized for display.',file=(file.path(PathToVisuaRAnalysis,paste(VisuaRProjectName,'_Analysis_log.txt',sep=''))),sep="\t",append=TRUE)

# # NMDS featuring sample names 
pdf(file.path(NMDSPlotsPath,paste(VisuaRProjectName,"_NMDS_Text.pdf",sep="")),height=20,width=20,useDingbats=F)
par(mar=c(3,3,3,3),xpd=T)
ordiplot(M.mMDS, type="t", display="sites",ylab="",xlab="",xaxt='n',yaxt='n')  # type='t': shows the sites as text, display='sites: shows samples, y and x axis removed
points (M.mMDS, col=M.colvec, pch=19, cex=1)                                 # cex= the point size of each sample represents Shannon Entropy.
dev.off()

# # NMDS featuring Groups
pdf(file.path(NMDSPlotsPath,paste(VisuaRProjectName,"_NMDS_dots.pdf",sep="")),height=5,width=6,useDingbats=F)
par(mar=c(2, 2, 2, 7), xpd=T)                                                                                     # mar: margin sizes in the following order: bottom, left, top, and right. There is more space at the right hand site for the figure legend, xpd='T': all plotting is clipped to the figure region (not only to the plot region). This allows to place figure legens outside of the plot
ordiplot(M.mMDS, type='n', display="sites",ylab='',xlab='',xaxt='n',yaxt='n')                                     # type='n': sites are invisible and can be filled with points command
points (M.mMDS, col=M.colvec, pch=19, cex=1)                                 # cex= the point size of each sample represents Shannon Entropy.
par(new=T)                                                                                                        # only using par(new=T) it is possible to plot the figure legend independent of the axis scaling to a fixed place inside the pdf.
plot.new()
if(length(M.projects.unique)>1){                                                                                  # only if more than one grouping
  legend(x=1.01,y=1.05,legend='Groups',bty='n',cex=0.75,text.font = 2)                                            # text.font=2: prints text in bold, bty='n': no box will be drawn around the legend. Alternative: bty='o'
  legend(1.045,1.00,legend=c(M.projects.unique.ord),fill=c(M.col),bty='n',cex=0.75)
  legend(1.01,0.46,legend=paste0("Community\nSimilarity [%]"),bty='n',cex=0.75,text.font = 2)
  legend(1.01,0.4,legend=paste0('max: ',round(Max.Com.Sim,1),'\nmean: ',round(Mean.Com.Sim,1),'\nmin: ',round(Min.Com.Sim,1)),bty='n',cex=0.75)
  legend(1.01,0.21,legend=paste0('ANOSIM'),bty='n',cex=0.75,text.font = 2)
  legend(1.01,0.17,legend=paste0('R: ',round(M.ano$statistic,3),'\np: ',M.ano$signif),bty='n',cex=0.75)
  legend(1.01,0.03,legend=paste0("Stress: ",round(M.mMDS.stress,3)),bty='n',cex=0.75)                             # cex: changes the size of the Stress value shown in the plot
} else {
  legend(1.01,0.46,legend=paste0("Community\nSimilarity [%]"),bty='n',cex=0.75,text.font = 2)
  legend(1.01,0.4,legend=paste0('max: ',round(Max.Com.Sim,1),'\nmean: ',round(Mean.Com.Sim,1),'\nmin: ',round(Min.Com.Sim,1)),bty='n',cex=0.75)
  legend(1.01,0.03,legend=paste0("Stress: ",round(M.mMDS.stress,3)),bty='n',cex=0.75) 
}
dev.off()

# # NMDS featuring Shannon Entropy
pdf(file.path(NMDSPlotsPath,paste(VisuaRProjectName,"_NMDS_Shannon.pdf",sep="")),height=5,width=6,useDingbats=F)
par(mar=c(2, 2, 2, 7), xpd=TRUE)
ordiplot (M.mMDS, display = 'si', type = 'n',ylab='',xlab='',xaxt='n',yaxt='n')
points (M.mMDS, col=M.colvec, pch=19, cex=M.shan.r.mean/(max(M.shan.r.mean)/2.7))                                 # cex= the point size of each sample represents Shannon Entropy.
par(new=T)                                                                                                        # cex sizes are scaled to fit the plot dimensions using a scaling factor based on the maximum average Shannon Entropy and a constant (2.7)
plot.new()                                                                                                        # circle sizes are meant to visualize relative differences in Shannon. Not absolute values. 
if(length(M.projects.unique)>1){ 
  legend(x=1.01,y=1.05,legend='Groups',bty='n',cex=0.75,text.font = 2) 
  legend(1.045,1.00,legend=c(M.projects.unique.ord),fill=c(M.col),bty='n',cex=0.75)
  legend(1.01,0.58,legend='Dotsize',bty='n',cex = 0.75,text.font = 2)
  legend(1.01,0.535,legend='Shannon Entropy',bty='n',cex = 0.75)
  legend(1.01,0.46,legend=paste0("Community\nSimilarity [%]"),bty='n',cex=0.75,text.font = 2)
  legend(1.01,0.4,legend=paste0('max: ',round(Max.Com.Sim,1),'\nmean: ',round(Mean.Com.Sim,1),'\nmin: ',round(Min.Com.Sim,1)),bty='n',cex=0.75)
  legend(1.01,0.21,legend=paste0('ANOSIM'),bty='n',cex=0.75,text.font = 2)
  legend(1.01,0.17,legend=paste0('R: ',round(M.ano$statistic,3),'\np: ',M.ano$signif),bty='n',cex=0.75)
  legend(1.01,0.03,legend=paste0("Stress: ",round(M.mMDS.stress,3)),bty='n',cex=0.75)
} else {
  legend(1.01,0.58,legend='Dotsize',bty='n',cex = 0.75,text.font = 2)
  legend(1.01,0.535,legend='Shannon Entropy',bty='n',cex = 0.75)
  legend(1.01,0.46,legend=paste0("Community\nSimilarity [%]"),bty='n',cex=0.75,text.font = 2)
  legend(1.01,0.4,legend=paste0('max: ',round(Max.Com.Sim,1),'\nmean: ',round(Mean.Com.Sim,1),'\nmin: ',round(Min.Com.Sim,1)),bty='n',cex=0.75)
  legend(1.01,0.03,legend=paste0("Stress: ",round(M.mMDS.stress,3)),bty='n',cex=0.75) 
}
dev.off()

if (SaveWholeworkspace=='N') {rm(M.shan.r.mean)}
gc()

# # NMDS featuring ASV Richness
pdf(file.path(NMDSPlotsPath,paste(VisuaRProjectName,"_NMDS_Richness.pdf",sep="")),height=5,width=6,useDingbats=F)
par(mar=c(2, 2, 2, 7), xpd=TRUE)
ordiplot (M.mMDS, display = 'si', type = 'n',ylab='',xlab='',xaxt='n',yaxt='n')
points (M.mMDS, col=M.colvec, pch=19, cex=M.sobs.r.mean/(max(M.sobs.r.mean)/2.7)) 
par(new=T)
plot.new()
if(length(M.projects.unique)>1){ # only if more than one grouping
  legend(x=1.01,y=1.05,legend='Groups',bty='n',cex=0.75,text.font = 2)
  legend(1.045,1.00,legend=c(M.projects.unique.ord),fill=c(M.col),bty='n',cex=0.75)
  legend(1.01,0.58,legend='Dotsize',bty='n',cex = 0.75,text.font = 2)
  legend(1.01,0.535,legend='ASV Richness',bty='n',cex = 0.75)
  legend(1.01,0.46,legend=paste0("Community\nSimilarity [%]"),bty='n',cex=0.75,text.font = 2)
  legend(1.01,0.4,legend=paste0('max: ',round(Max.Com.Sim,1),'\nmean: ',round(Mean.Com.Sim,1),'\nmin: ',round(Min.Com.Sim,1)),bty='n',cex=0.75)
  legend(1.01,0.21,legend=paste0('ANOSIM'),bty='n',cex=0.75,text.font = 2)
  legend(1.01,0.17,legend=paste0('R: ',round(M.ano$statistic,3),'\np: ',M.ano$signif),bty='n',cex=0.75)
  legend(1.01,0.03,legend=paste0("Stress: ",round(M.mMDS.stress,3)),bty='n',cex=0.75)
} else {
  legend(1.01,0.58,legend='Dotsize',bty='n',cex = 0.75,text.font = 2)
  legend(1.01,0.535,legend='ASV Richness',bty='n',cex = 0.75)
  legend(1.01,0.46,legend=paste0("Community\nSimilarity [%]"),bty='n',cex=0.75,text.font = 2)
  legend(1.01,0.4,legend=paste0('max: ',round(Max.Com.Sim,1),'\nmean: ',round(Mean.Com.Sim,1),'\nmin: ',round(Min.Com.Sim,1)),bty='n',cex=0.75)
  legend(1.01,0.03,legend=paste0("Stress: ",round(M.mMDS.stress,3)),bty='n',cex=0.75) 
}
dev.off()

gc()

# # NMDS featuring Inverse Simpson Diversity
pdf(file.path(NMDSPlotsPath,paste(VisuaRProjectName,"_NMDS_InvSimpson.pdf",sep="")),height=5,width=6,useDingbats=F)
par(mar=c(2, 2, 2, 7), xpd=TRUE)
ordiplot (M.mMDS, display = 'si', type = 'n',ylab="",xlab="",xaxt='n',yaxt='n')
points (M.mMDS, col=M.colvec, pch=19, cex=M.invs.r.mean/(max(M.invs.r.mean)/3.5)) 
par(new=T)
plot.new()
if(length(M.projects.unique)>1){ 
  legend(x=1.01,y=1.05,legend='Groups',bty='n',cex=0.75,text.font = 2) 
  legend(1.045,1.00,legend=c(M.projects.unique.ord),fill=c(M.col),bty='n',cex=0.75)
  legend(1.01,0.58,legend='Dotsize',bty='n',cex = 0.75,text.font = 2)
  legend(1.01,0.535,legend='Inverse Simpson',bty='n',cex = 0.75)
  legend(1.01,0.46,legend=paste0("Community\nSimilarity [%]"),bty='n',cex=0.75,text.font = 2)
  legend(1.01,0.4,legend=paste0('max: ',round(Max.Com.Sim,1),'\nmean: ',round(Mean.Com.Sim,1),'\nmin: ',round(Min.Com.Sim,1)),bty='n',cex=0.75)
  legend(1.01,0.21,legend=paste0('ANOSIM'),bty='n',cex=0.75,text.font = 2)
  legend(1.01,0.17,legend=paste0('R: ',round(M.ano$statistic,3),'\np: ',M.ano$signif),bty='n',cex=0.75)
  legend(1.01,0.03,legend=paste0("Stress: ",round(M.mMDS.stress,3)),bty='n',cex=0.75) 
} else {
  legend(1.01,0.58,legend='Dotsize',bty='n',cex = 0.75,text.font = 2)
  legend(1.01,0.535,legend='Inverse Simpson',bty='n',cex = 0.75)
  legend(1.01,0.46,legend=paste0("Community\nSimilarity [%]"),bty='n',cex=0.75,text.font = 2)
  legend(1.01,0.4,legend=paste0('max: ',round(Max.Com.Sim,1),'\nmean: ',round(Mean.Com.Sim,1),'\nmin: ',round(Min.Com.Sim,1)),bty='n',cex=0.75)
  legend(1.01,0.03,legend=paste0("Stress: ",round(M.mMDS.stress,3)),bty='n',cex=0.75) 
}
dev.off()

if (SaveWholeworkspace=='N') {rm(M.invs.r.mean)}
gc()

# # NMDS featuring Chao1 Richness 
pdf(file.path(NMDSPlotsPath,paste(VisuaRProjectName,"_NMDS_Chao1.pdf",sep="")),height=5,width=6,useDingbats=F)
par(mar=c(2, 2, 2, 7), xpd=TRUE) 
ordiplot (M.mMDS, display = 'si', type = 'n',ylab="",xlab="",xaxt='n',yaxt='n')
points (M.mMDS, col=M.colvec, pch=19, cex=M.chao1.r.mean/(max(M.chao1.r.mean)/3.5))
par(new=T)
plot.new()
if(length(M.projects.unique)>1){ 
  legend(x=1.01,y=1.05,legend='Groups',bty='n',cex=0.75,text.font = 2) 
  legend(1.045,1.00,legend=c(M.projects.unique.ord),fill=c(M.col),bty='n',cex=0.75)
  legend(1.01,0.58,legend='Dotsize',bty='n',cex = 0.75,text.font = 2)
  legend(1.01,0.535,legend='Chao1 Richness',bty='n',cex = 0.75)
  legend(1.01,0.46,legend=paste0("Community\nSimilarity [%]"),bty='n',cex=0.75,text.font = 2)
  legend(1.01,0.4,legend=paste0('max: ',round(Max.Com.Sim,1),'\nmean: ',round(Mean.Com.Sim,1),'\nmin: ',round(Min.Com.Sim,1)),bty='n',cex=0.75)
  legend(1.01,0.21,legend=paste0('ANOSIM'),bty='n',cex=0.75,text.font = 2)
  legend(1.01,0.17,legend=paste0('R: ',round(M.ano$statistic,3),'\np: ',M.ano$signif),bty='n',cex=0.75)
  legend(1.01,0.03,legend=paste0("Stress: ",round(M.mMDS.stress,3)),bty='n',cex=0.75) 
} else {
  legend(1.01,0.58,legend='Dotsize',bty='n',cex = 0.75,text.font = 2)
  legend(1.01,0.535,legend='Chao1 Richness',bty='n',cex = 0.75)
  legend(1.01,0.46,legend=paste0("Community\nSimilarity [%]"),bty='n',cex=0.75,text.font = 2)
  legend(1.01,0.4,legend=paste0('max: ',round(Max.Com.Sim,1),'\nmean: ',round(Mean.Com.Sim,1),'\nmin: ',round(Min.Com.Sim,1)),bty='n',cex=0.75)
  legend(1.01,0.03,legend=paste0("Stress: ",round(M.mMDS.stress,3)),bty='n',cex=0.75) 
}
dev.off()

if (SaveWholeworkspace=='N') {rm(M.chao1.r,M.chao1.r.mean)}
gc()

# # NMDS featuring Ordiellipse&Ordispider 
cat('\nNote: Ordispider: each Sample is connected to the weighted average mean of the within group distances (centroid).',file=(file.path(PathToVisuaRAnalysis,paste(VisuaRProjectName,'_Analysis_log.txt',sep=''))),sep="\t",append=TRUE)
cat('\nNote: Ordiellipse: visualizes one standard deviation from the groups centroids. ',file=(file.path(PathToVisuaRAnalysis,paste(VisuaRProjectName,'_Analysis_log.txt',sep=''))),sep="\t",append=TRUE)

pdf(file.path(NMDSPlotsPath,paste(VisuaRProjectName,"_NMDS_spider_ellipse.pdf",sep="")),height=5,width=6,useDingbats=F)
par(mar=c(2, 2, 2, 7), xpd=F)                                                                                             # first, xpd=F because if not the ellipses might be drawn outside of the plot region
ordiplot (M.mMDS, display = 'si', type = 'n',ylab="",xlab="",xaxt='n',yaxt='n')
for (i in seq (1, NR)) ordiellipse (M.mMDS, groups = M.groups, show.groups = i, col = M.col[i], label = F,lwd=1.5,draw='polygon',border='NA',alpha=60,kind='sd')
for (i in seq (1, NR)) ordispider (M.mMDS, groups = M.groups, show.groups = i, col = M.col[i], label = F,lwd=1.5)
points (M.mMDS, col=M.colvec, pch=19,cex=1) 
par(new=T,xpd=T)
plot.new()
if(length(M.projects.unique)>1){
  legend(x=1.01,y=1.05,legend='Groups',bty='n',cex=0.75,text.font = 2) 
  legend(1.045,1.00,legend=c(M.projects.unique.ord),fill=c(M.col),bty='n',cex=0.75)
  legend(1.01,0.46,legend=paste0("Community\nSimilarity [%]"),bty='n',cex=0.75,text.font = 2)
  legend(1.01,0.4,legend=paste0('max: ',round(Max.Com.Sim,1),'\nmean: ',round(Mean.Com.Sim,1),'\nmin: ',round(Min.Com.Sim,1)),bty='n',cex=0.75)
  legend(1.01,0.21,legend=paste0('ANOSIM'),bty='n',cex=0.75,text.font = 2)
  legend(1.01,0.17,legend=paste0('R: ',round(M.ano$statistic,3),'\np: ',M.ano$signif),bty='n',cex=0.75)
  legend(1.01,0.03,legend=paste0("Stress: ",round(M.mMDS.stress,3)),bty='n',cex=0.75) 
} else {
  legend(1.01,0.46,legend=paste0("Community\nSimilarity [%]"),bty='n',cex=0.75,text.font = 2)
  legend(1.01,0.4,legend=paste0('max: ',round(Max.Com.Sim,1),'\nmean: ',round(Mean.Com.Sim,1),'\nmin: ',round(Min.Com.Sim,1)),bty='n',cex=0.75)
  legend(1.01,0.03,legend=paste0("Stress: ",round(M.mMDS.stress,3)),bty='n',cex=0.75) 
}
dev.off()


# # NMDS featuring ASV Richness & Ordiellipse & Ordispider
pdf(file.path(NMDSPlotsPath,paste(VisuaRProjectName,"_NMDS_Richness_ellipse2.pdf",sep="")),height=5,width=6,useDingbats=F)
par(mar=c(2, 2, 2, 7), xpd=F)
vegan::ordiplot (M.mMDS, display = 'si', type = 'n',ylab='',xlab='',xaxt='n',yaxt='n')
for (i in seq (1, NR)) vegan::ordiellipse (M.mMDS, groups = M.groups, show.groups = i, col = M.col[i], label = F,lwd=1.5,draw='polygon',border='NA',alpha=100,kind='sd')
for (i in seq (1, NR)) vegan::ordispider (M.mMDS, groups = M.groups, show.groups = i, col = M.col[i], label = F,lwd=0.5)
points (M.mMDS, col=M.colvec, pch=19, cex=M.sobs.r.mean/(max(M.sobs.r.mean)/2.7)) 
par(new=T,xpd=T)
plot.new()
if(length(M.projects.unique)>1){ # only if more than one grouping
  legend(x=1.01,y=1.05,legend='Groups',bty='n',cex=0.75,text.font = 2)
  legend(1.045,1.00,legend=c(M.projects.unique.ord),fill=c(M.col),bty='n',cex=0.75)
  legend(1.01,0.58,legend='Dotsize',bty='n',cex = 0.75,text.font = 2)
  legend(1.01,0.535,legend='ASV Richness',bty='n',cex = 0.75)
  legend(1.01,0.46,legend=paste0("Community\nSimilarity [%]"),bty='n',cex=0.75,text.font = 2)
  legend(1.01,0.4,legend=paste0('max: ',round(Max.Com.Sim,1),'\nmean: ',round(Mean.Com.Sim,1),'\nmin: ',round(Min.Com.Sim,1)),bty='n',cex=0.75)
  legend(1.01,0.21,legend=paste0('ANOSIM'),bty='n',cex=0.75,text.font = 2)
  legend(1.01,0.17,legend=paste0('R: ',round(M.ano$statistic,3),'\np: ',M.ano$signif),bty='n',cex=0.75)
  legend(1.01,0.03,legend=paste0("Stress: ",round(M.mMDS.stress,3)),bty='n',cex=0.75)
} else {
  legend(1.01,0.58,legend='Dotsize',bty='n',cex = 0.75,text.font = 2)
  legend(1.01,0.535,legend='ASV Richness',bty='n',cex = 0.75)
  legend(1.01,0.46,legend=paste0("Community\nSimilarity [%]"),bty='n',cex=0.75,text.font = 2)
  legend(1.01,0.4,legend=paste0('max: ',round(Max.Com.Sim,1),'\nmean: ',round(Mean.Com.Sim,1),'\nmin: ',round(Min.Com.Sim,1)),bty='n',cex=0.75)
  legend(1.01,0.03,legend=paste0("Stress: ",round(M.mMDS.stress,3)),bty='n',cex=0.75) 
}
dev.off()

if (SaveWholeworkspace=='N') {rm(M.sobs.r.mean)}
gc()


# # NMDS featuring Ordihull 
pdf(file.path(NMDSPlotsPath,paste(VisuaRProjectName,"_NMDS_ellipse.pdf",sep="")),height=5,width=6,useDingbats=F)
par(mar=c(2, 2, 2, 7), xpd=F)                                                                                             # first, xpd=F because if not the ellipses might be drawn outside of the plot region
ordiplot (M.mMDS, display = 'si', type = 'n',ylab="",xlab="",xaxt='n',yaxt='n')
for (i in seq (1, NR)) ordiellipse (M.mMDS, groups = M.groups, show.groups = i, col = M.col[i], label = F,lwd=1.5,draw='polygon',border='NA',alpha=60,kind='sd')
points (M.mMDS, col=M.colvec, pch=19,cex=0.5) 
# for (i in seq (1, NR)) ordispider (M.mMDS, groups = M.groups, show.groups = i, col = M.col[i], label = F,lwd=1.5)
# points (M.mMDS, col=M.colvec, pch=19,cex=1) 
par(new=T,xpd=T)
plot.new()
if(length(M.projects.unique)>1){
  legend(x=1.01,y=1.05,legend='Groups',bty='n',cex=0.75,text.font = 2) 
  legend(1.045,1.00,legend=c(M.projects.unique.ord),fill=c(M.col),bty='n',cex=0.75)
  legend(1.01,0.46,legend=paste0("Community\nSimilarity [%]"),bty='n',cex=0.75,text.font = 2)
  legend(1.01,0.4,legend=paste0('max: ',round(Max.Com.Sim,1),'\nmean: ',round(Mean.Com.Sim,1),'\nmin: ',round(Min.Com.Sim,1)),bty='n',cex=0.75)
  legend(1.01,0.21,legend=paste0('ANOSIM'),bty='n',cex=0.75,text.font = 2)
  legend(1.01,0.17,legend=paste0('R: ',round(M.ano$statistic,3),'\np: ',M.ano$signif),bty='n',cex=0.75)
  legend(1.01,0.03,legend=paste0("Stress: ",round(M.mMDS.stress,3)),bty='n',cex=0.75) 
} else {
  legend(1.01,0.46,legend=paste0("Community\nSimilarity [%]"),bty='n',cex=0.75,text.font = 2)
  legend(1.01,0.4,legend=paste0('max: ',round(Max.Com.Sim,1),'\nmean: ',round(Mean.Com.Sim,1),'\nmin: ',round(Min.Com.Sim,1)),bty='n',cex=0.75)
  legend(1.01,0.03,legend=paste0("Stress: ",round(M.mMDS.stress,3)),bty='n',cex=0.75) 
}
dev.off()

# # NMDS featuring Ordihull 
pdf(file.path(NMDSPlotsPath,paste(VisuaRProjectName,"_NMDS_Ordihull.pdf",sep="")),height=5,width=6,useDingbats=F)
par(mar=c(2, 2, 2, 7), xpd=TRUE) 
ordiplot (M.mMDS, display = 'si', type = 'n',ylab="",xlab="",xaxt='n',yaxt='n')
points (M.mMDS, col=M.colvec, pch=19,cex=0.5) 
for (i in seq (1, NR)) ordihull (M.mMDS, groups = M.groups, show.groups = i, col = M.col[i], label = F)
par(new=T)
plot.new()
if(length(M.projects.unique)>1){ 
  legend(x=1.01,y=1.05,legend='Groups',bty='n',cex=0.75,text.font = 2)
  legend(1.045,1.00,legend=c(M.projects.unique.ord),fill=c(M.col),bty='n',cex=0.75)
  legend(1.01,0.46,legend=paste0("Community\nSimilarity [%]"),bty='n',cex=0.75,text.font = 2)
  legend(1.01,0.4,legend=paste0('max: ',round(Max.Com.Sim,1),'\nmean: ',round(Mean.Com.Sim,1),'\nmin: ',round(Min.Com.Sim,1)),bty='n',cex=0.75)
  legend(1.01,0.21,legend=paste0('ANOSIM'),bty='n',cex=0.75,text.font = 2)
  legend(1.01,0.17,legend=paste0('R: ',round(M.ano$statistic,3),'\np: ',M.ano$signif),bty='n',cex=0.75)
  legend(1.01,0.03,legend=paste0("Stress: ",round(M.mMDS.stress,3)),bty='n',cex=0.75) 
} else {
  legend(1.01,0.46,legend=paste0("Community\nSimilarity [%]"),bty='n',cex=0.75,text.font = 2)
  legend(1.01,0.4,legend=paste0('max: ',round(Max.Com.Sim,1),'\nmean: ',round(Mean.Com.Sim,1),'\nmin: ',round(Min.Com.Sim,1)),bty='n',cex=0.75)
  legend(1.01,0.03,legend=paste0("Stress: ",round(M.mMDS.stress,3)),bty='n',cex=0.75) 
}
dev.off()

closeAllConnections() # closes all currently open connections.

gc()

# #------- 5.2. Calculate and visualize grouped rarefaction -----------------------------------------------------------------------------------------------------------------------------

# # slow 
if (calcGroupedRaref == "Y") {
  MRR=M.grouped #sample-wise: M; group-wise: M.grouped
  
  M.rar <- rrarefy(MRR,sample=M.minreads) # data frame with estimated species richness using random subsamples of size M.minreads
  
  pdf(file.path(PathToVisuaRAnalysis,"Alpha_Diversity","Diversity_Metrics",paste(VisuaRProjectName,"_RarefactionCurve.pdf",sep="")),height=5,width=6,useDingbats=F)
  par(mar=c(5, 5, 5, 7), xpd=F) 
  M.rac <- rarecurve(MRR,sample=M.minreads,step=1000,col=M.col,lwd=3,main=c(VisuaRProjectName,'Rarefaction_Grouped'),cex.main=0.8,label=F) # rarefaction curve for each group. yields richness of all samples at M.minreads
  par(new=T,xpd=T)
  plot.new()
  legend(x=1.05,y=1.05,legend=c(M.projects.unique.ord),fill=c(M.col),bty ='n',cex=0.75,text.font = 2) 
  dev.off()
  
  # M.ras <- rareslope(MRR,sample=M.minreads-1) # calculates the slope of rarecurve for each group
  
  if (SaveWholeworkspace=='N') {rm(MRR,M.rar,M.rac)}
  gc()
}

# #------- 5.3. Calculates and visualizes Cluster Dendrograms by Groups ------------------------------------------------------------------------------------------------------------------
#? noch bearbeiten, fan: namen nicht abchneiden
# # Default metric is bray curtis distance and average linkage hierarchical clustering.

# # fast
dir.create(file.path(PathToVisuaRAnalysis,'Beta_Diversity','Dendrograms'))

if(length(M.projects.unique)>2){  
  M.grouped.dist=vegdist(M.grouped, method="bray")                              # Calculates distance matrix
  M.grouped.clust <- hclust(M.grouped.dist, method="average")                   # hierarchical cluster analysis on a set of dissimilarities
  M.grouped.clust$labels=c(M.projects.unique.ord)                               # renames the groups (names are removed while clustering)
  
  pdf(file.path(PathToVisuaRAnalysis,'Beta_Diversity','Dendrograms',paste(VisuaRProjectName,"_ClusterDendrogram_grouped.pdf",sep="")),height=5,width=5,useDingbats=F)
  M.grouped.cd=plot(M.grouped.clust,main=paste('Cluster Dendrogram\n',VisuaRProjectName),cex.main=0.8,xlab =NA,ylab = 'Distance' ,sub=NA)
  dev.off()
  pdf(file.path(PathToVisuaRAnalysis,'Beta_Diversity','Dendrograms',paste(VisuaRProjectName,"_ClusterDendrogram2_grouped.pdf",sep="")),height=5,width=5,useDingbats=F)
  M.grouped.phylo=plot(as.phylo(M.grouped.clust),label.offset = 0.01,cex=0.6,tip.color = c(M.col))
  dev.off()
  pdf(file.path(PathToVisuaRAnalysis,'Beta_Diversity','Dendrograms',paste(VisuaRProjectName,"_Cluster_fan_grouped.pdf",sep="")),height=5,width=5,useDingbats=F)
  M.grouped.phylo=plot(as.phylo(M.grouped.clust),type='fan',cex=0.4,label.offset = 0.02,tip.color = c(M.col))
  dev.off()
  if (SaveWholeworkspace=='N') {rm(M.grouped.dist,M.grouped.clust,M.grouped.cd,M.grouped.phylo)}
  gc()
}

# if (SaveWholeworkspace=='N') {rm(M.grouped)}
gc()

# #------- 5.4. Calculates and visualizes Cluster Dendrograms by samples -----------------------------------------------------------------------------------------------------------------
#? Gr??e mit anzahl samploes skalieren, und au?en nichts abschneiden
# # Creates different cluster diagrams of all samples colored in the selected Grouping1
# # fast

M.sample.clust <- hclust(M.dist, method="average")

pdf(file.path(PathToVisuaRAnalysis,'Beta_Diversity','Dendrograms',paste(VisuaRProjectName,"_ClusterDendrogram_samples.pdf",sep="")),height=5,width=5,useDingbats=F)
M.sample.cd=plot(M.sample.clust,main=paste('Cluster Dendrogram\n',VisuaRProjectName),cex.main=0.8,xlab =NA,ylab = 'Distance' ,sub=NA,hang=-1)
dev.off()
pdf(file.path(PathToVisuaRAnalysis,'Beta_Diversity','Dendrograms',paste(VisuaRProjectName,"_ClusterDendrogram3_samples.pdf",sep="")),height=5,width=5,useDingbats=F)
M.sample.phylo=plot(as.phylo(M.sample.clust),no.margin = T,cex=0.6,tip.color = c(M.colvec))
dev.off()
pdf(file.path(PathToVisuaRAnalysis,'Beta_Diversity','Dendrograms',paste(VisuaRProjectName,"_Cluster_fan_samples.pdf",sep="")),height=6.5,width=6.5,useDingbats=F)
M.sample.phylo=plot(as.phylo(M.sample.clust),type='fan',no.margin = T,cex=0.4,tip.color = c(M.colvec))
dev.off()
pdf(file.path(PathToVisuaRAnalysis,'Beta_Diversity','Dendrograms',paste(VisuaRProjectName,"_Cluster_radial_samples.pdf",sep="")),height=6.5,width=6.5,useDingbats=F)
M.sample.phylo=plot(as.phylo(M.sample.clust),type='radial',no.margin = T,cex=0.6,tip.color = c(M.colvec))
dev.off()

if (SaveWholeworkspace=='N') {rm(M.dist,M.sample.phylo)
  gc()}

save.image(file.path(PathToVisuaRAnalysis,paste(VisuaRProjectName,'_VisuaR','.RData',sep='')))

# #------- 5.4. Calculates de novo clusters based on the Bray-Curtis distance matrix for 2 to 8 clusters -----------------------------------------------------------------------------------------------------------------

# fast 
cluster.matrix <- matrix(nrow=length(M.sample.clust$labels),ncol = 0)
rownames(cluster.matrix) <- M.sample.clust$labels

for (num in 2:8) {
  clusters <- as.data.frame(cutree(M.sample.clust, k = num))
  cluster.matrix <- cbind(cluster.matrix,clusters)
  colnames(cluster.matrix)[ncol(cluster.matrix)] <- paste(num,"Clusters",sep="")
}
cluster.matrix <- cluster.matrix[order(cluster.matrix[,ncol(cluster.matrix)]),]

data.table::fwrite(as.data.frame(cluster.matrix),file=file.path(PathToVisuaRAnalysis,'Beta_Diversity',paste(VisuaRProjectName,"_Clusters_new.csv",sep="")),col.names = T,row.names = T)

if (SaveWholeworkspace=='N') {rm(M.sample.clust,cluster.matrix,clusters)
  gc()}


# # #------- 5.5. Create UpsetR turnover plot ---------------------------------------------------------------------------------------------------------------------------------------------
# # UpsetR is a tool to visualize overlaps in datasets, it is analogous to Venn diagrams, yet can plot more dimensions
# # Check Conway et al. 2017, doi: 10.1093/bioinformatics/btx364
# # It can be used with samples (M.pa) or groups (M.grouped.pa). For samplewise analysis nsets has to be changed to the number of samples
# # If only specific samples should be visualize use sets=c('name1','name2',...) and set nsets to the number of samples
# # nintersects: number of intersects to show, set to NA to show all

# # we want to create an Upset plot (and venn diagrams) which account for the different sample sizes per group
# # we need to randomly select 

# ok fast?

min.n.per.group <- min(as.numeric(M.group.count[,2])) # # minimum number of samples in a group
M.grouped.pa.norm <- matrix(nrow=length(unique(M.groups)),ncol=ncol(M)) 
# in M we have the counts matrix with samples on the x axis (alphabetically sorted) and ASVs on the Y axis
# M.groups is the right vector to go with this

for (groups in 1:length(unique(M.groups))) {
  M.subset <- as.matrix(M[M.groups==groups,]) # we select one group
  result <- matrix(ncol=ncol(M))
  for (i in 1:NI) {
    # # randomly subset to the lowest number of samples in one group (min.n.per.group)
    sample_indices <- sample(1:nrow(M.subset),size=min.n.per.group,replace=T)
    M.subset.subset <- as.matrix(M.subset[sample_indices,])
    # we create the columnsums(sums up the ASV counts over all samples)
    M.subset.subset.sum <- t(as.data.frame(colSums(M.subset.subset)))
    # we use deconstand on these summed up table (one row for the group)
    M.pa.subset.subset <- vegan::decostand(M.subset.subset.sum, method='pa')
    # we add the result to our group result table
    result <- rbind(result,M.pa.subset.subset)
  }
  # # remove first row in result table (empty)
  result <- result[-1,]
  result <- as.matrix(result)
  # # calculate column averages (ASVs) over all iterations
  result <- colSums(result)/nrow(result)
  # # round up to 1 if present in more than 50 % of the iterations, and down to 0 if in less
  result <- ifelse(result >= 0.5, 1, 0)
  # # add result to our final pa table
  M.grouped.pa.norm[groups,] <- result
}
rm(M.subset,result,sample_indices,groups,M.subset.subset,M.subset.subset.sum,M.pa.subset.subset)
colnames(M.grouped.pa.norm) <- colnames(M)

# # do upset plot with uncorrected data
if(length(M.projects.unique)>1){
  M.up=as.data.frame(t(M.grouped.pa))
  colnames(M.up)=M.projects.unique.ord
  M.upset=UpSetR::upset(M.up,mb.ratio = c(0.7, 0.3),nsets = length(M.projects.unique),sets=rev.default(M.projects.unique.ord),keep.order = T, nintersects = NA, order.by = c("freq", "degree"), decreasing = c(TRUE,FALSE),sets.bar.color = rev.default(M.col),empty.intersections = "on",point.size = 5,text.scale=2) # keep.order=T: Keep Groups in the order entered using the sets parameter. The default is FALSE, which orders the sets by their sizes.,mb.ratio: Ratio between matrix plot and main bar plot
  pdf(file.path(PathToVisuaRAnalysis,'Beta_Diversity',paste(VisuaRProjectName,"_UpsetPlot.pdf",sep="")),height=6,width=15,useDingbats=F)
  print(M.upset)
  M.upset
  q <- recordPlot()
  # print(M.upset)
  dev.off()
  gc()
}

# # do Upset plot with corrected data (previously named _ncor.pdf)
if(length(M.projects.unique)>1){
  M.up.cor=as.data.frame(t(M.grouped.pa.norm))
  colnames(M.up.cor)=M.projects.unique.ord
  M.upset.cor=UpSetR::upset(M.up.cor,mb.ratio = c(0.7, 0.3),nsets = length(M.projects.unique),sets=rev.default(M.projects.unique.ord),keep.order = T, nintersects = NA, order.by = c("freq", "degree"), decreasing = c(TRUE,FALSE),sets.bar.color = rev.default(M.col),empty.intersections = "on",point.size = 5,text.scale=2) # keep.order=T: Keep Groups in the order entered using the sets parameter. The default is FALSE, which orders the sets by their sizes.,mb.ratio: Ratio between matrix plot and main bar plot
  pdf(file.path(PathToVisuaRAnalysis,'Beta_Diversity',paste(VisuaRProjectName,"_UpsetPlot_cor.pdf",sep="")),height=6,width=15,useDingbats=F)
  print(M.upset.cor)
  M.upset.cor
  # q <- recordPlot()
  # print(M.upset.cor)
  dev.off()
  gc()
}


# #------- 5.6. Create Venn Diagram -----------------------------------------------------------------------------------------------------------------------------------------------------
# # fast

if(length(M.projects.unique)>1 && length(M.projects.unique)<8){
  pdf(file.path(PathToVisuaRAnalysis,'Beta_Diversity',paste(VisuaRProjectName,"_VennDiagram.pdf",sep="")),height=5,width=5,useDingbats=F)
  venn::venn(M.up,zcolor = M.col,opacity = 0.4,ilcs=0.85,sncs=1,scaled=T)
  dev.off()
  gc()
}

# #  with corrected data (previously named _ncor.pdf)
if(length(M.projects.unique)>1 && length(M.projects.unique)<8){
  pdf(file.path(PathToVisuaRAnalysis,'Beta_Diversity',paste(VisuaRProjectName,"_VennDiagram_cor.pdf",sep="")),height=5,width=5,useDingbats=F)
  venn::venn(M.up.cor,zcolor = M.col,opacity = 0.4,ilcs=0.85,sncs=1,scaled=T)
  dev.off()
  gc()
}

# #------- 5.6. Create Scaled Venn Diagram -----------------------------------------------------------------------------------------------------------------------------------------------------

# # fast 

if(length(M.projects.unique)>1 && length(M.projects.unique)<20){
  pdf(file.path(PathToVisuaRAnalysis,'Beta_Diversity',paste(VisuaRProjectName,"_VennDiagram_scaled.pdf",sep="")),height=5,width=5,useDingbats=F)
  vennscaled <- plot(eulerr::euler(M.up),fills=c(M.col),edges=T, legend=F,quantities=T,lw=3,alpha=0.6,col="black")
  print(vennscaled)
  dev.off()
  if (SaveWholeworkspace=='N') {rm(M.up)}
  gc()
}

# # with corrected data (previously named _ncor.pdf)
if(length(M.projects.unique)>1 && length(M.projects.unique)<20){
  pdf(file.path(PathToVisuaRAnalysis,'Beta_Diversity',paste(VisuaRProjectName,"_VennDiagram_scaled_cor.pdf",sep="")),height=5,width=5,useDingbats=F)
  vennscaled <- plot(eulerr::euler(M.up.cor),fills=c(M.col),edges=T, legend=F,quantities=T,lw=3,alpha=0.6,col="black")
  print(vennscaled)
  dev.off()
  if (SaveWholeworkspace=='N') {rm(M.up.cor)}
  gc()
}


# #------- 5.7. Pairwise comparison of groups -------------------------------------------------------------------------------------------------------------------------------------------

if(length(M.projects.unique)>1){
  M.compare=M.grouped.pa  # group-wise analysis, M.grouped.pa is a presence absence matrix from grouped matrix of M
} else {
  M.compare=M.pa          # Sample-wise analysis
}

if (SaveWholeworkspace=='N') {rm(M.pa,M.grouped.pa)}
gc()

if(length(M.projects.unique)>1){
  # # number of unique ASVs of GroupX when comapred with GroupY... 
  unique_for_group=matrix(NA, (nrow(M.compare)^2), 3) # creates a matrix of (number of Groups)^2 rows and 3 columns
  # # number of total unique ASVs 
  unique=matrix(NA,nrow(M.compare),nrow(M.compare))
  rownames(unique)= M.projects.unique.ord
  colnames(unique) = M.projects.unique.ord
  
  # # number of shared OTUs 
  common=matrix(NA,nrow(M.compare),nrow(M.compare)) 
  rownames(common)=M.projects.unique.ord
  colnames(common)=M.projects.unique.ord
  
  # # number of total ASVs 
  total=matrix(NA,nrow(M.compare),nrow(M.compare)) 
  rownames(total)= M.projects.unique.ord
  colnames(total) = M.projects.unique.ord
  
  # # Percentage of shared ASVs ((shared number of ASVs/total number of ASVs)* 100)
  percentage=matrix(NA,nrow(M.compare),nrow(M.compare)) 
  rownames(percentage)= M.projects.unique.ord
  colnames(percentage) = M.projects.unique.ord
  
  b=vector() # creates an empty vector
  p=vector()
  
  rowid = 0                                                               # sets rowid to 0. This will be used to identify the row to fill the values into (first round first row, second round second row....)
  for (r in (1 : (nrow(M.compare)))) {                                    # number of repeats = number of groups
    for (s in (1 : (nrow(M.compare)))) {                                  # number of repeats = number of groups
      for (c in (1 : (ncol(M.compare)))) {                                # number of repeats = number of ASVs
        b[c]=sum (M.compare[r,c],M.compare[s,c])                          # vector b is filled with the sum of cell [1,1] and cell [1,1] (=1 if only in one, =2 if in both, =0 if in none)
        p[c] = if (M.compare[r,c]==1 & M.compare[s,c]== 0) 1 else 0       # vector p is filled with 1 if the ASV occur in cell 1 but not in cell 2 and with 0 if 0 in both or 1 in both
      }
      rowid = rowid + 1;
      unique_for_group[rowid,1] = paste(M.projects.unique.ord[r])         # prints the name of the current sample[r] in the current row[rowid]
      unique_for_group[rowid,2] = paste(M.projects.unique.ord[s])         # prints the name of the current sample[s] in the current row[rowid]
      unique_for_group[rowid,3] = sum(p)                                  # prints the sum of the unique ASVs for sample [r] (do occur in sample r but not in sample [s])
      
      unique[r,s]=length (which (b==1) )                                  # fills unique table with the number of how many ASVs only occur in sample [r] or sample [s]
      common[r,s]= length (which (b==2) )                                 # fills common table with the number of how many ASVS occur in both samples ([r] and [s])
      total[r,s]= sum (unique[r,s], common[r,s])                          # fills total table with the sum of the unique and common ASVs for each sample
      percentage[r,s]=round(common[r,s]/total[r,s]*100,0)                 # fills percentage table with the percentage of shared ASVs for each sample
    }
  }
  if (SaveWholeworkspace=='N') {rm(M.compare)}
  gc()
  
  data.table::fwrite(as.data.frame(total),file=file.path(PathToVisuaRAnalysis,'Beta_Diversity',paste(VisuaRProjectName,"_Total_ASVs_Groups.csv",sep="")),col.names = T,row.names = T)
  data.table::fwrite(as.data.frame(unique),file=file.path(PathToVisuaRAnalysis,'Beta_Diversity',paste(VisuaRProjectName,"_Unique_ASVs_Groups.csv",sep="")),col.names = T,row.names = T)
  data.table::fwrite(as.data.frame(common),file=file.path(PathToVisuaRAnalysis,'Beta_Diversity',paste(VisuaRProjectName,"_Shared_ASVs_Groups.csv",sep="")),col.names = T,row.names = T)
  data.table::fwrite(as.data.frame(percentage),file=file.path(PathToVisuaRAnalysis,'Beta_Diversity',paste(VisuaRProjectName,"_Percent_Shared_ASVs_Groups.csv",sep="")),col.names = T,row.names = T)
  data.table::fwrite(as.data.frame(unique_for_group),file=file.path(PathToVisuaRAnalysis,'Beta_Diversity',paste(VisuaRProjectName,"_Unique_ASVs_per_Grouping.csv",sep="")),col.names = T,row.names = T) # results of "unique_for_group" should read as: sampleI (first column), when compared to sampleJ (second column), has "X Number" (third column) of unique ASVs
  
  if (SaveWholeworkspace=='N') {rm(total,unique,common,percentage,unique_for_group)}
  gc()

}


# #=== 5.8. Create Overview Plot ============================================================================================================================================================================
# # prepare NMDS for use with ggarrange
dev.off()
par(mar=c(2, 2, 2, 7), xpd=F,bg="transparent")                                                                                             # first, xpd=F because if not the ellipses might be drawn outside of the plot region
ordiplot (M.mMDS, display = 'si', type = 'n',ylab="",xlab="",xaxt='n',yaxt='n')
for (i in seq (1, NR)) ordiellipse (M.mMDS, groups = M.groups, show.groups = i, col = M.col[i], label = F,lwd=1.5,draw='polygon',border='NA',alpha=60,kind='sd')
for (i in seq (1, NR)) ordispider (M.mMDS, groups = M.groups, show.groups = i, col = M.col[i], label = F,lwd=1.5)
points (M.mMDS, col=M.colvec, pch=19,cex=1) 
par(new=T,xpd=T)
plot.new()
if(length(M.projects.unique)>1){
  legend(x=1.01,y=1.05,legend='Groups',bty='n',cex=0.75,text.font = 2) 
  legend(1.045,1.00,legend=c(M.projects.unique.ord),fill=c(M.col),bty='n',cex=0.75)
  legend(1.01,0.46,legend=paste0("Community\nSimilarity [%]"),bty='n',cex=0.75,text.font = 2)
  legend(1.01,0.4,legend=paste0('max: ',round(Max.Com.Sim,1),'\nmean: ',round(Mean.Com.Sim,1),'\nmin: ',round(Min.Com.Sim,1)),bty='n',cex=0.75)
  legend(1.01,0.21,legend=paste0('ANOSIM'),bty='n',cex=0.75,text.font = 2)
  legend(1.01,0.17,legend=paste0('R: ',round(M.ano$statistic,3),'\np: ',M.ano$signif),bty='n',cex=0.75)
  legend(1.01,0.03,legend=paste0("Stress: ",round(M.mMDS.stress,3)),bty='n',cex=0.75) 
} else {
  legend(1.01,0.46,legend=paste0("Community\nSimilarity [%]"),bty='n',cex=0.75,text.font = 2)
  legend(1.01,0.4,legend=paste0('max: ',round(Max.Com.Sim,1),'\nmean: ',round(Mean.Com.Sim,1),'\nmin: ',round(Min.Com.Sim,1)),bty='n',cex=0.75)
  legend(1.01,0.03,legend=paste0("Stress: ",round(M.mMDS.stress,3)),bty='n',cex=0.75) 
}

p <- recordPlot()

dev.off()

if(exists("M.upset")) {
  uu_c <- cowplot::plot_grid(NULL, M.upset$Main_bar, M.upset$Sizes, M.upset$Matrix,
                             nrow=2, align='hv', rel_heights = c(3,1),
                             rel_widths = c(0.7,3))
}


if (removeContaminants != "") {
  # # get rid of x axis labels
  perc.asv.nocontam <- perc.asv.nocontam + xlab(NULL) + theme(axis.text.x = element_blank())
  perc.reads.noncontam <- perc.reads.noncontam + xlab(NULL)
}

# # different plots depending on which plots are available
if (removeContaminants != "" & !exists("vennscaled") & !exists("M.upset")) {
  pdf(file.path(PathToVisuaRAnalysis,paste(VisuaRProjectName,"_CommunityOverview.pdf",sep="")),height=50,width=34,useDingbats=F) 
  overview.plot <- ggarrange(ncol=1,nrow=6,labels = c("A","B","C","","F","G"),heights = c(0.75,1,1,1,1,1.3),
                             perc.asv.nocontam,
                             perc.reads.noncontam,
                             M.bubble,
                             ggarrange(NULL,asvs.plot,reads.plot,NULL,ncol=4,labels=c("D","E")),
                             ggarrange(SOBS.plot,Shan.plot,Simp.plot,Chao1.plot,ncol=4,nrow=1,align='hv'),
                             ggarrange(NULL,p,NULL,ncol=3,nrow=1))
  annotate_figure(overview.plot,top = text_grob(paste(VisuaRProjectName," - Overview"), color = "black", size = 14))
  print(overview.plot)
  dev.off()
} else if(removeContaminants != "" & !exists("vennscaled") & exists("M.upset")) {
  pdf(file.path(PathToVisuaRAnalysis,paste(VisuaRProjectName,"_CommunityOverview.pdf",sep="")),height=50,width=34,useDingbats=F) 
  overview.plot <- ggarrange(ncol=1,nrow=7,labels = c("A","B","C","","F","G","H"),heights = c(0.75,1,1,1,1,1.3,1),
                             perc.asv.nocontam,
                             perc.reads.noncontam,
                             M.bubble,
                             ggarrange(NULL,asvs.plot,reads.plot,NULL,ncol=4,labels=c("D","E")),
                             ggarrange(SOBS.plot,Shan.plot,Simp.plot,Chao1.plot,ncol=4,nrow=1,align='hv'),
                             ggarrange(NULL,p,NULL,ncol=3,nrow=1),
                             uu_c)
  annotate_figure(overview.plot,top = text_grob(paste(VisuaRProjectName," - Overview"), color = "black", size = 14))
  print(overview.plot)
  dev.off()
} else if(removeContaminants != "" & exists("vennscaled") & exists("M.upset")) {
  pdf(file.path(PathToVisuaRAnalysis,paste(VisuaRProjectName,"_CommunityOverview.pdf",sep="")),height=58,width=34,useDingbats=F) 
  overview.plot <- ggarrange(ncol=1,nrow=7,labels = c("A","B","C","","F","G","H"),heights = c(0.75,1,1,1,1,1,1),
                             perc.asv.nocontam,
                             perc.reads.noncontam,
                             M.bubble,
                             ggarrange(NULL,asvs.plot,reads.plot,NULL,ncol=4,labels=c("D","E")),
                             ggarrange(SOBS.plot,Shan.plot,Simp.plot,Chao1.plot,ncol=4,nrow=1,align='hv'),
                             ggarrange(p,vennscaled,ncol=2,nrow=1),
                             uu_c)
  annotate_figure(overview.plot,top = text_grob(paste(VisuaRProjectName," - Overview"), color = "black", size = 14))
  print(overview.plot)
  dev.off()
} else if (removeContaminants == "" & !exists("vennscaled") & !exists("M.upset")) {
  pdf(file.path(PathToVisuaRAnalysis,paste(VisuaRProjectName,"_CommunityOverview.pdf",sep="")),height=50,width=34,useDingbats=F) 
  overview.plot <- ggarrange(ncol=1,nrow=4,labels = c("A","","D","E"),heights = c(1,1,1,1.3),
                             M.bubble,
                             ggarrange(NULL,asvs.plot,reads.plot,NULL,ncol=4,labels=c("B","C")),
                             ggarrange(SOBS.plot,Shan.plot,Simp.plot,Chao1.plot,ncol=4,nrow=1,align='hv'),
                             ggarrange(NULL,p,NULL,ncol=3,nrow=1))
  annotate_figure(overview.plot,top = text_grob(paste(VisuaRProjectName," - Overview"), color = "black", size = 14))
  print(overview.plot)
  dev.off()
} else if(removeContaminants == "" & !exists("vennscaled") & exists("M.upset")) {
  pdf(file.path(PathToVisuaRAnalysis,paste(VisuaRProjectName,"_CommunityOverview.pdf",sep="")),height=31,width=34,useDingbats=F) 
  overview.plot <- ggarrange(ncol=1,nrow=5,labels = c("A","","D","E","F"),heights = c(1,1,1,1.3,1),
                             M.bubble,
                             ggarrange(NULL,asvs.plot,reads.plot,NULL,ncol=4,labels=c("B","C")),
                             ggarrange(SOBS.plot,Shan.plot,Simp.plot,Chao1.plot,ncol=4,nrow=1,align='hv'),
                             ggarrange(NULL,p,NULL,ncol=3,nrow=1),
                             uu_c)
  annotate_figure(overview.plot,top = text_grob(paste(VisuaRProjectName," - Overview"), color = "black", size = 14))
  print(overview.plot)
  dev.off()
} else if(removeContaminants == "" & exists("vennscaled") & exists("M.upset")) {
  pdf(file.path(PathToVisuaRAnalysis,paste(VisuaRProjectName,"_CommunityOverview.pdf",sep="")),height=31,width=34,useDingbats=F) 
  overview.plot <- ggarrange(ncol=1,nrow=5,labels = c("A","","D","E",""),heights = c(1,1,1,1.3,1),
                             M.bubble,
                             ggarrange(NULL,asvs.plot,reads.plot,NULL,ncol=4,labels=c("B","C")),
                             ggarrange(SOBS.plot,Shan.plot,Simp.plot,Chao1.plot,ncol=4,nrow=1,align='hv'),
                             ggarrange(NULL,p,NULL,ncol=3,nrow=1),
                             ggarrange(uu_c,vennscaled,ncol=2,labels = c("F","G")))
  annotate_figure(overview.plot,top = text_grob(paste(VisuaRProjectName," - Overview"), color = "black", size = 14))
  print(overview.plot)
  dev.off()
}

# #=== 5.9. Save new seqtab-nochim and taxonomy ============================================================================================================================================================================

# # save new seqtab nochim with above filtering applied already
if(savenewseqtabandtax == "Y") {
  M.seq.tax.subset.save <- M.seq.tax.subset
  row_indices <-  match(rownames(M.seq.tax.subset),ASV.mapfile[,"ASV"])
  rownames(M.seq.tax.subset.save) <- ASV.mapfile[row_indices,"ASV sequence"]

  # # save  taxonomy
  saveRDS((M.seq.tax.subset.save[,(ncol(M.seq.tax.subset.save)-ncol.taxo.noNA+1):ncol(M.seq.tax.subset.save)]),file.path(PathToVisuaRAnalysis,paste(VisuaRProjectName,"_taxa_species_noNAs.rds",sep="")))
  # # save  seqtab
  saveRDS(t(M.seq.tax.subset.save[,1:(ncol(M.seq.tax.subset.save)-ncol.taxo.noNA)]),file.path(PathToVisuaRAnalysis,paste(VisuaRProjectName,"_seqtab_nochim.rds",sep="")))
  
  rm(M.seq.tax.subset.save,row_indices)
}
  
# #=== 6. Epilogue ============================================================================================================================================================================
save.image(file.path(PathToVisuaRAnalysis,paste(VisuaRProjectName,'_VisuaR','.RData',sep='')))
# rstudioapi::documentSave()
closeAllConnections() # closes all currently open connections.
# dev.off()
print(paste0(VisuaRProjectName," finished.",sep=""))
rm(list=ls())
gc()
```
