## Supplementary material for "A global atlas of subsurface microbiomes reveals phylogenetic novelty, large scale biodiversity gradients, and a marine-terrestrial divide": Dataset S5 - VisuaR User Input R script: DatasetS5_VisuaR_UserInput.R

```
# # To be used to provide input for VisuaR analysis. This file will be saved to the folder of the analysis.
# #=== 1. User INPUT =====================================================================================================================================================================
# #=== 1.1. Mandatory INPUT ===============================================================================================================================================================

VisuaRProjectName='20240402_CoDL_arc_envbiome' # Give your analysis a (short) name
# VisuaRProjectName='20240219_CoDL_arc_ter_depthbiome' # Give your analysis a (short) name
# VisuaRProjectName='20240219_CoDL_bac_rps3_kaiju_depthbiome' # Give your analysis a (short) name
# VisuaRProjectName='20240219_CoDL_metagenome_biome' # Give your analysis a (short) name

PathToVisuarOutput=file.path('C:/Users/i.hrabedeangelis/Dropbox/_CoDL_2020_2022/_VisuaR_2024_2')  # At this location the folder 'VisuaR' will be created containing subfolders with your analyses (named as provided in VisuaRProjectname)
isthiscodl <- "Y" # # set to N for metagenome based codl stuff
# # Provide your dada2 output
# # Used this pipeline for your dada2 analysis? https://astrobiomike.github.io/amplicon/dada2_workflow_ex#processing-with-dada2-in-r 
HappyBellydada2Input='N'          # set to 'Y' if you want to use the output generated from the happybelly workflow, set to 'N' if you want to use the output generated by the dada2-pipeline on the VisuaR github page
QIIMEInput='N'
MetagenomeInput='N'

IsthisCoDLrps3_kaiju="N"
IsthisCoDLrps3_cat="N"
IsthisCoDLrps3_uniprot="N"
IsthisCoDL_Metagenome="N"
IsthisCoDLphyloSSU="N"

KingdomOfInterest='Archaea'  # Which kingdom are you interested in? choose Bacteria/Archaea/Eukaryota/All/Archaea|Bacteria...

if (KingdomOfInterest=="Archaea") {
  if (IsthisCoDLrps3_kaiju == "Y") {
    PathToSeqtabNochim=file.path("C:/Users/i.hrabedeangelis/Dropbox/_CoDL_2020_2022/Metagenomes/MG_rps3_diversity/CODL.UniG.RPS3.CPM.csv") # provide the location and name of your seqtab_nochim.rds
    PathToTaxonomy=file.path("C:/Users/i.hrabedeangelis/Dropbox/_CoDL_2020_2022/Metagenomes/MG_rps3_diversity/CODL.UniG.RPS3.CPM.csv")      # provide the location and name of your taxonomy.rds
    MinimumAllowedReadCount=100             # MinimumAllowedReadcount depends on the overall quality of the run and targeted sequencing depth and is used to remove samples with low reads (failed samples, controls). All samples with less reads than this number will be excluded from the analysis. This is based on all reads derived from your dada2 analysis, no matter which kingdom they belong to.
    MinimumAllowedReadCount.Analysis=100    # This field can be set to any number including 0 (zero) depending on which samples (based on read counts) should be included in the analysis. This happens after all ASVs and samples have been excluded as set in KeepSamplesbyName, excludeSamplesbyName, CladeOfInterest, ExcludeClade.
    # excludeSamplesbyName="^dco_ins_str2$|dco_ins_str1$|^dco_boy_0$|^dco_boy_9$|^dco_boy_15$|^dco_slu_wt7$|^dco_wri_2a_day6_5$|^dco_mou_sample_06272012$|^dco_wri_3_day13_3$"
    ColumnForMerging <- "NewSampleID_rps3" 
    excludeSamplesbyName =""
    # for min9 analyses
    excludeSamplesbyName="^dco_ins_str2$|dco_ins_str1$|^dco_boy_0$|^dco_boy_9$|^dco_boy_15$|^dco_slu_wt7$|^dco_wri_2a_day6_5$|^dco_mou_sample_06272012$|^dco_wri_3_day13_3$"
  } else if (IsthisCoDLphyloSSU == "Y") {
    PathToSeqtabNochim=file.path("C:/Users/i.hrabedeangelis/Dropbox/_CoDL_2020_2022/Metagenomes/MG_SSU/Phyloflash_SSU_16S_Tables/all.phyloFlash.NTUfull_abundance-pivot.csv") # provide the location and name of your seqtab_nochim.rds
    PathToTaxonomy=file.path("C:/Users/i.hrabedeangelis/Dropbox/_CoDL_2020_2022/Metagenomes/MG_SSU/Phyloflash_SSU_16S_Tables/all.phyloFlash.NTUfull_abundance-pivot.csv")      # provide the location and name of your taxonomy.rds
    KingdomOfInterest='Archaea'  # Which kingdom are you interested in? choose Bacteria/Archaea/Eukaryota/All/Archaea|Bacteria...
    MinimumAllowedReadCount=100             # MinimumAllowedReadcount depends on the overall quality of the run and targeted sequencing depth and is used to remove samples with low reads (failed samples, controls). All samples with less reads than this number will be excluded from the analysis. This is based on all reads derived from your dada2 analysis, no matter which kingdom they belong to.
    MinimumAllowedReadCount.Analysis=100    # This field can be set to any number including 0 (zero) depending on which samples (based on read counts) should be included in the analysis. This happens after all ASVs and samples have been excluded as set in KeepSamplesbyName, excludeSamplesbyName, CladeOfInterest, ExcludeClade.
    excludeSamplesbyName =""
    ColumnForMerging <- "NewSampleID_phylo" 

  } else {
    # # codl arc
    PathToSeqtabNochim=file.path("C:/Users/i.hrabedeangelis/Dropbox/_CoDL_2020_2022/All_Dada2_v138/arc_merge_DCOwbrz-nonDCO_seqtab_nochim_all.rds") # provide the location and name of your seqtab_nochim.rds
    PathToTaxonomy=file.path("C:/Users/i.hrabedeangelis/Dropbox/_CoDL_2020_2022/All_Dada2_v138/arc_merge_DCOwbrz-nonDCO_taxa_species_v138.rds")      # provide the location and name of your taxonomy.rds
    # # codl arc
    excludeSamplesbyName='DCO_BRZ|DCO_GRD|^DCO_BRZ_Av4v5_Serp_LC_BB13$|^DCO_BRZ_Av4v5_Serp_CRO_8A_Aug11_Av4v5$|^DCO_BRZ_Av4v5_Serp_CRO_8A_Apr11_Av4v5$|^DCO_BRZ_Av4v5_Serp_CRO_8B_AprD11_Av4v5$|^DCO_BRZ_Av4v5_Serp_CRO_8B_AprC11_Av4v5$|^DCO_BRZ_Av4v5_Serp_CRO_CSW1_4A_Mar13$|^DCO_BRZ_Av4v5_Serp_CRO_CSW1_2A_Mar13$|^DCO_BRZ_Av4v5_Serp_CRO_CSW1_3B_Mar13$|^DCO_BRZ_Av4v5_Serp_CRO_CSW1_5A_Mar13$|^DCO_BRZ_Av4v5_Serp_CRO_CSW1_1A_Mar13$|^DCO_BRZ_Av4v5_Serp_CRO_CSWoldA_Mar13$|^DCO_BRZ_Av4v5_Serp_CRO_CSWoldT0_Mar13$|^DCO_BRZ_Av4v5_Serp_CRO_N08CA_Mar13$|^DCO_BRZ_Av4v5_Serp_CRO_N08AB_Mar13$|^DCO_BRZ_Av4v5_Serp_CRO_N08BB_Mar13$|^DCO_BRZ_Av4v5_Serp_CRO_N08BA_Mar13$|^DCO_BRZ_Av4v5_Serp_CRO_N08T0_Mar13$|^DCO_BRZ_Av4v5_Serp_CRO_QV1_3B_Mar13$|^DCO_BRZ_Av4v5_Serp_CRO_QV1_2B_Mar13$|^DCO_BRZ_Av4v5_Serp_CRO_QV1_1A_Mar13$|^DCO_BRZ_Av4v5_Serp_Lig_GORupA_2013$|^DCO_BRZ_Av4v5_Serp_TL_WHC2B_Aug10$|^DCO_BRZ_Av4v5_Serp_LC1408$|^DCO_BRZ_Av4v5_Serp_LC_BB35$|^DCO_BRZ_Av4v5_Serp_LC_BB31$|^DCO_BRZ_Av4v5_Serp_LC_BB22$|^DCO_BRZ_Av4v5_Serp_LC_BB20$|^DCO_BRZ_Av4v5_Serp_LC_BB21$|^KIE_CCL_Av4v5_CS_SL_4_B$|^DCO_BKR_Av4v5_Knox_65C_7H2$|^SBJ_BME_Av4v5_Dead_Sea_2$|^SBJ_BME_Av4v5_Dead_Sea_1$|^SBJ_BME_Av4v5_Lake_Fryxell_3$|^DCO_MAR_Av4v5_10A$|^DCO_MAR_Av4v5_9A$|^DCO_PIN_Av4v5_A3$|^DCO_TAS_Av4v5_S4_30$|^DCO_TAS_Av4v5_S4_10$|^DCO_PIN_Av4v5_A2$|^DCO_LZA_Av4v5_AHS_5$'
    ColumnForMerging <- "SAMPLE_ID_OTU_TABLE_withREseq_Av4v5" # column in contextdata with names to use to merge contextdata to sequencing data (should be the same names) , if left empty, the first column of the contextual data will be used
    MinimumAllowedReadCount=1000             # MinimumAllowedReadcount depends on the overall quality of the run and targeted sequencing depth and is used to remove samples with low reads (failed samples, controls). All samples with less reads than this number will be excluded from the analysis. This is based on all reads derived from your dada2 analysis, no matter which kingdom they belong to.
    MinimumAllowedReadCount.Analysis=1000    # This field can be set to any number including 0 (zero) depending on which samples (based on read counts) should be included in the analysis. This happens after all ASVs and samples have been excluded as set in KeepSamplesbyName, excludeSamplesbyName, CladeOfInterest, ExcludeClade.
  }
} else if(KingdomOfInterest=="Bacteria") {
  if (IsthisCoDLrps3_kaiju == "Y") {
    PathToSeqtabNochim=file.path("C:/Users/i.hrabedeangelis/Dropbox/_CoDL_2020_2022/Metagenomes/MG_rps3_diversity/CODL.UniG.RPS3.CPM.csv") # provide the location and name of your seqtab_nochim.rds
    PathToTaxonomy=file.path("C:/Users/i.hrabedeangelis/Dropbox/_CoDL_2020_2022/Metagenomes/MG_rps3_diversity/CODL.UniG.RPS3.CPM.csv")      # provide the location and name of your taxonomy.rds
    MinimumAllowedReadCount=100             # MinimumAllowedReadcount depends on the overall quality of the run and targeted sequencing depth and is used to remove samples with low reads (failed samples, controls). All samples with less reads than this number will be excluded from the analysis. This is based on all reads derived from your dada2 analysis, no matter which kingdom they belong to.
    MinimumAllowedReadCount.Analysis=100    # This field can be set to any number including 0 (zero) depending on which samples (based on read counts) should be included in the analysis. This happens after all ASVs and samples have been excluded as set in KeepSamplesbyName, excludeSamplesbyName, CladeOfInterest, ExcludeClade.
    # excludeSamplesbyName="^dco_ins_str2$|dco_ins_str1$|^dco_boy_0$|^dco_boy_9$|^dco_boy_15$|^dco_slu_wt7$|^dco_wri_2a_day6_5$|^dco_mou_sample_06272012$|^dco_wri_3_day13_3$"
    ColumnForMerging <- "NewSampleID_rps3" 
    excludeSamplesbyName=""
  } else if (IsthisCoDLphyloSSU == "Y") {
      PathToSeqtabNochim=file.path("C:/Users/i.hrabedeangelis/Dropbox/_CoDL_2020_2022/Metagenomes/MG_SSU/Phyloflash_SSU_16S_Tables/all.phyloFlash.NTUfull_abundance-pivot.csv") # provide the location and name of your seqtab_nochim.rds
      PathToTaxonomy=file.path("C:/Users/i.hrabedeangelis/Dropbox/_CoDL_2020_2022/Metagenomes/MG_SSU/Phyloflash_SSU_16S_Tables/all.phyloFlash.NTUfull_abundance-pivot.csv")      # provide the location and name of your taxonomy.rds
      KingdomOfInterest='Bacteria'  # Which kingdom are you interested in? choose Bacteria/Archaea/Eukaryota/All/Archaea|Bacteria...
      MinimumAllowedReadCount=100             # MinimumAllowedReadcount depends on the overall quality of the run and targeted sequencing depth and is used to remove samples with low reads (failed samples, controls). All samples with less reads than this number will be excluded from the analysis. This is based on all reads derived from your dada2 analysis, no matter which kingdom they belong to.
      MinimumAllowedReadCount.Analysis=100    # This field can be set to any number including 0 (zero) depending on which samples (based on read counts) should be included in the analysis. This happens after all ASVs and samples have been excluded as set in KeepSamplesbyName, excludeSamplesbyName, CladeOfInterest, ExcludeClade.
      excludeSamplesbyName =""
      ColumnForMerging <- "NewSampleID_phylo" 
      
  } else if (IsthisCoDL_Metagenome == "Y") {
    PathToSeqtabNochim=file.path("C:/Users/i.hrabedeangelis/Dropbox/_CoDL_2020_2022/Metagenomes/VisuaR/20200803_CoDL_Metagenomes/GenebySample_CoDL.txt") # provide the location and name of your seqtab_nochim.rds
    PathToTaxonomy=file.path("C:/Users/i.hrabedeangelis/Dropbox/_CoDL_2020_2022/Metagenomes/VisuaR/20200803_CoDL_Metagenomes/Taxonomy.txt")      # provide the location and name of your taxonomy.rds
    KingdomOfInterest='Bacteria'  # Which kingdom are you interested in? choose Bacteria/Archaea/Eukaryota/All/Archaea|Bacteria...
    MinimumAllowedReadCount=0             # MinimumAllowedReadcount depends on the overall quality of the run and targeted sequencing depth and is used to remove samples with low reads (failed samples, controls). All samples with less reads than this number will be excluded from the analysis. This is based on all reads derived from your dada2 analysis, no matter which kingdom they belong to.
    MinimumAllowedReadCount.Analysis=0    # This field can be set to any number including 0 (zero) depending on which samples (based on read counts) should be included in the analysis. This happens after all ASVs and samples have been excluded as set in KeepSamplesbyName, excludeSamplesbyName, CladeOfInterest, ExcludeClade.
    excludeSamplesbyName =""
    ColumnForMerging <- "SampleID_kegg" 
    
  } else {
    # # codl bac
    PathToSeqtabNochim=file.path("C:/Users/i.hrabedeangelis/Dropbox/_CoDL_2020_2022/All_Dada2_v138/bac_merge_DCOwbrz-nonDCO_seqtab_nochim_all.rds") # provide the location and name of your seqtab_nochim.rds
    PathToTaxonomy=file.path("C:/Users/i.hrabedeangelis/Dropbox/_CoDL_2020_2022/All_Dada2_v138/bac_merge_DCOwbrz-nonDCO_taxa_species_v138.rds")      # provide the location and name of your taxonomy.rds
    # # codl bac
    excludeSamplesbyName='^DCO_BRZ_Bv4v5_CRO_QV1_2B_Jul14$|^DCO_BRZ_Bv4v5_Serp_LC_BB20$|^DCO_BRZ_Bv4v5_Serp_LC_BB36$|^DCO_BRZ_Bv4v5_CRO_CSW1_1C_Aug13$|^DCO_BRZ_Bv4v5_CRO_QV1_1A_Aug13$|^DCO_BRZ_Bv4v5_CRO_CSW1_1A_Aug13$|^DCO_BRZ_Bv4v5_CRO_CSW1_1B_Aug13$|^DCO_BRZ_Bv4v5_CRO_QV1_1C_Aug13$|^DCO_BRZ_Bv4v5_Serp_LC_BB13$|^DCO_BRZ_Bv4v5_N08Bc_Aug12$|^DCO_BRZ_Bv4v5_CRO_QV1_1B_Aug13$|^DCO_BRZ_Bv4v5_N08A_B_Aug13$|^DCO_BRZ_Bv4v5_N08B_July11$|^DCO_BRZ_Bv4v5_N08C_A_Aug13$|^DCO_BRZ_Bv4v5_N08B_A_Aug13$|^DCO_BRZ_Bv4v5_CRO_QV1_2C_Aug13$|^DCO_BRZ_Bv4v5_Serp_CRO_QV1_1A_Mar13$|^DCO_BRZ_Bv4v5_N08Bb_Aug12$|^DCO_BRZ_Bv4v5_CRO_CSW1_3C_Aug13$|^DCO_BRZ_Bv4v5_Serp_CRO_CSWoldA_Mar13$|^DCO_BRZ_Bv4v5_Serp_LC_BB6$|^DCO_BRZ_Bv4v5_Serp_Lig_GOR1CDEF_2013$|^DCO_BRZ_Bv4v5_Serp_CRO_CSW1_1A_Mar13$|^DCO_BRZ_Bv4v5_Serp_CRO_8B_AprC11_Bv4v5$|^DCO_BRZ_Bv4v5_Serp_CRO_CSWoldB_Mar13$|^DCO_BRZ_Bv4v5_Serp_CRO_8A_Aug11_Bv4v5$|^DCO_BRZ_Bv4v5_Serp_TL_WHC2B_Aug10$|^DCO_BRZ_Bv4v5_Serp_Lig_GOR1AB_2013$|^DCO_BRZ_Bv4v5_Serp_CRO_8B_AprD11_Bv4v5$|^DCO_BRZ_Bv4v5_Serp_CRO_8A_Apr11_Bv4v5$|^DCO_BRZ_Bv4v5_Serp_CRO_CSWoldT0_Mar13$|^DCO_BRZ_Bv4v5_CRO_QV1_2B_Aug13$|^DCO_BRZ_Bv4v5_Serp_CRO_N08BA_Mar13$|^DCO_BRZ_Bv4v5_Serp_LIG_Gor_upB_Jul12_Bv4v5$|^DCO_BRZ_Bv4v5_Serp_CRO_CSW1_2A_Mar13$|^DCO_BRZ_Bv4v5_Serp_CRO_CSW1_4A_Mar13$|^DCO_BRZ_Bv4v5_Serp_CRO_N08A_Aug11_Bv4v5$|^DCO_BRZ_Bv4v5_CRO_CSW1_3A_Aug13$|^DCO_BRZ_Bv4v5_Serp_TL_WHC75_DNA2_Jun11_Bv4v5$|^DCO_BRZ_Bv4v5_Serp_CRO_CSWold_11_11_Bv4v5$|^DCO_BRZ_Bv4v5_Serp_LIG_L43_upstream_Oct11_Bv4v5$|^DCO_BRZ_Bv4v5_Serp_CRO_QV13A_Aug12_Bv4v5$|^DCO_BRZ_Bv4v5_Serp_LIG_BR2_back1_Jun12_Bv4v5$|^DCO_BRZ_Bv4v5_Serp_LIG_Gor_upA_Jul12_Bv4v5$|^DCO_BRZ_Bv4v5_Serp_LIG_BR2_back2_Jun12_Bv4v5$|^DCO_BRZ_Bv4v5_Serp_TL_WHC2C_DNA2_Jun11_Bv4v5$|^DCO_BRZ_Bv4v5_Serp_TL_WHC2A_DNA3_Jun11_Bv4v5$|^DCO_BRZ_Bv4v5_Serp_CRO_CSW1_3B_Mar13$|^DCO_BRZ_Bv4v5_Serp_CRO_QV1_3B_Mar13$|^DCO_BRZ_Bv4v5_Serp_CRO_CSW1_5A_Mar13$|^DCO_BRZ_Bv4v5_Serp_CRO_N08AB_Mar13$|^DCO_BRZ_Bv4v5_Serp_CRO_QV1_2B_Mar13$|^DCO_BRZ_Bv4v5_Serp_CRO_N08CA_Mar13$|^DCO_BRZ_Bv4v5_Serp_CRO_N08BB_Mar13$|^DCO_BRZ_Bv4v5_Serp_CRO_N08T0_Mar13$|^DCO_BRZ_Bv4v5_Serp_LC1408$|^DCO_BRZ_Bv4v5_Serp_LC_BB21$|^DCO_BRZ_Bv4v5_Serp_Lig_GORupA_2013$|^DCO_BRZ_Bv4v5_Serp_LC_BB22$|^DCO_BRZ_Bv4v5_Serp_LC_BB31$|^DCO_BRZ_Bv4v5_Serp_Lig_GORupB_2013$|^DCO_BRZ_Bv4v5_Serp_LC_BB35$|^DCO_BRZ_Bv4v5_N08A_A_Dec13$|^DCO_BRZ_Bv4v5_CRO_CSW1_1C_Jul14$|^DCO_BRZ_Bv4v5_N08Aa_Aug12$|^DCO_BRZ_Bv4v5_CRO_QV1_1C_Jul14$|^DCO_BRZ_Bv4v5_CSWold_A_Dec13$|^DCO_BRZ_Bv4v5_CRO_CSW1_1B_Jul14$|^DCO_BRZ_Bv4v5_N08Ba_Aug12$|^DCO_BRZ_Bv4v5_CRO_QV1_2A_Jul14$|^DCO_BRZ_Bv4v5_CRO_QV1_1B_Jul14$|^DCO_BRZ_Bv4v5_N5A_Aug12$|^DCO_BRZ_Bv4v5_N08B_A_Dec13$|^DCO_BRZ_Bv4v5_CRO_CSW1_3A_Jul14$|^DCO_BRZ_Bv4v5_CRO_CSW1_1A_Jul14$|^DCO_BRZ_Bv4v5_CRO_QV1_1A_Jul14$|^DCO_BRZ_Bv4v5_N1A_Aug12$|^DCO_BRZ_Bv4v5_N08C_A_Dec13$|^DCO_BRZ_Bv4v5_CRO_CSW1_3B_Jul14$|^DCO_BRZ_Bv4v5_CSWold_Mar12$|^DCO_BRZ_Bv4v5_CRO_CSW1_3B_Aug13$|^DCO_BRZ_Bv4v5_CSWold_A_Aug13$|^DCO_BRZ_Bv4v5_CRO_QV1_2A_Aug13$|DCO_GRD|^DCO_MAR_Bv4v5_8B$|^DCO_MAR_Bv4v5_7B$|^DCO_TAS_Bv4v5_S4_30$|^DCO_TAS_Bv4v5_S4_10$'
    ColumnForMerging <- "SAMPLE_ID_OTU_TABLE_withREseq" # column in contextdata with names to use to merge contextdata to sequencing data (should be the same names) , if left empty, the first column of the contextual data will be used
    MinimumAllowedReadCount=50000             # MinimumAllowedReadcount depends on the overall quality of the run and targeted sequencing depth and is used to remove samples with low reads (failed samples, controls). All samples with less reads than this number will be excluded from the analysis. This is based on all reads derived from your dada2 analysis, no matter which kingdom they belong to.
    MinimumAllowedReadCount.Analysis=2000    # This field can be set to any number including 0 (zero) depending on which samples (based on read counts) should be included in the analysis. This happens after all ASVs and samples have been excluded as set in KeepSamplesbyName, excludeSamplesbyName, CladeOfInterest, ExcludeClade.
    
  }
}


SaveWholeworkspace='Y'            # Set this variable to 'N' to save computational resources. R might run into memory issues if you have large datasets of e.g. 1000 samples. Set this variable to 'Y' to save all created tables in your workspace.

class.algorithm <- 'dada2' # dada2 or dec for dada2 or decipher algorithm used for taxonomiy assignment
tax.database <- 'silva1381'  # database used for taxonomy assignment (for dada2: silva1381, silva138, silva132, rdp18, pr2, unite. for decipher: silva138, rdp18, pr2v4, unite2021.)

# #=== 1.2. Optional INPUT - Sample selection  =============================================================================================================================================

# # when entering names with e.g. ^Terrabacter$ (exact match), be aware that you have to enter the whole name, i.e. ^Bacteria_Actinobacteriota_Actinobacteria_Micrococcales_Intrasporangiaceae_Terrabacter$
CladeOfInterest=''                # Enter one or several clade(s) or keywords as found in your taxa.rds file, e.g. 'Desulfo|Meth'. Only keeps ASVs belonging to lineages containing the term 'Desulfo' and/or 'Meth', i.e. 'Methylococcus', 'Methylotenera'. use "^Methylotenera$|^Methylococcus$" for exact matches
ExcludeClade=''                   # Enter one or several clade(s) or keywords as found in your taxa.rds file. e.g. 'Escherichia|Salmonella'. Excludes ASVs belonging to lineages containing the term 'Escherichia' and/or 'Salmonella', i.e. Escherichia coli.

# # after Salter et al. 2014,  doi:10.1186/s12915-014-0087-z:
# ExcludeClade='Afipia|Aquabacterium|Asticcacaulis|Aurantimonas|Beijerinckia|Bosea|Bradyrhizobium|Brevundimonas|Caulobacter|Craurococcus|Devosia|Hoeflea|Mesorhizobium|Methylobacterium|Novosphingobium|Ochrobactrum|Paracoccus|Pedomicrobium|Phyllobacterium|Rhizobium|Roseomonas|Sphingobium|Sphingomonas|Sphingopyxis|Acidovorax|Azoarcus|Azospira|Burkholderia|Comamonas|Cupriavidus|Curvibacter|Delftia|Duganella|Herbaspirillum|Janthinobacterium|Kingella|Leptothrix|Limnobacter|Massilia|Methylophilus|Methyloversatilis|Oxalobacter|Pelomonas|Polaromonas|Ralstonia|Schlegelella|Sulfuritalea|Undibacterium|Variovorax|Acinetobacter|Enhydrobacter|Enterobacter|Escherichia|Nevskia|Pseudomonas|Pseudoxanthomonas|Psychrobacter|Stenotrophomonas|Xanthomonas|Aeromicrobium|Arthrobacter|Beutenbergia|Brevibacterium|Corynebacterium|Curtobacterium|Dietzia|Geodermatophilus|Janibacter|Kocuria|Microbacterium|Micrococcus|Microlunatus|Patulibacter|Propionibacterium|Rhodococcus|Tsukamurella|Abiotrophia|Bacillus|Brevibacillus|Brochothrix|Facklamia|Paenibacillus|Streptococcus|Chryseobacterium|Dyadobacter|Flavobacterium|Hydrotalea|Niastella|Olivibacter|Pedobacter|Wautersiella|Deinococcus'                   # Enter one or several clade(s) or keywords as found in your taxa.rds file. e.g. 'Escherichia|Salmonella'. Excludes ASVs belonging to lineages containing the term 'Escherichia' and/or 'Salmonella', i.e. Escherichia coli.
# # potential human contaminants after Lang-Yona et al., 2022 doi: https://doi.org/10.1038/s43247-022-00441-6
# ExcludeClade="Bacteroides|Bifidobacterium|Corynebacterium|Cutibacterium|Escherichia|Shigella|Faecalibacterium|Haemophilus|Klebsiella|Lactobacillus|Listeria|Moraxella|Neisseria|Porphyromonas|Prevotella|Salmonella|Staphylococcus|Veillonella"

# #=== 1.3. Optional INPUT - Sample selection - removal of selected samples =============================================================================================================================================
# # in case that you are renaming the samples with the contextual data provided (see sectin 1.4), you have to use the "new" assigned sample names here
KeepSamplesbyName=''              # Enter names of samples that you want to keep. Leave blank to keep all samples. For an exact match use e.g. '^SampleName1$|^SampleName2$'. The use of general terms is possible too, e.g. 'control|blank|enrichment|cont|cntrl|enr|extrctrl|Control|Blank|Enrichment|Cont|Cntrl|Enr|Extrctrl'.


# #=== 1.4. Optional INPUT - Contextual data ==============================================================================================================================================

contextdata='Y' # Do you have contextual data? If not write 'N' , if yes write 'Y' and select a grouping category (Grouping1) as found in your contextdata header.

# # Useful notes on how to prepare your contextdata file:
# # replace missing values by NA
# # replace spaces or special symbols by underscores
# # use unique row names, rownames have to be exactly the same as your sample names from your sequencing files
# # dates: use 01/01/2016 instead of 01/01/16
# # do not start row or column names with a number
# # use a .txt file

if (IsthisCoDLrps3_kaiju == "Y") {
  contextdatafilepath=file.path("C:/Users/i.hrabedeangelis/Dropbox/_CoDL_2020_2022/_CoDL_2022_MG_Context_Data/MetadataForVisuaR_Feb2024.xlsx")  # provide the file path to your contextual data. i.e. 'C:/Users/contextdata.txt'
  sheetname <- "Nov2022_sort"
} else if (IsthisCoDLphyloSSU == "Y") {
  contextdatafilepath=file.path("C:/Users/i.hrabedeangelis/Dropbox/_CoDL_2020_2022/_CoDL_2022_MG_Context_Data/MetadataForVisuaR_Feb2024.xlsx")  # provide the file path to your contextual data. i.e. 'C:/Users/contextdata.txt'
  sheetname <- "Nov2022_sort"
} else if (IsthisCoDL_Metagenome == "Y") {
  contextdatafilepath=file.path("C:/Users/i.hrabedeangelis/Dropbox/_CoDL_2020_2022/_CoDL_2022_MG_Context_Data/MetadataForVisuaR_Feb2024.xlsx")  # provide the file path to your contextual data. i.e. 'C:/Users/contextdata.txt'
  sheetname <- "Nov2022_sort"
} else {
  contextdatafilepath=file.path("C:/Users/i.hrabedeangelis/Dropbox/_CoDL_2020_2022/Context_Data/_CoDL_Contextdata_All_Apr2024.xlsx")  # provide the file path to your contextual data. i.e. 'C:/Users/contextdata.txt'
  sheetname <- "ContextData_clean_2024" # provide the name of the sheet inside your .xlsx file in case you provided one
  
}
excelstartrow <- 1 # only needed if input is an excel dataframe, set to 1 if you want to read in the whole file or to the first row you want to be kept

ColumnForAnalysisNames <- "" # optional, If not provided the same as the column for merging will be used to refer to samples within the analysis


# Grouping1='DEPTH_BIOME_FEATURE'
# Grouping1='DEPTH_BIOME'
Grouping1='ENV_BIOME'
Grouping2=""                    # The samples within Grouping1 will be sorted using this continuous grouping parameter (e.g. concentration, depth profile) for visualization, smalles value is set first

VariableToExclude="FALSE"            # Enter a contextual data variable to exclude. E.g. '4C'
ColumnOfVariableToExclude="INCLUDE"    # Enter column name where the VariableToExclude can be found. E.g. 'Temperature'
  
# VariableToKeep="marine"            # Enter a contextual data variable to keep E.g. '4C'
# VariableToKeep="terrestrial"            # Enter a contextual data variable to keep E.g. '4C'
VariableToKeep=""            # Enter a contextual data variable to keep E.g. '4C'
# ColumnOfVariableToKeep="ENV_BIOME"    # Enter column name where the VariableToKeep can be found. E.g. 'Temperature'
ColumnOfVariableToKeep=""    # Enter column name where the VariableToKeep can be found. E.g. 'Temperature'

CalculateIndicators='N'          # If you have provided a Grouping 1 you can set this to 'Y' and VisuaR will calculate IndicatorSpecies on all taxonomic levels for you. Be aware that this might be time-consuming for a large dataset, if you are not interested in Indicator Species set it to 'N'


# #=== 1.5. Specify VisuaR output =========================================================================================================================================================

NumberOfTOPClades <- 20      # Choose the number of most abundant clades that should be included in the relative abundance plot (bubbleplot). Remaining clades will be summed up and shown as 'Others'. A number between 10 and 30 is recommended.
NumberOfTOPClades.box <- 30  # Choose the number of most abundant clades that should be included in the relative abundance plot (boxplots). A number of 5 to 20 is recommended.
NumberOfWilcoxTtestClades <- 20 # Number or "all" Choose the number of clades you want to do wilcoxon test on. This takes quite long depending on your data

NI <- 10                     # number of iterations used for the calculation of alpha diversity indices (iterations will take long for big datasets, 10-50 are recommended based on number of samples analysed).

savenewseqtabandtax <- "N"

# M.col <- ""
if (Grouping1=='DEPTH_BIOME' && (IsthisCoDLphyloSSU=="Y" | IsthisCoDLrps3_kaiju=="Y" | IsthisCoDLrps3_cat=="Y" | IsthisCoDLrps3_uniprot=="Y") && KingdomOfInterest=="Archaea" ) {
  M.col <- c("#6fe4f2","#2575e5","#de9423","#17258d","#f41f1c") # no samples with surface terrestrial
} else if (Grouping1=='DEPTH_BIOME'&& ColumnOfVariableToKeep=="ENV_BIOME" && VariableToKeep=="terrestrial") {
  M.col <- c("#feea4b","#de9423","#f41f1c") # light blue, yellow, blue, orange, dark blue, red
} else if (Grouping1=='DEPTH_BIOME'&& ColumnOfVariableToKeep=="ENV_BIOME" && VariableToKeep=="marine") {
  M.col <- c("#6fe4f2","#2575e5","#17258d") # light blue, yellow, blue, orange, dark blue, red
} else if (Grouping1=='DEPTH_BIOME') {
 M.col <- c("#6fe4f2","#feea4b","#2575e5","#de9423","#17258d","#f41f1c") # light blue, yellow, blue, orange, dark blue, red
} else if (Grouping1=='DEPTH_BIOME_FEATURE' && KingdomOfInterest =='Bacteria') {
    M.col <- c("#6fe4f2","#59B6C2","#47929B", # 3 times blue for surface marine
               "#feea4b","#CBBB3C","#A29630","#827826", # 4x yellow for surface terrestrial
               "#2575e5","#184B92", #2x blue for interface marine
               "#de9423","#B2761C","#8E5E16", #3x orange for interface terrestrial
               "#17258d", #1x blue for subsurface marine
               "#f41f1c") # 1x red for subsurface terrestrail
} else if (Grouping1=='DEPTH_BIOME_FEATURE' && KingdomOfInterest=='Archaea') {
    M.col <- c("#6fe4f2",# 1x blue for surface marine
               "#feea4b","#CBBB3C", # 2x yellow for surface terrestrial
               "#2575e5","#184B92", #2x blue for interface marine
               "#de9423","#B2761C","#8E5E16", #3x orange for interface terrestrial
               "#17258d", #1x blue for subsurface marine
               "#f41f1c") # 1x red for subsurface terrestrail
} else if (Grouping1 == "ENV_BIOME") {
  M.col <- c("#9A56C8","#C7B299") # marine vs terrestrial
}
# if (Grouping1=='DEPTH_BIOME') {
#   M.col <- c("#6fe4f2","#feea4b","#2575e5","#de9423","#17258d","#f41f1c") # light blue, yellow, blue, orange, dark blue, red
# }
# # M.col <- c("")               # You can provide a color vector, e.g. c('blue','orange') or c('#CC79A7','#999999'). The vector needs as many colors as unique variables found in 'Grouping1'. The colors will be assigned to the Grouping1 variables in alphabetical order.
# if (Grouping1=='DEPTH_BIOME_FEATURE' && KingdomOfInterest =='Bacteria') {
#   M.col <- c("#6fe4f2","#59B6C2","#47929B", # 3 times blue for surface marine
#              "#feea4b","#CBBB3C","#A29630","#827826", # 4x yellow for surface terrestrial
#              "#2575e5","#184B92", #2x blue for interface marine
#              "#de9423","#B2761C","#8E5E16", #3x orange for interface terrestrial
#              "#17258d", #1x blue for subsurface marine
#              "#f41f1c") # 1x red for subsurface terrestrail
# }
# 
# 
# if (Grouping1=='DEPTH_BIOME_FEATURE' && KingdomOfInterest=='Archaea') {
#   M.col <- c("#6fe4f2",# 1x blue for surface marine
#              "#feea4b","#CBBB3C", # 2x yellow for surface terrestrial
#              "#2575e5","#184B92", #2x blue for interface marine
#              "#de9423","#B2761C","#8E5E16", #3x orange for interface terrestrial
#              "#17258d", #1x blue for subsurface marine
#              "#f41f1c") # 1x red for subsurface terrestrail
# }

# #=== 1.6. Optional INPUT - removal of potential contaminants =============================================================================================================================================

VisuaRContaminants="N" # (N: normal analysis). If set to Y this vizualizes the "contaminant" fraction distinguished by the parameters set below. All ASVs/Reads which are not contaminants will be excluded from the analysis and only contaminants analyzed, to investigate potential sources.
removeContaminants="" # leave empty if no removal is wanted!
# # Set to "remove" if you want to eliminate all ASVs which occur in your blanks. If you have more than one sample-blank group in your contextual data, only the ASVs occuring in the respective blank group will be eliminated from their sample group
# # set to "subtract-average" if you want to average the ASV counts in your controls and then subtract the average ASV count from your samples. If you provide sample-blank groupings below, the ASV counts of different controls will be averaged and then subtracted from the respective selection of samples (e.g. water controls get subtracted from water samples, air controls get subtracted from air samples)
# # set to "subtract-max" if you want to subtract the maximum read count occuring for the respective ASVs in the blanks from your samples instead of subtracting the average
# # set to "decontam" if you want to use the functions of the package "decontam" (https://github.com/benjjneb/decontam)
# # leave empty if you do not want to remove any potentially contaminant ASVs


# # if you want to normalize your read counts before removing/subtracting them set to Y (reads in samples are converted to frequencies by summing read counts per sample to 1)
normalizeMseqtax <- "Y" # Y/N

# # add some probability testing
stat.test <- "Y" # set to Y if you want to perform a statistical test before the ASVs are removed/subtracted. Cant be used together with "ASVcountInBlanksToExclude" 
testtouse <- "t" # set to "t" or "w" to use t-test or wilcoxon test (does not assume normal distribution)
signiflevel <- 0.05

# # Provide a column defining blank batches (e.g. different sequencing runs) of blanks/samples, provide column name with grouping of samples (and blanks!). If all belong to the same blank set e.g. all values to a, if to group, set to e.g. "a" and "b" in your provided contextual data. Important: also set the group here for the blank
ColumnWithBlankGroupingSamples <- 'Blank_group-samples' 

# # only needed if using "remove", "subtract-average" or "subtract-max"
ColumnWithBlankGrouping <- 'Blank_group-blanks' # column where the same grouping code as above is assigned to each blank belonging to the respective group (also if only one group!). All samples have to be set to NA in the contextual data
ASVcountInBlanksToExclude = 0 # Number of counts an ASV must occur in blanks (in average per blank). If set to 0, even if the ASV only occurs once in one blank it will be excluded/subtracted. If set to 1 or more, the ASV has to occur in average once in each blank to be excluded/subtracted. 

# # If using package decontam
# # when using decontam you can include the DNA concentrations of samples and blanks.
### https://doi.org/10.1186/s40168-018-0605-2
columnfordecontam <-"decontam-groups" # if using "decontam", define column in contextual data classifying samples with "FALSE" and blanks with "TRUE"
# # for low-biomass samples 
decontam_isnotcontaminant <- "N" # set to Y/N. Can't be used with IncludeDNAconc set to "Y" but works with batches

IncludeDNAconc<-"Y"  # set to Y or N
columnwithDNAconcentrations<-"DNA-Total-DNA-recovered-ng" # column in contextual data with DNA concentration
decontam.method <- "auto" # set to "auto", "frequency", "prevalence", "combined", "minimum", "either","both"

# # In case you want to save the decontaminated seqtab nochim and taxonomy table to continue working with that
SaveTables <- "Y"
```
